# The Megachilidae (Hymenoptera, Apoidea, Apiformes) of the Democratic Republic of Congo curated at the Royal Museum for Central Africa (RMCA, Belgium)

**DOI:** 10.1101/2022.12.17.520875

**Authors:** Alain Tshibungu Nkulu, Alain Pauly, Achik Dorchin, Nicolas J. Vereecken

**Author notes:** Corresponding authors: Alain Tshibungu Nkulu; Nicolas J. Vereecken.

## Abstract

Natural history collections (NHCs) are a cornerstone of entomology, and the conservation of specimens is the essential prerequisite for the development of research into systematics, biogeography, ecology, evolution and other disciplines. Yet, specimens collected during decades of entomological research conducted in less developed countries across Sub-Saharan Africa on pests, beneficial insects and insect biodiversity in general have largely been exported to be permanently preserved in developed countries, primarily in South Africa, Europe and the United States of America.

This is particularly true for the Democratic Republic of the Congo’s (DRC) diverse wild bee fauna, which has been investigated throughout the colonial period by visiting or resident entomologists and missionaries who have then transferred their collected material primarily to Belgium as part of a wider legacy of scientific exploration and colonialism. Digitizing NHC is one way to mitigate this current bias, by making samples accessible to researchers from the target post-colonial countries as well as to the wider international scientific community.

In this study, we compiled and digitized 6,490 specimens records relevant to 195 wild bee species grouped in 18 genera within the biodiverse family Megachilidae, essentially from the colonial era (i.e., mostly between 1905-1960, with additional records up to 1978), and curated at the Royal Museum for Central Africa (RMCA) in Belgium. We provide a detailed catalogue of all records with updated locality and province names, including 26 species only available as type specimens. We also explore the historical patterns of diversity and distribution across DRC, and we provide a list of the research entomologists involved. This study is an important first step that uses digital technologies to democratize and repatriate important aspects of DRC’s natural heritage of insect biodiversity, to stimulate more contemporary field surveys, as well as to identify and characterize research gaps and biodiversity shortfalls in little-explored regions of Sub-Saharan Africa.

## Introduction

The collection and preservation of biological specimens has traditionally been at the core of most research dealing with the systematics, biogeography, ecology and evolution of living organisms, and the establishment of natural history collections (NHCs) has proven an immensely valuable tool to stimulate scientific research long before Darwin and Wallace. Today, important new perspectives are provided for NHCs, as they already confirmed their key role as infrastructures allowing the study of global change, of temporal (i.e., time series) changes in communities, systematic revisions and the more recent development of various “-omics” approaches to the study of biodiversity. By and large, and despite the increasing regulations in place to control specimen collection (Sikes *et al*. 2016), it seems that NHCs still have a bright future (Suarez & Tsustui 2004; Pyke & Ehrlich 2010) and that hitherto unexpected societal challenges related to environmental conservation, to food security, and to human health will find at least part of their answers through the adequate use and valorisation of NHCs (Heberling 2020; Hedrick *et al*. 2020; Lendemer *et al*. 2020; Belitz *et al*. 2022). Nowadays, the availability of research-grade biodiversity data is pressing to better evaluate extinction risks and conservation needs along with the multiple threats on ecosystem services (Rodger *et al*. 2004; Enke *et al*. 2012; Potts *et al*. 2016a, Díaz *et al*. 2019 Ganzevoort *et al*. 2017; Mkenda *et al*. 2019).

Over the past century or more, entomological surveys on pests, beneficial insects and insect biodiversity in general conducted in lower-income but biodiverse countries have yielded countless specimens that have largely been exported outside their native range to be permanently preserved in higher-income countries. Sub-Saharan African specimens have primarily been exported to South Africa, Europe and the United States of America (USA). As a consequence, the long tradition of establishing NHCs has been primarily associated with higher-income countries, leaving lower-income countries lagging behind in terms of infrastructure. Up to the present day, many countries in (sub-)tropical and biodiverse regions of the world still lack even the most basic NHCs, as well as the financial support and logistic means to curate specimens appropriately and sustainably in airtight boxes, in temperature and humidity-controlled spaces, with back-up generators and with no exposure to sunlight. This, in turn, has been a major obstacle to knowledge transfer, local capacity building and the development of scientific research on biodiversity conservation in lower-income countries. Furthermore, the overall lack of digitization efforts targeting specimens collected in lower-income but biodiverse countries is also responsible for the current gaps and spatial patterns in sampling biases (i.e., more data and studies on trends in biodiversity in higher-income vs. lower-income countries), as well as for the major distortion in our understanding of the contemporary state of biodiversity at the worldwide scale (Hughes *et al*. 2021; Raja *et al*. 2022). Addressing such cases of “scientific colonialism” (when the knowledge base relevant to a certain country is by default located or curated outside of that specific country) is increasingly becoming an important societal issue fuelled by shifting social and political priorities, the global economics, and the socio-political context of countries and their colonial history among other key drivers (Zizka *et al*. 2021).

The colonial history of the Democratic Republic of the Congo (DRC) in Central Africa has been described extensively (Freund 1998; Ewans 2001; Van Reybrouck 2015), and the country is also renowned for its unparalleled and threatened biodiversity within the African continent, and within the Sub-Saharan region in particular. A significant part of our contemporary understanding of biodiversity in DRC stems directly from research and surveys made during the colonial era (i.e., until 1960), through fieldwork typically supervised and carried out by visiting or resident, non-indigenous researchers (from Belgium and other countries) enrolling the local indigenous populations on colonized territories then controlled by the Belgian Ministry of Colonies. These historical surveys and more contemporary biodiversity research have highlighted that DRC harbours has the highest number of species for almost all groups of organisms with the exception of plants, whose diversity is at its peak in South Africa. This, along with high numbers of endemic species, make DRC a most important target for biodiversity conservation in Africa. However, biodiversity in DRC and more largely in the Congo Basin region is also severely threatened by agriculture, the collection of fuelwood (Megevand *et al*. 2013; Aquilas *et al*. 2022; Miao *et al*. 2022) and urbanization (Parnell & Walawege 2011). These factors, considered as the major drivers of anthropogenic land use change, resulting in the deforestation and forest degradation of biodiverse habitats, have plagued this region of the world since precolonial times and still persist to the present day (Larcom *et al*. 2016).

Among the biodiverse organisms of conservation interest in DRC are wild bees (Hymenoptera, Apoidea, Apiformes), an important group of pollinators for wildflowers and insect-pollinated crops alike (Klein *et al*. 2007) that significantly contribute to human well-being (Potts *et al*. 2016b) and the generation of income in rural areas (Wratten *et al*. 2012; Sabino *et al*. 2022). The bees of DRC have received attention during the end of the 19th and the first two decades of the 20th century, through the collecting efforts of non-indigenous, resident or non-resident entomologists. This has resulted in several landmark publications (Smith 1853; Schletterer 1891; Friese 1903, 1905, 1909; Vachal 1910; Strand 1921 among others) and the collection of more specimens by missionaries, anonymous, amateur and professional entomologists (the latter originally and primarily working on pest control) during the first half of the 20th century. A famous introductory paper on the bees of DRC was entitled “Collecting bees in the Belgian Congo”, published by Cockerell (1932), a British-born professor of systematic zoology at the University of Colorado who led the “Mission Cockerell” with a team of “Biluluists” (“bilulu” is a term used in DRC, Rwanda and Burundi to designate insects or insect pests; the term is likely derived from “kilulu” in Swahili with the same meaning). Over 16,000 insect specimens were collected during this expedition, which also resulted in a series of landmark papers on the bees of the then-Belgian Congo (Cockerell 1930, 1931, 1935a, 1936a, 1936b among others; see Eardley & Urban’s 2010 “Catalogue of Afrotropical Bees” for more details and references). Collectively, these publications have stimulated the interest of the non-indigenous, resident and non-resident entomologists in DRC, resulting in increasing sampling activity and export of wild bee specimens to the Museum of Belgian Congo in Belgium (now Royal Museum of Central Africa (RMCA)) up until just under two decades after the independence of the Belgian Congo in 1960 (that is, in 1978 for the last records). These collections have also proven useful as they have been used for taxonomic revisions in the second half of the 20th century (Pasteels 1965, 1968, 1970, 1984), but they have never been complied at a higher taxonomic level or digitized to date.

In this paper, we provide a catalogue of all wild bee specimens belonging to the family Megachilidae collected in DRC (with their updated locality, provinces names and taxonomy) and curated at the Royal Museum of Central Africa (RMCA) in Belgium. We focused on the family Megachilidae because it is one of the most species-rich bee families in Sub-Saharan Africa (841 spp. according to Eardley & Urban (2010)), only second to the family Halictidae (915 spp. according to Eardley & Urban (2010)). We also explore and analyze the historical patterns of diversity and distribution of specimen records across DRC, as well as those of the entomologists involved. Collectively, our results will shed new light on the underreported entomlogical biodiversity of DRC and help establish a robust baseline for future studies, and they also contribute to democratize the access to important aspects of DRC’s natural heritage - an essential prerequisite to stimulate more research into the taxonomy, the ecology and the evolution of wild bees in the Afrotropical region.

## Material and methods

### Digitization of RMCA specimen records

The digitization of the wild bee data curated at RMCA was carried out following the organization of the 79 numbered boxes in the collection and the alphabetical sorting of genera, subgenera and species within these boxes. The RMCA Megachilidae collection was identified in the 1960’s and 1970’s by the late Jean J. Pasteels who worked on Sub-Saharan African bees and published several taxonomic revisions. For example, on genera *Creightonella, Chalicodoma, Megachile* (s. str.) (Pasteels 1965, 1966), *Coelioxys* (Pasteels 1968), and Anthidiinae (Pasteels 1984), as classified by him. Another important revision was devoted to the Lithurginae (Eardley 1988), wich has only 4 species in Sub-Saharian Africa and was not published by Pasteels. The corresponding box number of each specimen and species is retained in the catalogue to facilitate their future examination in the RMCA collections. For each specimen record, we retained its taxonomic affiliation (both as it was used by Pasteels and the current taxonomic update as described above), the contemporary name of the collection locality as well as the former name between parathesis and its geographic coordinates (degrees and minutes in WGS84 at 1 km resolution, obtained from GoogleMaps), the date of collection (dd.month.yyyy) and the name of the collector. It should be noted here that the catalog data follows the recommendations of the European Journal of Taxonomy (https://europeanjournaloftaxonomy.eu/index.php/ejt/fairopenscience).

### Preparation of the RMCA Megachilid bees catalogue

All of the above-mentioned digitized specimen records are compiled and detailed in the form of a catalog and we have followed the higher taxonomic updates for each group. Specifically, Tribe Lithurgini was inspired by Eardley (1988), while Megachile sensu lato, Genus Gronoceras, and Genus Noteriades were reorganized as informed by Praz et al. (2008), Gonzalez et al. (2012, 2019), Litman et al. (2012), Litman et al. (2016), Trunz et al. (2016). The remaining groups (i.e., genus Coelioxys, tribe Anthidiini, and tribe Osmiini) were taken up following contemporary work on bees in sub-Saharan Africa by Schulten (1977), Liongo li Enkulu (1988), Eardley et al. (2010), Eardley & Urban (2010), Eardley (2012a, 2012b). Overall, the groups referred to the most recent checklist by Ascher & Pickering (2020) as described above to classify the Megachilidae into the following seven taxonomic groups:

1. Tribe Lithurgini: including the genus *Lithurgus*;
2. Megachile *sensu lato*: comprising “Leafcutter bees” (i.e., subgenera *Megachile* s. str., *Amegachile*, *Anodonteutricharaea*, *Creightonella*, *Digitella*, *Eurymella*, *Eutricharaea*, *Megella*) and “(Resin-)dauber bees” (i.e., subgenera *Callomegachile*, *Carinula*, *Chalicodoma*, *Cesacongoa*, *Maximegachile*, *Neglectella*, *Pseudomegachile* and *Stenomegachile*);
3. Genus *Coelioxys*: including subgenera *Coelioxys* and *Liothyrapis*;
4. Genus *Gronoceras*;
5. Genus *Noteriades*;
6. Tribe Anthidiini: including the genera *Anthidium*, *Pseudoanthidium*, *Eoanthidium*, *Euaspis*, *Immanthidium*, *Pachyanthidium*, *Serapista*, *Trachusa*, and *Afrostelis*;
7. Tribe Osmiini: including the genera *Heriades* and *Heriadopsis*.

### Analysis of spatial and temporal biodiversity patterns

We first mapped the distribution of all Megachilidae specimen and species records using QGIS ver. 3.6 (QGIS Development Team 2020) using a 50 km x 50 km grid. We used three different administrative boundaries, namely (i) the six historical provinces (valid up to the Independence on June 30, 1960), (ii) the 26 present-day provinces, and (iii) the 10 known phytogeographical districts defined within the country (Coastal, Mayumbe, Low-Congo, Kasaï, Low-Katanga, Central Forestry, Ubangi-Uele, Lake Albert, Lakes Edouard and Kivu, Upper-Katanga). This allowed us to plot and discuss the abundance and the diversity of DRC Megachilidae records from the RMCA collection according to different spatial scales and contexts.

To characterize the taxonomic patterns of specimen records and the associated entomological collectors, we then used the *rankabundance* and *rankabunplot* functions in the “BiodiversityR” package (v. 2.14-2; Kindt & Coe 2005). This allowed us to compute a top 10 (or rank abundance) of the most commonly recorded Megachilidae species, as well as to identify the entomologists associated with the highest numbers of specimen records. We also computed a kind of diversity index for each of these 10 major collectors, in order to assess their contribution to the variety of Megachilidae species among all reported specimens. This was done by using the *diversitycomp* function in the “BiodiversityR” package (v. 2.11-3; Kindt 2019) by computing the Simpson, Shannon (alpha diversity) and Pielou (species evenness) diversity indices (Borcard *et al*. 2018). To further explore the database, we then visualized the temporal trends in the collection of various taxonomic groups within the Megachilidae across DRC using specimen accumulation curves and the *specaccum* function in the “vegan” package (v. 2.5-7; Oksanen *et al*. 2020). All these analyses were performed in RStudio (RStudio Team 2020) for R (R Core Team 2020).

We then tested whether there is a significant difference in community composition among (i) the six historical provinces in DRC, (ii) the 26 present-day provinces, and (iii) the 10 known phytogeographical districts defined within the country, using analyses of similarities (ANOSIM) with the vegan package (version 2.0–5) (Oksanen *et al*. 2019). We computed average Bray-Curtis distances among samples and used 10,000 permutations to statistically test the extent to which an increase in the number of administrative or natural units (i.e., 6 vs. 10 vs. 26) was statistically associated with an increase in community differentiation. This analysis allows to discuss the scale of a putative spatial concentration of records around key localities preferentially visited by entomologists to collect Megachilidae species during the targeted period in DRC.

To examine in greater detail, the species and phylogenetic structure of wild bee communities among increasing numbers of administrative units in DRC, we followed the analytical procedure described in Vereecken *et al*. (2021) by first aggregating the dataset to retain a single line of species occurrence records for each region, and then used the “BAT” package (v. 1.6.0.; Cardoso *et al*. 2015) to explore the species and phylogenetic turnover. For the phylogenetic approach, we built up a phylogeny based on the hierarchical Linnaean classification following the higher-rank classification adopted here; the R-package ape (v. 5.0) (Paradis *et al*. 2004) was used to build a polytomous, ultra-metric tree comprising all species, and branch lengths were calculated for the whole “taxonomy-based phylogeny” by setting the p-parameter to 1, following Hoiss *et al*. (2012). For both the species and phylogenetic approaches, we computed the Sørensen index of beta diversity, βsør, a measure of total dissimilarity, and its partitioning into its two major components: (1) species replacement among habitats, i.e., turnover: βsim; and (2) species loss/gain among habitats, i.e., nestedness: βsne; following the formula βsør = βsim + βsne (Baselga 2010, Legendre 2014). The relative values of βsim and βsne allow to quantitatively evaluate the relative importance of species turnover versus nestedness among habitats. This “taxonomy-based phylogeny” was used as an input for analyses of phylogenetic turnover; for both approaches, we investigated the differentiation among (i) the six historical provinces in DRC, (ii) the 26 present-day provinces, and (iii) the 10 known phytogeographical districts defined within the country as described above. Thus, the values of the two components (βsim and βsne) allow to quantify the relative contribution of turnover and nestedness (respectively) to the overall taxonomic and phylogenetic differentiation with an increase in the number of administrative or natural units (i.e., 6 vs. 10 vs. 26).

## Results

### Catalogue

The catalogue presented below includes, for each species, the number of female and male specimens, the collection details, including the locality, conventional, and date, and the name of collectors. The collections of the Megachilidae of the RMCA were all identified by Professor Jean Jules Pasteels, who was the last author to publish inclusive taxonomic revisions for Afrotropical Megachilid bees based on the RMCA collections, until about the 1980s and later reorganized by the museum technician (? De Bakker). For this reason, and to allow easy tracking of the collection holdings, we use as titles taxon names based on the updated taxonomy (in bold font), and also provide below the species-level taxonomy as understood by Pasteels and as it appears in the collection boxes.

#### Sub-family Lithurginae, Tribe Lithurgini, *Lithurgus acanthurus* Vachal, 1910

**Material.** Cupboard 110, Box 62 (1♀ and 1♂). **D.R. CONGO**. Holotype – **Haut-Katanga** • 1 ♂; Kambove-Bunkeya; 10°24’ S, 26°58’ E; Oct.1907; Dr. Sheffield-Neave leg.; RMCA. Allotype • 1 ♀; Lukafu; 10°31’ S, 27°33’ E; 1909; Dr. Sheffield-Neave leg.; RMCA.

#### *Lithurgus pullatus* Vachal, 1903

**Material.** Cupboard 110, Box 62 (2♀♀ and 1♂). **D.R. CONGO**. – **Ituri** • 1 ♂; Mahagi-Niarembe; 02°15’ N, 31°07’ E; 1935; Ch. Scops leg.; RMCA. – **Tanganyika** • 2 ♀♀; Tanganyika: Mpala; 06°45’ S, 29°31’ E, alt. 780 m; Jul. – Aug. 1953; H. Bomans leg.; RMCA.

#### *Lithurgus sparganotes* (Schletterer, 1891)

**Material.** Cupboard 110, Box 62 (130♀♀ and 13♂♂). **D.R. CONGO**. – **Bas-Uele** • 1 ♀; Bambesa; 03°28’ N, 25°43’ E; Dec. 1933; H.J. Brédo leg.; RMCA • 1 ♀; same location as for preceding; 4 Jun. 1937; J. Vrijdagh leg.; RMCA • 1 ♂; same location as for preceding; 4 Jun. 1937; J. Vrijdagh leg.; RMCA • 1 ♀; same location as for preceding; Jun. 1937; J. Vrijdagh leg.; RMCA. – **Equateur** • 1 ♀; Coquilhatville (Mbandaka); 00°03’ N, 18°15’ E; 1927; Dr. Strada leg.; RMCA • 1 ♀; Eala (Mbandaka); 00°03’ N, 18°19’ E; Jun. 1932; A. Corbisier leg.; RMCA • 2 ♀♀; same location as for preceding; Sep. 1930; Dr. Staner leg.; RMCA • 2 ♀♀; same location as for preceding; 15 Aug. 1937; G. Couteaux leg.; RMCA • 1 ♀; same location as for preceding; 16 Sep. 1936; G. Couteaux leg.; RMCA • 1 ♀; same location as for preceding; 16 Sep. 1936; G. Couteaux leg.; RMCA • 2 ♀♀; same location as for preceding; 21 Nov. 1931; H.J. Brédo leg.; RMCA • 5 ♀♀; same location as for preceding; 11 Nov. 1936; J. Ghesquière leg.; RMCA • 5 ♀♀; same location as for preceding; 14 Jul. 1936; J. Ghesquière leg.; RMCA • 1 ♀; same location as for preceding; Jan. 1936; J. Ghesquière leg.; RMCA • 1 ♀; same location as for preceding; Jul. 1936; J. Ghesquière leg.; RMCA • 1 ♂; same location as for preceding; Jul. 1936; J. Ghesquière leg.; RMCA • 1 ♀; same location as for preceding; Aug. 1935; J. Ghesquière leg.; RMCA • 3 ♀♀; same location as for preceding; Aug. 1936; J. Ghesquière leg.; RMCA • 1 ♀; same location as for preceding; Oct. 1935; J. Ghesquière leg.; RMCA • 1 ♀; Equateur; 00°00’ N, 18°14’ E; 15 Nov. 1911; Cap. Van Gile leg.; RMCA • 1 ♀; Flandria; 00°20’ S, 19°06’ E; 1 Sep. 1932; R.P. Hulstaert leg.; RMCA • 1 ♀; same location as for preceding; Sep. 1932; R.P. Hulstaert leg.; RMCA • 1 ♀; Ubangi: Nouvelle Anvers (Bomongo); 00°03’ N, 18°19’ E; 9 Dec. 1952; P. Basilewsky leg.; RMCA. – **Haut-Uele** • 1 ♀; Mayumbe: Ganda Sundi; 05°30’ S, 12°53’ E; 1915; R. Mayné leg.;

RMCA. – **Kinshasa** • 7 ♀♀; Léopoldville (Kinshasa); 04°19’ S, 15°19’ E; 1933; A. Tinant leg.; RMCA • 1 ♂; same location as for preceding; May 1911; Dr. Dubois leg.; RMCA • 1 ♀; same location as for preceding; Dr. Houssiaux; leg.; RMCA. – **Kongo Central** • 1 ♀; Kisantu (Madimba); 05°08’ S, 15°06’ E; 12 Jan. 1924; Dr. M. Bequaert leg.; RMCA • 22 ♀♀; same location as for preceding; 1927; R.P. Vanderyst; leg.; RMCA • 3 ♂♂; same location as for preceding; 1927; R.P. Vanderyst; leg.; RMCA • 8 ♀♀; same location as for preceding; 1931; R.P. Vanderyst; leg.; RMCA • 1 ♀; same location as for preceding; 1932; R.P. Vanderyst; leg.; RMCA • 4 ♀♀; same location as for preceding; 1 Apr. 1930; R.P. Vanderyst; leg.; RMCA • 1 ♂; same location as for preceding; Jan. – Apr. 1930; R.P. Vanderyst; leg.; RMCA • 2 ♀♀; same location as for preceding; Dec. 1927; R.P. Vanderyst; leg.; RMCA • 1 ♀; same location as for preceding; Dec. 1927; R.P. Vanderyst; leg.; RMCA • 1 ♀; Thysville (Mbanza Ngungu); 05°15’ S, 14°52’ E; Jan. 1953; J. Sion leg.; RMCA. – **Kwilu**• 2 ♀♀; Moyen Kwilu: Leverville (Bulungu); 04°50’ S, 18°44’ E; Jan. 1914; P. Vanderijst leg.; RMCA. – **Lomami** • 1 ♀; Lomami: Kabwe; 08°47’ S, 26°52’ E; Jul. – Aug. 1931; P. Quarré leg.; RMCA. – **Lualaba** • 1 ♀; Kafakumba (Sandoa); 09°41’ S, 23°44’ E; Apr. 1933; G.F. Overlaet leg.; RMCA. – **Maniema** • 1 ♂; Kibombo; 03°54’ S, 25°55’ E; 3 Nov. 1910; Dr. J. Bequaert leg.; RMCA • 1 ♂; same location as for preceding; 6 Nov. 1910; Dr. J. Bequaert leg.; RMCA • 1 ♀; same location as for preceding; Sep. – Nov. 1930; H.J. Brédo leg.; RMCA • 1 ♀; Lokandu; 02°31’ S, 25°47’ E; Dec. 1939; Lt Vissers leg.;

RMCA. – **Nord-Kivu** • 1 ♂; Bafula; 04°09’ N, 27°54’ E; Dr. M. Bequaert leg.; RMCA. – **Sud-Ubangi** • 1 ♂; Libenge; 03°39’ N, 18°38’ E;1933; J. Van Gils leg.; RMCA. – **Tshopo** • 2 ♀♀; Miss. St. Gabriel; 00°33’ N, 25°05’ E; M. Torley leg.; RMCA • 1 ♂; Stanleyville (Kisangani); 00°31’ N, 25°11’ E; 10 Feb. 1932; J. Vrijdagh leg.; RMCA. – **Tshuapa** • 3 ♀♀; Bokuma; 00°06’ S, 18°41’ E; 1953; R.P. Lootens leg.; RMCA • 1 ♀; same location as for preceding; Sep. 1952; R.P. Lootens leg.; RMCA • 6 ♀♀; same location as for preceding; Jul. 1952; R.P. Lootens leg.; RMCA • 1 ♀; same location as for preceding; Aug. - Sep. 1951; R.P. Lootens leg.; RMCA • 8 ♀♀; Tshuapa: Mayumbe: Ganda Sundi; 05°30’ S, 12°53’ E; 1953; R.P. Lootens leg.; RMCA • 2 ♀♀; same location as for preceding; 1954; R.P. Lootens leg.; RMCA • 1 ♀; same location as for preceding; Jan. 1954; R.P. Lootens leg.; RMCA • 1 ♀; same location as for preceding; Feb. 1952; R.P. Lootens leg.; RMCA • 2 ♀♀; same location as for preceding; Feb. 1954; R.P. Lootens leg.; RMCA • 6 ♀♀; same location as for preceding; Mar. 1954; R.P. Lootens leg.; RMCA • 1 ♀; same location as for preceding; Jan. – Feb. 1954; R.P. Lootens leg.; RMCA • 1 ♂; same location as for preceding; Jan. – Feb. 1954; R.P. Lootens leg.; RMCA • 1 ♀; same location as for preceding; Apr. 1952; R.P. Lootens leg.; RMCA • 2 ♀♀; same location as for preceding; Apr. 1954; R.P. Lootens leg.; RMCA • 1 ♀; same location as for preceding; Sep. 1952; R.P. Lootens leg.; RMCA • 1 ♀; same location as for preceding; Jun. 1952; R.P. Lootens leg.; RMCA • 2 ♀♀; Tshuapa: Bokuma; 00°06’ S, 18°41’ E; 9 Sep. 1952; R.P. Hulstaert leg.; RMCA.

#### Sub-family Megachiliinae, Tribe Megachilini, Leafcutter bees, *Megachile (Amegachile) acraensis* Friese 1903

**Material.** Cupboard 110, Box 10 (76♀♀ and 41♂♂). **D.R. CONGO**. – **Bas-Uele** • 1 ♂; Bambesa; 03°28’ N, 25°43’ E; 1 Jun. 1934; H.J. Brédo leg.; RMCA • 1 ♀; same location as for preceding; 10 Aug. 1933; J.V. Leroy leg.; RMCA • 1 ♀; same location as for preceding; 14 May 1938; P. Henrard leg.; RMCA • 1 ♀; same location as for preceding; 15 Sep. 1933; H.J. Brédo leg.; RMCA • 1 ♂; same location as for preceding; 30 Oct. 1933; H.J. Brédo leg.; RMCA • 1 ♂; same location as for preceding; Feb. 1934; H.J. Brédo leg.; RMCA • 3 ♀♀; same location as for preceding; Dec. 1933; H.J. Brédo leg.; RMCA • 5 ♀♀; Mobwasa; 02°40’ N, 23°11’ E; Sep. 1911; De Giorgi leg.; RMCA • 1 ♀; Uele; 04°06’ N, 22°23’ E; Degreef leg.; RMCA • 3 ♀♀; Uele: Bambesa; 03°28’ N, 25°43’ E; 10 Oct. 1933; J.V. Leroy leg.; RMCA. – **Equateur** • 1 ♀; Coquilhatville (Mbandaka); 00°03’ N, 18°15’ E; 23 Nov. 1923; Dr. M. Bequaert leg.; RMCA • 4 ♀♀; Eala (Mbandaka); 00°03’ N, 18°19’ E; 11 Nov. 1936; J. Ghesquière leg.; RMCA • 5 ♀♀; same location as for preceding; 14 Jul. 1936; J. Ghesquière leg.; RMCA • 1 ♂; same location as for preceding; 14 Nov. 1931; H.J. Brédo leg.; RMCA • 1 ♂; same location as for preceding; 22 Nov. 1931; H.J. Brédo leg.; RMCA • 1 ♀; same location as for preceding; 4 Nov. 1932; H.J. Brédo leg.; RMCA • 1 ♂; same location as for preceding; 5 Nov. 1931; H.J. Brédo leg.; RMCA • 2 ♀♀; same location as for preceding; Mar. 1932; H.J. Brédo leg.; RMCA • 1 ♂; same location as for preceding; Mar. 1932; H.J. Brédo leg.; RMCA • 1 ♂; same location as for preceding; Mar. 1935; A. Corbisier leg.; RMCA • 1 ♀; same location as for preceding; Apr. 1933; A. Corbisier leg.; RMCA • 2 ♂♂; same location as for preceding; Apr. 1933; A. Corbisier leg.; RMCA • 1 ♂; same location as for preceding; May 1932; H.J. Brédo leg.; RMCA • 1 ♂; same location as for preceding; Jun. 1932; A. Corbisier leg.; RMCA • 1 ♀; same location as for preceding; Nov. 1931; H.J. Brédo leg.; RMCA • 2 ♂♂; same location as for preceding; Nov. 1931; H.J. Brédo leg.; RMCA • 1 ♀; same location as for preceding; Nov. 1932; A. Corbisier leg.; RMCA • 1 ♀; same location as for preceding; Dec. 1932; A. Corbisier leg.;

RMCA. – **Haut-Uele** • 1 ♀; Wamba; 00°20’ S, 29°50’ E; 1932; Putnam leg.; RMCA. – **Ituri** • 1 ♀; Mahagi-Niarembe; 02°15’ N, 31°07’ E; Nov. 1935; M. & Me. Ch. Scops leg.; RMCA • 1 ♀; Mahagi-Port; 02°09’ N, 31°14’ E; Oct. 1934; H.J. Brédo leg.; RMCA • 1 ♀; Irumu-Penge; 01°27’ N, 02°09’ N, 31°14’ E; 1 Mar. 1914; Dr. J. Bequaert leg.; RMCA. – **Kasaï** • 1 ♂; Ilebo; 04°20’ S, 20°36’ E; 14-15 Jul. 1925; S.A.R. Prince Léopold leg.; RMCA • 1 ♂; Kasaï: Lutzkwadi (Ilebo); 04°20’ S, 20°36’ E; 17 Jun. 1946; V. Lagae leg.; RMCA. – **Kasaï Central** • 2 ♀♀; Lula (Kasaï); 07°11’ S, 22°24’ E; 1958; A.J. Jobaert leg.; RMCA • 1 ♀; Luluabourg (Kananga); 05°54’ S, 21°52’ E; 20 Apr.1939; J.J. Deheyn leg.; RMCA • 1 ♂; same location as for preceding; P. Callewaert leg.; RMCA. – **Kongo Central** • 1 ♀; Camp de Lukula; 05°23’ S, 12°57’ E; 1911; Dr. Daniel leg.; RMCA • 4 ♀♀; Kisantu (Madimba) (Madimba); 05°08’ S, 15°06’ E; Rév. P. Regnier leg.; RMCA • 3 ♀♀; Mayidi (Madimba); 05°11’ S, 15°09’ E; 1942; Rév. P. Van Eyen leg.; RMCA. – **Lomami** • 1 ♂; Lomami-Luputa; 11°00’ S, 26°44’ E; Apr. 1934; Dr. Bouvier leg.; RMCA. – **Lualaba** • 1 ♀; Kafakumba (Sandoa); 09°41’ S, 23°44’ E; Apr. 1933; F.G. Overlaet leg.; RMCA • 1 ♂; Kapanga; 08°21’ S, 22°34’ E; Aug. 1934; F.G. Overlaet leg.; RMCA • 1 ♂; same location as for preceding; Dec. 1932; F.G. Overlaet leg.; RMCA • 1 ♀; Lulua: Kapanga; 08°21’ S, 22°34’ E; Sep. 1932; F.G. Overlaet leg.; RMCA • 1 ♀; same location as for preceding; Sep. 1933; F.G. Overlaet leg.; RMCA • 1 ♀; same location as for preceding; Oct. 1932; F.G. Overlaet leg.; RMCA • 1 ♂; same location as for preceding; Oct. 1932; F.G. Overlaet leg.; RMCA • 1 ♂; same location as for preceding; Nov. 1932; F.G. Overlaet leg.; RMCA • 2 ♀♀; same location as for preceding; Nov. 1932; F.G. Overlaet leg.; RMCA • 1 ♀; same location as for preceding; Dec. 1932; F.G. Overlaet leg.;

RMCA. – **Maniema** • 1 ♀; Kasongo; 04°27’ S, 26°40’ E; Apr. 1954; Dr. J. Claessens leg.; RMCA • 2 ♀♀; Kibombo; 03°54’ S, 25°55’ E; Apr. 1954; Dr. M. Bequaert leg.; RMCA. – **Nord-Kivu** • 5 ♀♀; Beni à Lesse; 00°45’ N, 29°46’ E; Jul. 1911; Dr. Murtula leg.; RMCA • 1 ♀; N’Guli (Lubero); 00°10’ S, 29°20’ E; 3 Sep. 1913; Dr. Rodhain leg.; RMCA. – **Nord-Ubangi** • 1 ♀; Abumombazi (Mobayi); 03°34’ N, 22°03’ E; 18 – 26 Feb. 1932; H.J. Brédo leg.; RMCA • 1 ♀; Ubangi: Karawa (Businga); 03°21’ N, 20°18’ E; 1937; Rév. Wallin leg.; RMCA • 1 ♀; Yakoma; 04°06’ N, 22°23’ E; 5 – 17 Feb. 1932; H.J. Brédo leg.; RMCA. – **Sankuru** • 1 ♀; Lusambo: route Batempa; 04°58’ S, 23°26’ E; 17 Jan. 1950; Dr. M. Fontaine leg.; RMCA • 1 ♀; Sankuru: Benadibele; 04°07’ S, 22°50’ E; 1925; R. Mayné leg.; RMCA • 1 ♂; Sankuru: Djeka; 02°57’ S, 23°33’ E; 1955-1956; R. Roiseux leg.; RMCA • 1 ♀; Sankuru: Komi; 03°23’ S, 23°46’ E; Apr. 1930; J. Ghesquière leg.; RMCA 1 ♀; same location as for preceding; Jun. 1930; J. Ghesquière leg.; RMCA • 1 ♂; same location as for preceding; Jun. 1930; J. Ghesquière leg.; RMCA. – **Sud-Kivu** • 1 ♀; Kavumu à Kabunga, 82 km (Mingazi); 02°18’ S, 28°49’ E; May – Jun. 1951; H. Bomans leg.; RMCA • 1 ♀; Kivu; 05°29’ S, 28°51’ E; Sep. – Oct. 1925; S.A.R. Prince Léopold leg.; RMCA. – **Tanganyika** • 1 ♀; Bassin Lukuga; 05°40’ S, 26°55’ E; Apr. – Jul. 1934; H. De Saeger leg.; RMCA • 1 ♂; Kipiri; 07°35’ S, 29°53’ E; Sep. 1912; Miss. Agric. leg.; RMCA • 1 ♂; Kongolo; 05°24’ S, 27°00’ E; 23 Jan. 1911; Dr. J. Bequaert leg.; RMCA • 1 ♂; same location as for preceding; 30 Jan. 1911; Dr. J. Bequaert leg.; RMCA • 1 ♀; Lukulu (Manono); 07°08’ S, 28°06’ E; 1 – 3 May 1931; G.F. de Witte leg.; RMCA • 1 ♂; Lulua: Kalenge; 06°33’ S, 26°35’ E; Feb. 1934; F.G. Overlaet leg.; RMCA. – **Tshopo** • 1 ♂; Basoko; 01°13’ N, 23°36’ E; 6 Jun. 1909; Prince Albert leg.; RMCA • 1 ♂; Ponthierville; (Ubundu); 00°22’ S, 25°29’ E; 22 Oct. 1910; Dr. J. Bequaert leg.;

RMCA • 1 ♂; same location as for preceding; 22 Oct. 1910; Lang & Chapin leg.; RMCA • 1 ♂; Stanleyville (Kisangani); 00°31’ N, 25°11’ E; 8 Apr. 1915; Lang & Chapin leg.; RMCA. – **Tshuapa** • 1 ♀; Boende; 00°13’ S, 20°52’ E; Jan. 1952; Rév. P. Lootens leg.; RMCA • 3 ♂♂; same location as for preceding; Jan. 1952; Rév. P. Lootens leg.; RMCA • 4 ♂♂; Tshuapa: Bokuma; 00°06’ S, 18°41’ E; Mar. 1952; Rév. P. Lootens leg.; RMCA • 1 ♀; Bokuma; 00°06’ S, 18°41’ E; Jul. 1951; Rév. P. Lootens leg.; RMCA • 2 ♀♀; same location as for preceding; 00°06’ S, 18°41’ E; Jul. 1952; Rév. P. Lootens leg.; RMCA • 1 ♂; same location as for preceding; Jul. 1952; Rév. P. Lootens leg.; RMCA • 1 ♂; same location as for preceding; Aug. – Sep. 1951; Rév. P. Lootens leg.; RMCA • 1 ♀; Tshuapa: Ikela; 01°11’ S, 01°11’ S, 1956; Rév. P. Lootens leg.; RMCA • 1 ♂; same location as for preceding; 1956; Rév. P. Lootens leg.; RMCA.

#### *Megachile (Amegachile) bituberculata* Ritsema, 1889

**Material.** Cupboard 110, Box 9 (184♀♀ and 97♂♂). **D.R. CONGO**. – **Bas-Uele** • 2 ♀♀; Bambesa; 03°28’ N, 25°43’ E; 1938; J. Vrydagh leg.; RMCA • 1 ♀; same location as for preceding; 11 May 1938; P. Henrard leg.; RMCA • 5 ♂♂; same location as for preceding; 11 May 1938; P. Henrard leg.; RMCA • 1 ♀; same location as for preceding; 11 May 1939; P. Henrard leg.; RMCA • 10 ♀♀; same location as for preceding; 14 May 1938; P. Henrard leg.; RMCA • 5 ♂♂; same location as for preceding; 14 May 1938; P. Henrard leg.; RMCA • 1 ♀; same location as for preceding; 14 May 1940; P. Henrard leg.; RMCA • 1 ♀; same location as for preceding; 14 May 1941; P. Henrard leg.; RMCA • 4 ♀♀; same location as for preceding; 25 May 1938; P. Henrard leg.; RMCA • 1 ♂; same location as for preceding; 4 Jun. 1937; P. Henrard leg.; RMCA • 9 ♀♀; same location as for preceding; 9 May 1938; P. Henrard leg.; RMCA • 2 ♂♂; same location as for preceding; 9 May 1938; P. Henrard leg.; RMCA • 3 ♀♀; same location as for preceding; May 1938; P. Henrard leg.; RMCA • 1 ♀; same location as for preceding; Jul. 1952; P. Henrard leg.; RMCA • 1 ♂; same location as for preceding; Dec. 1933; P. Henrard leg.; RMCA • 1 ♀; same location as for preceding; 30 Aug. 1938; H.J. Brédo leg.; RMCA • 1 ♀; same location as for preceding; Feb. 1934; H.J. Brédo leg.; RMCA • 1 ♀; Lebo Paigba; 04°27’ N, 23°57’ E; 14 Oct.1913; Dr. Rodhain leg.; RMCA • 1 ♀; Mobwasa; 02°40’ N, 23°11’ E; Sep.1911; De Giorgi leg.; RMCA • 1 ♂; Uele: van Kerkhovenville; 04°06’ N, 22°23’ E; Degreef leg.; RMCA • 1 ♀; Uele: Bambesa; 03°28’ N, 25°43’ E; 10 Oct.1933; P. Henrard leg.; RMCA • 2 ♀♀; Uele: Tukpwo; 04°26’ N, 25°51’ E; Aug.1937; J. Vrydagh leg.; RMCA. – **Equateur** • 2 ♂♂; Beneden-Congo: Tumba; 00°50’ S, 18°00’ E; Sep. 1933; Mevr. Bequaert leg.; RMCA • 1 ♂; Eala (Mbandaka); 00°03’ N, 18°19’ E; 1932; A. Corbisier leg.; RMCA • 1 ♀; same location as for preceding, 22 Apr.1932; H.J. Brédo leg.; RMCA • 2 ♀♀; same location as for preceding, Mar.1932; H.J. Brédo leg.; RMCA • 1 ♂; same location as for preceding; Mar.1932; H.J. Brédo leg.; RMCA • 2 ♀♀; same location as for preceding; Apr.1932; H.J. Brédo leg.; RMCA • 1 ♂; same location as for preceding; Apr.1932; H.J. Brédo leg.; RMCA • 1 ♂; same location as for preceding; Apr.1933; A. Corbisier leg.; RMCA • 1 ♀; same location as for preceding; May1932; H.J. Brédo leg.;

RMCA • 2 ♂♂; same location as for preceding; May1932; H.J. Brédo leg.; RMCA • 1 ♀; same location as for preceding; Jun.1932; A. Corbisier leg.; RMCA • 1 ♀; same location as for preceding; Jun.1932; H.J. Brédo leg.; RMCA • 1 ♂; same location as for preceding; Jun.1932; H.J. Brédo leg.; RMCA • 1 ♂; same location as for preceding; Jul.1932; A. Corbisier leg.; RMCA • 1 ♀; same location as for preceding; Nov.1931; H.J. Brédo leg.; RMCA • 1 ♂; same location as for preceding; Nov.1932; H.J. Brédo leg.; RMCA • 1 ♂; same location as for preceding; 4 Apr.1932; H.J. Brédo leg.; RMCA • 2 ♂♂; same location as for preceding; Apr.1932; H.J. Brédo leg.; RMCA • 1 ♂; same location as for preceding; Nov.1932; A. Corbisier leg.; RMCA • 1 ♀; Ubangi: Yakoma; 04°06’ N, 22°23’ E; 12 Nov. 1932; H.J. Brédo leg.; RMCA. – **Haut-Katanga** • 1 ♂; Elisabethville (Lubumbashi); 11°40’ S, 27°28’ E; 1933; De Loose leg.; RMCA • 2 ♂♂; same location as for preceding; 18 Apr. 1933; De Loose leg.; RMCA • 1 ♀; same location as for preceding; 29 Jan. 1924; Ch. Seydel leg.; RMCA • 1 ♀; same location as for preceding; 30 Jan. 1921; Dr. M. Bequaert leg.; RMCA • 1 ♀; same location as for preceding; 6 Jun. 1920; Dr. M. Bequaert leg.; RMCA • 1 ♂; same location as for preceding; 7 Jan. 1924; Ch. Seydel leg.; RMCA • 1 ♂; same location as for preceding; 7 Sep. 1932; De Loose leg.; RMCA • 1 ♀; same location as for preceding; Mar. 1926; Ch. Seydel leg.; RMCA • 1 ♂; same location as for preceding; De Loose leg.; RMCA • 1 ♀; same location as for preceding; Jun. 1932; De Loose leg.; RMCA • 1 ♀; same location as for preceding; 1925; Dr. M. Bequaert leg.; RMCA • 1 ♂; same location as for preceding; 18 Apr. 1933; Dr. M. Bequaert leg.; RMCA • 1 ♂; same location as for preceding; 2 Apr. 1923; Dr. M. Bequaert leg.; RMCA • 1 ♀; same location as for preceding; Aug. 1932; Dr. M. Bequaert leg.; RMCA • 1 ♀; Elisabethville (Kimilolo); 11°40’ S, 27°28’ E; 2 Apr. 1923; Dr. M. Bequaert leg.; RMCA • 1 ♀; Elisabethville; riv. Kimilolo; 11°40’ S, 27°28’ E; 2 Apr. 1923; Dr. M. Bequaert leg.; RMCA • 1 ♀; Kambove-Bunkeya; 10°52’ S, 26°38’ E; Sep. 1907; Dr. Sheffield Neave leg.; RMCA • 1 ♀; Kampemba (Elisabethville); 11°40’ S, 27°28’ E; 16 Jun. 1953; Ch. Seydel leg.; RMCA • 1 ♀; Katanga; 11°40’ S, 27°28’ E; 1915; A. Smaelen leg.; RMCA • 1 ♂; Katumba; 08°32’ S, 25°59’ E; Oct. 1907; Dr. Sheffield Neave leg.; RMCA • 1 ♀; Lubumbashi; 11°40’ S, 27°28’ E; 10 May 1953; Ch. Seydel leg.; RMCA • 1 ♂; same location as for preceding; 9 Jan. 1921; Dr. M. Bequaert leg.; RMCA • 1 ♀; same location as for preceding; May 1953; Ch. Seydel leg.; RMCA • 1 ♂; same location as for preceding; May 1953; Ch. Seydel leg.; RMCA • 1 ♂; Mbiliwa-Wantu; 11°40’ S, 27°28’ E; Oct. 1907; Dr. Sheffield Neave leg.; RMCA • 1 ♀; Muelushi; 08°59’ S, 26°45’ E; 1931; H.J. Brédo leg.; RMCA • 1 ♀; Mufunga; 09°21’ S, 27°27’ E; 20 Oct. 1911; Dr. J. Bequaert leg.; RMCA • 2 ♀♀; PNU Kabwe sur Muye; 08°47’ S, 26°52’ E, alt. 1320 m; 11 May 1948; Mis. G.F. de Witte leg.; RMCA • 1 ♀; same location as for preceding; 30 Apr.-10. May 1948; Mis. G.F. de Witte leg.; RMCA • 1 ♀; PNU Kilwezi; 09°05’ S, 26°45’ E, alt. 750 m, 6 – 7 Sep.1948; Mis. G.F. de Witte leg.;

RMCA • 1 ♀; PNU Lusinga, Rivière Kamitungulu; 08°55’ S, 27°12’ E; 13 Jun.1943; Mis. G.F. de Witte leg.; RMCA • 1 ♂; same location as for preceding; 3 Jun.1945; Mis. G.F. de Witte leg.; RMCA • 1 ♀; PNU Masombwe s. Grde; Kafwe; 09°05’ S, 27°12’ E, alt. 1120 m; 16 Apr. 1948; Mis. G.F. de Witte leg.; RMCA • 1 ♂; same location as for preceding; 16. Apr. 1948; Mis. G.F. de Witte leg.; RMCA. – **Haut-Lomami** • 1 ♂; Bukama; 09°12’ S, 25°51’ E; 9 Mar. 1911; Dr. M. Bequaert leg.; RMCA • 1 ♂; Kadiamapanga; 08°16’ S, 26°36’ E; 30 Sep. 1952; H.J. Brédo leg.; RMCA • 1 ♂; Lualaba: Kabongo; 07°20’ S, 25°35’ E; 30 Sep. 1952; Ch. Seydel leg.; RMCA • 2 ♀♀; Luashi; 07°31’ S, 24°10’ E; 1935; Freyne leg.; RMCA • 2 ♀♀; PNU Gorge de la Petenge; 08°56’ S, 27°12’ E, alt. 1150 m, 28 May 1947; Mis. G.F. de Witte leg.; RMCA • 2 ♀♀; PNU Kanonga; 09°16’ S, 26°08’ E, alt. 675 m,14-23 Feb. 1949; Mis. G.F. de Witte leg.; RMCA • 1 ♀; PNU Kaswabilenga; 08°51’ S, 26°43’ E, alt. 700 m; 15 Sep. – 6 Nov. 1947; Mis. G.F. de Witte leg.; RMCA • 1 ♂; same location as for preceding; 15 Sep. – 6 Nov. 1947; Mis. G.F. de Witte leg.; RMCA • 1 ♀; PNU Mabwe; 08°47’ S, 26°52’ E, alt. 585 m; 1-11 Jan. 1949; Mis. G.F. de Witte leg.; RMCA. – **Haut-Uele** • 1 ♂; Faradje; 03°44’ N, 29°42’ E; 1909 – 1915; Dr. J. Bequaert leg.; RMCA • 1 ♀; Haut-Uele: Moto; 03°03’ N, 29°28’ E; 1920; L. Burgeon leg.; RMCA • 2 ♀♀; same location as for preceding; 1923; L. Burgeon leg.; RMCA • 1 ♀; Mayumbe: Buende-Irundi; 04°43’ S, 13°00’ E; 15 May 1926; A. Collart leg.; RMCA • 2 ♀♀; Mayumbe: Zobe; 05°08’ S, 12°29’ E; Jan. 1916; R. Mayné leg.; RMCA • 1 ♂; PNG [Parc National de Garamba]; 03°40’ N, 29°00’ E; 19 May 1950; H. De Saeger leg.; RMCA • 1 ♀; same location as for preceding; 22 Jun. 1950; G. Demoulin leg.; RMCA • 1 ♀; same location as for preceding; 5 Oct. 1950; G. Demoulin leg.; RMCA • 1 ♂; Uele: Dungu; 03°37’ N, 28°34’ E; Degreef leg.; RMCA. – **Ituri** • 1 ♀; Ituri: Blukwa; 01°45’ N, 30°36’ E; 22 Nov. 1928; A. Collart leg.; RMCA • 1 ♀; Ituri: Bunia; 01°34’ N, 30°15’ E; 28 Feb. 1934; J.V. Leroy leg.; RMCA • 1 ♀; Ituri: Kilo (Djugu); 01°50’ S, 30°09’ E; Jul. 1914; Dr. J. Bequaert leg.; RMCA • 1 ♀; Ituri: Mahagi; 02°18’ N, 30°59’ E; 15 May 1925; Dr. H. Schouteden leg.; RMCA • 1 ♂; Mahagi-Port; 02°09’ N, 31°14’ E; 1935; Ch. Scops leg.; RMCA • 1 ♀; same location as for preceding; Sep. 1934; H.J. Brédo leg.; RMCA • 1 ♀; Mongbwalu (Kilo); 01°50’ N, 30°09’ E; 1938; Mme. Scheitz leg.; RMCA • 1 ♀; same location as for preceding; Jul. 1938; Mme. Scheitz leg.; RMCA. – **Kasaï** • 1 ♀; Kasaï: Tshikapa; 06°24’ S, 20°48’ E; 16 Mar. 1939; J.J. Deheyn leg.; RMCA. – **Kasaï Central** • 1 ♀; Limbala; 06°07’ S, 22°20’ E; 3 – 8 Aug. 1913; Dr. Rodhain leg.; RMCA • 1 ♀; Lula (Kasaï); 07°11’ S, 22°24’ E; 1958; A.J. Jobaert leg.;

RMCA. – **Kinshasa** • 1 ♂; Léopoldville (Kinshasa); 04°19’ S, 15°19’ E; 1933; A. Tinant leg.; RMCA. – **Kongo Central** • 1 ♂; Congo da Lemba (Songololo); 05°42’ S, 13°42’ E; Jun. 1911; R. Mayné leg.; RMCA • 2 ♀♀; Kisantu (Madimba); 05°08’ S, 15°06’ E; 1931; R.P. Vanderyst leg.; RMCA • 1 ♀; same location as for preceding; 1931; R.P. Vanderyst leg.; RMCA • 2 ♂♂; same location as for preceding; 1932; R.P. Vanderyst leg.; RMCA • 3 ♀♀; same location as for preceding; Rév. P. Regnier leg.; RMCA • 1 ♂; same location as for preceding; Jun. 1921; P. Van Wing leg.; RMCA • 3 ♂♂; same location as for preceding; Dec. 1927; R.P. Vanderyst leg.; RMCA • 1 ♂; Lemfu (Madimba); 05°18’ S, 15°13’ E; Jan. 1945; Rév. P. De Beir leg.; RMCA • 4 ♀♀; same location as for preceding; 1942; Rév. P. Van Eyen leg.; RMCA • 2 ♂♂; same location as for preceding; Jan. 1953; J. Sion leg.; RMCA. – **Kwango** • 1 ♀; Kasaï: Samsonge; 06°07’ S, 27°28’ E; 18 Mar. 1939; Mevr. Bequaert leg.; RMCA. – **Kwilu** • 1 ♀; Kolo-Kwilu-Madiata; 05°27’ S, 14°52’ E; Sep. 1913; R. Verschueren leg.; RMCA • 1 ♀; Leverville (Bulungu); 04°50’ S, 18°44’ E; 1927; Mme. J. Tinant leg.; RMCA. – **Lomami** • 3 ♀♀; Kabinda; 06°08’ S, 24°29’ E; Dr. Schwetz leg.; RMCA • 2 ♀♀; Lomami: Kabwe; 08°47’ S, 26°52’ E; Jul.-Aug. 1931; P. Quarré leg.; RMCA • 1 ♂; same location as for preceding; Jul. – Aug. 1931; P. Quarré leg.; RMCA • 1 ♂; Lomami-Luputa; 11°00’ S, 26°44’ E; Apr. 1934; Dr. Bouvier leg.; RMCA • 1 ♂; Sankuru: M’Pemba Zeo (Gandajika); 06°49’ S, 23°58’ E; 19 Oct. 1958; R. Maréchal leg.; RMCA. – **Lualaba** • 2 ♂♂; Biano; 10°20’ S, 27°00’ E; Oct. 1931; Ch. Seydel leg.; RMCA • 1 ♂; Bunkeya; 10°24’ S, 26°58’ E; Oct. 1907; Dr. Sheffield Neave leg.; RMCA • 1 ♀; Dilolo; 10°41’ S, 22°21’ E; Sep. – Oct. 1933; H. De Saeger leg.; RMCA • 1 ♀; same location as for preceding; Apr. 1933; F.G. Overlaet leg.; RMCA • 1 ♀; Kalule Nord; 09°45’ S, 25°55’ E; 1934; Ch. Seydel leg.; RMCA • 1 ♂; Kanzenze; 10°31’ S, 25°12’ E; Aug. 1931; G.F. de Witte leg.; RMCA • 1 ♂; Kapanga; 08°21’ S, 08°21’ S, Mar. 1933; F.G. Overlaet leg.; RMCA • 1 ♂; Kayambo-Dikuluwe; 10°45’ S, 25°22’ E; Jun. 1907; Dr. Sheffield Neave leg.; RMCA ; • 1 ♂; Lubudi; 09°55’ S, 25°58’ E; 29 Jan. 1924; Ch. Seydel leg.; RMCA • 1 ♀; same location as for preceding; Mar. 1933; F.G. Overlaet leg.; RMCA • 1 ♂; Lulua: Kapanga; 08°21’ S, 22°34’ E; 1 Oct. 1932; F.G. Overlaet leg.; RMCA • 2 ♀♀; same location as for preceding; Mar. 1933; F.G. Overlaet leg.; RMCA • 2 ♀♀; same location as for preceding; Apr. 1933; F.G. Overlaet leg.; RMCA • 1 ♀; same location as for preceding; Sep. 1932; F.G. Overlaet leg.; RMCA • 1 ♂; same location as for preceding; Sep. 1932; F.G. Overlaet leg.; RMCA • 1 ♀; same location as for preceding; May 1933; F.G. Overlaet leg.; RMCA • 1 ♂; same location as for preceding; May 1933; F.G. Overlaet leg.; RMCA • 4 ♀♀; same location as for preceding; Oct. 1932; F.G. Overlaet leg.; RMCA • 4 ♂♂; same location as for preceding; Oct. 1932; F.G. Overlaet leg.; RMCA • 2 ♀♀; same location as for preceding; Dec. 1932; F.G. Overlaet leg.; RMCA • 1 ♂; same location as for preceding; Dec. 1933; F.G. Overlaet leg.; RMCA • 1 ♀; Sandoa; 09°41’ S, 22°53’ E; 28 Nov. 1918; F.G. Overlaet leg.;

RMCA. – **Maï-Ndombe** • 1 ♀; Kwamouth (Mushie); 03°11’ S, 16°12’ E; 20 Jun. 1913; Dr. J. Maes leg.; RMCA. – **Maniema** • 1 ♀; Kibombo; 03°54’ S, 25°55’ E; Sep. – Oct. 1930; H.J. Brédo leg.; RMCA • 1 ♀; same location as for preceding; Sep. – Nov.1930; H.J. Brédo leg.; RMCA • 1 ♀; Maniema; 05°00’ S, 29°00’ E; R. Mayné leg.; RMCA • 1 ♀; Terr. de Kasongo, Lupaya; 04°23’ S, 26°46’ E; 8 Feb. 1960; P.L.G. Benoit leg.; RMCA. – **Mongala** • 1 ♂; Binga (Lisala); 02°28’ N, 20°31’ E; 1 Mar. 1932; H.J. Brédo leg.; RMCA • 1 ♀; Itimbiri; 02°03’ N, 22°45’ E; 23 May 1918; Dr. Rodhain leg.; RMCA • 2 ♂♂; same location as for preceding; 23 May 1918; Dr. Rodhain leg.; RMCA • 1 ♂; same location as for preceding; 28 May 1915; Dr. Rodhain leg.; RMCA • 1 ♂; Ubangi: Binga (Lisala); 02°28’ N, 20°31’ E; 5 – 12 Mar. 1932; H.J. Brédo leg.; RMCA. – **Nord-Kivu** • 1 ♀; Beni; 00°29’ N, 29°28’ E, alt. 1150 m; Jul. 1911; Mme. L. Lebrun leg.; RMCA •9 ♀♀; Beni à Lesse; 00°45’ N, 29°46’ E, Jul. 1911; Dr. Murtula leg.; RMCA • 1 ♀; N. Kivu: Kisenyi; 01°41’ S, 29°14’ E; Feb. 1928; Ch. Seydel leg.; RMCA • 1 ♂; PNA: Ngoma (lac Biuniu); 01°41’ S, 29°14’ E; 3 – 20 Apr. 1935; Dr. H. Damas leg.; RMCA. – **Nord-Ubangi** • 1 ♂; Abumombazi (Mobayi); 03°34’ N, 22°03’ E; 18 – 26 Feb. 1932; H.J. Brédo leg.; RMCA • 1 ♀; Banzyville (Mobayi); 04°18’ N, 21°11’ E; Royaux leg.; RMCA • 4 ♀♀; Ubangi: Bosobolo; 04°11’ N, 19°53’ E; 8-11 Jan. 1932; H.J. Brédo leg.; RMCA • 1 ♂; Ubangi: Karawa (Businga); 03°21’ N, 20°18’ E; 1936; Rév. Wallin leg.; RMCA • 1 ♀; Yakoma; 04°06’ N, 22°23’ E; 5 – 17 Feb. 1932; H.J. Brédo leg.; RMCA. – **Sud-Kivu** • 1 ♀; Ibanda; 02°29’ S, 28°51’ E; 1952; M. Vandelannoite leg.; RMCA • 1 ♀; Katana (Kabare); 02°14’ S, 28°49’ E; Oct. 1932; P. Van der Houd leg.; RMCA • 1 ♀; Mulungu; 02°20’ S, 28°47’ E; 19 Sep. 1938; Hendrickx leg.; RMCA • 1 ♀; Uvira: riv. Kalimabenge; 03°25’ S, 29°08’ E, alt. 780 m; 23 Mar. 1949; G. Marlier leg.;

RMCA. – **Sud-Ubangi**• 1 ♀; Kunungu; 02°45’ N, 18°26’ E; 1932; Dr. H. Schouteden leg.; RMCA • 3 ♀♀; M’Paka, Terr. Libenge; 03°39’ N, 18°38’ E; Jul. – Aug.1959; M. Pecheur leg.; RMCA • 2 ♀♀; same location as for preceding; Dec 1959; M. Pecheur leg.; RMCA • 2 ♀♀; Ubangi: Boma-Motenge; 03°15’ N, 18°39’ E; Dec. 1931; H.J. Brédo leg.; RMCA • 1 ♀; Ubangi: Motenge-Boma; 03°15’ N, 18°39’ E; 14 Dec. 1931; H.J. Brédo leg.; RMCA • 2 ♀♀; same location as for preceding; 22 Dec. 1931; H.J. Brédo leg.; RMCA • 2 ♂♂; same location as for preceding; Dec. 1931; H.J. Brédo leg.; RMCA • 1 ♂; Ubangi: Nzali; 03°15’ N, 19°47’ E; 3 – 4 Feb. 1932; H.J. Brédo leg.; RMCA. – **Tanganyika** • 1 ♀; Bassin Lukuga; 05°40’ S, 26°55’ E; Apr. – Jul. 1934; H. De Saeger leg.; RMCA • 1 ♂; Kabalo; 06°03’ S, 26°55’ E; Jan. 1929; Ch. Seydel leg.; RMCA • 1 ♀; Katompe (Kabalo); 06°11’ S, 26°20’ E; Feb. 1929; Ch. Seydel leg.; RMCA • 1 ♀; Kongolo; 05°24’ S, 27°00’ E; 12 Feb. 1911; Dr. J. Bequaert leg.; RMCA • 1 ♀; same location as for preceding; 22 Feb. 1911; Dr. J. Bequaert leg.; RMCA • 2 ♂♂; same location as for preceding; 23 Jan. 1911; Dr. J. Bequaert leg.; RMCA • 1 ♂; same location as for preceding; 23 Feb. 1911; Dr. J. Bequaert leg.; RMCA • 1 ♀; Lukuga r. Niemba; 05°57’ S, 28°26’ E; Dec. 1917 – Jan. 1918; Dr. Pons leg.; RMCA • 1 ♀; Lulua: Kalenge; 06°33’ S, 26°35’ E; Dec. 1934; F.G. Overlaet leg.; RMCA • 1 ♀; Vallée Lukuga; 05°40’ S, 26°55’ E; Nov. 1911; Dr. Schwetz leg.; RMCA. – **Tshopo** • 1 ♀; Miss. St. Gabriel; 00°33’ N, 25°05’ E; M. Torley leg.; RMCA • 1 ♀; Stanleyville (Kisangani); 00°31’ N, 25°11’ E; 29 Apr. 1915; Lang & Chapin leg.; RMCA. – **Tshuapa** • 1 ♀; Bokuma; 00°06’ S, 18°41’ E; 00°06’ S, 18°41’ E; Jan. – Feb. 1952; Rév. P. Lootens leg.; RMCA • 3 ♀♀; same location as for preceding; 00°06’ S, 18°41’ E; Jul. 1952; Rév. P. Lootens leg.; RMCA • 1 ♀; Tshuapa: Bokuma; 00°06’ S, 18°41’ E; Mar. 1952; Rév. P. Lootens leg.; RMCA • 2 ♀♀; same location as for preceding; Jul. 1952; Rév. P. Lootens leg.; RMCA.

#### *Megachile (Amegachile) fimbriata* Smith, 1853

**Material.** Cupboard 110, Box 11 (4♀♀ and 21♂♂). **D.R. CONGO**. – **Haut-Katanga** • 1 ♂; Kambove; 10°52’ S, 26°38’ E; Nov. 1907; Dr. Sheffield Neave leg.; RMCA • 3 ♂♂; Mbiliwa-Wantu; 11°40’ S, 27°28’ E; Oct. 1907; Dr. Sheffield Neave leg.; RMCA • 1 ♂; PNU Kiamakoto entre Masombwe et Mukana; 09°09’ S, 27°10’ E; 2 Sep. 1948; Mis. G.F. de Witte leg.; RMCA • 1 ♂; PNU Kiamakoto-Kiwakishi; 09°09’ S, 27°10’ E, alt. 1070 m; 4 – 16 Oct. 1948; Mis. G.F. de Witte leg.; RMCA • 2 ♂♂; PNU Masombwe s. Grde; Kafwe; 09°05’ S, 27°12’ E, alt. 1120 m; 16 Apr. 1948; Mis. G.F. de Witte leg.; RMCA. – **Haut-Lomami** • 1 ♂; Bukama; 09°12’ S, 25°51’ E; Jun. 1911; Dr. J. Bequaert leg.; RMCA • 1 ♂; same location as for preceding; Aug. 1923; Ch. Seydel leg.; RMCA • 1 ♂; Kalongwe; 11°01’ S, 25°13’ E; 19 Sep. 1911; Dr. J. Bequaert leg.; RMCA • 1 ♂; Sankisia; 09°24’ S, 25°48’ E; 28 Sep. 1911; Dr. J. Bequaert leg.; RMCA • 1 ♂; same location as for preceding; 4 Sep. 1911; Dr. J. Bequaert leg.; RMCA – **Ituri** • 2 ♂♂; Kibimbi; 01°22’ N, 28°55’ E; 2 Feb. 1911; Dr. J. Bequaert leg.; RMCA. – **Lualaba** • 1 ♂; Sandu; 09°41’ S, 22°53’ E; Apr. 1931; H.J. Brédo leg.; RMCA. – **Tanganyika** • 1 ♀; Kiambi (Manono); 07°19’ S, 28°01’ E; 22 – 24 Apr. 1931; G.F. de Witte leg.; RMCA^1^ • 1 ♂; Kongolo; 05°24’ S, 27°00’ E; Jan. 1911; Dr. J. Bequaert leg.; RMCA • 2 ♂♂; Lukulu (Manono); 07°08’ S, 28°06’ E; 1 – 3 May 1931; G.F. de Witte leg.; RMCA • 1 ♂; Tanganyika-Kabalo; 06°03’ S, 26°55’ E; 1 Jul. 1947; Dr. M. Poll leg.; RMCA • 3 ♀♀; same location as for preceding; 1 Feb. 1934; H. De Saeger leg.; RMCA • 1 ♂; same location as for preceding; 1 Feb. 1934; H. De Saeger leg.; RMCA.

#### *Megachile (Amegachile) geoffreyi* Cockerell, 1919

**Material.** Cupboard 110, Box 11 (1 ♂). **D.R. CONGO**. – **Haut-Uele** • 1 ♂; PNG, Akam; 03°40’ N, 29°00’ E; 21 Apr. 1950; H. De Saeger leg.; RMCA.

#### *Megachile (Amegachile) nasalis* Smith, 1879

**Material.** Cupboard 110, Box 11 (1 ♀ and 8 ♂♂). **D.R. CONGO**. – **Haut-Katanga** • 1 ♂; Elisabethville (Lubumbashi); 11°40’ S, 27°28’ E; Oct. 1929; Dr. M. Bequaert leg.; RMCA • 2 ♂♂; Mbiliwa-Wantu; 11°40’ S, 27°28’ E; Oct. 1907; Dr. Sheffield Neave leg.; RMCA • 1 ♂; Panda (Likasi); 11°00’ S, 26°44’ E; 16 Oct. 1920; Dr. M. Bequaert leg.; RMCA. – **Haut-Lomami** • 1 ♂; PNU Kaswabilenga; 08°51’ S, 26°43’E, alt. 700 m; 1 – 4 Nov. 1947; Mis. G.F. de Witte leg.; RMCA • 1 ♂; Sankisia; 09°24’ S, 25°48’ E; Sep. 1911; Dr. J. Bequaert leg.; RMCA. – **Lualaba** • 1 ♀; Bunkeya; 10°24’ S, 26°58’ E; Oct. 1907; Dr. Sheffield Neave leg.; RMCA • 1 ♂; Bunkeya-Lukafu; 10°24’ S, 26°58’ E; Oct. 1907; Dr. Sheffield Neave leg.; RMCA. – **Tanganyika** • 1 ♂; Kanikiri; 00°09’ S, 29°32’ E; Oct. 1907; Dr. Sheffield Neave leg.; RMCA.

#### *Megachile* (*Anodonteutricharaea*) *abongana* Strand, 1911, *Megachile* (*Paracella*) *abongana* Strand, 1911

**Material.** Cupboard 110, Box 16 (5♀♀). **D.R. CONGO**. – **Bas-Uele** • 1 ♀; Uele: Bambesa; 03°28’ N, 25°43’ E; 20 Oct.1933; J. V. Leroy leg.; RMCA. – **Kasaï**, • 1 ♀; Kamayembi (Luebo); 05°20’ S, 21°24’ E; 19 Sep.1921; Dr. J. Schouteden leg.; RMCA. – **Tshopo** • 1 ♀; Aruwimi: Panga; 01°51’ N, 26°25’ E; 19 Sep.1921; Dr. J. Schouteden leg.; RMCA. – **Tshuapa** • 2 ♀♀; Moma; 00°45’ S, 21°56’ E; Jun.1925; Lt. J. Ghesquière leg.; RMCA.

#### *Megachile (Anodonteutricharaea) apiformis* Smith, 1953, *Megachile* (*Paracella*) *apiformis* Smith, 1953

**Material.** Cupboard 110, Box 17 (7♀♀ and 3♂♂). **D.R. CONGO**. – **Nord-Kivu** • 1 ♀; Beni à Lesse; 00°45’ N, 29°46’ E; Jul.1911; Dr. Martula leg.; RMCA ; • 1 ♂; same location as for preceding; Jul. 1911; Dr. Martula leg.; RMCA ; • 1 ♀; Butembo : Tuwali; 00°09’ N, 29°17’ E; Jan-Apr. 1954, C. Léontovich leg.; RMCA ; • 1 ♀; Lac Kivu : Rwankwi; 01°20’ S, 29°22’ E; May 1948; J.V. Leroy leg.; RMCA ; • 1 ♂; Rutshuru; 01°11’ S, 29°27’ E; Apr. 1937; J. Ghesquière leg.; RMCA • 1 ♀; Mt. Ruwenzori; 00°26’ S, 29°50’ E; alt. 1400 m, 5. Jun.1914; Dr. J. Bequaert leg.; RMCA • 1 ♂; Terr. Lubero, Mulo; 00°10’ S, 29°14’ E; 1960, June-Jul. 1953, R.P.M. J. Célis leg.; RMCA • 1 ♀; Tshibinda; 02°19’ S, 28°45’ E; 21-27. Aug. 1931; T.D. Cockerell leg.; RMCA – **Haut-Katanga** • 1 ♀; Kambove-Bunkeya; 10°24’ S, 26°58’ E; Oct. 1907; Dr. Sheffield-Neave leg.; RMCA • 1 ♀ Kiamanwa; 09°30’ S, 27°57’ E; Feb. 1931; H.J. Brédo leg.; RMCA.

#### *Megachile* (*Anodonteutricharaea*) *bangana* Cockerell, 1920, *Megachile* (*Paracella*) *bangana* Cockerell, 1920

**Material.** Cupboard 110, Box 16 (3♀♀). **D.R. CONGO**. – **Bas-Uele** • 1 ♀; Bambesa; 03°28’ N, 25°43’ E; Feb. 1934; H.J. Brédo leg.; RMCA. – **Tshuapa** • 2 ♀♀; Bokuma; 00°06’ S, 18°41’ E; Jul. 1952, R.P. Lootens leg.; RMCA.

#### *Megachile (Anodonteutricharaea*) *chrysopogon* Vachal, 1910, *Megachile* (*Paracella*) *chrysopogon* Vachal, 1910

**Material.** Cupboard 110, Box 15 (1♂). **D.R. CONGO**. Holotype – **Haut-Katanga** • 1 ♂; Kiamokosa; 11°40’ S, 27°28’ E; Oct. 1907; Dr. Sheffield Neave leg.; RMCA.

#### *Megachille (Anodonteutricharaea*) *curtula* Gerstaecker, 1857, *Megachille* (*Paracella*) *curtula* Gerstaecker, 1857

**Material.** Cupboard 110, Box 16 (14♀♀ and 25♂♂). **D.R. CONGO**. – **Equateur** • 1 ♀; Coquilhatville (Mbandaka); 00°03’ N, 18°15’ E; 1927; Dr. Strada leg.; RMCA • 1 ♂; Eala (Mbandaka); 00°03’ N, 18°19’ E; 1933, A. Corbisier leg.; RMCA • 2 ♂♂; same location as for preceding; Mar. 1935, A. Corbisier leg.; RMCA • 1 ♂; same location as for preceding; Apr. 1933, A. Corbisier leg.; RMCA • 1 ♀; same location as for preceding; May 1933, A. Corbisier leg.; RMCA • 1 ♀; same location as for preceding; Jun. 1933, A. Corbisier leg.; RMCA • 1 ♂; same location as for preceding; Dec. 1932, A. Corbisier leg.; RMCA • 1 ♂; same location as for preceding; 17 Nov. 1931; H.J. Brédo leg.; RMCA • 1 ♀; same location as for preceding; 3 Oct. 1931; H.J. Brédo leg.; RMCA • 1 ♂; same location as for preceding; 3 Oct. 1931; H.J. Brédo leg.; RMCA • 1 ♂; same location as for preceding; 4 Nov. 1932; H.J. Brédo leg.; RMCA • 1 ♂; same location as for preceding; Mar. 1932; H.J. Brédo leg.; RMCA • 1 ♀; same location as for preceding; Sep. 1935; J. Ghesquière leg.; RMCA • 1 ♀; same location as for preceding; Oct. 1935; J. Ghesquière leg.; RMCA • 1 ♀; same location as for preceding; Nov. 1935; J. Ghesquière leg.; RMCA. – **Haut-Katanga** • 1 ♂; Elisabethville (Lubumbashi); 11°40’ S, 27°28’ E; 1928-1933, De Loose leg.; RMCA • 1 ♂; same location as for preceding; Feb. 1912; Dr. J. Bequaert leg.; RMCA • 1 ♀; same location as for preceding; Feb. 1935; Dr. M. Bequaert leg.; RMCA • 1 ♂; Lubumbashi; 11°40’ S, 27°28’ E; 30 Jan. 1915; Dr. J. Bequaert leg.; RMCA • 1 ♂; PNU Kolwezi (Affl. Dr. Lufira); 09°05’ S, 26°45’ E, alt. 750 m; 9-14 Aug. 1948; G.F. de Witte leg.; RMCA – **Haut-Lomami** • 2 ♀♀; PNU Munoi, Bifure Lupiala; 08°50’ S, 26°44’ E, alt. 890 m; 28 May-15 Jun. 1948; G.F. de Witte leg.; RMCA. – **Haut-Uele** • 1 ♂; Haut-Uele: Mauda; 04°46’ N, 27°18’ E; Mar. 1935; Dr. H. Schouteden leg.; RMCA • 1 ♂; Haut-Uele: Moto; 03°03’ N, 29°28’ E; Oct.-Nov. 1923; L. Burgeon leg.; RMCA. – **Ituri** • 1 ♀; Mahagi-Port; 02°09’ N, 31°14’ E; 1935; Ch. Scops leg.; RMCA. – **Kinshasa** • 1 ♂; Léopoldville (Kinshasa); 04°19’ S, 15°19’ E; Aug. 1949; R. Dubois leg.; RMCA – **Lomami** • 1 ♂; Lomami: Kabwe; 08°47’ S, 26°52’ E; Jul.-Aug. 1931; P. Quarré leg.; RMCA – **Lualaba** • 1 ♂; Haut-Luapula: Kansenia; 10°19’ S, 26°04’ E; 15 Oct. 1929, Dom de Montpellier leg.; RMCA • 1 ♂; Kapiri; 10°18’ S, 26°11’ E; Sep. 1912; Miss. Agric. leg.; RMCA – **Mongala** • 1 ♂; Ubangi: Binga (Lisala); 02°28’ N, 20°31’ E; 5-12 Mar. 1932; H.J. Brédo leg.; RMCA • 1 ♂; same location as for preceding; Feb. 5-12 Mar. 1935; H.J. Brédo leg.; RMCA. – **Nord-Kivu** • 1 ♂; Rutshuru; 01°11’ S, 29°27’ E; 17 May 1936; L. Lippens leg.; RMCA. – **Nord-Ubangi** • 2 ♂♂; Abumombazi (Mobayi); 03°34’ N, 22°03’ E; 18-26 Feb. 1932; H.J. Brédo leg.; RMCA –**Sankuru** • 1 ♂; Sankuru: Lukeni; 03°32’ S, 24°19’ E; Jan. 1928; J. Ghesquière leg.; RMCA. – **Sud-Ubangi** • 1 ♂; Ubangi: Boma-Motenge; 03°15’ N, 18°39’ E; Dec. 1931; H.J. Brédo leg.; RMCA. – **Tshopo** • 1 ♀; Basoko; 01°13’ N, 23°36’ E; Oct. 1948; P.L.G. Benoit leg.; RMCA. – **Tshuapa**• 1 ♀; Tshuapa: Bokungu; 00°41’ S, 22°19’ E; 1949; Dupuis leg.; RMCA.

#### *Megachile (Anodonteutricharaea*) *fulvitarsis* Friese, 1910, *Megachile* (*Paracella*) *fulvitarsis* Friese, 1910

**Material.** Cupboard 110, Box 16 (16♀♀and 4♂♂). **D.R. CONGO**. – **Haut-Katanga** • 1 ♀; Elisabethville (Lubumbashi); 11°40’ S, 27°28’ E; Nov. 1928; Ch. Seydel leg.; RMCA • 1 ♀; same location as for preceding; Feb. 1933, De Loose leg.; RMCA • 1 ♀; same location as for preceding; 1 Feb. 1935; P. Quarré leg.; RMCA • 7 ♀♀; same location as for preceding; Feb. 1935; P. Quarré leg.; RMCA • 1 ♀; same location as for preceding; Mar. 1935; P. Quarré leg.; RMCA • 1 ♀; same location as for preceding; Jun. 1935; P. Quarré leg.; RMCA • 1 ♀; same location as for preceding; Oct. 1934; P. Quarré leg.; RMCA • 1 ♀; same location as for preceding; Nov. 1934; P. Quarré leg.; RMCA • 1 ♀; same location as for preceding; Dec. 1935; P. Quarré leg.; RMCA • 1 ♂; PNU Kaziba; 09°09’ S, 26°57’ E, alt. 1140 m; 1-6 Feb. 1948; G.F. De Witte leg.; RMCA • 1 ♂; same location as for preceding; 16 Apr. 1948; G.F. De Witte leg.; RMCA. – **Haut-Lomami** • 1 ♂; Bukama; 09°12’ S, 25°51’ E; Aug. 1923; Ch. Seydel leg.; RMCA. – **Maniema** • 1 ♀; Kasongo; 04°27’ S, 26°40’ E; 1954; Dr. J. Claessens leg.; RMCA. – **Sud-Kivu** • 1 ♂; Luofu; 00°37’ S, 29°07’ E, alt. 1700 m; 10 Dec. 1934; G.F. De Witte leg.; RMCA.

#### *Megachile* (*Anodonteutricharaea*) *granulicauda* Cockerell, 1931, *Megachile* (*Paracella*) *granulicauda* Cockerell, 1931

**Material.** Cupboard 110, Box 16 (1 ♀). **D.R. CONGO**. Holotype – **Haut-Katanga** • 1 ♀; Lubumbashi; 11°40’ S, 27°28’ E; 18 Mar. 1921; Dr. M. Bequaert leg.; RMCA.

#### *Megachile* (*Anodonteutricharaea*) *guineae* Strand, 1912, *Megachile* (*Paracella*) *guineae* Strand, 1912

**Material.** Cupboard 110, Box 16 (2♀♀). **D.R. CONGO**. – **Haut-Katanga** • 1 ♀; PNU Kabwe sur Muye; 08°47’ S, 26°52’ E, alt. 1320 m; 11 May 1948; Mis. G.F. de Witte leg.; RMCA. – **Tanganyika** • 1 ♀; Vallée Lukuga; 05°40’ S, 26°55’ E; Nov. 1911; Dr. Schwetz leg.; RMCA.

#### *Megachile* (*Anodonteutricharaea*) *harrarensis* Friese, 1915, *Megachile* (*uicertae sedis*) *harrarensis* Friese, 1915

**Material.** Cupboard 110, Box 17 (1♀♂). **D.R. CONGO**. – **Tanganyika** • 1 ♀♂; Kongolo; 05°24’ S, 27°00’ E; 23 May 1911; Dr. J. Bequaert leg.; RMCA^2^.

#### *Megachile* (*Anodonteutricharaea*) *heteroscopa* Cockerell, 1937, *Megachile* (*Paracella*) *heteroscopa* Cockerell, 1937

**Material.** Cupboard 110, Box 16 (1♀). **D.R. CONGO**. – **Ituri** • 1 ♀; Mongbwalu (Kilo); 01°50’ N, 30°09’ E; 1939; Mme Scheitz leg.; RMCA.

#### *Megachile* (*Anodonteutricharaea*) *maculosella* Pasteels, 1965, *Megachile* (*Parcella*) *maculosella* Pasteels, 1965

**Material.** Cupboard 110, Box 16 (1♀). **D.R. CONGO**. Holotype – **Tshuapa** • 1 ♀; Bokuma; 00°06’ S, 18°41’ E; Jul. 1952; Rév. P. Lootens leg.; RMCA.

#### *Megachile* (*Anodonteutricharaea*) *meruensis* Friese, 1910, *Megachile* (*Paracella*) *meruensis* Friese, 1910

**Material.** Cupboard 110, Box 17 (1♀). **D.R. CONGO**. – **Tanganyika** • 1 ♀; Bassin Lukuga; 05°40’ S, 26°55’ E; 4 Jul. 1934; H. De Saeger leg.; RMCA.

#### *Megachile* (*Anodonteutricharaea*) *niveicauda* Cockerell, 1931, *Megachile* (*Paracella*) *meruensis* Friese, 1910

**Material.** Cupboard 110, Box 16 (2♂♂). **D.R. CONGO**. Holotype – **Haut-Katanga** • 1 ♂; Panda (Likasi); 11°00’ S, 26°44’ E; 9 Sep. 1920; Dr. M. Bequaert leg.; RMCA. – **Lualaba** • 1 ♂; Tenke; 10°35’ S, 26°06’ E; 30 Jul. – 9 Aug. 1931; J. Ogilvie leg.; RMCA.

#### *Megachile* (*Anodonteutricharaea*) *polychroma* Cockerell, 1937, *Megachile* (*Paracella*) *polychroma* Cockerell, 1937

**Material.** Cupboard 110, Box 15 (6 ♀♀). **D.R. CONGO**. – **Bas-Uele** • 1 ♀; Uele: Dingila; 03°37’ N, 26°03’ E; May 1933; H.J. Brédo leg.; RMCA. – **Equateur** • 1 ♀; Eala (Mbandaka); 00°03’ N, 18°19’ E; Dec 1932; A. Corbisier leg.; RMCA • 1 ♀; same location as for preceding; Mar. 1935; J. Ghesquière leg.; RMCA. – **Haut-Katanga** • 1 ♀; Elisabethville (Lubumbashi); 11°40’ S, 27°28’ E; Mar. 1935; P. Carré leg.; RMCA. – **Maniema** • 1 ♀; Kibombo; 03°54’ S, 25°55’ E; 3 Nov. 1910; Dr. J. Bequaert leg.; RMCA. – **Tshopo** • 1 ♀; Stanleyville (Kisangani); 00°31’ N, 25°11’ E; 27 Apr. 1915; Exp Lang-Chapin leg.; RMCA.

#### *Megachile* (*Anodonteutricharaea*) *selenostoma* Cockerell, 1931, *Megachile* (*Paracella*) *selenostoma* Cockerell, 1931

**Material.** Cupboard 110, Box 16 (1♀). **D.R. CONGO**. Holotype – **Nord-Kivu** • 1 ♀; Mt. Ruwenzori; 00°26’ S, 29°50’ E, alt. 1400 m; 5 Jun. 1914; Dr. J. Bequaert leg.; RMCA.

#### *Megachile* (*Anodonteutricharaea*) *semivenusta* Cockerell, 1931, *Megachile* (*Paracella*) *semivenusta* Cockerell, 1931

**Material.** Cupboard 110, Box 15 (45♀♀ and 102♂♂). **D.R. CONGO**. Paratypes – **Equateur** • 1 ♀; Coquilhatville (Mbandaka); 00°03’ N, 18°15’ E; 4 Feb. 1923; Dr. M. Bequaert leg.; RMCA. – **Haut-Katanga** • 2 ♀♀; Lubumbashi; 11°40’ S, 27°28’ E; 13 Mar. 1921; Dr. M. Bequaert leg.; RMCA • 1 ♀; same location as for preceding; 4 Feb. 1921; Dr. M. Bequaert leg.; RMCA.

– **Bas-Uele** • 1 ♀; Bambesa; 03°28’ N, 25°43’ E; 15 Oct. 1933; H.J. Brédo leg.; RMCA • 2 ♀♀; same location as for preceding; 30 Oct. 1933; H.J. Brédo leg.; RMCA • 1 ♀; same location as for preceding; Sep. 1933; J.V. Leroy leg.; RMCA. – **Equateur** • 1 ♂; Beneden-Congo: Tumba; 00°50’ S, 18°00’ E; Sep. 1938; Mevr. Bequaert leg.; RMCA • 1 ♂; Eala (Mbandaka); 00°03’ N, 18°19’ E; 1932; H.J. Brédo leg.; RMCA • 1 ♀; same location as for preceding; 21 Nov. 1931; H.J. Brédo leg.; RMCA • 1 ♀; same location as for preceding; 22 Nov. 1931; H.J. Brédo leg.; RMCA • 1 ♀; same location as for preceding; 7 Oct. 1931; H.J. Brédo leg.; RMCA • 1 ♀; same location as for preceding; May 1933; A. Corbisier leg.; RMCA • 1 ♂; same location as for preceding; May 1933; A. Corbisier leg.; RMCA • 1 ♀; same location as for preceding; Jun. 1935; J. Ghesquière leg.; RMCA • 1 ♀; same location as for preceding; Nov. 1935; J. Ghesquière leg.; RMCA • 1 ♂; Ubangi: Mokolo; 01°35’ N, 29°05’ E; 5 Dec. 1931; H.J. Brédo leg.; RMCA • 1 ♂; Ubangi: Yakoma; 04°06’ N, 22°23’ E; 17 Feb. 1932; H.J. Brédo leg.; RMCA. – **Haut-Katanga** • 1 ♂; Elisabethville (Lubumbashi); 11°40’ S, 27°28’ E; 11 – 17 Sep. 1931; J. Ogilvie leg.; RMCA • 1 ♂; same location as for preceding; 11°40’ S, 27°28’ E; 11 – 17 Sep. 1931; Mrs. W.P. Cockerell leg.; RMCA • 1 ♂; same location as for preceding; 16 Aug. 1932; De Loose leg.; RMCA • 1 ♂; same location as for preceding; 2 May 1920; Dr. M. Bequaert leg.; RMCA • 1 ♀; same location as for preceding; 27 Aug. 1932; De Loose leg.; RMCA • 1 ♀; same location as for preceding; Sep. 1928; Ch. seydel leg.; RMCA • 2 ♂♂; same location as for preceding; Nov. 1928; Ch. Seydel leg.; RMCA • 1 ♀; Elisabethville (60km ruiss. Kasepa); 11°40’ S, 27°28’ E; 23 Sep. 1923; Dr. M. Bequaert leg.; RMCA • 1 ♂; Elisabethville (Nord Tshisangwe); 11°40’ S, 27°28’ E; 4 Feb. 1923; Dr. M. Bequaert leg.; RMCA • 1 ♀; Kalunkumia; 10°49’ S, 26°51’ E; 3 Mar. 1925; Ch. Seydel leg.; RMCA • 1 ♂; Lubumbashi; 11°40’ S, 27°28’ E; 10 Feb. 1920; Dr. M. Bequaert leg.; RMCA • 2 ♂♂; same location as for preceding; 11 Apr. 1921; Dr. M. Bequaert leg.; RMCA • 1 ♂; same location as for preceding; 2 May 1920; Dr. M. Bequaert leg.; RMCA • 2 ♂♂; same location as for preceding; 23 May 1920; Dr. M. Bequaert leg.; RMCA • 1 ♂; same location as for preceding; 27 May 1920; Dr. M. Bequaert leg.; RMCA • 1 ♂; same location as for preceding; 28 May 1920; Dr. M. Bequaert leg.; RMCA • 3 ♂♂; same location as for preceding; 4 Feb. 1921; Dr. M. Bequaert leg.; RMCA • 1 ♂; same location as for preceding; 8 Dec. 1920; Dr. M. Bequaert leg.; RMCA • 1 ♀; Lukafu; 10°31’ S, 27°33’ E; 15 Sep. 1930; Dr. M. Bequaert leg.; RMCA • 2 ♀♀; PNU Kabwe sur Muye; 08°47’ S, 26°52’ E; alt. 1320 m; 11 May 1948; G.F. de Witte leg.; RMCA • 1 ♂; PNU Kolwezi (Affl. Dr. Lufira); 09°05’ S, 26°45’ E, alt. 750 m; 27 Aug. – 8 Sep. 1948; G.F. de Witte leg.; RMCA • 1 ♂; PNU Masombwe s. Grde, Kafwe; 09°05’ S, 27°12’ E, alt. 1120 m; 16 Apr. 1948; G.F. de Witte leg.;

RMCA. – **Haut-Lomami** • 1 ♂; Bukama; 09°12’ S, 25°51’ E; 14 Jul. 1911; Dr. J. Bequaert leg.; RMCA • 1 ♀; same location as for preceding; 17 Jul. 1911; Dr. J. Bequaert leg.; RMCA • 1 ♀; same location as for preceding; 19 Aug. 1911; Dr. J. Bequaert leg.; RMCA • 1 ♂; same location as for preceding; 19 Aug. 1911; Dr. J. Bequaert leg.; RMCA • 1 ♂; same location as for preceding; 22 Mar. 1911; Dr. J. Bequaert leg.; RMCA • 1 ♂; same location as for preceding; 29 May 1911; Dr. J. Bequaert leg.; RMCA • 1 ♂; same location as for preceding; 7 Aug. 1911; Dr. J. Bequaert leg.; RMCA • 1 ♂; Kadiamapanga; 08°16’ S, 26°36’ E; Nov. 1931; H.J. Brédo leg.; RMCA • 1 ♂; PNU Kaswabilenga; 08°51’ S, 26°43’ E, alt. 700 m; 21 Oct. 1947; G.F. de Witte leg.; RMCA • 1 ♂; PNU Mabwe (Lac Upemba); 08°39’ S, 26°31’ E, alt. 585 m; 28 Aug. 1947; G.F. de Witte leg.; RMCA • 1 ♀; same location as for preceding; 4 Sep. 1947; G.F. de Witte leg.; RMCA • 3 ♂♂; Sankisia; 09°24’ S, 25°48’ E; 1 Sep. 1911; Dr. J. Bequaert leg.; RMCA • 2 ♂♂; same location as for preceding; 3 Sep. 1911; Dr. J. Bequaert leg.; RMCA • 1 ♂; same location as for preceding; 30 Sep. 1911; Dr. J. Bequaert leg.; RMCA • 3 ♂♂; same location as for preceding; 4 Sep. 1911; Dr. J. Bequaert leg.; RMCA. – **Haut-Uele** • 1 ♂; Haut-Uele: Abimva; 03°44’ N, 29°42’ E; 1925; L. Burgeon leg.; RMCA • 1 ♂; Haut-Uele: Moto; 03°03’ N, 29°28’ E; 1920; L. Burgeon leg.; RMCA • 2 ♂♂; PNG [Parc National de Garamba]; 03°40’ N, 29°00’ E; 18 Dec. 1951; H. De Saeger leg.; RMCA • 1 ♀; same location as for preceding; 24 Jun. 1952; H. De Saeger leg.; RMCA • 1 ♀; same location as for preceding; 25 Jun. 1952; H. De Saeger leg.; RMCA • 1 ♂; same location as for preceding; 25 Jun. 1952; H. De Saeger leg.; RMCA • 1 ♂; same location as for preceding; 3 Jan. 1952; H. De Saeger leg.; RMCA • 1 ♂; same location as for preceding; 4 Aug. 1952; H. De Saeger leg.; RMCA • 1 ♂; same location as for preceding; 9 Nov. 1950; H. De Saeger leg.; RMCA • 3 ♀♀; PNG Akam; 03°40’ N, 29°00’ E; 22 Apr. 1950; H. De Saeger leg.; RMCA • 5 ♂♂; same location as for preceding; 22 Apr. 1950; H. De Saeger leg.; RMCA • 1 ♀; PNG Mabanga; 04°22’ N, 29°47’ E; 29 Sep. 1952; H. De Saeger leg.; RMCA • 2 ♀♀; PNG Mt Tungu (S); 03°40’ N, 29°00’E; 9 Jun. 1952; H. De Saeger leg.; RMCA • 1 ♀; PNG Utukuru; 03°40’ N, 29°00’ E; 22 Jul. 1952; H. De Saeger leg.; RMCA • 1 ♂; Watsa à Niangara; 03°02’ N, 29°32’ E; Jul. 1920; L. Burgeon leg.; RMCA. – **Ituri** • 1 ♂; Ituri: La Moto (Madyu); 01°34’ N, 30°14’ E; L. Burgeon leg.; RMCA • 1 ♂; Kibimbi; 01°22’ N, 28°55’ E; 2 Feb. 1911; Dr. J. Bequaert leg.; RMCA. – **Kasaï** • 1 ♀; Luluabourg; 05°54’ S, 21°52’ E; P. Callewaert leg.;

RMCA. – **Kinshasa** • 3 ♂♂; Léopoldville (Kinshasa); 04°19’ S, 15°19’ E; 16 Sep. 1910; Dr. J. Bequaert leg.; RMCA • 2 ♂♂; same location as for preceding; 18 Sep. 1910; Dr. J. Bequaert leg.; RMCA • 1 ♂; same location as for preceding; 25 Sep. 1910; Dr. J. Bequaert leg.; RMCA • 1 ♀; Stanley Pool; 04°12’ S, 15°33’ E; 13 Aug. 1958; Dr. J. Pasteels leg.; RMCA. – **Kongo Central** • 1 ♀; Banana; 06°00’ S, 12°24’ E; Aug. 1910; Dr. Etienne leg.; RMCA • 1 ♂; Congo da Lemba (Songololo); 05°42’ S, 13°42’ E; Mar. 1913; R. Mayné leg.; RMCA • 1 ♂; same location as for preceding; Jan – Feb. 1913; R. Mayné leg.; RMCA • 1 ♀; Kisantu (Madimba); 05°08’ S, 15°06’ E; 1927; R.P. Vanderyst leg.; RMCA • 1 ♂; same location as for preceding; 27 Sep. 1910; Dr. J. Bequaert leg.; RMCA. – **Lomami** • 4 ♂♂; Lomami: Kabwe; 08°47’ S, 26°52’ E; Jul. – Aug. 1931; P. Quarré leg.; RMCA. – **Lualaba** • 1 ♂; Biano; 10°20’ S, 27°00’ E; 8 – 11 Jul. 1931; P. Quarré leg.; RMCA • 1 ♂; Dilolo; 10°41’ S, 22°21’ E; 24 – 27 Jul. 1931; G.F. de Witte leg.; RMCA • 1 ♂; same location as for preceding; Aug. 1931; J. Ogilvie leg.; RMCA • 1 ♂; Kalule Sud; 09°36’ S, 25°37’E; Oct. 1931; Ch. Seydel leg.; RMCA • 1 ♂; Kanzenze; 10°31’ S, 25°12’ E; Aug. 1931; G.F. de Witte leg.; RMCA • 1 ♂; La Mulonga; 09°56’ S, 25°53’ E; Aug. 1931; Miss. H. De Saeger leg.; RMCA • 1 ♀; Lualaba: Kolwezi; 10°44’ S, 25°28’ E; 1 – 6 Nov. 1952; Mme. L. Gilbert leg.; RMCA • 2 ♂♂; Lukata; 10°05’ S, 25°55’ E; Oct. 1931; Ch. Seydel leg.; RMCA • 2 ♂♂; Tenke; 10°35’ S, 26°06’ E; 30 Jul. – 9 Aug. 1931; J. Ogilvie leg.; RMCA • 1 ♀; same location as for preceding; 30 Jul. – 9 Aug. 1931; T.D. Cockerell leg.; RMCA. – **Maï-Ndombe** • 1 ♂; Bolobo (Mushie); 02°10’ S, 16°14’ E 24 Apr. 1924; Dr. H. Schouteden leg.; RMCA. – **Maniema** • 1 ♀; Kibombo; 03°54’ S, 25°55’ E; 10 Nov. 1910; Dr. M. Bequaert leg.; RMCA. – **Mongala** • 1 ♀; Bumba; 02°11’ N, 22°32’ E; Dec. 1939 – Jan. 1940; H. De Saeger leg.; RMCA • 1 ♀; Ubangi: Binga (Lisala); 02°28’ N, 20°31’ E; 5-12 Mar. 1932; H.J. Brédo leg.; RMCA. – **Nord-Kivu** • 1 ♂; PNA SL Edouard: Bitshumbi; 00°20’ S, 29°30’ E, alt. 925 m; 15 Apr. 1936; L. Lippens leg.; RMCA • 2 ♂♂; same location as for preceding; 3 Apr. 1936; L. Lippens leg.; RMCA • 2 ♂♂; Rutshuru; 01°11’ S, 29°27’ E; 11 May 1936; L. Lippens leg.; RMCA • 1 ♂; same location as for preceding; 15 May 1936; L. Lippens leg.; RMCA • 1 ♀; same location as for preceding; Feb. – May 1936; L. Lippens leg.; RMCA • 1 ♂; same location as for preceding; Dec 1937; J. Ghesquière leg.; RMCA • 1 ♀; Walikale; 01°25’ S, 28°03’ E; 7 Jan. 1915; Dr. J. Bequaert leg.;

RMCA. – **Nord-Ubangi** • 1 ♂; Ubangi: Bosobolo; 04°11’ N, 19°53’ E; 8 – 11 Jan. 1932; H.J. Brédo leg.; RMCA. – **Sud-Kivu** • 1 ♂; Kalembelembe-Baraka; 04°32’ S, 28°47’ E; Aug. 1918; R. Mayné leg.; RMCA. – **Sud-Ubangi** • 1 ♀; Libenge; 03°39’ N, 18°38’ E; 9 Dec. 1931; H.J. Brédo leg.; RMCA. • 1 ♂; Ubangi: Nzali; 03°15’ N, 19°47’ E; 3 – 4 Feb. 1932; H.J. Brédo leg.; RMCA. – **Tanganyika** • 1 ♂; Albertville (Kalemie); 05°57’ S, 29°12’ E Dec. 1918; Miss. H. De Saeger leg.; RMCA • 1 ♂; Kongolo; 05°24’ S, 27°00’ E; 16 Aug. 1921; Mlle. Brunard leg.; RMCA • 2 ♂♂; Lomami: Katompe (Kabalo); 06°11’ S, 26°20’ E; 20 Dec. 1923; Dr. M. Bequaert leg.; RMCA • 3 ♂♂; Tanganyika-Moero: Nyunzu; 05°57’ S, 28°01’ E; 1935; H. De Saeger leg.; RMCA. – **Tshopo** • 2 ♂♂; Stanleyville (Kisangani); 00°31’ N, 25°11’ E; 20 Oct. 1910; Dr. J. Bequaert leg.; RMCA.

#### *Megachile* (*Anodonteutricharaea*) *ungulata* Smith, 1953, *Megachile* (*Paracella*) *ungulata* Smith, 1953

**Material.** Cupboard 110, Box 17 (1♂). **D.R. CONGO**. – **Haut-Katanga** • 1 ♂; PNU Lusinga, Rivière Dipidi; 08°55’ S, 27°12’ E; 12 Jun. 1945; G.F. de Witte leg.; RMCA.

#### *Megachile* (*Creightonella*) *aculeata* Vachal, 1910, *Creightonella aculeata* (Vachal, 1910)

**Material.** Cupboard 110, Box 51 (30♂♂). **D.R. CONGO**. Paratypes – **Haut-Katanga** • 1 ♂; Kiamokosa; 11°40’ S, 27°28’ E; Sep. 1907; Dr. Sheffield Neave leg.; RMCA • 1 ♂; Kipaila-Kisinga; 10°47’ S, 28°30’ E; Sep. 1907; Dr. Sheffield Neave leg.; RMCA. – **Tanganyika** • 1 ♂; Kanikiri; 00°09’ S, 29°32’ E; Sep. 1907; Dr. Sheffield Neave leg.; RMCA.

– **Haut-Katanga** • 1 ♂; Elisabethville (Lubumbashi); 11°40’ S, 27°28’ E; 1931; De Loose.; RMCA • 1 ♂; same location as for preceding; 11 Sep. 1932; Dr. M. Bequaert leg.; RMCA • 1 ♂; same location as for preceding; 12 Nov. 1933; Ch. Seydel leg.; RMCA • 1 ♂; same location as for preceding; 2 Jul. 1933; Dr. M. Bequaert leg.; RMCA • 2 ♂♂; same location as for preceding; 20 Oct. 1929; Dr. M. Bequaert leg.; RMCA • 1 ♂; same location as for preceding; Jun. 1932; De Loose leg.; RMCA • 1 ♂; same location as for preceding; Dec. 1925; Ch. Seydel leg.; RMCA • 1 ♂; Gombela; 10°47’ S, 27°48’ E; Jun. 1929; Ch. Seydel leg.; RMCA • 1 ♂; Kasenga; 10°22’ S, 28°38’ E; Apr. 1931; H.J. Brédo leg.; RMCA • 2 ♂♂; Lubumbashi; 11°40’ S, 27°28’ E; 10 May 1953; Ch. Seydel leg.; RMCA • 1 ♂; same location as for preceding; 7 Jan. 1921; Dr. M. Bequaert leg.; RMCA • 1 ♂; PNU Lusinga (riv. Kamitungulu); 08°55’ S, 08°55’ S, 13 Jun. 1945; Mis. G.F. de Witte leg.; RMCA • 2 ♂♂; PNU Masombwe s. Grde. Kafwe; 09°05’ S, 27°12’ E, alt. 1120 m; 16 Apr. 1948; Mis. G.F. de Witte leg.; RMCA. – **Haut-Lomami** • 1 ♂; Bukama; 09°12’ S, 25°51’ E; 23 Mar. 1911; Dr. J. Bequaert leg.; RMCA • 1 ♂; Sankisia; 09°24’ S, 25°48’ E; 4 Sep. 1911; Dr. J. Bequaert leg.; RMCA. – **Lualaba** • 1 ♂; Biano; 10°20’ S, 27°00’ E; Apr. 1927; R. Mayné leg.; RMCA • 1 ♂; Sandu; 09°41’ S, 22°53’ E; Apr. 1921; H.J. Brédo leg.; RMCA. – **Maniema** • 1 ♂; Nyangwe; 04°13’ S, 26°11’ E; 8 Dec. 1910; Dr. J. Bequaert leg.; RMCA. – **Nord-Kivu** • 1 ♂; PNA: S.L. Kitembo; 00°20’ S, 29°30’ E; 13 Apr. 1936; L. Lippens leg.; RMCA. – **Sud-Kivu** • 1 ♂; Kalembelembe-Baraka; 04°32’ S, 28°47’ E; Jul. 1918; R. Mayné leg.; RMCA. – **Tanganyika** • 1 ♂; Kiambi (Manono); 07°19’ S, 28°01’ E; 27 Apr. 1931; G.F. de Witte leg.; RMCA • 1 ♂; Kongolo; 05°24’ S, 27°00’ E; 23 Jan. 1911; G.F. de Witte leg.; RMCA • 1 ♂; same location as for preceding; 30 Feb. 1911; G.F. de Witte leg.; RMCA • 1 ♂; Tanganyika-Moero: Nyunzu; 05°57’ S, 28°01’ E; Jan. – Feb. 19134; H. Bomans leg.; RMCA.

#### *Megachile* (*Creightonella*) *adeloptera* Schletterer, 1891, *Creightonella adeloptera* (Schletterer, 1891)

**Material.** Cupboard 110, Box 46 (53♀♀ and 3♂♂). **D.R. CONGO**. – **Bas-Uele** • 1 ♂; Bambesa; 03°28’ N, 25°43’ E; 14 May 1938; P. Henrard leg.; RMCA • 1 ♀; same location as for preceding; 6 Aug. 1937; J. Vrydagh leg.; RMCA • 1 ♀; same location as for preceding; Nov. 1933; H.J. Brédo leg.; RMCA • 1 ♀; Mobwasa; 02°40’ N, 23°11’ E; Sep. 1911; De Giorgi leg.; RMCA. – **Equateur** • 1 ♀; Coquilhatville (Mbandaka); 00°03’ N, 18°15’ E; 11 May 1925; Dr. M. Bequaert leg.; RMCA • 1 ♀; Eala (Mbandaka); 00°03’ N, 18°19’ E; 14 Mar. 1933; A. Corbisier leg.; RMCA • 1 ♀; same location as for preceding; 7 Jul. 1932; A. Corbisier leg.; RMCA • 1 ♂; same location as for preceding; Mar. 1932; H.J. Brédo leg.; RMCA • 1 ♂; same location as for preceding; Apr. 1933; A. Corbisier leg.; RMCA • 1 ♀; same location as for preceding; May 1932; H.J. Brédo leg.; RMCA • 2 ♀♀; same location as for preceding; Jun. 1932; A. Corbisier leg.; RMCA • 1 ♀; Flandria (Ingende); 00°20’ S, 19°06’ E; Jan. 1932; R.P. Hulstaert leg.; RMCA. – **Ituri** • 18 ♀♀; Haut-Ituri; 01°53’ N, 29°03’ E; 1 – 3 May. 1931;1906; H. Wilmin leg.; RMCA. – **Maniema** • 1 ♀; Kibombo; 03°54’ S, 25°55’ E; 18 Jan. 1910; Dr. J. Bequaert leg.; RMCA • 1 ♀; Lokandu (Biawa); 02°31’ S, 25°47’ E; Dec. 1939; R.P. Vankerckhoven leg.; RMCA • 1 ♀; Malela; 04°22’ S, 26°08’ E; 12-18 Dec. 1913; L. Burgeon leg.; RMCA. – **Mongala** • 1 ♀; Congo: Bolombo; 02°38’ N, 21°36’ E; ex coll. Breuning leg.; RMCA • 1 ♀; Itimbiri; 02°03’ N, 22°45’ E; 23 May 1913; Dr. Rodhain leg.; RMCA. – **Nord-Ubangi** • 2 ♀♀; Ubangi: Karawa (Businga); 03°21’ N, 20°18’ E; 1936; Rév. Wallin leg.; RMCA. – **Sankuru** • 1 ♀; Sankuru: Komi; 03°23’ S, 23°46’ E; 3 Feb. 1930; J. Ghesquière leg.; RMCA • 1 ♀; same location as for preceding; Feb. 1930; J. Ghesquière leg.; RMCA • 2 ♀♀; same location as for preceding; Jul. 1930; J. Ghesquière leg.; RMCA. – **Sud-Kivu** • 1 ♀; Kivu; 05°29’ S, 28°51’ E; Sep.-Oct. 1925; S.A.R. Prince Léopold leg.; RMCA. – **Tanganyika** • 2 ♀♀; Lukulu (Manono); 07°08’ S, 28°06’ E; 1 – 3 May 1931; G.F. de Witte leg.; RMCA. – **Tshopo** • 1 ♀; Lula (Kisangani); 00°27’ N, 25°12’ E; 1958; A.J. Jobaert leg.; RMCA • 1 ♀; Miss. St. Gabriel; 00°33’ N, 25°05’ E; M. Torley leg.; RMCA • 1 ♀; Utisongo (Stanleyville); 00°31’ N, 25°11’ E; 22 Apr. 1932; J. Vrydagh leg.; RMCA. – **Tshuapa** • 4 ♀♀; Boende; 00°13’ S, 20°52’ E; Sep. – Oct. 1925; Rév. P. Lootens leg.; RMCA • 1 ♂; Bokuma; 00°06’ S, 18°41’ E; Jul. 1952; Rév. P. Lootens leg.; RMCA • 2 ♀♀; same location as for preceding; Dec. 1951; Rév. P. Lootens leg.; RMCA • 1 ♀; Moma; 00°45’ S, 21°56’ E; May 1925; J. Ghesquière leg.; RMCA.

#### *Megachile* (*Creightonella*) *angulata* Smith, 1853, *Creightonella angulata* (Smith, 1853)

**Material.** Cupboard 110, Box 47 (24♀♀ and 19♂♂). **D.R. CONGO**. – **Haut-Katanga** • 1 ♂; Elisabethville(Lubumbashi); 11°40’ S, 27°28’ E; De Loose leg.; RMCA • 1 ♀; Elisabethville (60km ruiss. Kasepa); 11°40’ S, 27°28’ E; 22 Sep. 1923; Dr. M. Bequaert leg.; RMCA • 1 ♀; Kasenga; 10°22’ S, 28°38’ E; Oct. 1929; H.J. Brédo leg.; RMCA • 3 ♂♂; Katumba; 08°32’ S, 25°59’ E; Oct. 1907; Dr. Sheffield Neave leg.; RMCA • 3 ♂♂; PNU Lusinga (riv. Kamitungulu); 08°55’ S, 08°55’ S, 13 Jun. 1945; Mis. G.F. de Witte leg.; RMCA. – **Haut-Lomami** • 2 ♀♀; Lomami: Mutombo-Mukulu; 07°58’ S, 24°00’ E; Jun. 1931; P. Quarré leg.; RMCA • 1 ♀; Lualaba: Kabongo; 07°20’ S, 25°35’ E; 31 Dec. 1952; Ch. Seydel leg.; RMCA • 1 ♀; same location as for preceding; 5 Jan. 1953; Ch. Seydel leg.; RMCA • 1 ♀; Sankisia; 09°24’ S, 25°48’ E; 15 Sep. 1911; Dr. J. Bequaert leg.; RMCA • 1 ♀; same location as for preceding; 21 Sep. 1911; Dr. J. Bequaert leg.; RMCA • 2 ♀♀; same location as for preceding; 24 Sep. 1911; Dr. J. Bequaert leg.; RMCA • 2 ♂♂; same location as for preceding; 3 Sep. 1911; Dr. J. Bequaert leg.; RMCA. – **Kasaï Central** • 2 ♀♀; Luluabourg (Kananga); 05°54’ S, 21°52’ E; P. Callewaert leg.; RMCA. – **Kasaï Oriental** • 1 ♂; Sankuru: Bakwanga (Dibindi); 06°08’ S, 23°38’ E; 18 Jun. 1939; Mevr. Bequaert leg.; RMCA. – **Kongo Central** • 2 ♀♀; Mayidi (Madimba); 05°11’ S, 15°09’ E; 1942; Rév. P. Van Eyen leg.; RMCA. – **Lualaba** • 2 ♂♂; Bunkeya; 10°24’ S, 26°58’ E; Oct. 1907; Dr. Sheffield Neave leg.; RMCA • 2 ♂♂; Dilolo; 10°41’ S, 22°21’ E; Aug. 1931; G.F. de Witte leg.; RMCA • 1 ♀; Ditanto; 10°15’ S, 25°53’ E; Oct. 1925; Ch. Seydel leg.; RMCA • 1 ♂; Haut-Luapula: Kansenia; 10°19’ S, 26°04’ E; 18 Oct. 1929; Dom de Montpellier leg.; RMCA • 1 ♀; Lulua: Kapanga; 08°21’ S, 22°34’ E; Oct. 1932; F.G. Overlaet leg.; RMCA. – **Maï-Ndombe** • 1 ♀; Bumbuli; 03°24’ S, 20°31’ E; Jan.-Apr. 1915; R. Mayné leg.; RMCA. – **Maniema** • 1 ♂; Kindu; 02°57’ S, 25°55’ E; 7 Jun. 1912; L. Burgeon leg.; RMCA • 1 ♀; Nyangwe; 04°13’ S, 26°11’ E; 1918; R. Mayné leg.; RMCA. – **Nord-Kivu** • 1 ♂; Kanikiri; 00°09’ S, 29°32’ E; Oct. 1909; Dr. J. Schouteden leg.; RMCA. – **Sankuru** • 1 ♀; Sankuru: Djeka; 02°57’ S, 23°33’ E; 1955-1956; R. Roiseux leg.; RMCA • 1 ♀; Sankuru: Komi; 03°23’ S, 23°46’ E; May 1930; J. Ghesquière leg.; RMCA. – **Tanganyika** • 1 ♀; Bassin Lukuga; 05°40’ S, 26°55’ E; Apr.-Jul. 1934; H. De Saeger leg.; RMCA • 1 ♀; Katanga: Kabinda (Moba); 07°02’ S, 27°47’ E; Jan. 1926; Ch. Seydel leg.; RMCA • 1 ♀; Tanganyika-Moero: Nyunzui; 05°57’ S, 28°01’ E; 1935; H. Bomans leg.; RMCA • 2 ♂♂; same location as for preceding; 1931; H. Bomans leg.; RMCA.

#### *Megachile* (*Creightonella*) *aurivillii* Friese, 1901, *Creightonella aurivillii* (Friese, 1901)

**Material.** Cupboard 110, Box 46 (8♀♀ and 2♂♂). **D.R. CONGO**. – **Haut-Katanga** • 1 ♂; PNU : Lusinga (riv. Kamitungulu); 08°55’ S, 27°12’ E; 13 Jun. 1945; Mis. G.F. de Witte leg.; RMCA. – **Ituri** • 3 ♀♀; Haut-Ituri, 29°03’, 29°03’ E; 1906; H. Wilmin leg.; RMCA. – **Lomami** • 1 ♀; Lomami: Lusuku, 07°19’ S, 23°53’ E; Nov. 1930; R. Massart leg.; RMCA. –**Lualaba** • 1 ♀; Dilolo; 10°41’ S, 22°21’ E; Sep. – Oct.1933; H. De Saeger leg.; RMCA • 1 ♀; Kanzenze; 10°31’ S, 25°12’ E; Aug.1931; G.F. de Witte leg.; RMCA • 1 ♀; Lulua : Kapanga; 08°21’ S, 22°34’ E; Dec. 1932; F.G. Overlaet leg.; RMCA ; 1 ♂ ; same collection data as for preeceding ; Dec. 1932; F.G. Overlaet leg.; RMCA. – **Tshuapa** • 1 ♀; Moma; 00°45’ S, 21°56’ E; Jun. 1925; J. Ghesquière leg.; RMCA.

#### *Megachile* (*Creightonella*) *cognata* Smith, 1853, *Creightonella cognata* (Smith, 1853)

**Material.** Cupboard 110, Box 47 (28♀♀ and 17♂♂). **D.R. CONGO**. – **Haut-Katanga** • 1 ♂; Elisabethville (Lubumbashi); 11°40’ S, 27°28’ E; 7 Sep. 1932; De Loose leg.; RMCA • 1 ♀; same location as for preceding; Jun. 1932; De Loose leg.; RMCA • 1 ♂; same location as for preceding; Mar. 1921; Dr. M. Bequaert leg.; RMCA • 1 ♂; Kambove-Bunkeya; 10°52’ S, 26°38’ E; Oct. 1907; Dr. Sheffield Neave leg.; RMCA • 1 ♂; Kasenga; 10°22’ S, 28°38’ E; Oct. 1925; H.J. Brébo leg.; RMCA • 1 ♂; Katanga: Mwera; 11°09’ S, 27°19’ E; 1956; R.P.Th. de Caters leg.; RMCA • 1 ♀; Kundelungu; 10°40’ S, 28°00’ E; Sep. 1907; Dr. Sheffield Neave leg.; RMCA • 1 ♀; same location as for preceding; Jun. 1949; A. Zielinski leg.; RMCA • 1 ♂; Lubumbashi; 11°40’ S, 27°28’ E; 29 Jun. 1953; Ch. Seydel leg.; RMCA • 2 ♀♀; PNU Kabwe sur Muye; 08°47’ S, 26°52’ E, Alt. 1320 m; 11 May 1948; Mis. G.F. de Witte leg.; RMCA • 1 ♀; PNU Lusinga; 08°55’ S, 27°12’ E, alt. 1760 m; 25 Mar. 1947; Mis. G.F. de Witte leg.; RMCA • 8 ♀♀; PNU Lusinga (riv. Kamitungulu); 08°55’ S, 08°55’ S, 13 Jun. 1945; Mis. G.F. de Witte leg.; RMCA. – **Haut-Lomami** • 1 ♂; Kamina; 08°44’ S, 25°00’ E; 27 Nov. 1925; Ch. Seydel leg.; RMCA • 1 ♀; Lomami: Mutombo-Mukulu; 07°58’ S, 24°00’ E; Mar. 1931; P. Quarré leg.; RMCA • 1 ♀; Lualaba: Kabongo; 07°20’ S, 25°35’ E; 27 Dec. 1952; Ch. Seydel leg.; RMCA • 1 ♂; same location as for preceding; 27 Dec. 1952; Ch. Seydel leg.; RMCA • 1 ♀; same location as for preceding; 3 Jan. 1953; F.G. Overlaet leg.; RMCA • 1 ♂; same location as for preceding; 30 Dec. 1952; Ch. Seydel leg.; RMCA • 1 ♀; same location as for preceding; 31 Dec. 1952; Ch. Seydel leg.; RMCA • 1 ♂; same location as for preceding; 7 Jan. 1953; Ch. Seydel leg.; RMCA • 1 ♂; Katanga: Luashi; 07°31’ S, 24°10’ E; 1935; Freyne leg.; RMCA • 1 ♂; Lulua: Luashi; 07°31’ S, 24°10’ E; Mar. 1936; Freyne leg.; RMCA • 1 ♀; PNU Gorge de la Petenge; 08°56’ S, 27°12’ E, alt. 1150 m; 22 May-22 Jun. 1947; Mis. G.F. de Witte leg.; RMCA • 1 ♀; PNU Munoi, bifure Lupiala; 08°50’ S, 26°44’ E, alt. 890 m; 28 May-15 Jun. 1948; Mis. G.F. de Witte leg.; RMCA. – **Ituri**, • 1 ♂; Mahagi-Port; 02°09’ N, 31°14’ E; Oct. 1934; H.J. Brédo leg.; RMCA. – **Kasaï-Central** • 1 ♀; Luluabourg (Kananga); 05°54’ S, 21°52’ E; 18 Mar. 1919; P. Callewaert leg.; RMCA. – **Kongo Central** • 1 ♀; Kisantu (Madimba); 05°08’ S, 15°06’ E; 1927; R. P. Vanderyst leg.; RMCA • 1 ♀; Mayidi (Madimba); 05°11’ S, 15°09’ E; 1942; Rév. P. Van Eyen leg.; RMCA 1 ♂; same location as for preceding; 1942; Rév. P. Van Eyen leg.; RMCA. – **Kwilu** • 1 ♀; Leverville (Bulungu); 04°50’ S, 18°44’ E; 1927; Mme. J. Tinant leg.; RMCA. – **Lualaba** • 1 ♂; Kansenia; 10°19’ S, 26°04’ E; 15 Sep.-15 Oct. 1930; G.F. de Witte leg.; RMCA • 1 ♀; Lulua: Kapanga; 08°21’ S, 22°34’ E; Jan. 1933; F.G. Overlaet leg.; RMCA 1 ♀; same location as for preceding; Apr. 1933; F.G. Overlaet leg.; RMCA 2 ♂♂; same location as for preceding; Oct. 1932; F.G. Overlaet leg.; RMCA. – **Sankuru** • 1 ♀; Sankuru: Tshumbe S. Marie; 04°02’ S, 22°41’ E; May 1948; Mevr. Bequaert leg.; RMCA. – **Sud-Ubangi** • 1 ♀; Kunungu; 02°45’ N, 18°26’ E; 1932; Dr. H. Schouteden leg.; RMCA.

#### *Megachile* (*Creightonella*) *discolor* Smith, 1853, *Creightonella discolor* (Smith, 1853)

**Material.** Cupboard 110, Box 47 (5♀♀ and 11♂♂). **D.R. CONGO**. – **Haut-Katanga** • 1 ♀; Elisabethville (Lubumbashi); 11°40’ S, 27°28’ E; 12 Jul. 1951; Ch. Seydel leg.; RMCA • 1 ♂; same location as for preceding; 12 Jul. 1951; Ch. Seydel leg.; RMCA • 2 ♀♀; Kasenga; 10°22’ S, 28°38’ E; Apr. 1931; Ch. Seydel leg.; RMCA • 1 ♂; Kiswishi; 11°30’ S, 27°26’ E; De Loose leg.; RMCA • 1 ♂; Lubumbashi; 11°40’ S, 27°28’ E; 16 Jan. 1921; Dr. M. Bequaert leg.; RMCA • 1 ♀; Lukafu-Bunkeya; 10°31’ S, 27°33’ E; Oct. 1907; Dr. Sheffield Neave leg.; RMCA • 1 ♂; Sankisia; 09°24’ S, 25°48’ E; 3 Sep. 1911; Dr. J. Bequaert leg.; RMCA. – **Haut-Lomami** • 1 ♂; PNU Mabwe (R.P. Lac Upemba); 08°47’ S, 26°52’ E, alt. 585 m; 17-20 Nov. 1948; Mis. G.F. de Witte leg.; RMCA. – **Maniema** • 1 ♂; Nyangwe; 04°13’ S, 26°11’ E; Apr. – May 1918; R. Mayné leg.; RMCA. – **Sud-Kivu** • 2 ♂♂; Kalembelembe-Baraka; 04°32’ S, 28°47’ E; Jul. 1918; R. Mayné leg.; RMCA. – **Tanganyika** • 1 ♂; Albertville (Kalemie); 05°57’ S, 29°12’ E; Dec. 1918; Miss. H. De Saeger leg.; RMCA • 1 ♂; Kanikiri; 00°09’ S, 29°32’ E; Oct. 1907; Dr. Sheffield Neave leg.; RMCA • 1 ♂; Kiambi (Manono); 07°19’ S, 28°01’ E; 27 Apr. 1931; G.F. de Witte leg.; RMCA • 1 ♀; Kongolo; 05°24’ S, 27°00’ E; 25 Jan. 1911; Dr. J. Bequaert leg.; RMCA.

#### *Megachile* (*Creightonella*) *gastracantha* Cockerell, 1931, *Creightonella gastracantha* (Cockerell, 1931)

**Material.** Cupboard 110, Box 46 (1♂). **D.R. CONGO**. – **Haut-Katanga** • 1 ♂; PNU Kilwezi affl. dr. Lufira; 09°05’ S, 26°45’ E, alt. 750 m; 9 – 14 Jul. 1948; Mis. G.F. de Witte leg.; RMCA.

#### *Megachile* (*Creightonella*) *hoplitis* Vachal, 1903, *Creightonella hoplitis* (Vachal, 1903)

**Material.** Cupboard 110, Box 47 (2♀♀ and 13♂♂). **D.R. CONGO**. – **Haut-Lomami** • 1 ♂; Lomami: Lulumbo; 05°24’ S, 25°19’ E; Jul. 1930; P. Quarré leg.; RMCA • 1 ♀; same location as for preceding; 7 Jan. 1953; Ch. Seydel leg.; RMCA. – **Kasaï** • 2 ♂♂; Luebo; 05°20’ S, 21°24’ E; Jan.-Apr. 1959; F. François leg.; RMCA. – **Kasaï Central** • 1 ♂; Luluabourg (Kananga); 05°54’ S, 21°52’ E; Jul. 1930; R.P. Vanderyst leg.; RMCA^3^ • 1 ♂; same location as for preceding; 1939; R.P. Vankerckhoven leg.; RMCA. – **Kongo Central** • 1 ♂; Kisantu (Madimba); 05°08’ S, 15°06’ E; 1927; R. P. Vanderyst leg.; RMCA • 1 ♂; same location as for preceding; Dec. 1927; R. P. Vanderyst leg.; RMCA • 1 ♂; Lemfu (Madimba); 05°18’ S, 15°13’ E; 1931; Rév. P. Van Eyen leg.; RMCA • 1 ♀; Mayidi (Madimba); 05°11’ S, 15°09’ E; 1942; Rév. P. Van Eyen leg.; RMCA • 2 ♂♂; same location as for preceding; Dec. 1942; Rév. P. Van Eyen leg.; RMCA. – **Lomami** • 1 ♂; Kabinda; 06°08’ S, 24°29’ E; 1935; Mme. Gillardin leg.; RMCA • 1 ♂; Lomami: Kambaye; 06°53’ S, 23°44’ E; Jul. 1930; P. Quarré leg.; RMCA. – **Sud-Kivu** • 1 ♂; Kalembelembe-Baraka; 04°32’ S, 28°47’ E; Jul. 1918; R. Mayné leg.; RMCA.

#### *Megachile* (*Creightonella*) *ianthoptera* Smith, 1853, *Creightonella ianthoptera* (Smith, 1853)

**Material.** Cupboard 110, Box 48 (184♀♀ and 97♂♂). **D.R. CONGO**. – **Bas-Uele** • 1 ♂; Uele: Tukpwo; 04°26’ N, 25°51’ E; Sep. 1937; J. Vrydagh leg.; RMCA • 1 ♀; same location as for preceding; Jul. 1937; J. Vrydagh leg.; RMCA. –**Equateur**• 1 ♂; Bamania (Mbandaka); 00°01’ N, 18°19’ E; 1934; Dr. J. Schouteden leg.; RMCA • 1 ♀; Coquilhatville (Mbandaka); 00°03’ N, 18°15’ E; 1946; Ch. Scops leg.; RMCA • 1 ♀; same location as for preceding; 16 Jun. 1924; Dr. M. Bequaert leg.; RMCA • 3 ♀♀; Eala (Mbandaka); 00°03’ N, 18°19’ E; 1932; A. Corbisier leg.; RMCA • 4 ♂♂; same location as for preceding; 1932; A. Corbisier leg.; RMCA • 1 ♀; same location as for preceding; G. Couteaux leg.; RMCA •1 ♀; same location as for preceding; 1933; A. Corbisier leg.; RMCA •1 ♂; same location as for preceding; 1933; A. Corbisier leg.; RMCA •1 ♂; same location as for preceding; 1933; H.J. Brédo leg.; RMCA • 8 ♀♀; same location as for preceding; 14 Mar.1931; A. Corbisier leg.; RMCA • 1 ♂; same location as for preceding; 14 Mar.1933; A. Corbisier leg.; RMCA • 1 ♂; same location as for preceding; 14 Nov.1931; H.J. Brédo leg.; RMCA • 1 ♀; same location as for preceding; 15 Nov.1931; H.J. Brédo leg.; RMCA • 2 ♀♀; same location as for preceding; 2 Oct. 1931; H.J. Brédo leg.; RMCA • 1 ♂; same location as for preceding; 2 Oct. 1931; H.J. Brédo leg.; RMCA • 2 ♀♀; same location as for preceding; 21 Nov. 1931; H.J. Brédo leg.; RMCA • 1 ♂; same location as for preceding; 21 Nov. 1931; H.J. Brédo leg.; RMCA • 2 ♀♀; same location as for preceding; 22 Jul. 1914; R. Mayné leg.; RMCA • 1 ♂; same location as for preceding; 22 Jul. 1914; R. Mayné leg.; RMCA • 1 ♀; same location as for preceding; 22 Nov. 1931; H.J. Brédo leg.; RMCA • 1 ♂; same location as for preceding; 29 Aug. 1914; R. Mayné leg.; RMCA • 1 ♀; same location as for preceding; 3 Oct. 1931; H.J. Brédo leg.; RMCA • 1 ♀; same location as for preceding; 30 Sep. 1939; G. Couteaux leg.; RMCA • 1 ♀; same location as for preceding; 30 Nov. 1939; G. Couteaux leg.; RMCA • 1 ♂; same location as for preceding; 4 Apr. 1932; H.J. Brédo leg.; RMCA • 1 ♀; same location as for preceding; 7 Jul. 1932; A. Corbisier leg.; RMCA • 1 ♂; same location as for preceding; 7 Jul. 1932; A. Corbisier leg.; RMCA • 7 ♀♀; same location as for preceding; Mar. 1932; H.J. Brédo leg.; RMCA • 1 ♀; same location as for preceding; Mar. 1933; H.J. Brédo leg.; RMCA • 3 ♀♀; same location as for preceding; Mar. 1935; J. Ghesquière leg.; RMCA • 3 ♂♂; same location as for preceding; Mar. 1935; J. Ghesquière leg.;

RMCA • 12 ♀♀; same location as for preceding; Apr. 1933; A. Corbisier leg.; RMCA • 4 ♂♂; same location as for preceding; Apr. 1933; A. Corbisier leg.; RMCA • 1 ♀; same location as for preceding; Sep. 1912; R. Mayné leg.; RMCA • 1 ♂; same location as for preceding; Sep. 1912; R. Mayné leg.; RMCA • 4 ♀♀; same location as for preceding; Sep. 1930; H.J. Brédo leg.; RMCA • 2 ♀♀; same location as for preceding; May. 1932; H.J. Brédo leg.; RMCA • 2 ♀♀; same location as for preceding; May. 1935; J. Ghesquière leg.; RMCA • 38 ♀♀; same location as for preceding; Jun. 1932; A. Corbisier leg.; RMCA • 8 ♂♂; same location as for preceding; Jun. 1932; A. Corbisier leg.; RMCA • 1 ♀; same location as for preceding; Jun. 1935; J. Ghesquière leg.; RMCA • 1 ♂; same location as for preceding; Oct. 1929; H.J. Brédo leg.; RMCA • 1 ♀; same location as for preceding; Oct. 1931; H.J. Brédo leg.; RMCA • 2 ♀♀; same location as for preceding; Nov. 1931; H.J. Brédo leg.; RMCA • 1 ♂; same location as for preceding; Nov. 1931; H.J. Brédo leg.; RMCA • 3 ♀♀; same location as for preceding; Nov. 1932; H.J. Brédo leg.; RMCA • 1 ♀; same location as for preceding; Nov. 1934; J. Ghesquière leg.; RMCA • 1 ♂; same location as for preceding; Nov. 1934; J. Ghesquière leg.; RMCA • 3 ♀♀; Equateur; 00°00’ N, 18°14’ E; Verlaine leg.; RMCA • 1 ♀; Flandria (Ingende); 00°20’ S, 19°06’ E; 15 Mar. 1932 R.P. Hulstaert leg.; RMCA • 1 ♀; same location as for preceding; Jan.-Feb. 1928; R.P. Hulstaert leg.; RMCA • 1 ♀; Ikenge; 00°06’ S, 18°46’ E; 18 Sep. 1912; R. Mayné leg.; RMCA. – **Haut-Katanga** • 1 ♀; Elisabethville (Lubumbashi); 11°40’ S, 27°28’ E; 15 Nov. 1957; Ch. Seydel leg.; RMCA • 1 ♂; same location as for preceding; De Loose leg.; RMCA • 1 ♀; Elisabethville; riv. Kimilolo; 11°40’ S, 27°28’ E; 2 Apr. 1923; Dr. M. Bequaert leg.; RMCA • 1 ♀; Lubumbashi; 11°40’ S, 27°28’ E; 2 May 1921; Dr. M. Bequaert leg.; RMCA • 1 ♀; PNU Kabwe sur Muye; 08°47’ S, 26°52’ E, alt. 1320 m; 11 May 1948; Mis. G.F. de Witte leg.; RMCA • 1 ♀; PNU Lusinga; 08°55’ S, 27°12’ E, alt. 1760 m; 12-17 Dec. 1947; Mis. G.F. de Witte leg.; RMCA • 1 ♂; PNU Lusinga (riv. Kamitungulu); 08°55’ S, 08°55’ S, 13 Jun. 1945; Mis. G.F. de Witte leg.; RMCA • 1 ♀; Ruwe-Kambove; 10°40’ S, 25°33’ E; 3 Apr. 1907; Dr. Sheffield Neave leg.; RMCA. – **Haut-Lomami** • 1 ♀; Lomami-Kaniama; 07°31’ S, 24°11’ E; 1932; R. Massart leg.; RMCA • 1 ♂; same location as for preceding; 1932; R. Massart leg.; RMCA • 1 ♀; Lualaba: Kabongo; 07°20’ S, 25°35’ E; 31 Dec. 1952; Ch. Seydel leg.;

RMCA • 1 ♀; same location as for preceding; 5 Jan. 1953; Ch. Seydel leg.; RMCA • 1 ♀; same location as for preceding; Nov. 1953; Ch. Seydel leg.; RMCA. – **Haut-Uele** • 1 ♀; Haut-Uele: Abimva; 03°44’ N, 29°42’ E; 19-22 Jun. 1925; Dr. H. Schouteden leg.; RMCA • 1 ♀; Haut-Uele: Moto; 03°03’ N, 29°28’ E; 1923; L. Burgeon leg.; RMCA. – **Ituri** • 1 ♀; Ituri: Bunia; 01°34’ N, 30°15’ E; 1938; P. Lefèvre leg.; RMCA • 1 ♀; same location as for preceding; 23 Feb. 1934; Ch. Scops leg.; RMCA • 1 ♂; same location as for preceding; 23 Feb. 1934; Ch. Scops leg.; RMCA • 1 ♂; same location as for preceding; Feb. 1934; Ch. Scops leg.; RMCA • 1 ♂; same location as for preceding; Jul. 1934; Ch. Scops leg.; RMCA • 1 ♀; Ituri: Mahagi; 02°18’ N, 30°59’ E; 1931; Dr. H. Schouteden leg.; RMCA • 1 ♀; same location as for preceding; 25 May 1925; Dr. H. Schouteden leg.; RMCA • 1 ♀; Kibali-Ituri: Nioka; 02°10’ N, 30°40’ E; 16-28 Aug. 1931; H.J. Brédo leg.; RMCA • 1 ♀; Kilo: Kere-Kere; 02°42’ N, 30°33’ E; Jan. 1935; Dr. Turco leg.; RMCA • 4 ♂♂; same location as for preceding; Jan. 1935; Dr. Turco leg.; RMCA • 1 ♂; Lac Albert: Kasenyi; 01°24’ N, 30°26’ E; 15 May 1935; H.J. Brédo leg.; RMCA • 1 ♂; Mahagi-Niarembe; 02°15’ N, 31°07’ E; 1935; Ch. Scops leg.; RMCA • 1 ♂; same location as for preceding; Nov. 1935; Dr. J. Schouteden leg.; RMCA • 1 ♂; Région d’Abok; 02°00’ N, 31°00’ E; Oct. 1935; Ch. Scops leg.; RMCA. – **Kongo Central** • 1 ♀; Kisantu (Madimba); 05°08’ S, 15°06’ E; 1927; R.P. Vanderyst leg.; RMCA • 4 ♀♀; same location as for preceding; 1927; R.P. Vanderyst leg.; RMCA • 1 ♀; same location as for preceding; 1931; R.P. Vanderyst leg.; RMCA • 1 ♂; same location as for preceding; 1931; R.P. Vanderyst leg.; RMCA • 3 ♀♀; same location as for preceding; Dec. 1927; R.P. Vanderyst leg.; RMCA • 2 ♂♂; same location as for preceding; Dec. 1927; R.P. Vanderyst leg.; RMCA • 1 ♀; Lemfu (Madimba); 05°18’ S, 15°13’ E; May 1945; Rév. P. De Beir leg.; RMCA • 2 ♀♀; same location as for preceding; May 1945; Rév. P. De Beir leg.; RMCA • 3 ♀♀; Mayidi (Madimba); 05°11’ S, 15°09’ E; 1942; Rév. P. Van Eyen leg.; RMCA • 4 ♂♂; same location as for preceding; 1942; Rév. P. Van Eyen leg.; RMCA • 2 ♂♂; same location as for preceding; 1945; Rév. P. Van Eyen leg.; RMCA • 1 ♀; Sunba (Boma); 05°11’ S, 15°00’ E; Dec. 1907; Dr. Sheffield Neave leg.; RMCA. – **Lomami** • 1 ♂; Gandajika (Station INEAC 167); 06°45’ S, 23°57’ E; 1957; G. Schmitz leg.; RMCA • 1 ♂; Lomami: Kabwe; 08°47’ S, 26°52’ E; Jul. – Aug. 1931; P. Quarré leg.; RMCA. – **Lualaba** • 1 ♀; Katentenia; 10°19’ S, 25°54’ E; May 1923; Ch. Seydel leg.; RMCA • 1 ♀; Lulua: Kanzenze; 10°31’ S, 25°12’ E; May 1932; F.G. Overlaet leg.; RMCA • 5 ♀♀; Lulua: Kapanga; 08°21’ S, 22°34’ E; Oct. 1932; F.G. Overlaet leg.; RMCA • 3 ♀♀; same location as for preceding; Dec. 1933; F.G. Overlaet leg.; RMCA • 1 ♂; same location as for preceding; Dec. 1933; F.G. Overlaet leg.; RMCA • 1 ♀; Sandu; 09°41’ S, 22°53’ E; Apr. 1931; H.J. Brédo leg.; RMCA • 1 ♀; Tenke; 10°35’ S, 26°06’ E; Mar. 1925; Ch. Seydel leg.; RMCA. – **Maï-Ndombe** • 1 ♂; Kwamouth (Mushie); 03°11’ S, 16°12’ E; 13 Dec. 1926; Dr. M. Bequaert leg.; RMCA • 1 ♀; Lac Léopold II (Lac Maï-Ndombe); 02°00’ S, 18°15’ E; 1925; R.P. Vankerckhoven leg.; RMCA • 1 ♀; Mushie; 03°11’ S, 16°12’ E; Jan. – Mar. 1915; R. Mayné leg.;

RMCA. – **Maniema** • 1 ♀; Manyema; 05°00’ S, 29°00’ E; R. Mayné leg.; RMCA • 1 ♀; Nyangwe; 04°13’ S, 26°11’ E; Apr. 1918; R. Mayné leg.; RMCA • 1 ♀; same location as for preceding; Apr.-May 1918; R. Mayné leg.; RMCA • 2 ♂♂; same location as for preceding; Apr.-May 1918; R. Mayné leg.; RMCA. – **Nord-Kivu** • 1 ♀; Lac Vert; 01°41’ S, 29°14’ E; 16 Sep. 1951; A.E. Bertrand leg.; RMCA • 1 ♀; PNA [Parc National Albert]; 00°03’ S, 29°30’ E; 10 Oct. 1951; P. Vanschuytbroeck & H. Synave leg.; RMCA • 1 ♀; same location as for preceding; 9-12 Jul. 1955; P. Vanschuytbroeck & H. Synave leg.; RMCA • 1 ♀; Ruwenzori; 00°20’ N, 29°50’ E, alt. 1000 m; 6 Dec. 1931; Mme. L. Lebrun leg.; RMCA • 1 ♀; Ruwenzori: Vall. Butagu; 00°20’ N, 29°50’ E, alt. 2000 m; 3 Dec. 1931; Mme. L. Lebrun leg.; RMCA • 1 ♀; Tshibinda; 02°19’ S, 28°45’ E; 26 Nov. 1932; T.D. Cockerell leg.; RMCA. – **Nord-Ubangi** • 2 ♀♀; Yakoma; 04°06’ N, 22°23’ E; 5 – 17 Feb. 1932; H.J. Brédo leg.; RMCA. – **Sankuru** • 1 ♂; Sankuru: Komi; 03°23’ S, 23°46’ E; 27 Jan. 1930; J. Ghesquière leg.; RMCA • 1 ♂; Sankuru: Lodja; 03°29’ S, 23°26’ E; 17 Feb. 1930; J. Ghesquière leg.; RMCA. – **Sud-Kivu** • 1 ♂; Kalembelembe-Baraka; 04°32’ S, 28°47’ E; Jul. 1918; R. Mayné leg.; RMCA • 1 ♀; Mulungu: Tshibinda; 02°19’ S, 28°45’ E; 19 Sep. 1938; Hendrickx leg.; RMCA • 2 ♀♀; same location as for preceding; Nov. 1951; P.C. Lefèvre leg.; RMCA • 4 ♂♂; same location as for preceding; Nov. 1951; P.C. Lefèvre leg.; RMCA • 1 ♂; Région des lacs, 02°29’ S, 28°51’ E; Dr. Sagona leg.; RMCA. – **Sud-Ubangi** • 1 ♀; Kunungu; 02°45’ N, 18°26’ E; 1937; Dr. H. Schouteden leg.; RMCA • 1 ♂; same location as for preceding; 1937; Dr. H. Schouteden leg.; RMCA. – **Tanganyika** • 1 ♀; Kabalo; 06°03’ S, 26°55’ E; Jan. 1929; Ch. Seydel leg.; RMCA • 1 ♀; Kiambi (Manono); 07°19’ S, 28°01’ E; 15 Mar. 1911; Dr. Valdonio leg.; RMCA • 1 ♀; Lukuga r. Niemba; 05°57’ S, 28°26’ E; Sep. 1917-Jan. 1918; Dr. Pons leg.; RMCA • 1 ♂; Tanganyika; 06°20’ S, 29°00’ E; 31 Aug. 1927; H. Bomans leg.; RMCA • 1 ♀; Tanganyika: Moba; 07°02’ S, 29°47’ E; Mar. 1954; H. Bomans leg.; RMCA. – **Tshopo** • 1 ♀; Isangi: Région du Lomami; 00°47’ N, 24°16’ E; Oct. 1905; H. Wilmin leg.; RMCA • 1 ♀; Ponthierville (Ubundu); 00°22’ S, 25°29’ E; 22 Oct. 1910; Dr. J. Bequaert leg.; RMCA • 1 ♂; Stan. Yangambi; 00°46’ N, 24°27’ E; 14-15 Dec. 1952; P. Basilewsky leg.; RMCA • 1 ♀; Stanleyville (Kisangani); 00°31’ N, 25°11’ E; 10 – 13 Sep. 1928; A. Collart leg.; RMCA • 1 ♂; same location as for preceding; Feb. 1948; Dr. R. Mouchamps leg.; RMCA • 1 ♀; Yangambi; 00°46’ N, 24°27’ E; Dec. 1939; Dr. Parent leg.;

RMCA. – **Tshuapa** • 2 ♂♂; Bokuma; 00°06’ S, 18°41’ E; Jul. 1952; Rév. P. Lootens leg.; RMCA • 1 ♂; Mondombe (Ikela); 00°54’ S, 22°49’ E; Oct. 1912; R. Mayné leg.; RMCA • 2 ♀♀; Tshuapa: Bokuma; 00°06’ S, 18°41’ E; 1949; Dupuis leg.; RMCA • 2 ♂♂; same location as for preceding, Jun. 1952; Rév. P. Lootens leg.; RMCA • 1 ♀; same location as for preceding, Jul. 1952; Rév. P. Lootens leg.; RMCA • 1 ♀; Tshuapa: Bokungu; 00°41’ S, 22°19’ E; 1949; Dupuis leg.; RMCA • 7 ♂♂; same location as for preceding; 1949; Dupuis leg.; RMCA • 1 ♂; same location as for preceding; 1950; M. Dupuis leg.; RMCA.

#### *Megachile* (*Creightonella*) *ikuthaensis* Friese, 1903, *Creightonella ikuthaënsis* (Friese, 1903)

**Material.** Cupboard 110, Box 50 (1♂)**. D.R. CONGO**. – **Haut-Uele** • 1 ♂ ; PNG [Parc National de Garamba]; 03°40’ N, 29°00’ E; 21 Apr. 1950; H. De Saeger leg.; RMCA.

#### *Megachile* (*Creightonella*) *paracantha* (Pasteels, 1965), *Creightonella paracantha* Pasteels, 1965

**Material.** Cupboard 110, Box 46 (1♂)**. D.R. CONGO**. Holotype – **Haut-Katanga** • 1 ♂; Katanga: Mwema; 08°13’ S, 27°28’ E; Aug. 1927; A. Bayet leg.; RMCA.

#### *Megachile* (*Creightonella*) *ruda* (Pasteels, 1965), *Creightonella ruda* Pasteels, 1965

**Material.** Cupboard 110, Box 46 (1♂)**. D.R. CONGO**. Holotype – **Bas-Uele** • 1 ♀; Mobwasa; 02°40’ N, 23°11’ E; Sep. 1911; De Giorgi leg.; RMCA.

#### *Megachile* (*Creightonella*) *rufa* Friese, 1903, *Creightonella rufa* (Friese, 1903)

**Material.** Cupboard 110, Box 50 (72♀♀ and 36♂♂)**. D.R. CONGO**. – **Bas-Uele** • 1 ♀; Bambesa; 03°28’ N, 25°43’ E; 1938; P. Henrard leg.; RMCA • 1 ♂; same location as for preceding; 11 May 1938; P. Henrard leg.; RMCA • 1 ♂; same location as for preceding; 14 May 1938; P. Henrard leg.; RMCA • 2 ♀♀; same location as for preceding; Feb. 1934; H.J. Brédo leg.; RMCA. – **Haut-Katanga** • 1 ♀; Elisabethville (Lubumbashi); 11°40’ S, 27°28’ E; 1928-1929; P. Quarré leg.; RMCA • 1 ♀; same location as for preceding; Mar. 1930; Dr. M. Bequaert leg.; RMCA • 1 ♀; same location as for preceding; Apr. 1933; De Loose leg.; RMCA • 1 ♀; same location as for preceding; Jun. 1932; De Loose leg.; RMCA • 1 ♂; Elisabethville (Lubumbashi); 11°40’ S, 27°28’ E; 16 Jun. 1953; Ch. Seydel leg.; RMCA • 1 ♀; same location as for preceding; 5 – 9 Oct. 1951; Ch. Seydel leg.; RMCA • 1 ♀; Kundelungu; 10°40’ S, 28°00’ E; Jun. 1949; A. Zielinski leg.; RMCA • 1 ♀; Lukafu; 10°31’ S, 27°33’ E; 6-22 Dec. 1930; Dr. M. Bequaert leg.; RMCA • 1 ♀; PNU Kabwe sur Muye; 08°47’ S, 26°52’ E, alt. 1320 m; 11 May 1948; Mis. G.F. de Witte leg.; RMCA • 1 ♀; PNU Kilwezi affl. dr. Lufira; 09°05’ S, 26°45’ E, alt. 750 m; 16 – 21 Mar. 1948; Mis. G.F. de Witte leg.; RMCA • 1 ♀; PNU Lusinga (riv. Kamitungulu); 08°55’ S, 08°55’ S; 13 Jun. 1945; Mis. G.F. de Witte leg.; RMCA. – **Haut-Lomami** • 2 ♀♀; Lualaba: Kabongo; 07°20’ S, 25°35’ E; 27 Dec. 1952; Ch. Seydel leg.; RMCA • 1 ♀; same location as for preceding; 3 Jan. 1953; F.G. Overlaet leg.; RMCA • 1 ♀; same location as for preceding; 31 Dec. 1952; Ch. Seydel leg.; RMCA • 1 ♀; Lulua: Luashi; 07°31’ S, 24°10’ E; 1935; Freyne leg.; RMCA • 2 ♀♀; same location as for preceding; 1936; Freyne leg.; RMCA. – **Haut-Uele** • 1 ♀; PNG [Parc National de Garamba]; 03°40’ N, 29°00’ E; 23 May 1951; H. De Saeger leg.; RMCA • 1 ♀; same location as for preceding; 24 May 1951; H. De Saeger leg.; RMCA • 1 ♀; same location as for preceding; 5 May 1950; H. De Saeger leg.; RMCA • 2 ♀♀; PNG Napokomweli; 03°40’ N, 29°00’ E; 18 Oct. 1950; H. De Saeger leg.; RMCA – **Ituri** • 1 ♀; Kibali-Ituri: Mahagi; 02°18’ N, 30°59’ E; 1932; Ch. Scops leg.; RMCA. – **Kasaï Central** • 1 ♂; Luluabourg (Kananga); 05°54’ S, 21°52’ E; 15 Apr. 1939; J.J. Deheyn leg.; RMCA. – **Kinshasa**• 1 ♂; Congo belge; 04°19’ S, 15°19’ E; Carpentier leg.;

RMCA. – **Kongo Central** • 1 ♂; Boma; 05°51’ S, 13°03’ E; 28 Mar. 1913; Lt. Styczynski leg.; RMCA • 1 ♂; Kisantu (Madimba); 05°08’ S, 15°06’ E; 1921; P. Van Wing leg.; RMCA • 3 ♀♀; same location as for preceding; 1927; R. P. Vanderyst leg.; RMCA • 5 ♂♂; same location as for preceding; 1927; R. P. Vanderyst leg.; RMCA • 1 ♂; same location as for preceding; 1932; R. P. Vanderyst leg.; RMCA • 1 ♂; same location as for preceding; Jan. – Apr. 1930; R. P. Vanderyst leg.; RMCA • 1 ♂; same location as for preceding; Dec. 1927; R. P. Vanderyst leg.; RMCA • 2 ♂♂; Lemfu (Madimba); 05°18’ S, 15°13’ E; 1945; Rév. P. De Beir leg.; RMCA • 1 ♂; same location as for preceding; Jan. 1945; Rév. P. De Beir leg.; RMCA • 2 ♂♂; same location as for preceding; May 1945; Rév. P. De Beir leg.; RMCA • 1 ♂; same location as for preceding; Oct. – Dec. 1944; Rév. P. De Beir leg.; RMCA • 2 ♀♀; Mayidi (Madimba); 05°11’ S, 15°09’ E; 1942; 1942; Rév. P. Van Eyen leg.; RMCA • 3 ♂♂; same location as for preceding; 1942; Rév. P. Van Eyen leg.; RMCA. – **Lomami** • 1 ♀; Gandajika; 06°45’ S, 23°57’ E; 22 Nov. 1950; P. de Francquen leg.; RMCA • 1 ♀; same location as for preceding; 24 Nov. 1950; P. de Francquen leg.; RMCA • 1 ♀; Kanda Kanda; 06°56’ S, 23°37’ E; 1935; J. Drion leg.; RMCA • 1 ♀; Lomami-Luputa; 11°00’ S, 26°44’ E; May 1935; Dr. Bouvier leg.; RMCA • 2 ♀♀; Sankuru: Mwene-Ditu; 07°00’ S, 23°27’E; 25 Nov. 1952; R. Maréchal leg.; RMCA. – **Lualaba** • 1 ♂; Biano; 10°20’ S, 27°00’ E; Apr. 1927; R. Mayné leg.; RMCA – **Lualaba** • 1 ♂; Kapanga; 08°21’ S, 22°34’ E; Aug. 1934; F.G. Overlaet leg.; RMCA • 3 ♀♀; same location as for preceding; Oct. 1933; F.G. Overlaet leg.; RMCA • 1 ♀; same location as for preceding; Nov. 1933; F.G. Overlaet leg.; RMCA • 1 ♀; Kayembe Mukulu; 09°03’ S, 23°57’ E; 29 Nov. 1911; Dr. J. Bequaert leg.; RMCA • 1 ♂; Lulua: Kapanga; 08°21’ S, 22°34’ E; Sep. 1932; F.G. Overlaet leg.; RMCA • 3 ♀♀; same location as for preceding; Oct. 1932; F.G. Overlaet leg.; RMCA • 3 ♂♂; same location as for preceding; Oct. 1932; F.G. Overlaet leg.; RMCA • 4 ♀♀; same location as for preceding; Nov. 1932; F.G. Overlaet leg.; RMCA • 2 ♀♀; same location as for preceding; Dec. 1932; F.G. Overlaet leg.; RMCA • 1 ♀; Lulua: r. Kapelekese; 08°21’S, 22°34’ E; 16 Nov. 1933; F.G. Overlaet leg.; RMCA. – **Maniema** • 1 ♂; Kibombo; 03°54’ S, 25°55’ E; 18 Jan. 1910; Dr. J. Bequaert leg.; RMCA • 1 ♂; same location as for preceding; Oct. 1930; H.J. Brédo leg.; RMCA • 1 ♂; Malela; 04°22’ S, 26°08’ E; Jan. 1948; L. Burgeon leg.; RMCA • 1 ♂; Nyangwe; 04°13’ S, 26°11’ E; Apr. – May 1918; R. Mayné leg.; RMCA. – **Nord-Kivu** • 1 ♂; PNA: S.L. Edouard Kitembo; 00°20’ S, 29°30’ E; 4 Apr. 1936; L. Lippens leg.; RMCA. – **Sankuru** • 1 ♀; Lusambo: Sangale; 04°58’ S, 23°26’ E; Nov. 1934; Mme. Gillardin leg.; RMCA. – **Sud-Kivu** • 1 ♀; Rég. Des Grands Lacs; 02°29’ S, 28°51’ E; Thys leg.; RMCA. – **Tanganyika** • 1 ♂; Shausele N. Kabemba; 05°57’ S, 29°12’ E; Mar. 1939; Mevr. Bequaert leg.; RMCA.

#### *Megachile* (*Creightonella*) *rufoscopacaea* Friese, 1903, *Creightonella rufoscopacaea* (Friese, 1903)

**Material.** Cupboard 110, Box 51 (42♀♀)**. D.R. CONGO**. – **Haut-Katanga** • 2 ♀♀; Elisabethville (Lubumbashi); 11°40’ S, 27°28’ E; 26 Sep. 1952; Ch. Seydel leg.; RMCA • 1 ♀; same location as for preceding; 29 Apr. 1920; Dr. M. Bequaert leg.; RMCA • 1 ♀; same location as for preceding; Mar. 1924; Dr. M. Bequaert leg.; RMCA • 1 ♀; same location as for preceding; De Loose leg.; RMCA • 1 ♀; Kambove-Bunkeya; 10°52’ S, 26°38’ E; Oct. 1907; Dr. Sheffield Neave leg.; RMCA • 1 ♀; Kapema-Kipaila; 10°42’ S, 28°40’ E; Oct. 1907; Dr. Sheffield Neave leg.; RMCA • 1 ♀; Kipopo; 11°33’ S, 27°21’ E; 11 Sep. 1961; R. Maréchal leg.; RMCA • 1 ♀; same location as for preceding; 30 Nov. 1961; R. Maréchal leg.; RMCA • 1 ♀; Lubumbashi; 11°40’ S, 27°28’ E; 10 May 1953; Ch. Seydel leg.; RMCA • 2 ♀♀; same location as for preceding; 5 – 9 Oct. 1951; Ch. Seydel leg.; RMCA • 1 ♀; Mwema; 08°13’ S, 27°28’ E; 1956; R.P.Th. de Caters leg.; RMCA • 1 ♀; PNU Lusinga (riv. Kamitungulu); 08°55’ S, 08°55’ S; 13 May 1945; Mis. G.F. de Witte leg.; RMCA • 1 ♀; PNU Masombwe s. Grde. Kafwe; 09°05’ S, 27°12’ E, alt. 1120 m; 16 Apr. 1948; Mis. G.F. de Witte leg.; RMCA. – **Haut-Lomami** • 1 ♀; Bukama; 09°12’ S, 25°51’ E; 11 May 1911; Dr. J. Bequaert leg.; RMCA • 1 ♀; same location as for preceding; 16 Apr. 1911; Dr. J. Bequaert leg.; RMCA • 1 ♀; same location as for preceding; 24 May 1911; Dr. J. Bequaert leg.; RMCA • 1 ♀; same location as for preceding; 26 Apr. 1911; Dr. J. Bequaert leg.; RMCA • 1 ♀; Kadiamapanga; 08°16’ S, 26°36’ E; Nov. 1931; H.J. Brédo leg.; RMCA • 1 ♀; Luashi; 07°31’ S, 24°10’ E; 1935; Freyne leg.; RMCA • 1 ♀; PNU Munoi, bifure Lupiala; 08°50’ S, 26°44’E, alt. 890 m; 25 May-15 Jun. 1948; Mis. G.F. de Witte leg.; RMCA. – **Ituri** • 1 ♀; Ituri: Blukwa; 01°45’ N, 30°36’ E; 3-4 Dec. 1928; A. Collart leg.; RMCA. – **Lualaba** • 1 ♀; Bunkeya-Lukafu; 10°24’ S, 26°58’ E; Oct. 1927; Dr. Sheffield Neave leg.; RMCA • 1 ♀; Kando; 10°42’ S, 26°23’ E; Apr. 1931; G.F. de Witte leg.; RMCA • 1 ♀; Kayambo-Dikuluwe; 10°45’ S, 25°22’ E; Jun. 1907; Dr. Sheffield Neave leg.; RMCA • 1 ♀; Lulua: Kafakumba (Sandoa); 09°41’ S, 23°44’ E; Dec. 1932; F.G. Overlaet leg.; RMCA • 1 ♀; Lulua: Kanzenze; 10°31’ S, 25°12’ E; Dec. 1932; F.G. Overlaet leg.; RMCA • 1 ♀; Tenke; 10°35’ S, 26°06’ E; 30 Jul.-9Aug. 1931; T.D. Cockerell leg.; RMCA. – **Maniema** • 1 ♀; Kakinga; 03°27’ S, 26°32’ E; Nov. 1931; T.D. Cockerell leg.; RMCA. – **Nord-Kivu** • 1 ♀; Tshibinda; 02°19’ S, 28°45’ E; Nov. 1927; Ch. Seydel leg.; RMCA. – **Sud-Kivu** • 1 ♀; Bobandana; 01°42’ S, 29°01’ E; 1928; O. Douce leg.; RMCA. • 1 ♀; Kahuzi; 02°15’ S, 28°41’ E; 25 Sep. 1938; Dr. J. Schouteden leg.; RMCA. – **Tanganyika** • 1 ♀; Kanikiri; 00°09’ S, 29°32’ E; Oct. 1907; Dr. Sheffield Neave leg.; RMCA • 1 ♀; Lukuga r. Niemba; 05°57’ S, 28°26’ E; Nov. 1917-Jan. 1918; Dr. Pons leg.; RMCA • 2 ♀♀; Lukulu (Manono); 07°08’ S, 28°06’ E; 1 – 3 May 1931; G.F. de Witte leg.; RMCA • 2 ♀♀; Tanganyika; 05°57’ S, 29°12’ E; Hecq leg.; RMCA • 1 ♀; Tanganyika-Moero: Nyunzu; 05°57’ S, 28°01’ E; Jan. – Feb. 1934; H. De Saeger leg.; RMCA. – **Tshopo** • 1 ♀; Mfungwe-Kayumbe; 01°17’ S, 26°16’ E; Jun. 1907; Dr. Sheffield Neave leg.; RMCA.

#### *Megachile* (*Creightonella*) *sternintegra* (Pasteels, 1965), *Creightonella sternintegra* Pasteels, 1965

**Material.** Cupboard 110, Box 50 (8♀♀and 2♂♂)**. D.R. CONGO**. Holotype – **Haut-Katanga** • 1 ♀; Kapema; 10°42’ S, 28°40’ E; Nov. 1924; Ch. Seydel leg.; RMCA. Paratypes – **Haut-Katanga** • 2 ♀♀; PNU Kankunda (riv. dr. Lupiala); 08°56’ S, 27°12’ E, alt. 1300 m; 13-27 Nov. 1947; Mis. G.F. de Witte leg.; RMCA • 5 ♀♀; PNU Lusinga (riv. Kamitungulu); 08°55’ S, 08°55’ S; 13 Jun. 1945; G.F. de Witte leg.; RMCA. – **Lualaba** • 1 ♂; Kafakumba (Sandoa); 09°41’ S, 23°44’ E; Dec. 1932; F.G. Overlaet leg.; RMCA. – **Tanganyika** • 1 ♂; Shausele N. Kabemba; 05°57’ S, 29°12’ E; Mar. 1939; Mevr. Bequaert leg.; RMCA.

#### *Megachile* (*Creightonella*) *trichroma* Friese, 1922, *Creightonella trichroma* Friese, 1922

**Material.** Cupboard 110, Box 46 (1♀)**. D.R. CONGO**. – **Kasaï Central** • 1 ♀; Lula, Terr. Luiza; 07°11’ S, 22°24’ E; Aug. 1956; A.J. Jobaert leg.; RMCA.

#### *Megachile (Digitella) digiticauda* Cockerell, 1937

**Material.** Cupboard 110, Box 11 (5♂♂)**. D.R. CONGO**. Paratype – **Lualaba** • 1 ♂; Biano; 10°20’ S, 27°00’ E; 8 – 11 Aug. 1931; T.D. Cockerell leg.; RMCA.

– **Haut-Katanga** • 2 ♂♂; Elisabethville (Lubumbashi); 11°40’ S, 27°28’ E; 23 Sep. 1923; Dr. M. Bequaert leg.; RMCA • 1 ♂; same location as for preceding; 1928 – 1933; De Loose leg.; RMCA • 1 ♂; Lubombo; 08°10’ S, 29°45’ E; Aug. 1928; Ch. Seydel leg.; RMCA.

#### *Megachile (Eurymella) akamiella* Pasteels, 1965

**Material.** Cupboard 110, Box 7 (1♀)**. D.R. CONGO**. Holotype – **Haut-Uele** • 1 ♀; PNG Akam; 03°40’ N, 29°00’ E; 3 May 1950; H. De Saeger leg.; RMCA.

#### *Megachile (Eurymella) atroalbida* Pasteels, 1965

**Material.** Cupboard 110, Box 3 (1♀)**. D.R. CONGO**. Holotype – **Sud-Kivu** • 1 ♀; Uvira; 03°25’ S, 29°08’ E; Sep.1958, J. Pasteels leg.; RMCA.

#### *Megachile (Eurymella) aurifera* Cockerell, 1935

**Material.** Cupboard 110, Box 6 (2♀♀)**. D.R. CONGO**. – **Bas-Uele** • 1 ♀; Uele: Bambesa; 03°28’ N, 25°43’ E; Jan. 1934; J.V. Leroy leg.; RMCA. – **Equateur** • 1 ♀; Eala (Mbandaka); 00°03’ N, 18°19’ E; May 1932; H.J. Brédo leg.; RMCA.

#### *Megachile (Eurymella) basalis* Smith, 1853

**Material.** Cupboard 110, Box 1 (1♀ and 1♂)**. D.R. CONGO**. – **Haut-Lomami** • 1 ♀; Lovoi; 08°50’ S, 25°00’ E; 18 Oct. 1911; Dr. J. Bequaert leg.; RMCA • 1 ♂; PNU Mabwe (r. E. lac Upemba); 08°47’ S, 26°52’ E, alt. 585 m; 17-20. Nov.1948; G.F. de Witte leg.; RMCA.

#### Megachile (Eurymella) bredoi Cockerell, 1935

**Material.** Cupboard 110, Box 4 (4 ♀♀)**. D.R. CONGO**. Holotype – **Haut-Katanga** • 1 ♀; Muelushi; 08°59’ S, 26°45’ E; Nov. 1931; H.J. Brédo leg.; RMCA. Paratype – **Haut-Katanga** • 3 ♀♀; Muelushi; 08°59’ S, 26°45’ E; Nov. 1931; H.J. Brédo leg.; RMCA.

#### *Megachile (Eurymella) brochidens* Vachal, 1903

**Material.** Cupboard 110, Box 7 (3♀♀)**. D.R. CONGO**. – **Equateur** • 1 ♀; Coquilhatville (Mbandaka); 00°03’ N, 18°15’ E; 14 Oct. 1922; Dr. M. Bequaert leg.; RMCA. – **Haut-Lomami** • 1 ♀; Bukama; 09°12’ S, 25°51’ E; Apr. 1911; Dr. J. Bequaert leg.; RMCA. – **Haut-Lomami**• 1 ♀; Bukama; 09°12’ S, 25°51’ E; 18 Feb. 1911; Dr. J. Bequaert leg.; RMCA.

#### *Megachile (Eurymella) bucephala* Fabricius, 1793

**Material.** Cupboard 110, Box 4 (28 ♀♀ and 24 ♂♂)**. D.R. CONGO**. – **Equateur** • 1 ♂; Coquilhatville (Mbandaka); 00°03’ N, 18°19’ E; 10 Nov. 1931; Lt. Dorman leg.; RMCA • 1 ♂; Eala (Mbandaka); 00°03’ N, 18°19’ E; 1933; A. Corbisier leg.; RMCA • 1 ♂; same location as for preceding; 19 Oct. 1931; H.J. Brédo leg.; RMCA • 3 ♂♂; same location as for preceding; 20 Oct. 1931; H.J. Brédo leg.; RMCA • 1 ♀; same location as for preceding; 5 Nov. 1931; H.J. Brédo leg.; RMCA • 1 ♀; same location as for preceding; May 1935; J. Ghesquière leg.; RMCA • 1 ♀; same location as for preceding; Jun. 1932; A. Corbisier leg.; RMCA • 1 ♂; same location as for preceding; Apr. 1935; J. Ghesquière leg.; RMCA • 1 ♀; same location as for preceding; Jun. 1935; J. Ghesquière leg.; RMCA • 2 ♀♀; same location as for preceding; Nov. 1931; H.J. Brédo leg.; RMCA • 1 ♀; same location as for preceding; Nov. 1932; A. Corbisier leg.; RMCA • 1 ♀; Eala-Bokatola-Bikoro; 00°03’ N, 18°19’ E; Sep. – Oct. 1930; Dr. P. Staner leg.; RMCA • 1 ♀; Ubangi: Nouvelle Anvers (Bomongo); 00°03’ N, 18°19’ E; 9 Dec. 1952; P. Basilewsky leg.; RMCA. – **Haut-Katanga** • 1 ♀; Kalunkumia; 10°49’ S, 26°51’ E; 3 Apr. 1925; Ch. Seydel leg.; RMCA • 1 ♀; Mwema; 08°13’ S, 27°28’ E; Jul. 1927; A. Bayet leg.; RMCA • 1 ♀; Point E; 11°40’ S, 27°28’ E; 18 Feb. 1911; Dr. J. Bequaert leg.; RMCA. – **Haut-Lomami** • 1 ♂; Kalele; 02°07’ S, 28°55’ E; Feb. 1931; H.J. Brédo leg.; RMCA • 1 ♂; Kalele; 02°07’ S, 28°55’ E; Feb. 1931; H.J. Brédo leg.; RMCA • 1 ♀; Lualaba: Kaniama; 07°31’ S, 24°11’ E; 19 Dec. 1952; Ch. Seydel leg.; RMCA • 1 ♂; same location as for preceding; 6 Apr. 1949; Mis. G.F. de Witte leg.; RMCA • 1 ♂; same location as for preceding; Jan. – Feb. 1949; Mis. G.F. de Witte leg.; RMCA. – **Haut-Uele** • 1 ♂; PNG [Parc National de Garamba]; 03°40’ N, 29°00’ E; Nov. 1950; H. De Saeger leg.; RMCA. – **Kongo Central** • 1 ♀; Banana; 06°00’ S, 12°24’ E; Aug. 1910; Dr. Etienne leg.; RMCA • 1 ♀; Boma; 05°51’ S, 13°03’ E; 28 Mar. 1913; Lt. Styczynski leg.; RMCA • 1 ♂; Congo da Lemba (Songololo); 05°42’ S, 13°42’ E; Jan. – Feb. 1913; R. Mayné leg.; RMCA • 1 ♀; Lemfu (Madimba); 05°18’ S, 15°13’ E; Jun. 1945; Rév. P. De Beir leg.; RMCA • 1 ♀; Mayidi (Madimba); 05°11’ S, 15°09’ E; 1942; Rév. P. Van Eyen leg.; RMCA • 1 ♂; same location as for preceding; 1942; Rév. P. Van Eyen leg.;

RMCA. – **Maniema** • 1 ♀; K.300 de Kindu; 02°57’ S, 25°55’ E; 9 Apr. 1911; L. Burgeon leg.; RMCA • 1 ♀; Kibombo; 03°54’ S, 25°55’ E; 6 Nov. 1910; Dr. J. Bequaert leg.; RMCA • 1 ♀; Terr. de Kasongo, Riv. Lumami; 04°27’ S, 26°40’ E; 8 Nov. 1960; P.L.G. Benoit leg.; RMCA • 1 ♀; Vieux Kassongo (Kasongo); 04°27’ S, 26°40’ E; 22 Dec. 1910; Dr. J. Bequaert leg.; RMCA. – **Sud-Kivu** • 1 ♂; Kalembelembe-Baraka; 04°32’ S, 28°47’ E; Jul. 1918; R. Mayné leg.; RMCA. – **Tanganyika** • 1 ♀; Albertville (Kalemie); 05°57’ S, 29°12’ E; 1 – 20 Jan. 1919; R. Mayné leg.; RMCA • 1 ♀; same location as for preceding; Dec. 1918; R. Mayné leg.; RMCA • 1 ♀; Bassin Lukuga; 05°40’ S, 26°55’ E; Apr. – Jul. 1934; H. De Saeger leg.; RMCA • 1 ♂; Kongolo; 05°24’ S, 05°40’ S, 26°55’ E; 25 Feb. 1911; Dr. J. Bequaert leg.; RMCA • 1 ♂; same location as for preceding; 6 Feb. 1911; Miss. H. De Saeger leg.; RMCA. – **Tshuapa** • 2 ♀♀; Bokuma; 00°06’ S, 18°41’ E; 00°06’ S, 18°41’ E; Jun. 1952; Rév. P. Lootens leg.; RMCA • 1 ♂; same location as for preceding; Jul. 1952; Rév. P. Lootens leg.; RMCA • 1 ♂; Boma; 05°51’ S, 13°03’ E; Dec. 1951; Rév. P. Lootens leg.; RMCA • 1 ♂; Tshuapa: Bokuma; 00°06’ S, 18°41’ E; Mar. 1954; Rév. P. Lootens leg.; RMCA • 3 ♀♀; same location as for preceding; Jun. 1952; Rév. P. Lootens leg.; RMCA • 2 ♂♂; same location as for preceding; Jun. 1952; Rév. P. Lootens leg.; RMCA • 1 ♀; same location as for preceding; Jul. 1952; Rév. P. Lootens leg.; RMCA.

#### *Megachile (Eurymella) caricina* Cockerell, 1907

**Material.** Cupboard 110, Box 6 (8♀♀ and 3♂♂)**. D.R. CONGO**. – **Haut-Katanga** • 1 ♂; Elisabethville (Lubumbashi); 11°40’ S, 27°28’ E; 1924; Dr. M. Bequaert leg.; RMCA • 2 ♀♀; PNU Kaziba; 09°09’ S, 26°57’ E, alt. 1140 m; 11 – 15 Feb. 1948; G.F. de Witte leg.; RMCA • 1 ♀; PNU Masombwe s. Grde; Kafwe; 09°05’ S, 27°12’ E, alt. 1120 m; 16 Apr. 1948; G.F. de Witte leg.; RMCA. – **Haut-Lomami** • 2♀♀; Kadiamapanga; 08°16’ S, 26°36’ E; Nov. 1931; H.J. Brédo leg.; RMCA. – **Haut-Uele** • 1 ♀; Haut-Uele: Abimva; 03°44’ N, 29°42’ E; 1925; L. Burgeon leg.; RMCA. – **Ituri** • 1 ♀; Ituri: Bunia; 01°34’ N, 30°15’ E; Jun. 1938; P. Lefèvre leg.; RMCA. – **Lualaba** • 1 ♂; Haut-Luapula: Kansenia; 10°19’ S, 26°04’ E; 14 Oct. 1929; Dom de Montpellier leg.; RMCA. – **Nord-Kivu** • 1 ♂; PNA SL Edouard: Kamande; 00°20’ S, 29°30’ E, alt. 925 m; 4 Apr. 1935; L. Lippens leg.; RMCA. – **Tanganyika** • 1 ♂; Vallée Lukuga; 05°40’ S, 26°55’ E; Nov. 1911; Dr. Schwetz leg.; RMCA.

#### *Megachile (Eurymella) crassitarsis* Cockerell, 1920

**Material.** Cupboard 110, Box 6 (1♀ and 7♂♂)**. D.R. CONGO**. – **Haut-Katanga** • 1 ♂; Kiamanwa; 09°30’ S, 27°57’ E; Feb. 1931; H.J. Brédo leg.; RMCA • 1 ♂; PNU Lusinga (riv. Kamitungulu); 08°55’ S, 08°55’ S, 13 Jun. 1945; G.F. de Witte leg.; RMCA • 2 ♂♂; PNU Masombwe s. Grde; Kafwe; 09°05’ S, 27°12’ E, alt. 1120 m; 16 Apr. 1948; G.F. de Witte leg.; RMCA • 1 ♀; Lubumbashi; 11°40’ S, 27°28’ E; 8 Mar. 1921; Dr. M. Bequaert leg.; RMCA • 1 ♂; same location as for preceding; 8 Mar. 1921; Dr. M. Bequaert leg.; RMCA • 1 ♂; Wabishamba; 11°40’ S, 27°28’ E; 3 Dec. 1926; Ch. Seydel leg.; RMCA. – **Haut-Lomami** • 1 ♂; Sankisia; 09°24’ S, 25°48’ E; 4 Sep. 1911; Dr. J. Bequaert leg.; RMCA.

#### *Megachile (Eurymella) dolichognatha* Cockerell, 1931

**Material.** Cupboard 110, Box 4 (2♀♀)**. D.R. CONGO**. Holotype – **Nord-Kivu** • 1 ♀; Ruwenzori; 00°20’ N, 29°50’ E, alt. 1900 m; 23 May 1915; Miss. H. De Saeger leg.; RMCA.

– **Haut-Katanga** • 1 ♀; Lusinga; 08°56’ S, 27°12’ E; 9 Apr. 1917; G.F. de Witte leg.; RMCA.

#### *Megachile (Eurymella) eurymera* Smith, 1854

**Material.** Cupboard 110, Box 1 (39♀♀ and 9♂♂)**. D.R. CONGO**. – **Bas-Uele** •1 ♀; Bambesa; 03°28’ N, 25°43’ E; 11 May 1938; P. Henrard leg.; RMCA • 1 ♀; same location as for preceding; 14 May 1938; P. Henrard leg.; RMCA. – **Equateur** • 1 ♀; Bokote; 00°06’ S, 20°08’ E; 6 Mar. 1926; R.P. Hulstaert leg.; RMCA • 2 ♀♀; Eala (Mbandaka); 00°03’ N, 18°19’ E; 1932, A. Corbisier leg.; RMCA • 2 ♀♀; same location as for preceding; Jun. 1932, A. Corbisier leg.; RMCA • 3 ♀♀; same location as for preceding; 4 Apr. 1932; H.J. Brédo leg.; RMCA • 7 ♀♀; same location as for preceding; Mar. 1932; H.J. Brédo leg.; RMCA. – **Haut-Katanga** • 1 ♀; Kilwa; 09°18’ S, 28°25’ E; Apr. 1931; H.J. Brédo leg.; RMCA • 1 ♀; PNU Kaswabilenga; 08°51’ S, 26°43’ E, alt. 700 m; 21 Oct. 1947; G.F. De Witte leg.; RMCA. – **Haut-Lomami** • 1 ♀; Lualaba: Kabongo; 07°20’ S, 25°35’ E; 3 Jan. 1953; Ch. Seydel leg.; RMCA • 2 ♀♀; PNU Mabwe (R.P. Lac Upemba); 08°47’ S, 26°52’ E, alt. 585 m; 17-20 Nov. 1948; G.F. De Witte leg.; RMCA • 5 ♂♂; same location as for preceding; 4 Apr. 1932; G.F. De Witte leg.; RMCA • 2 ♀♀; Sankisia; 09°24’ S, 25°48’ E; 1 Sep. 1911; Dr. J. Bequaert leg.; RMCA. – **Haut-Uele** • 1 ♀; PNG [Parc National de Garamba]; 03°40’ N, 29°00’ E; 16 Aug. 1951; H. De Saeger leg.; RMCA. – **Ituri** • 1 ♀; Lac Albert: Kasenyi; 01°24’ N, 30°26’ E; 1935; H.J. Brédo leg.; RMCA • 1 ♂; same location as for preceding; 1935; H.J. Brédo leg.; RMCA • 1♂; Mahagi-Niarembe; 02°15’ N, 31°07’ E; 1935; Ch. Scops leg.; RMCA • 1 ♀; Penge; 01°20’ N, 28°09’ E; Jan. 1926; Ch. Seydel leg.; RMCA. – **Kinshasa** • 1 ♀; Léopoldville (Kinshasa); 04°19’ S, 15°19’ E; 25 Sep. 1925; Dr. M. Bequaert leg.; RMCA. – **Kongo Central** • 1 ♀; Kalina; 04°18’ S, 15°17’ E; Jul. 1945; Mme. Delsaut leg.; RMCA • 1 ♀; Kisantu (Madimba); 05°08’ S, 15°06’ E; Rév. P. Regnier leg.; RMCA • 2 ♀♀; Lemfu (Madimba); 05°18’ S, 15°13’ E; May 1945; Rév. P. De Beir leg.; RMCA • 2 ♀♀; Mayidi (Madimba); 05°11’ S, 15°09’ E; 1942; Rév. P. Van Eyen leg.; RMCA • 1 ♀; same location as for preceding; 1945; Rév. P. Van Eyen leg.; RMCA. – **Lualaba**• 1 ♂; Sandu; 09°41’ S, 22°53’ E; Apr. 1931; H.J. Brédo leg.; RMCA. – **Maï-Ndombe** • 1 ♀; Bokolaka (Bolobo); 02°09’ S, 16°14’ E; 1954, R.C. Eloy leg.; RMCA. –**Maniema** • 1 ♂; Nyangwe; 04°13’ S, 26°11’ E; Apr. – May 1918; R. Mayné leg.; RMCA.

#### Megachile (Eurymella) flavopilosa Pasteels, 1965

**Material.** Cupboard 110, Box 3 (4♂♂)**. D.R. CONGO**. Holotype – **Haut-Uele** • 1 ♂; PNG [Parc National de Garamba]; 03°40’ N, 29°00’ E; 23 Aug. 1951; H. De Saeger leg.; RMCA. <u>Paratypes</u> DRC – **Haut-Uele** • 3 ♂♂; PNG [Parc National de Garamba]; 03°40’ N, 29°00’ E; 27 Aug. 1951; H. De Saeger leg.; RMCA.

#### *Megachile (Eurymella) garambana* Pasteels, 1965

**Material.** Cupboard 110, Box 6 (2♀♀and 3♂♂)**. D.R. CONGO**. Holotype – **Haut-Uele** • 1 ♂; PNG Akam; 03°40’ N, 29°00’ E; 21 Apr. 1950; H. De Saeger leg.; RMCA. Allotype – **Haut-Uele** • 1 ♀; PNG Nagero (Dungu); 03°45’ N, 29°31’ E; 1 – 23 Apr. 1954; C. Nebay leg.; RMCA. <u>Paratypes</u> DRC – **Haut-Uele** • 2 ♂♂; PNG Akam; 03°40’ N, 29°00’ E; 21 Apr. 1950; H. De Saeger leg.; RMCA • 1 ♀; PNG Mt. Mboyo; 03°40’ N, 29°00’ E; 25 Sep. 1952; H. De Saeger leg.; RMCA.

#### *Megachile* (*Eurymella) kimilolana* Cockerell, 1931 *Megachile* (*Eurymella*) *kimilonana* Cockerell, 1931

**Material.** Cupboard 110, Box 6 (12♂♂)**. D.R. CONGO**. Holotype – **Haut-Katanga** • 1 ♂; Elisabethville; riv. Kimilolo; 11°40’ S, 27°28’ E; 6 Nov. 1920; Dr. M. Bequaert leg.; RMCA.

– **Haut-Katanga** • 1 ♂; Elisabethville (Lubumbashi); 11°40’ S, 27°28’ E; 11 – 17 Sep. 1931; T.D. Cockerell leg.; RMCA • 1 ♂; Kapema; 10°42’ S, 28°40’ E; Sep. 1924; Ch. Seydel leg.; RMCA • 1 ♂; PNU Kabwe sur Muye; 08°47’ S, 26°52’ E, alt. 1320 m; 11 May 1948; G.F. de Witte leg.; RMCA • 3 ♂♂; PNU Masombwe s. Grde; Kafwe; 09°05’ S, 27°12’ E, alt. 1120 m; 16 Apr. 1948; G.F. de Witte leg.; RMCA. – **Haut-Lomami** • 1 ♂; Kalongwe; 11°01’ S, 25°13’ E; 15 Sep. 1911; Dr. J. Bequaert leg.; RMCA • 2 ♂♂; Sankisia; 09°24’ S, 25°48’ E; 4 Sep. 1911; Dr. J. Bequaert leg.; RMCA – **Lualaba** • 1 ♂; Haut-Luapula: Kansenia; 10°19’ S, 26°04’ E; 18 Oct. 1929; Dom de Montpellier leg.; RMCA • 1 ♂; same location as for preceding; Nov. 1929; Dom de Montpellier leg.; RMCA.

#### *Megachile (Eurymella) konowiana* Friese, 1903

**Material.** Cupboard 110, Box 7 (6♂♂)**. D.R. CONGO**. – **Haut-Katanga** • 6 ♂♂; PNU Lusinga, Rivière Kamitungulu; 08°55’ S, 27°12’ E; 13 Jun. 1945; G.F. de Witte leg.; RMCA.

#### *Megachile (Eurymella) michaelis* Cockerell, 1931

**Material.** Cupboard 110, Box 4 (4♀♀)**. D.R. CONGO**. Holotype – **Haut-Katanga** • 1 ♀; Lubumbashi; 11°40’ S, 27°28’ E; 8 Aug. 1921; Dr. M. Bequaert leg.; RMCA.

– **Haut-Katanga** • 1 ♀; Elisabethville (Lubumbashi); 11°40’ S, 27°28’ E; Jun. 1932; De Loose leg.; RMCA.

– **Haut-Uele** • 1 ♀; PNG [Parc National de Garamba]; 03°40’ N, 29°00’ E; 21 Apr. 1950; H. De Saeger leg.; RMCA. – **Sud-Kivu** • 1 ♀; Mulungu: Tshibinda; 02°19’ S, 28°45’ E; Nov. 1951; Miss. H. De Saeger leg.; RMCA.

#### Megachile (Eurymella) nigripollex Vachal, 1910

**Material.** Cupboard 110, Box 2 (51♀♀and 139♂♂)**. D.R. CONGO**. Holotype – **Lualaba** • 1 ♂; Bunkeya; 10°24’ S, 26°58’ E; Oct. 1907; Dr. Sheffield-Neave leg.; RMCA.

– **Equateur** • 1 ♂; Beneden-Congo: Tumba; 00°50’ S, 18°00’ E; 20 Sep. 1938; Mevr. Bequaert leg.; RMCA • 2 ♀♀; Bokote; 00°06’ S, 20°08’ E; 5 Mar. 1926; R.P. Hulstaert leg.; RMCA • 1 ♀; Cité de Kombo (Ingende); 00°05’ S, 18°37’ E; 21 Mar. 1939; Mevr. Bequaert leg.; RMCA • 5 ♂♂; Eala (Mbandaka); 00°03’ N, 18°19’ E; Apr. 1933; A. Corbisier leg.; RMCA • 5 ♂♂; same location as for preceding; Jun. 1932; A. Corbisier leg.; RMCA • 1 ♂; same location as for preceding; Nov. 1932; A. Corbisier leg.; RMCA • 1 ♀; same location as for preceding; 20 Oct. 1931; H.J. Brédo leg.; RMCA • 1 ♀; same location as for preceding; 4 Apr. 1932; H.J. Brédo leg.; RMCA • 1 ♀; same location as for preceding; Mar. 1932; H.J. Brédo leg.; RMCA • 2 ♂♂; same location as for preceding; Mar. 1932; H.J. Brédo leg.; RMCA • 1 ♀; same location as for preceding; Nov. 1931; H.J. Brédo leg.; RMCA • 1 ♂; same location as for preceding; Nov. 1931; H.J. Brédo leg.; RMCA • 1 ♀; Coquilhatville (Mbandaka); 00°03’ N, 18°15’ E; 14 Oct. 1922; Dr. M. Bequaert leg.; RMCA • 1 ♂; same location as for preceding; 25 Nov. 1924; Dr. M. Bequaert leg.; RMCA. • 2 ♂♂; Ubangi: Nouvelle Anvers (Bomongo); 00°03’ N, 18°19’ E; 9 Dec. 1952; P. Basilewsky leg.; RMCA. – **Haut-Katanga** • 1 ♂; Elisabethville (Lubumbashi); 11°40’ S, 27°28’ E; 1-6 Sep. 1932; De Loose leg.; RMCA • 1 ♂; same location as for preceding; De Loose leg.; RMCA • 2 ♂♂; Kapema; 10°42’ S, 28°40’ E; Sep. 1924; Ch. Seydel leg.; RMCA • 1 ♀; Lubumbashi; 11°40’ S, 27°28’ E; 10 Apr. 1921; Dr. M. Bequaert leg.; RMCA. – **Haut-Lomami**• 1 ♂; Bukama; 09°12’ S, 25°51’ E; Aug. 1923; Ch. Seydel leg.; RMCA • 1♀; Kikondja; 08°11’ S, 26°26’ E; 28 Nov. 1911; Dr. J. Bequaert leg.; RMCA • 1 ♂; Lualaba: Kabongo; 07°20’ S, 25°35’ E; 27 Dec. 1952; Ch. Seydel leg.; RMCA • 2 ♂♂; same location as for preceding; 3 Jan. 1953; Ch. Seydel leg.; RMCA • 2 ♂♂; same location as for preceding; 30 Dec. 1952; Ch. Seydel leg.; RMCA • 4 ♂♂; same location as for preceding; 7 Jan. 1953; Ch. Seydel leg.; RMCA. – **Haut-Uele** • 1 ♂; Mayumbe: Tshela; 04°59’ S, 12°56’ E; Jun. 1925; A. Collart leg.; RMCA • 2 ♂♂; PNG [Parc National de Garamba]; 03°40’ N, 29°00’ E; 21 Apr. 1950; Mis. H. De Saeger leg.; RMCA • 1 ♀; Uele: Gangala-na-Bodio; 03°41’ N, 29°08’ E; Oct. 1956; Dr. M. Poll leg.; RMCA. – **Ituri** • 1 ♂; Kilo: Kere-Kere; 02°42’ N, 30°33’ E; Jan. 1935; Dr. Turco leg.; RMCA. – **Kasaï** • 1 ♀; Luebo rivière; 05°20’ S, 21°24’ E; 25 May 1938; Mevr. Bequaert leg.; RMCA • 1 ♂; same location as for preceding; 25 May 1938; Mevr. Bequaert leg.; RMCA. – **Kasaï Central** • 1 ♀; Lula (Kasaï); 07°11’ S, 22°24’ E; 1958; A.J. Jobaert leg.; RMCA • 1♂; same location as for preceding; 1958; A.J. Jobaert leg.; RMCA • 1 ♀; Lula: Terr. Luiza; 07°11’ S, 22°24’ E; Aug. 1958; Dr. M. Poll leg.; RMCA • 3 ♀♀; Luluabourg (Kananga); 05°54’ S, 21°52’ E; P. Callewaert leg.; RMCA • 1 ♂; same location as for preceding; P. Callewaert leg.; RMCA • 2 ♂♂; Luluabourg: Katoka; 05°54’ S, 21°52’ E; 1939; R.P.N. Vankerckhoven leg.; RMCA. – **Kinshasa** • 1 ♀; Kimwenza (Kinshasa); 04°28’ S, 15°17’ E; Sep. 1962; M.J. Deheegher leg.; RMCA • 1 ♂; Léopoldville (Kinshasa); 04°19’ S, 15°19’ E; 13 Mar. 1911; Dr. Mouchet leg.; RMCA • 1 ♂; same location as for preceding; 17 Dec. 1925; R.P. Hulstaert leg.; RMCA • 1 ♂; Léopold IX (Kinshasa); 04°19’ S, 15°19’ E; 1910; Dr. J. Bequaert leg.;

RMCA. – **Kongo Central** • 1 ♂; Camp de Lukula; 05°23’ S, 12°57’ E; 1911; Dr. Daniel leg.; RMCA • 1 ♀; Kisantu (Madimba); 05°08’ S, 15°06’ E; Apr. 1921; P. Van Wing leg.; RMCA • 4 ♀♀; same location as for preceding; 1927; R.P. Vanderyst leg.; RMCA • 15 ♂♂; same location as for preceding; 1927; R.P. Vanderyst leg.; RMCA • 1 ♂; same location as for preceding; 1928; R.P. Vanderyst leg.; RMCA • 1 ♀; same location as for preceding; 1931; R.P. Vanderyst leg.; RMCA • 2 ♂♂; same location as for preceding; 1931; R.P. Vanderyst leg.; RMCA • 1 ♀; same location as for preceding; 1 Apr. 1930; R.P. Vanderyst leg.; RMCA • 2 ♂♂; same location as for preceding; 1 Apr. 1930; R.P. Vanderyst leg.; RMCA • 1 ♀; same location as for preceding; Dec. 1927; R.P. Vanderyst leg.; RMCA • 3 ♂♂; same location as for preceding; Dec. 1927; R.P. Vanderyst leg.; RMCA • 1 ♂; same location as for preceding; Dec. 1928; R.P. Vanderyst leg.; RMCA • 1 ♂; same location as for preceding; Rév. P. Regnier leg.; RMCA • 1 ♀; Lemfu (Madimba); 05°18’ S, 15°13’ E; 13 Apr. 1905; R.P. Van Eyen leg.; RMCA • 1 ♀; same location as for preceding; Mar. 1945; Rév. P. De Beir leg.; RMCA • 1 ♂; same location as for preceding; Mar. 1945; Rév. P. De Beir leg.; RMCA • 2 ♂♂; same location as for preceding; Jun. 1945; Rév. P. De Beir leg.; RMCA • 1 ♂; same location as for preceding; Oct. – Dec. 1944; Rév. P. De Beir leg.; RMCA • 1 ♂; Mayidi (Madimba); 05°11’ S, 15°09’ E; 1942; 1945; R.P. Van Eyen leg.; RMCA • 1 ♀; same location as for preceding; 1942; Rév. P. Van Eyen leg.; RMCA • 1 ♂; same location as for preceding; 1942; Rév. P. Van Eyen leg.; RMCA • 1 ♂; Thysville (Mbanza Ngungu); 05°15’ S, 14°52’ E; 1 Dec. 1952; P. Basilewsky leg.; RMCA. – **Kwango** • 1 ♀; Kwango; 04°48’ S, 17°02’ E; 1925; P. Vanderijst leg.; RMCA • 1 ♀; Kwango: Atene; 05°23’S, 19°24’E; Charlier leg.; RMCA • 1 ♂; Kwango: Kahemba; 07°17’ S, 19°00’ E; 1 Mar. 1939; Mevr. Bequaert leg.; RMCA. – **Kwilu** • 1 ♂; Leverville (Bulungu); 04°50’ S, 18°44’ E; 1925; Mme. Tinant leg.; RMCA. – **Lomami** • 1 ♀; Sankuru: Gandajika; 06°45’ S, 23°57’ E; 1954; JP. de Francquen leg.; RMCA • 1 ♂; Sankuru: M’Pemba Zeo (Gandajika); 06°49’ S, 23°58’ E; 9 Nov. 1959; Don R. Maréchal leg.; RMCA. – **Lualaba** • 1 ♂; Dilolo; 10°41’ S, 22°21’ E; 9 oct. 1933; H. De Saeger leg.; RMCA • 1 ♂; Haut-Luapula: Kansenia; 10°19’ S, 26°04’ E; 17 oct. 1929; Dom de Montpellier leg.; RMCA • 5 ♂♂; Kapanga; 08°21’ S, 22°34’ E; Aug. 1934; G.F. Overlaet leg.; RMCA • 1 ♀; Lulua: Kapanga; 08°21’ S, 22°34’ E; Oct. 1932; G.F. Overlaet leg.; RMCA • 1 ♂; same location as for preceding; Nov. 1932; G.F. Overlaet leg.; RMCA • 1 ♂; same location as for preceding; Dec. 1932; G.F. Overlaet leg.; RMCA • 1 ♂; Lulua: Tshibalaka; 08°32’ S, 23°12’ E; Oct. 1933; G.F. Overlaet leg.; RMCA • 5 ♂♂; Sandu; 09°41’ S, 22°53’ E; Apr. 1931; H.J. Brédo leg.;

RMCA. – **Maï-Ndombe** • 1 ♀; Wombali (Mushie); 03°16’ S, 17°22’ E; Jul. 1913; P. Vanderijst leg.; RMCA. – **Maniema** • 1 ♂; Lualaba: Kindu; 02°57’ S, 25°55’ E; 13 Jul. 1947; Dr. M. Poll leg.; RMCA • 2 ♀♀; Nyangwe; 04°13’ S, 26°11’ E; Apr.-May 1918; R. Mayné leg.; RMCA • 3 ♂♂; same location as for preceding; Apr.-May 1918; R. Mayné leg.; RMCA • 1 ♂; Terr. De Kasongo, Riv. Lumami; 04°27’ S, 26°40’ E; Feb. 1960; P.L.G. Benoit leg.; RMCA • 1 ♂; same location as for preceding; Sep. 1959; P.L.G. Benoit leg.; RMCA. – **Mongala** • 2 ♂♂; Bumba; 02°11’ N, 22°32’ E; 1935; R.P. Lootens leg.; RMCA. – **Nord-Ubangi** • 1 ♂; Abumombazi (Mobayi); 03°34’ N, 22°03’ E; 18-26 Feb. 1932; H.J. Brédo leg.; RMCA • 1 ♂; Ubangi: Karawa (Businga); 03°21’ N, 20°18’ E; 1936; Rév. Wallin leg.; RMCA • 3 ♂♂; Yakoma; 04°06’ N, 22°23’ E; 5-17 Feb. 1932; H.J. Brédo leg.; RMCA. – **Sankuru** • 1 ♂; Kaolele; 05°21’ S, 24°02’ E; Mar. 1939; Mevr. Bequaert leg.; RMCA • 4 ♀♀; N’Kolé; 03°27’ S, 22°26’ E; May 1923; A. Pilette leg.; RMCA • 2 ♂♂; same location as for preceding; May 1923; A. Pilette leg.; RMCA • 1 ♀; Sankuru; 04°57’ S, 23°26’ E; Apr. 1930; J. Ghesquière leg.; RMCA • 1 ♂; Sankuru: Kondue; 04°58’ S, 23°16’ E; 1934; Puissant leg.; RMCA. – **Sud-Ubangi** • 1 ♂; Kunungu; 02°45’ N, 18°26’ E; 1941; N’Kele (Col. Sch.) leg.; RMCA • 1 ♂; Kunungu (Nkele); 02°45’ N, 18°26’ E; 17 Apr. 1905; Schouteden leg.; RMCA • 1 ♀; Libenge; 03°39’ N, 18°38’ E; 8 Dec. 1931; Schouteden leg.; RMCA • 1 ♂; M’Paka, Terr. Libenge; 03°39’ N, 18°38’ E; Dec. 1959; M. Pecheur leg.; RMCA. – **Tanganyika** • 1 ♂; Katompe (Kabalo); 06°11’ S, 26°20’ E; Feb. 1929; Ch. Seydel leg.; RMCA • • 1 ♂; Kiambi (Manono); 07°19’ S, 28°01’ E; 22 – 24 Apr. 1931; Ch. Seydel leg.; RMCA • 1 ♂; same location as for preceding; 27 Apr. 1931; Ch. Seydel leg.; RMCA • 3 ♂♂; Kongolo; 05°24’ S, 27°00’ E; 23 Jan. 1911; Dr. J. Bequaert leg.; RMCA • 1 ♀; Tanganyika-Kabalo; 06°03’ S, 26°55’ E; 5 Jul. 1947; Dr. M. Poll leg.; RMCA • 1 ♂; Tanganyika-Moero: Nyunzu; 05°57’ S, 28°01’ E; Jan. – Feb. 1934; H. De Saeger leg.; RMCA. – **Tshopo** • 1 ♀; Lula; 00°27’ N, 25°12’ E; 1958; A.J. Jobaert leg.; RMCA. – **Tshuapa** • 1 ♀; Moma (Equateur); 00°45’ S, 21°56’ E; Jun. 1925; J. Ghesquière leg.; RMCA • 1 ♂; Tshuapa: Bokuma; 00°06’ S, 18°41’ E; 1953; R.P. Lootens leg.; RMCA • 1 ♂; same location as for preceding; Feb. 1952; R.P. Lootens leg.; RMCA • 4 ♂♂; same location as for preceding; Mar. 1952; R.P. Lootens leg.; RMCA • 1 ♂; same location as for preceding; Feb.-Mar. 1954; R.P. Lootens leg.; RMCA • 1 ♂; same location as for preceding; Jun. 1952; R.P. Lootens leg.; RMCA • 1 ♀; same location as for preceding; 00°06’ S, 18°41’ E; 1953; R.P. Lootens leg.; RMCA • 3 ♂♂; same location as for preceding; 1953; R.P. Lootens leg.; RMCA • 2 ♂♂; same location as for preceding; Jan.-Feb. 1952; R.P. Lootens leg.; RMCA • 2 ♀♀; same location as for preceding; Jul. 1952; R.P. Lootens leg.; RMCA • 1 ♂; same location as for preceding; Jul. 1952; R.P. Lootens leg.; RMCA.

#### *Megachile (Eurymella) opaculina* Cockerell, 1937

**Material.** Cupboard 110, Box 1 (1♂)**. D.R. CONGO**. – **Sud-Kivu** • 1 ♂; Uvira; 03°25’ S, 29°08’ E; Sep. 1958; Dr. J. Pasteels leg.; RMCA.

#### *Megachile (Eurymella) ornaticoxis* Cockerell, 1935

**Material.** Cupboard 110, Box 3 (3♂♂)**. D.R. CONGO**. Holotype – **Tshopo** • 1 ♂; Ponthierville (Ubundu); 00°22’ S, 25°29’ E; 22 Oct. 1910; Dr. J. Bequaert leg.; RMCA

– **Haut-Uele** • 1 ♂; PNG [Parc National de Garamba]; 03°40’ N, 29°00’ E; 9 Nov. 1950; H. De Saeger leg.; RMCA. – **Tshopo** • 1 ♂; Stanleyville (Kisangani); 00°31’ N, 25°11’ E; 25 Nov. 1928; A. Collart leg.; RMCA.

#### *Megachile (Eurymella) paupera* Pasteels, 1965

**Material.** Cupboard 110, Box 6 (3♂♂)**. D.R. CONGO**. Holotype – **Haut-Uele** • 1 ♂; PNG [Parc National de Garamba]; 03°40’ N, 29°00’ E; 9 Nov. 1950; H. De Saeger leg.; RMCA. Paratypes – **Maniema** • 1 ♂; Kibombo; 03°54’ S, 25°55’ E; 8 Nov. 1910; Dr. J. Bequaert leg.; RMCA • 1 ♂; Nyangwe; 04°13’ S, 26°11’ E; Apr. – May 1919; R. Mayné leg.; RMCA.

#### *Megachile (Eurymella) perfimbriata* Cockerell, 1920

**Material.** Cupboard 110, Box 6 (2♂♂)**. D.R. CONGO.** Holotype – **Haut-Katanga** • 1 ♂; Elisabethville, riv. Kimilolo; 11°40’ S, 27°28’ E; 6 Nov. 1920; Dr. M. Bequaert leg.; RMCA.

– **Haut-Katanga** • 1 ♂; PNU Masombwe s. Grde, Kafwe; 09°05’ S, 27°12’ E, alt. 1120 m; 16 Apr. 1948; G.F. de Witte leg.; RMCA.

#### *Megachile (Eurymella) planatipes* Cockerell, 1931

**Material.** Cupboard 110, Box 4 (2♀♀)**. D.R. CONGO**. Holotype – **Kasaï Central** • 1 ♀; Limbala; 06°07’ S, 22°20’ E; 5 – 8 Aug. 1913; Miss. H. De Saeger leg.; RMCA. Paratype – **Haut-Katanga** • 1 ♀; Elisabethville (Lubumbashi); 11°40’ S, 27°28’ E; 1928 – 1929; P. Quarré leg.; RMCA.

#### *Megachile (Eurymella) platystoma* Pasteels, 1965

**Material.** Cupboard 110, Box 8 (4♀♀and 1♂)**. D.R. CONGO**. Paratypes DRC – **Haut-Uele** • 1 ♀; PNG [Parc National de Garamba]; 03°40’ N, 29°00’ E 10 Aug. 1951; H. De Saeger leg.; RMCA • 2 ♀♀; same location as for preceding; 23 Aug. 1951; H. De Saeger leg.; RMCA • 1 ♂; same location as for preceding; 30 Jul. 1951; H. De Saeger leg.; RMCA.

– **Haut-Uele** • 1 ♀; PNG [Parc National de Garamba]; 03°40’ N, 29°00’ E; 23 Aug. 1951; H. De Saeger leg.; RMCA.

#### *Megachile (Eurymella) pyrrhothorax* Schletterer, 1891

The species *Megachile (Eurymella) pyrrothorax* was partitioned into several distinct forms by Pasteels (1965). The two following forms are represented in the RMCA collections in addition to the typical form, *burgeoni* Cockerell (dedicated to M. Burgeon) and *dentata* Friese.

#### *Megachile* (*Eurymella*) *pyrrhothorax* Schletterer, 1891, Cupboard 110, Box 3 (5♀♀ and 3♂♂)

**Material.** Cupboard 110, Box 3 (5♀♀ and 3♂♂)**. D.R. CONGO.** – **Bas-Uele** • 1 ♂; Bambesa; 03°28’ N, 25°43’ E; Feb. 1934; H.J. Brédo leg.; RMCA • 1 ♀; Uele: Bambesa; 03°28’ N, 25°43’ E; 30 Oct. 1933; J.V. Leroy leg.; RMCA. – **Equateur** • 1 ♂; Eala (Mbandaka); 00°03’ N, 18°19’ E; Dec. 1932; A. Corbisier leg.; RMCA • 1 ♀; same location as for preceding; 21 Nov. 1931; H.J. Brédo leg.; RMCA • 1 ♀; same location as for preceding; 3 Oct. 1931; H.J. Brédo leg.; RMCA • 1 ♀; same location as for preceding; Oct. 1935; J. Ghesquière leg.; RMCA. – **Haut-Lomami** • 1 ♀; PNU Kamitungulu; 08°55’ S, 27°12’ E, alt. 1700 m; 3 Apr. 1947; Mis. G.F. de Witte leg.; RMCA. – **Nord-Ubangi** • 1 ♂; Abumombazi (Mobayi); 03°34’ N, 22°03’ E; 18-26 Feb. 1932; H.J. Brédo leg.; RMCA.

#### *Megachile (Eurymella) pyrrhothorax, f. burgeoni* Cockerell, 1933

**Material.** Cupboard 110, Box 3 (4♀♀ and 3♂♂)**. D.R. CONGO.** Holotype – **Maniema** • 1 ♀; Malela; 04°22’ S, 26°08’ E; Dec. 1913; L. Burgeon leg.; RMCA.

– **Bas-Uele** • 1 ♀; Bambesa; 03°28’ N, 25°43’ E; Jan. 1934; H.J. Brédo leg.; RMCA. – **Equateur** • 1 ♂; Eala (Mbandaka); 00°03’ N, 18°19’ E; Mar. 1932; H.J. Brédo leg.; RMCA • 1 ♀; same location as for preceding; May 1932; H.J. Brédo leg.; RMCA • 1 ♀; same location as for preceding; Sep. 1935; J. Ghesquière leg.; RMCA • 1 ♂; same location as for preceding; Sep. 1935; J. Ghesquière leg.; RMCA • 1 ♂; Ubangi: Yakoma; 04°06’ N, 22°23’ E; 17 Feb. 1932; H.J. Brédo leg.; RMCA.

#### *Megachile (Eurymella) pyrrhothorax, f. dentata* Friese, 1909

**Material.** Cupboard 110, Box 3 (4♀♀ and 6♂♂)**. D.R. CONGO.** – **Bas-Uele** • 1 ♀; Bambesa; 03°28’ N, 25°43’ E; 1 Jun. 1934; H.J. Brédo leg.; RMCA. – **Equateur** • 1 ♂; Eala (Mbandaka); 00°03’ N, 18°19’ E; Apr. 1933, A. Corbisier leg.; RMCA • 1 ♂; same location as for preceding; 21 Nov. 1931; H.J. Brédo leg.; RMCA • 2 ♂♂; same location as for preceding; Sep. 1935; J. Ghesquière leg.; RMCA. – **Haut-Katanga** • 1 ♂; Elisabethville (Lubumbashi); 11°40’ S, 27°28’ E; Nov. 1926; Ch. Seydel leg.; RMCA • 1 ♂; La Panda (Likasi); 11°00’ S, 26°44’ E; 1 Oct. 1920; Dr. M. Bequaert leg.; RMCA. – **Kasaï Central** • 1 ♀; Luluabourg (Kananga); 05°54’ S, 21°52’ E; P. Callewaert leg.; RMCA. – **Lomami** • 1 ♀; Kabinda; 06°08’ S, 24°29’ E; Nov. 1926; Dr. Schwetz leg.; RMCA^4^. – **Tanganyika** • 1 ♀; Kampunda; 08°14’ S, 29°46’ E; 10 Nov. 1914; Dr. Mouchet leg.; RMCA.

#### *Megachile (Eurymella) riggenbachiana* Strand, 1911

**Material.** Cupboard 110, Box 6 (1♂)**. D.R. CONGO.** – **Haut-Lomami** • 1 ♂; Kulu-Mwanza; 07°54’ S, 26°45’ E; May 1927; A. Bayet leg.; RMCA.

#### *Megachile (Eurymella) salsburyana* Friese, 1922

**Material.** Cupboard 110, Box 6 (3♂♂)**. D.R. CONGO.** – **Haut-Katanga** • 1 ♂; PNU Lusinga, Rivière Kamitungulu; 08°55’ S, 27°12’ E; 13 Jun. 1945; G.F. de Witte leg.; RMCA • 1 ♂; Ruashi (Lubumbashi); 11°37’ S, 27°32’ E; 30 Jun. 1924; Ch. Seydel leg.; RMCA. – **Lualaba** • 1 ♂; Katentenia; 10°19’ S, 25°54’ E; May 1923; Ch. Seydel leg.; RMCA.

#### *Megachile (Eurymella) seclusa* Cockerell, 1931

**Material.** Cupboard 110, Box 3 (3♂♂)**. D.R. CONGO.** Holotype – **Haut-Katanga** • 1 ♂; Lubumbashi; 11°40’ S, 27°28’ E; 3 Feb. 1921; Dr. M. Bequaert leg.; RMCA. Paratype – **Haut-Katanga** • 1 ♂; Elisabethville (Lubumbashi); 11°40’ S, 27°28’ E; 29 Apr. 1920; Dr. M. Bequaert leg.; RMCA.

– **Lomami** • 1 ♂; Lomami: Luputa; 11°00’ S, 26°44’ E; 1935; Dr. Bouvier leg.; RMCA.

#### *Megachile (Eurymella) semierma* Vachal, 1903

**Material.** Cupboard 110, Box 5 (103♀♀ and 88♂♂)**. D.R. CONGO.** – **Bas-Uele** • 1 ♀; Bambesa; 03°28’ N, 25°43’ E; 11 May 1938; P. Henrard leg.; RMCA • 1 ♀; Uele: Tukpwo; 04°26’ N, 25°51’ E; Jul. 1937; J. Vrydagh leg.; RMCA. – **Equateur** • 1 ♀; Coquilhatville (Mbandaka); 00°03’ N, 18°15’ E; 15 Oct. 1922; Dr. M. Bequaert leg.; RMCA • 2 ♀♀; same location as for preceding; 15 Oct. 1922; Dr. M. Bequaert leg.; RMCA • 1 ♀; Eala (Mbandaka); 00°03’ N, 18°19’ E; 1921; H.J. Brédo leg.; RMCA • 2 ♂♂; same location as for preceding; 1932; A. Corbisier leg.; RMCA • 1 ♀; same location as for preceding; 1933; Dr. M. Bequaert leg.; RMCA • 1 ♀; same location as for preceding; 14 Mar. 1933; A. Corbisier leg.; RMCA • 1 ♂; same location as for preceding; 14 Mar. 1933; A. Corbisier leg.; RMCA • 1 ♂; same location as for preceding; 15 Oct. 1931; H.J. Brédo leg.; RMCA • 1 ♀; same location as for preceding; 17 Nov. 1931; H.J. Brédo leg.; RMCA • 1 ♂; same location as for preceding; 19 Oct. 1931; H.J. Brédo leg.; RMCA • 2 ♀♀; same location as for preceding; 21 Nov. 1931; H.J. Brédo leg.; RMCA • 1 ♂; same location as for preceding; 21 Nov. 1931; H.J. Brédo leg.; RMCA • 1 ♀; same location as for preceding; 28 Nov. 1931; H.J. Brédo leg.; RMCA • 1 ♂; same location as for preceding; 3 Oct. 1931; H.J. Brédo leg.; RMCA • 1 ♀; same location as for preceding; 4 Apr. 1932; H.J. Brédo leg.; RMCA • 1 ♂; same location as for preceding; 7 Nov. 1931; H.J. Brédo leg.; RMCA • 1 ♀; same location as for preceding; Mar. 1932; H.J. Brédo leg.; RMCA • 1 ♂; same location as for preceding; Mar. 1932; H.J. Brédo leg.; RMCA • 1 ♂; same location as for preceding; Apr. 1933; H.J. Brédo leg.; RMCA • 1 ♀; same location as for preceding; Apr. 1936; J. Ghesuière leg.; RMCA • 1 ♀; same location as for preceding; May 1933; A. Corbisier leg.; RMCA • 7 ♀♀; same location as for preceding; Jun. 1932; A. Corbisier leg.; RMCA • 2 ♂♂; same location as for preceding; Jun. 1932; A. Corbisier leg.; RMCA • 1 ♀; same location as for preceding; Jun. 1935; J. Ghesuière leg.; RMCA • 1 ♂; same location as for preceding; May 1932; H.J. Brédo leg.; RMCA • 1 ♂; same location as for preceding; Nov. 1931; H.J. Brédo leg.; RMCA • 2 ♂♂; same location as for preceding; Nov. 1932; H.J. Brédo leg.; RMCA • 1 ♀; same location as for preceding; Dec. 1932; A. Corbisier leg.; RMCA • 1 ♂; same location as for preceding; Dec. 1932; A. Corbisier leg.; RMCA • 1 ♀; same location as for preceding; Jun. 1935; J. Ghesquière leg.; RMCA • 14 ♀♀; Ubangi: Nouvelle Anvers (Bomongo); 00°03’ N, 18°19’ E; 19 Dec. 1952; P. Basilewsky leg.; RMCA • 9 ♂♂; same location as for preceding; 19 Dec. 1952; P. Basilewsky leg.;

RMCA. – **Haut-Katanga** • 1 ♀; Elisabethville (Lubumbashi); 11°40’ S, 27°28’ E; 3 Nov. 1933; Dr. M. Bequaert leg.; RMCA • 1 ♂; same location as for preceding; 4 May 1933; Dr. M. Bequaert leg.; RMCA • 1♂; same location as for preceding; De Loose leg.; RMCA • 1 ♂; same location as for preceding; Jul. 1928; Ch. Seydel leg.; RMCA • 1 ♂; same location as for preceding; Nov. 1938; H.J. Brédo leg.; RMCA • 1 ♀; Katanga: Geleka; 11°40’ S, 27°28’ E; 5 Apr. 1925; Ch. Seydel leg.; RMCA. – **Haut-Lomami** • 2 ♀♀; Kadiamapanga; 08°16’ S, 26°36’ E; Nov. 1931; H.J. Brédo leg.; RMCA • 1 ♀; Kapwasa; 07°33’ S, 26°27’ E; Nov. 1931; H.J. Brédo leg.; RMCA • 1 ♂; Lomami: Kaniama; 07°31’ S, 24°11’ E; 3 Jul. 1932; R. Massart leg.; RMCA • 1 ♂; Lualaba: Kabongo; 07°20’ S, 25°35’ E; 7 Jan. 1953; Ch. Seydel leg.; RMCA • 1 ♀; Lualaba: Kabongo; 07°20’ S, 25°35’ E; Mar. 1954; R. P. Th. De Caters leg.; RMCA. – **Haut-Uele** • 1 ♂; PNG [Parc National de Garamba]; 03°40’ N, 29°00’ E; 7 May 1952; H. De Saeger leg.; RMCA. – **Ituri** • 1♂; Ituri: Bunia; 01°34’ N, 30°15’ E; 1938; P. Lefèvre leg.; RMCA • 1 ♂; Kilo: Kere-Kere; 02°42’ N, 30°33’ E; Jan. 1935; Dr. Turco leg.; RMCA. – **Kasaï** • 1 ♀; Luebo rivière; 05°20’ S, 21°24’ E; 25 May 1938; Mevr. Bequaert leg.; RMCA. – **Kasaï Central** • 1 ♂; Luluabourg (Kananga); 05°54’ S, 21°52’ E; 18 Mar. 1919; P. Callewaert leg.; RMCA • 1 ♂; same location as for preceding; P. Callewaert leg.; RMCA. – **Kongo Central** • 1 ♀; Boma; 05°51’ S, 13°03’ E; 1937; Dr. Schlesser leg.; RMCA • 1 ♂; same location as for preceding; 20 Mar. 1913; Lt. Styczynski leg.; RMCA • 1 ♂; Congo da Lemba (Songololo); 05°42’ S, 13°42’ E; Apr. 1913; R. Mayné leg.; RMCA • 13 ♂♂; Kisantu (Madimba); 05°08’ S, 15°06’ E; 1927; R.P. Vanderyst leg.; RMCA • 2 ♀♀; same location as for preceding; Dec. 1927; R.P. Vanderyst leg.; RMCA • 6 ♂♂; same location as for preceding; Dec. 1927; R.P. Vanderyst leg.; RMCA • 3 ♀♀; Lemfu (Madimba); 05°18’ S, 15°13’ E; Jan. 1945; Rév. P. De Beir leg.; RMCA • 2 ♀♀; Mayidi (Madimba); 05°11’ S, 15°09’ E; 1942; Rév. P. Van Eyen leg.;

RMCA. – **Lomami** • 1 ♂; Lomami: Kambaye; 06°53’ S, 23°44’ E; Aug. 1930; P. Quarré leg.; RMCA. – **Lualaba** • 1 ♀; Dilolo; 10°41’ S, 22°21’ E; Aug. 1931; G.F. de Witte leg.; RMCA • 2 ♂♂; same location as for preceding; 1931; G.F. de Witte leg.; RMCA • 3 ♀♀; Ditanto; 10°15’ S, 25°53’ E; Oct. 1925; Ch. Seydel leg.; RMCA • 1 ♂; Funda Biabo; 09°50’ S, 25°33’ E; 15 – 18 Mar. 1914; L. Charliers leg.; RMCA • 2 ♂♂; Kapanga; 08°21’ S, 22°34’ E; Aug. 1934; F.G. Overlaet leg.; RMCA • 1 ♀; Lualaba: Kolwezi; 10°44’ S, 25°28’ E; 1 – 6 Nov. 1952; Mme. L. Gilbert leg.; RMCA • 1 ♀; Lulua: Kapanga; 08°21’ S, 22°34’ E; 8 Dec. 1932; F.G. Overlaet leg.; RMCA • 1 ♂; same location as for preceding; Sep. 1921; F.G. Overlaet leg.; RMCA • 1 ♀; same location as for preceding; Nov. 1932; F.G. Overlaet leg.; RMCA • 1 ♀; same location as for preceding; Dec. 1932; F.G. Overlaet leg.; RMCA • 1 ♀; Manika (Kolwezi); 10°20’ S, 27°00’ E; Oct. 1931; Ch. Seydel leg.; RMCA. – **Nord-Kivu** • 2 ♀♀; Beni; 00°29’ N, 29°28’ E; Aug. 1914; Dr. J. Bequaert leg.; RMCA • 1 ♀; same location as for preceding; Lt. Borgerhoff leg.; RMCA. – **Sankuru** • 1 ♀; Sankuru: Komi; 03°23’ S, 23°46’ E; Jul. 1930; J. Ghesquière leg.; RMCA. – **Sud-Kivu** • 1 ♀; Mulungu: Tshibinda; 02°19’ S, 28°45’ E; Nov. 1951; P.C. Lefèvre leg.; RMCA • 2 ♂♂; same location as for preceding; Nov. 1951; P.C. Lefèvre leg.; RMCA. – **Sud-Ubangi** • 1 ♂; Kunungu (Nkele); 02°45’ N, 18°26’ E; 1938; Dr. H. Schouteden leg.; RMCA • 1 ♂; Libenge; 03°39’ N, 18°38’ E; 10 Dec. 1931; H.J. Brédo leg.; RMCA. – **Tanganyika** • 1 ♂; Baudouinville (Moba); 07°02’ S, 29°47’ E; 19 Jan. 1933; L. Burgeon leg.; RMCA • 1 ♂; Kongolo; 05°24’ S, 27°00’ E; 6 Nov. 1911; Dr. J. Bequaert r leg.; RMCA • 1 ♀; Vallée Lukuga; 05°40’ S, 26°55’ E; Nov. 1911; Dr. Schwetz leg.;

RMCA. – **Tshopo** • 1 ♂; Stanleyville (Kisangani); 00°31’ N, 25°11’ E; 20 Oct. 1910; Dr. J. Bequaert leg.; RMCA. – **Tshuapa** • 1 ♂; Bamania (Mbandaka); 00°01’ N, 18°19’ E; 1934; R. Fr. Longinus leg.; RMCA • 2 ♀♀; Bokuma; 00°06’ S, 18°41’ E; 00°06’ S, 18°41’ E; 1951; Rév. P. Lootens leg.; RMCA • 11 ♀♀; same location as for preceding; 1953; Rév. P. Lootens leg.; RMCA • 4 ♂♂; same location as for preceding; 1953; Rév. P. Lootens leg.; RMCA • 1 ♀; same location as for preceding; 1954; Rév. P. Lootens leg.; RMCA • 1 ♀; same location as for preceding; 1958; Rév. P. Lootens leg.; RMCA • 1 ♀; same location as for preceding; Mar. 1954; Rév. P. Lootens leg.; RMCA • 1 ♀; same location as for preceding; Jul. 1952; Rév. P. Lootens leg.; RMCA • 1 ♀; same location as for preceding; Aug. 1951; Rév. P. Lootens leg.; RMCA • 1 ♀; same location as for preceding; Dec. 1951; Rév. P. Lootens leg.; RMCA • 2 ♂♂; same location as for preceding; Dec. 1951; Rév. P. Lootens leg.; RMCA • 6 ♀♀; Tshuapa: Bokuma; 00°06’ S, 18°41’ E; 1953; Rév. P. Lootens leg.; RMCA • 2 ♂♂; same location as for preceding; 1953; Rév. P. Lootens leg.; RMCA • 2 ♀♀; same location as for preceding; 1954; Rév. P. Lootens leg.; RMCA • 1 ♀; same location as for preceding; Mar. 1954; Rév. P. Lootens leg.; RMCA • 1 ♂; same location as for preceding; Mar. 1954; Rév. P. Lootens leg.; RMCA • 2 ♀♀; same location as for preceding; Apr. 1954; Rév. P. Lootens leg.; RMCA • 1 ♂; same location as for preceding; Apr. 1954; Rév. P. Lootens leg.; RMCA • 4 ♀♀; same location as for preceding; Jun. 1952; Rév. P. Lootens leg.; RMCA • 5 ♂♂; same location as for preceding; Jun. 1952; Rév. P. Lootens leg.; RMCA • 1 ♀; same location as for preceding; Dec. 1951; Rév. P. Lootens leg.; RMCA.

#### *Megachile (Eurymella) vittatula* Cockerell, 1920

**Material.** Cupboard 110, Box 1 (1♀)**. D.R. CONGO.** – **Sud-Kivu** 1 ♀; Uvira; 03°25’ S, 29°08’ E; Sep. 1958; Dr. Pasteels leg.; RMCA.

#### *Megachile (Eurymella) wahlbergi* Friese, 1901

**Material.** Cupboard 110, Box 7 (5♀♀ and 8♂♂)**. D.R. CONGO.** – **Kwango** • 4 ♀♀; Kasaï: Samsonge; 06°07’ S, 27°28’ E; 18 Mar. 1939; Mevr. Bequaert leg.; RMCA. – **Kwango** • 1 ♂; Lomami-Luputa; 11°00’ S, 26°44’ E; Apr. 1934; Dr. Bouvier leg.; RMCA – **Lualaba** • 4 ♂♂; Sandu; 09°41’ S, 22°53’ E; Apr. 1931; H.J. Brédo leg.; RMCA • 1 ♂; same location as for preceding; Feb. 1931; H.J. Brédo leg.; RMCA. – **Lualaba** • 1 ♂; Sandu; 09°41’ S, 22°53’ E; Apr. 1931; H.J. Brédo leg.; RMCA. – **Lomami** • 1 ♀; Sankuru: Pemba Zeo (Gandajika); 06°49’ S, 23°58’ E; 7 Feb. 1960; R. Maréchal leg.; RMCA • 1 ♂; same location as for preceding; 7 Feb. 1960; R. Maréchal leg.; RMCA.

#### *Megachile (Eurymella) waterbergensis* Strand, 1911

**Material.** Cupboard 110, Box 1 (1♀ and 4♂♂)**. D.R. CONGO.** – **Bas-Uele** • 1 ♂; Uele; 04°06’ N, 22°23’ E; Degreef leg.; RMCA. – **Haut-Lomami** • 2 ♂♂; Sankisia; 09°24’ S, 25°48’ E; 3 Sep. 1911; Dr. J. Bequaert leg.; RMCA • 1 ♀; same location as for preceding; 4 Sep. 1911; Dr. J. Bequaert leg.; RMCA. – **Haut-Uele**• 1 ♂; PNG Nagero (Dungu); 03°45’ N, 29°31’ E; 24 Mar. 1952; H. De Saeger leg.; RMCA.

#### *Megachile (Eutricharaea) admixta* Cockerell, 1931

**Material.** Cupboard 110, Box 14 (11♂♂)**. D.R. CONGO.** –**Haut-Katanga**, • 1 ♂; Elisabethville (Lubumbashi); 11°40’ S, 27°28’ E; 16 Nov. 1920; Dr. M. Bequaert leg.; RMCA.

– **Haut-Katanga** • 1 ♂; Elisabethville (Lubumbashi); 11°40’ S, 27°28’ E; 20 Oct. 1929; Dr. M. Bequaert leg.; RMCA • 1 ♂; same location as for preceding; Oct. 1928; Ch. seydel leg.; RMCA • 1 ♂; Elisabethville (60km ruiss. Kasepa); 11°40’ S, 27°28’ E; 23 Sep. 1923; Dr. M. Bequaert leg.; RMCA. – **Haut-Lomami** • 1 ♂; Bukama; 09°12’ S, 25°51’ E; 14 Jul. 1911; Dr. J. Bequaert leg.; RMCA • 1 ♂; same location as for preceding; Aug. 1923; Ch. Seydel leg.; RMCA • 1 ♂; Sankisia; 09°24’ S, 25°48’ E; 1 Sep. 1945; Dr. M. Bequaert leg.; RMCA • 1 ♂; same location as for preceding; 4 Sep. 1911; Dr. J. Bequaert leg.; RMCA. – **Lualaba** • 1 ♂; Dilolo; 10°41’ S, 22°21’ E; Oct – Nov. 1933; H. De Saeger leg.; RMCA • 1 ♂; same location as for preceding; Aug. 1931; G.F. de Witte leg.; RMCA.

#### *Megachile (Eutricharaea) boswendica* Cockerell, 1920

**Material.** Cupboard 110, Box 14 (10♀♀)**. D.R. CONGO.** Holotype – **Nord-Kivu** • 1 ♀; Boswenda; 01°20’ S, 29°25’ E, alt. 1900 m; 22 Oct. 1914; Dr. J. Bequaert leg.; RMCA. Paratypes – **Nord-Kivu** • 2 ♀♀; Masisi; 01°24’ S, 28°49’ E; 30 Dec. 1914; Dr. J. Bequaert leg.; RMCA.

– **Haut-Katanga** • 1 ♀; Haut Elila: Terr. Mwanga; 08°23’ S, 28°56’ E; Froideline leg.; RMCA. – **Ituri** • 2 ♀♀; Mahagi-Djugu; 01°56’ N, 30°30’ E; 7 – 9 Sep. 1931; Mme. L. Lebrun leg.; RMCA. – **Nord-Kivu** • 1 ♀; Beni à Lesse; 00°45’ N, 29°46’ E; Jul. 1911; Dr. Murtula leg.; RMCA • 2 ♀♀; Tshibinda; 02°19’ S, 28°45’ E; 21 – 27 Aug. 1931; T.D. Cockerell leg.; RMCA • 1 ♀; same location as for preceding; 24 Aug. 1931; T.D. Cockerell leg.; RMCA.

#### *Megachile (Eutricharaea) burungana* Cockerell, 1931

**Material.** Cupboard 110, Box 14 (2♂♂)**. D.R. CONGO.** Holotype – **Nord-Kivu** • 1 ♂; Burunga; 01°20’ S, 29°02’ E; Dr. M. Bequaert leg.; RMCA.

– **Nord-Kivu** • 1 ♂; PNA [Parc National Albert]; 00°03’ S, 29°30’ E; 16 – 29 Mar. 1954; P. Vanschuytbroeck & H. Synave leg.; RMCA.

#### *Megachile (Eutricharaea) derelictula* Cockerell, 1937

**Material.** Cupboard 110, Box 13 (3♀♀)**. D.R. CONGO.** – **Haut-Lomami** • 1 ♀; Kaniama; 07°31’ S, 24°11’ E; 19 Dec. 1952; Ch. Seydel leg.; RMCA. – **Maniema** • 1 ♀; Kasongo: Mwanakusu; 04°27’ S, 26°40’ E; Aug. 1959; P.L.G. Benoit leg.; RMCA. – **Nord-Kivu** • 1 ♀; Rutshuru; 01°11’ S, 29°27’ E; Nov. 1937; J. Ghesquière leg.; RMCA.

#### *Megachile (Eutricharaea) ekuivella* Cockerell, 1909

**Material.** Cupboard 110, Box 13 (18♀♀ and 10♂♂)**. D.R. CONGO.** – **Equateur** • 1 ♀; Eala (Mbandaka); 00°03’ N, 18°19’ E; 1939; G. Couteaux leg.; RMCA. – **Haut-Katanga** • 1 ♂; Elisabethville (Lubumbashi); 11°40’ S, 27°28’ E; 11 Sep. 1931; T.D. Cockerell leg.; RMCA • 1 ♀; same location as for preceding; 17 Sep. 1931; T.D. Cockerell leg.; RMCA • 1 ♂; same location as for preceding; 17 Sep. 1931; T.D. Cockerell leg.; RMCA • 1 ♂; same location as for preceding; 3 Nov. 1923; Ch. Seydel leg.; RMCA • 1 ♀; same location as for preceding; 4 Nov. 1920; Dr. M. Bequaert leg.; RMCA • 1 ♂; same location as for preceding; Sep. 1928; Ch. Seydel leg.; RMCA • 1 ♀; same location as for preceding; Aug. 1928; Ch. Seydel leg.; RMCA • 1 ♀; same location as for preceding; Oct. 1928; Ch. Seydel leg.; RMCA • 1 ♀; same location as for preceding; Oct. 1931; J. Ogilvie leg.; RMCA • 1 ♂; Elisabethville (60 Km ruiss. Kasepa); 11°40’ S, 27°28’ E; 23 Sep. 1923; Dr. M. Bequaert leg.; RMCA • 1 ♂; Elisabethville ruiss. Kasepa; 11°40’ S, 27°28’ E; 23 Sep. 1923; Dr. M. Bequaert leg.; RMCA • 1 ♀; La Kasepa (Lubumbashi); 11°40’ S, 27°28’ E; 23 Sep. 1923; Ch. Seydel leg.; RMCA • 1 ♀; Lubumbashi; 11°40’ S, 27°28’ E; 25 May 1920; Dr. M. Bequaert leg.; RMCA • 1 ♀; same location as for preceding; 27 May 1920; Dr. M. Bequaert leg.; RMCA • 1 ♀; same location as for preceding; 4 Oct. 1921; Dr. M. Bequaert leg.; RMCA • 1 ♀; Panda (Likasi); 11°00’ S, 26°44’ E; 9 Sep. 1920; Dr. M. Bequaert leg.; RMCA • 1 ♀; PNU Kabwe sur Muye; 08°47’ S, 26°52’ E, alt. 1320 m; 11 May 1948; Mis. G.F. de Witte leg.; RMCA • 1 ♂; PNU Masombwe s. Grde; Kafwe; 09°05’ S, 27°12’ E, alt. 1120 m; 16 Apr. 1948; Mis. G.F. de Witte leg.; RMCA. – **Haut-Lomami** • 1 ♂; PNU Kafwi Af. Dr. Lufwa; 08°56’ S, 27°10’ E; alt. 1780 m; 2 May 1929; Mis. G.F. de Witte leg.; RMCA. – **Kasaï Central**• 1 ♂; Luluabourg (Kananga); 05°54’ S, 21°52’ E; 14-17 May. 1919; P. Callewaert leg.; RMCA. – **Lualaba** • 1 ♀; Biano; 10°20’ S, 27°00’ E; 11 Aug. 1931; T.D. Cockerell leg.; RMCA • 3 ♀♀; Tenke; 10°35’ S, 26°06’ E; 30 Jul. – 9 Aug. 1931; T.D. Cockerell leg.; RMCA • 1 ♀; same location as for preceding; 30 Aug. 1931; T.D. Cockerell leg.; RMCA. – **Nord-Kivu** • 1 ♀; PNA: Ngoma (lac Biunu); 01°41’ S, 29°14’ E; 3 – 10 May 1935; Dr. H. Damas leg.; RMCA. – **Tanganyika** • 1 ♀; Kongolo; 05°24’ S, 27°00’ E; 30 Jan. 1911; Dr. J. Bequaert leg.; RMCA.

#### *Megachile (Eutricharaea) frontalis* Smith, 1853

**Material.** Cupboard 110, Box 13 (3♀♀)**. D.R. CONGO.** – **Haut-Katanga** • 1 ♀; Elisabethville (60km ruiss. Kasepa); 11°40’ S, 27°28’ E; 23 Sep. 1923; Dr. M. Bequaert leg.; RMCA. – **Lualaba** • 1 ♀; Lulua: Kapanga; 08°21’ S, 22°34’ E; 3 Dec. 1932; F.G. Overlaet leg.; RMCA. – **Maniema** • 1 ♀; Nyangwe; 04°13’ S, 26°11’ E; 1918; R. Mayné leg.; RMCA.

#### *Megachile (Eutricharaea) gratiosa* Gerstaecker, 1857

**Material.** Cupboard 110, Box 12 (75♀♀ and 26♂♂)**. D.R. CONGO.** – **Bas-Uele** • 1 ♀; Buta; 02°48’ N, 24°47’ E; 25 Jun. 1941; R. Fr. Hutsebaut leg.; RMCA • 1 ♀; Uele: Ibembo; 02°38’ N, 23°36’ E; Sep. 1949; R. Fr. Hutsebaut leg.; RMCA. – **Equateur** • 1 ♀; Basankusu; 01°13’ N, 19°49’ E; 1949; Zusters O.L.V. ten Bunderen leg.; RMCA • 2 ♀♀; Eala (Mbandaka); 00°03’ N, 18°19’ E; 1937; G. Couteaux leg.; RMCA • 2 ♂♂; same location as for preceding; 1937; G. Couteaux leg.; RMCA • 2 ♀♀; same location as for preceding; 1939; G. Couteaux leg.; RMCA • 1 ♂; same location as for preceding; 1939; G. Couteaux leg.; RMCA • 1 ♀; same location as for preceding; 5 Nov. 1931; H.J. Brédo leg.; RMCA • 1 ♂; same location as for preceding; 8 Nov. 1932; A. Corbisier leg.; RMCA • 4 ♂♂; same location as for preceding; Mar. 1932; H.J. Brédo leg.; RMCA • 2 ♀♀; same location as for preceding; May 1932; H.J. Brédo leg.; RMCA • 2 ♂♂; same location as for preceding; Nov. 1932; A. Corbisier leg.; RMCA • 1 ♀; Flandria (Ingende); 00°20’ S, 19°06’ E; 24 Mar. 1928; Rév. P. Hulstaert leg.; RMCA • 1 ♀; Lukolela (Bikoro); 01°03’ S, 17°12’ E; Nov. 1934; J. Ghesquière leg.; RMCA. – **Haut-Katanga** • 1 ♀; Elisabethville (Lubumbashi); 11°40’ S, 27°28’ E; 1 Nov. 1928; Dr. M. Bequaert leg.; RMCA • 1 ♀; same location as for preceding; 14 Mar. 1928; Dr. M. Bequaert leg.; RMCA • 1 ♂; same location as for preceding; 2 Oct. 1926; Dr. M. Bequaert leg.; RMCA • 1♀; same location as for preceding; 25 Apr. 1928; Dr. M. Bequaert leg.; RMCA • 1 ♀; same location as for preceding; Feb. 1933; De Loose leg.; RMCA • 1 ♀; same location as for preceding; Nov. 1926; Ch. Seydel leg.; RMCA • 1 ♀; same location as for preceding; Dec. 1934; P. Quarré leg.; RMCA • 1 ♀; Elisabethville (60km ruiss. Kasepa); 11°40’ S, 27°28’ E; 23 Sep. 1923; Dr. M. Bequaert leg.; RMCA • 1 ♀; La Kasepa (Lubumbashi); 11°40’ S, 27°28’ E; 23 Sep. 1923; Ch. Seydel leg.; RMCA • 1 ♀; Lubumbasi; 11°40’ S, 27°28’ E; 1 Apr. 1921; Dr. M. Bequaert leg.; RMCA • 1 ♀; same location as for preceding; 7 Jan. 1920; Dr. M. Bequaert leg.; RMCA • 1 ♂; same location as for preceding; 20 Dec. 1920; Dr. M. Bequaert leg.; RMCA • 1 ♀; same location as for preceding; 23 Feb. 1921; Dr. M. Bequaert leg.; RMCA • 1 ♀; same location as for preceding; 23 May 1921; Dr. M. Bequaert leg.; RMCA • 1 ♀; same location as for preceding; 27 Jan. 1921; Dr. M. Bequaert leg.; RMCA • 1 ♀; same location as for preceding; 28 Apr. 1921; Dr. M. Bequaert leg.; RMCA • 1 ♂; Mbiliwa-Wantu; 11°40’ S, 27°28’ E; Oct. 1907; Dr. Sheffield Neave leg.; RMCA • 1 ♂; PNK [Parc Nationa des Kundelungu]; 10°15’ S, 27°36’ E; Jan. – Feb. 1946; R. Verhulst leg.; RMCA • 1 ♂; PNU Kabwe sur Muye; 08°47’ S, 26°52’ E, alt. 1320 m; 11 May 1948; Mis. G.F. de Witte leg.;

RMCA. – **Haut-Lomami** • 4 ♀♀; Lomami: Kaniama; 07°31’ S, 24°11’ E; 1931; R. Massart leg.; RMCA • 1 ♀; Sankisia; 09°24’ S, 25°48’ E; 11 Aug. 1912; Dr. J. Bequaert leg.; RMCA • 1 ♀; same location as for preceding; 13 Sep. 1911; Dr. J. Bequaert leg.; RMCA • 1 ♂; same location as for preceding; 4 Sep. 1911; Dr. J. Bequaert leg.; RMCA. – **Haut-Uele** • 1 ♀; Haut-Uele: Moto; 03°03’ N, 29°28’ E; 1923; L. Burgeon leg.; RMCA • 1 ♂; same location as for preceding; 1923; L. Burgeon leg.; RMCA • 1 ♀; Uele: Bayenga, terr. Wamba; 02°09’ N, 28°00’ E, alt. 810 m; 12 – 22 Aug. 1956; R. Castelain leg.; RMCA • 1 ♀; Uele: Dungu; 03°37’ N, 28°34’ E; Degreef leg.; RMCA • 1 ♀; Watsa à Niangara; 03°02’ N, 29°32’ E; Jul. 1920; L. Burgeon leg.; RMCA. – **Ituri** • 1 ♀; Ituri: Lubero; 00°10’ S, 29°14’ E; 1928; Mme. Van Riel leg.; RMCA • 1 ♀; Ituri: Mahagi; 02°18’ N, 30°59’ E; 25 May 1925; Dr. H. Schouteden leg.; RMCA • 1 ♀; Kilo (Djugu); 01°50’ S, 30°09’ E; 1930; G. du Soleil leg.; RMCA • 1 ♀; Lac Albert: Kasenyi; 01°24’ N, 30°26’ E; 15 May 1935; H.J. Brédo leg.; RMCA • 1 ♀; Mongbwalu; 01°55’ N, 30°02’ E; 1939; Mme. Scheitz leg.; RMCA • 1 ♀; same location as for preceding; 4 Apr. 1937; Mme. Scheitz leg.; RMCA • 1 ♀; Nioka; 02°10’ N, 30°40’ E; 20 Aug. 1931; H.J. Brédo leg.; RMCA. – **Kasaï** • 1 ♀; Luebo; 05°20’ S, 21°24’ E; 2 May 1919; P. Callewaert leg.; RMCA. – **Kinshasa** • 1 ♀; Léopoldville (Kinshasa); 04°19’ S, 15°19’ E; 15 Sep. 1910; Dr. J. Bequaert leg.; RMCA. – **Kwilu** • 2 ♀♀; Leverville (Bulungu); 04°50’ S, 18°44’ E; 1928; Mme. J. Tinant leg.; RMCA. – **Lomami •** 1 ♀; Sankuru: M’Pemba Zeo (Gandajika); 06°49’ S, 23°58’ E; Nov. 1957; R. Maréchal leg.; RMCA. – **Lualaba** • 1 ♀; Kapanga; 08°21’ S, 22°34’ E, Nov. 1933; F.G. Overlaet leg.; RMCA • 1 ♂; Lualaba: Kolwezi; 10°44’ S, 25°28’ E; 16 Nov. 1952; Mme. L. Gilbert leg.; RMCA. – **Maniema** • 1 ♀; Kasongo: Mwanakusu; 04°27’ S, 26°40’ E; Aug. 1960; P.L.G. Benoit leg.; RMCA • 1 ♂; Kibombo; 03°54’ S, 25°55’ E; 1 Nov. 1910; Dr. J. Bequaert leg.; RMCA • 1 ♀; same location as for preceding; 10 Nov. 1910; Dr. J. Bequaert leg.; RMCA • 1 ♀; Nyangwe; 04°13’ S, 26°11’ E; 29 Nov. 1910; Dr. J. Bequaert leg.; RMCA • 1 ♀; Terr. de Kasongo, Riv. Lumami; 04°27’ S, 26°40’ E; 04°27’ S, 26°40’ E; Oct. – Dec. 1959; P.L.G. Benoit leg.; RMCA. – **Nord-Kivu** • 1 ♀; N. Lac Kivu: Rwankwi; 01°20’ S, 29°22’ E; May 1948; J.V. Leroy leg.; RMCA • 1 ♀; PNA SL Edouard: Kamande; 00°20’ S, 29°30’ E, alt. 925 m; 8 Apr. 1936; L. Lippens leg.; RMCA • 1 ♀; Rutshuru; 01°11’ S, 29°27’ E; 1937; J. Ghesquière leg.; RMCA • 1 ♂; same location as for preceding; Nov. 1937; J. Ghesquière leg.; RMCA • 1 ♀; same location as for preceding; Dec 1914; J. Ghesquière leg.; RMCA • 1 ♂; same location as for preceding; Dec 1937; J. Ghesquière leg.; RMCA • 1 ♂; Walikale; 01°25’ S, 28°03’ E; 7 Jan. 1915; Dr. J. Bequaert leg.; RMCA. – **Sankuru** • 2 ♀♀; Sankuru: Lodja; 03°29’ S, 23°26’ E; Jan. – May 1925; J. Ghesquière leg.; RMCA • 1 ♂; same location as for preceding; Jan. – May 1929; J. Ghesquière leg.;

RMCA. – **Sud-Kivu** • 2 ♀♀; Ibanda; 02°29’ S, 28°51’ E; 1952; M. Vandelannoite leg.; RMCA • 1 ♀; Kalehe Makwe; 02°07’ S, 28°55’ E; Feb. 1950; H. Bomans leg.; RMCA • 1 ♀; Kavumu à Kabunga, 82 km (Mingazi); 02°18’ S, 28°49’ E; May – Jun 1951; H. Bomans leg.; RMCA • 1 ♀; Uvira; 03°25’ S, 29°08’ E; Sep. 1958; Dr. J. Pasteels leg.; RMCA. – **Sud-Ubangi** • 1 ♀; Libenge; 03°39’ N, 18°38’ E; 29 Dec. 1931; H.J. Brédo leg.; RMCA. – **Tanganyika** • 1 ♀; Buli (Kabalo); 05°50’ S, 26°55’ E; 18 Feb. 1911; Dr. J. Bequaert leg.; RMCA • 1 ♀; Lukuga r. Niemba; 05°57’ S, 28°26’ E; Nov. 1917 – Jan. 1918; Dr. Pons leg.; RMCA • 1 ♂; Tanganyika: Moba; 07°02’ S, 29°47’ E, alt. 780 m; Aug.-Sep. 1953; H. Bomans leg.; RMCA • 1 ♂; same location as for preceding; Oct. – Nov. 1953; H. Bomans leg.; RMCA • 1 ♂; Vallée Lukuga; 05°40’ S, 26°55’ E; Nov. 1911; Dr. Schwetz leg.; RMCA. – **Tshopo** • 1 ♀; Ponthierville (Ubundu); 00°22’ S, 25°29’ E; 21 Oct. 1910; Dr. J. Bequaert leg.; RMCA • 2 ♀♀; Ponthierville (Ubundu); 00°22’ S, 25°29’ E; 25 Oct. 1910; Dr. J. Bequaert leg.; RMCA • 1 ♂; same location as for preceding; 25 Oct. 1910; Dr. J. Bequaert leg.; RMCA • 1 ♀; Yangambi; 00°46’ N, 24°27’ E; 1939; Dr. Parent leg.; RMCA. – **Tshuapa** • 1 ♀; Bokuma; 00°06’ S, 18°41’ E; Juillet 1952; Rév. P. Lootens leg.; RMCA.

#### *Megachile (Eutricharaea) gratiosella* Cockerell, 1935

**Material.** Cupboard 110, Box 12 (6♂♂)**. D.R. CONGO.** – **Haut-Lomami** • 1 ♂; Bukama; 09°12’ S, 25°51’ E; 10 Jul. 1911; Dr. J. Bequaert leg.; RMCA • 1 ♂; same location as for preceding; 14 Jul. 1911; Dr. J. Bequaert leg.; RMCA • 2 ♂♂; same location as for preceding; 28 Aug. 1931; T.D. Cockerell leg.; RMCA.

– **Haut-Uele** • 1 ♂; PNG Nagero (Dungu); 03°45’ N, 29°31’ E; 24 Mar. 1952; H. De Saeger leg.; RMCA.

– **Sud-Kivu** • 1 ♂; PNKB: Luofu; 00°37’ S, 29°07’ E, alt. 1700 m; 10 Dec. 1934; G.F. de Witte leg.; RMCA.

#### *Megachile (Eutricharaea) hypopyrrha* Cockerell, 1937

**Material.** Cupboard 110, Box 12 (2♂♂)**. D.R. CONGO.** – **Haut-Katanga**• 1 ♂; PNU Lusinga; 08°56’ S, 27°12’ E; 13 Jun. 1945; G.F. de Witte leg.; RMCA. – **Sud-Kivu** • 1 ♂; Ibanda; 02°29’ S, 28°51’ E; 1952; M. Vandelannoite r leg.; RMCA.

#### *Megachile (Eutricharaea) ituriella* Pasteels, 1965

**Material.** Cupboard 110, Box 12 (1♀)**. D.R. CONGO.** Holotype – **Ituri** • 1 ♀; Ituri: Kawa; 01°34’ N, 30°32’ E; 2 May 1929; A. Collart leg.; RMCA.

#### *Megachile (Eutricharaea) luteoalba* Pasteels, 1973

**Material.** Cupboard 110, Box 14A (1♀)**. D.R. CONGO.** Holotype – **Haut-Katanga** • 1 ♀; Kipopo; 11°33’ S, 27°21’ E; 27 Aug. 1961; R. Maréchal leg.; RMCA.

#### *Megachile (Eutricharaea) malangensis* Friese, 1904

**Material.** Cupboard 110, Box 14 (1 ♂)**. D.R. CONGO.** – **Haut-Katanga** • 1 ♂; PNU Lusinga; 08°55’ S, 27°12’ E; 13 Jun. 1911; G.F. de Witte leg.; RMCA.

#### *Megachile (Eutricharaea) multidens* Fox, 1891

**Material.** Cupboard 110, Box 13 (1♂)**. D.R. CONGO.** – **Haut-Katanga** • 1 ♂; Elisabethville, riv. Kimilolo; 11°40’ S, 27°28’ E; 10 Nov. 1920; Dr. M. Bequaert leg.; RMCA.

#### *Megachile (Eutricharaea) natalica* Cockerell, 1920

**Material.** Cupboard 110, Box 14 (2♀♀)**. D.R. CONGO.** – **Haut-Lomami** • 1 ♀; Kalongwe; 11°01’ S, 25°13’ E; 11 Aug. 1911; Dr. M. Bequaert leg.; RMCA. – **Nord-Kivu** • 1 ♀; PNA [Parc National Albert]; 00°03’ S, 29°30’ E; 16-24 Mar. 1954; P. Vanschuytbroeck leg.; RMCA.

#### *Megachile (Eutricharaea) panda* Cockerell, 1931

**Material.** Cupboard 110, Box 14 (1♂)**. D.R. CONGO.** Holotype – **Haut-Katanga** • 1 ♂; Panda (Likasi); 11°00’ S, 26°44’ E; 9 Nov. 1911; Dr. M. Bequaert leg.; RMCA.

#### *Megachile (Eutricharaea) rhodesica* Cockerell, 1920

**Material.** Cupboard 110, Box 12 (3♀♀)**. D.R. CONGO.** – **Haut-Katanga** • 1 ♀; Elisabethville (Lubumbashi); 11°40’ S, 27°28’ E; 20 Sep. 1923; Dr. M. Bequaert leg.; RMCA. – **Haut-Lomami** • 1 ♀; Sankisia; 09°24’ S, 25°48’ E; 13 Sep. 1911; Dr. J. Bequaert leg.; RMCA. – **Ituri** • 1 ♀; Ituri: Blukwa; 01°45’ N, 30°36’ E; 28 Dec. 1928; A. Collart leg.; RMCA.

#### Megachile (Eutricharaea) ruficheloides Strand, 1911

**Material.** Cupboard 110, Box 12 (14♀♀)**. D.R. CONGO.** – **Haut-Lomami** • 4 ♀♀; Bukama; 09°12’ S, 25°51’ E; 10 Jul. 1911; Dr. J. Bequaert leg.; RMCA • 1 ♀; same location as for preceding; 14 Jul. 1911; Dr. J. Bequaert leg.; RMCA • 1 ♀; same location as for preceding; 7 Jul. 1911; Dr. J. Bequaert leg.; RMCA • 1 ♀; Kalongwe; 11°01’ S, 25°13’ E; 15 Sep. 1911; Dr. J. Bequaert leg.; RMCA • 1 ♀; Sankisia; 09°24’ S, 25°48’ E; 20 Sep. 1911; Dr. J. Bequaert leg.; RMCA • 1 ♀; same location as for preceding; 29 Sep. 1911; Dr. M. Bequaert leg.; RMCA • 3 ♀♀; same location as for preceding; 4 Sep. 1911; Dr. J. Bequaert leg.; RMCA • 1 ♀; same location as for preceding; 4 Oct. 1911; Dr. J. Bequaert leg.; RMCA. – **Lualaba**• 1 ♀; Dilolo; 10°41’ S, 22°21’ E; Aug. 1931; G.F. de Witte leg.; RMCA.

#### *Megachile (Eutricharaea) semiflava* Cockerell, 1935

**Material.** Cupboard 110, Box 12 (11♀♀)**. D.R. CONGO.** – **Haut-Katanga** • 1 ♀; Elisabethville (Lubumbashi); 11°40’ S, 27°28’ E; 13 May 1920; Dr. M. Bequaert leg.; RMCA • 1 ♀; same location as for preceding; 5 Apr. 1912; Dr. J. Bequaert leg.; RMCA • 1 ♀; Lukonzolwa; 08°47’ S, 28°38’ E; 31 Dec. 1941; Dr. M. Bequaert leg.; RMCA. – **Haut-Uele** • 1 ♀; PNG Akam; 03°40’ N, 29°00’ E; 21 Apr. 1950; H. De Saeger leg.; RMCA. – **Ituri** • 1 ♀; Lac Albert: Kasenyi; 01°24’ N, 30°26’ E; 15 May 1935; H.J. Brédo leg.; RMCA. – **Maniema** • 1 ♀; Nyangwe; 04°13’ S, 26°11’ E; 10 Dec. 1910; Dr. J. Bequaert leg.; RMCA. –

**Sankuru** • 1 ♀; Makarikari; 03°27’ S, 22°26’ E; 6 – 23 Aug. 1930; T.D. Cockerell leg.; RMCA. – **Sud-Kivu** • 1 ♀; Bukavu; 02°29’ S, 28°51’ E; 28 Aug. 1931; T.D. Cockerell leg.; RMCA • 2 ♀♀; Uvira; 03°25’ S, 29°08’ E; 29 Aug. 1931; T.D. Cockerell leg.; RMCA. – **Tanganyika** • 1 ♀; Tanganyika: Kabalo; 06°03’ S, 26°55’ E; 7 Jul. 1947; Dr. M. Poll leg.; RMCA.

#### *Megachile (Eutricharaea) stellarum* Cockerell, 1920

**Material.** Cupboard 110, Box 14 (1♀)**. D.R. CONGO.** – **Haut-Lomami** • 1 ♀; Sankisia; 09°24’ S, 25°48’ E; 8 Apr. 1911; Dr. J. Bequaert leg.; RMCA.

#### *Megachile (Megella) bouyssoui* Vachal, 1903

**Material.** Cupboard 110, Box 8 (25♀♀ and 8♂♂)**. D.R. CONGO.** – **Equateur** • 2 ♀♀; Eala (Mbandaka); 00°03’ N, 18°19’ E; May 1932; H.J. Brédo leg.; RMCA • 3 ♂♂; same location as for preceding; May 1932; H.J. Brédo leg.; RMCA • 3 ♀♀; same location as for preceding; Jun. 1932; A. Corbisier leg.; RMCA • 1 ♀; same location as for preceding; 14 Mar. 1933; H.J. Brédo leg.; RMCA • 2 ♀♀; same location as for preceding; Nov. 1932; A. Corbisier leg.; RMCA • 2 ♀♀; same location as for preceding; 28 Nov. 1931; H.J. Brédo leg.; RMCA • 3 ♀♀; same location as for preceding; Mar. 1932; H.J. Brédo leg.; RMCA • 1 ♀; same location as for preceding; Dec. 1932; A. Corbisier leg.; RMCA. – **Ituri** • 2 ♀♀; Haut-Ituri; 29°03’, 29°03’ E, alt. 1150 m; 1906; H. Wilmin leg.; RMCA – **Lualaba** • 2 ♂♂; same location as for preceding; Oct. 1932; F.G. Overlaet leg.; RMCA. – **Maï-Ndombe** • 1 ♀; Bena Bendi (Oshwe); 04°18’ S, 20°22’ E; May 1915; R. Mayné leg.; RMCA. – **Maniema** • 1 ♀; Lokandu: Ile Biawa; 02°31’ S, 25°47’ E; Jul. 1939; Lt. Vissers leg.; RMCA. – **Mongala** • 1 ♀; Congo: Bolombo; 02°38’ N, 21°36’ E; ex coll. Breuning leg.; RMCA • 1 ♂; same location as for preceding; ex coll. Breuning leg.; RMCA • 1 ♂; Itimbiri; 02°03’ N, 22°45’ E; 23 May 1918; Dr. Rodhain leg.; RMCA. – **Nord-Kivu** • 1 ♀; Gamangui. Congo; 00°20’ N, 29°45’ E; Feb. 1910; Lang & Chapin leg.; RMCA. – **Nord-Ubangi** • 1 ♀; Ubangi: Karawa (Businga); 03°21’ N, 20°18’ E; 1936; Rév. Wallin leg.; RMCA • 1 ♀; Yakoma; 04°06’ N, 22°23’ E; 5 – 17 Feb. 1932; H.J. Brédo leg.; RMCA. – **Tshuapa** • 1 ♀; Bokuma; 00°06’ S, 18°41’ E; Jul. 1952; Rév. P. Lootens leg.; RMCA • 2 ♀♀; Moma; 00°45’ S, 21°56’ E; Jun. 1925; J. Ghesquière leg.; RMCA • 1 ♂; Tshuapa: Bokuma; 00°06’ S, 18°41’ E; Mar. 1952; Rév. P. Lootens leg.; RMCA.

#### *Megachile (Megella) exsecta* Pasteels, 1960

**Material.** Cupboard 110, Box 8 (1♂)**. D.R. CONGO.** Holotype – **Haut-Lomami** • 1 ♂; Sankisia; 09°24’ S, 25°48’ E; 20 Aug. 1911; Dr. J. Bequaert leg.; RMCA.

#### (*Resin*-)dauber bees, *Megachile* (*Callomegachile*) *chrysorrhoea* Gerstaecker, 1857, *Chalicodoma* (*Callomegachile*) *chrysorrhaea* (Gerstaecker, 1857)

**Material.** Cupboard 110, Box 27 (1♀). **D.R. CONGO.** – **Sankuru** • 1 ♀; Lusambo; 04°58’ S, 23°26’ E; 1936; Dir. Gén. M. Gol leg.; RMCA.

#### *Magachile* (*Callomegachile*) *devexa* Vachal, 1903, *Chalicodoma* (*Callomegachile*) *devexa* (Vachal, 1903)

Following Pasteels (1965), *Megachile* (*Callomegachile*) *devexa* Vachal, 1903 includes *Chalicodoma* (*Callomegachile*) *duponti* Vachal, 1903 by synonymy. We list them separetly below as it was organized in the RMCA collections.

#### *Magachile* (*Callomegachile*) *devexa* Vachal, 1903, *Chalicodoma* (*Callomegachile*) *devexa* (Vachal, 1903)

**Material.** Cupboard 110, Box 26 (4 ♀♀ and 14♂♂). **D.R. CONGO.** – **Kasaï** • 1 ♀; Luebo; 05°20’ S, 21°24’ E; 25 May 1938; Mevr. Bequaert leg.; RMCA. – **Kwango** • 1 ♀; Kwango; 04°48’ S, 17°02’ E; 1925; P. Vanderijst leg.; RMCA. – **Lualaba** • 12 ♂♂; Lulua: Kapanga; 08°21’ S, 22°34’ E; Oct. 1932; F.G. Overlaet leg.; RMCA. – **Maniema** • 1 ♂; Kibombo; 03°54’ S, 25°55’ E; Sep.-Nov. 1930; H.J. Brédo leg.; RMCA • 1 ♂; Malela; 04°22’ S, 26°08’ E; Jan. 1914; L. Burgeon leg.; RMCA • 1 ♀; same location as for preceding; Dec. 1913; L. Burgeon leg.; RMCA. – **Tshuapa** • 1 ♀; Bokuma; 00°06’ S, 18°41’ E; Dec. 1951; Rév. P. Lootens leg.; RMCA.

#### *Magachile* (*Callomegachile*) *duponti* Vachal, 1903, *Chalicodoma* (*Callomegachile*) *duponti* (Vachal, 1903)

**Material.** Cupboard 110, Box 26A (70♀♀ and 110♂♂). **D.R. CONGO.** – **Equateur** • 1 ♂; Bamania (Mbandaka); 00°01’ N, 18°19’ E; 8 May 1924; Dr. M. Bequaert leg.; RMCA • 1 ♂; Boleke wa Bondele; 00°01’ S, 19°36’ E; 24 Nov. 1926; Rév. P. Hulstaert leg.; RMCA • 1 ♀; Eala (Mbandaka); 00°03’ N, 18°19’ E; 11 Oct. 1931; H.J. Brédo leg.; RMCA • 1 ♀; same location as for preceding; 14 Nov. 1931; H.J. Brédo leg.; RMCA • 1 ♀; same location as for preceding; 15-30 Nov. 1929; H.J. Brédo leg.; RMCA • 1 ♀; same location as for preceding; 17 Nov. 1931; H.J. Brédo leg.; RMCA • 1 ♀; same location as for preceding; 21 Nov. 1931; H.J. Brédo leg.; RMCA • 1 ♀; same location as for preceding; 22 Apr. 1932; H.J. Brédo leg.; RMCA • 1 ♀; same location as for preceding; 4 Apr. 1932; H.J. Brédo leg.; RMCA • 1 ♂; same location as for preceding; 7 Jul. 1932; A. Corbisier leg.; RMCA • 1 ♀; same location as for preceding; 5 Nov. 1931; H.J. Brédo leg.; • 1 ♀; same location as for preceding; 7 Nov. 1931; H.J. Brédo leg.; RMCA • 2 ♀♀; same location as for preceding; Mar. 1932; H.J. Brédo leg.; RMCA • 1 ♀; same location as for preceding; 7 Nov. 1931; H.J. Brédo leg.; RMCA • 3 ♂♂; same location as for preceding; Mar. 1932; H.J. Brédo leg.; RMCA • 3 ♀♀; same location as for preceding; Apr. 1932; H.J. Brédo leg.; RMCA leg.; RMCA • 3 ♀♀; same location as for preceding; May 1932; H.J. Brédo leg.; RMCA • 3 ♂♂; same location as for preceding; May 1932; H.J. Brédo leg.; RMCA • 2 ♀♀; same location as for preceding; Jun. 1932; H.J. Brédo leg.; RMCA • 4 ♂♂; same location as for preceding; Jun. 1932; A. Corbisier leg.; RMCA • 3 ♀♀; same location as for preceding; Nov. 1932; H.J. Brédo leg.; RMCA • 2 ♀♀; same location as for preceding; Dec. 1932; H.J. Brédo leg.; RMCA • 1 ♀; Ubangi: Mokolo; 01°35’ N, 29°05’ E; 5 Dec. 1931; H.J. Brédo leg.; RMCA RMCA. – **Ituri** • 5 ♀♀; Haut-Ituri, 29°03’, 29°03’ E; 1906; H. Wilmin leg.; RMCA. – **Kasaï** • 1 ♀; Ilebo; 04°20’ S, 20°36’ E; 14-15 Jul. 1925; S.A.R. Prince Léopold leg.; RMCA • 1 ♀; Lutzkwadi (Ilebo); 04°20’ S, 20°36’ E; 17 Jun. 1946; V. Lagae leg.; RMCA. – **Kasaï Central** • 1 ♀; Luluabourg (Kananga); 05°54’ S, 21°52’ E; Apr. 1939; J.J. Deheyn leg.; RMCA. – **Kinshasa** • 1 ♀; Barumbu; 01°14’ S, 23°31’ E; Dec. 1920; J. Ghesquière leg.; RMCA • 1 ♀; Léopoldville (Kinshasa); 04°19’ S, 15°19’ E; 6 Dec. 1925; Rév. P. Hulstaert leg.; RMCA. – **Kongo Central** • 2 ♀♀; Camp de Lukula; 05°23’ S, 12°57’ E; 1911; Dr. Daniel leg.; RMCA • 1 ♀; Lemfu (Madimba); 05°18’ S, 15°13’ E; Apr. 1945; Rév. P. De Beir leg.; RMCA. – **Kwango** • 1 ♀; Kwango; 08°00’ S, 20°00’ E; Jun. 1925; P. Vanderijst leg.; RMCA. – **Kwilu** • 1 ♀; Kikwit; 05°02’ S, 18°49’ E; 1920; P. Vanderijst leg.; RMCA. – **Lualaba** • 1 ♀; Lulua: Kapanga; 08°21’ S, 22°34’ E; Oct. 1932; F.G. Overlaet leg.; RMCA • 92 ♂♂; same location as for preceding; Oct. 1932; F.G. Overlaet leg.; RMCA. – **Maï-Ndombe** • 1 ♀; Bena Bendi (Oshwe); 04°18’ S, 20°22’ E; May 1915; R. Mayné leg.; RMCA. – **Maniema** • 1 ♀; Kibombo; 03°54’ S, 25°55’ E; Sep. – Oct. 1930; H.J. Brédo leg.; RMCA • 1 ♀; Lokandu: Ile Biawa; 02°31’ S, 25°47’ E; Jul. 1939; Lt. Vissers leg.; RMCA • 1 ♂; Malela; 04°22’ S, 26°08’ E; Jan. 1914; L. Burgeon leg.; RMCA. – **Mongala** • 1 ♀; Congo: Bolombo; 02°38’ N, 21°36’ E; ex coll. Breuning leg.; RMCA • 1 ♂; Itimbiri; 02°03’ N, 22°45’ E; 23 May 1913; Dr. Rodhain leg.; RMCA. – **Nord-Ubangi** • 1 ♂; Ubangi: Yakoma; 04°06’ N, 22°23’ E; 15 Feb. 1931; H.J. Brédo leg.; RMCA • 1 ♂; Yakoma; 04°06’ N, 22°23’ E; 5-17 Feb. 1932; H.J. Brédo leg.; RMCA. – **Sankuru** • 1 ♂; Sankuru: Komi; 03°23’ S, 23°46’ E; Apr.1930; J. Ghesquière leg.; RMCA • 5 ♀♀; same location as for preceding; May 1930; J. Ghesquière leg.; RMCA • 4 ♀♀; same location as for preceding; Jun. 1930; J. Ghesquière leg.; RMCA • 4 ♀♀; same location as for preceding; Jun. 1930; J. Ghesquière leg.; RMCA • 6 ♀♀; same location as for preceding; Jul. 1930; J. Ghesquière leg.; RMCA. – **Sud-Ubangi** • 1 ♀; Ubangi: Boma-Motenge; 03°15’ N, 18°39’ E; Dec.1931; H.J. Brédo leg.; RMCA. – **Tshopo** • 1 ♂; Ponthierville (Ubundu); 00°22’ S, 25°29’ E 22 Oct. 1910; Dr. J. Bequaert leg.; RMCA. – **Tshuapa** • 1 ♀; Boende; 00°13’ S, 20°52’ E; Jan. 1952; Rév. P. Lootens leg.; RMCA • 1 ♀; Bokuma; 00°06’ S, 18°41’ E; Dec. 1951; Rév. P. Lootens leg.; RMCA • 1 ♀; Tshuapa: Bokuma; 00°06’ S, 18°41’ E; Dec. 1951; Rév. P. Lootens leg.; RMCA.

#### *Megachile* (*Callomegachile*) *excavata* Cockerell, 1937, *Chalicodoma* (*Callomegachile*) *excavata* (Cockerell, 1937)

**Material.** Cupboard 110, Box 32 (6♀♀). **D.R. CONGO.** Holotype – **Tanganyika** • 1 ♀; Kiambi (Manono); 07°19’ S, 28°01’ E; 5 – 15 May 1931; G.F. de Witte leg.; RMCA.

– **Haut-Katanga** • 1 ♀; Elisabethville (Lubumbashi); 11°40’ S, 27°28’ E; 20 Sep. 1923; Dr. M. Bequaert leg.; RMCA • 1 ♀; same location as for preceding; Sep. 1934; Dr. M. Bequaert leg.; RMCA • 1 ♀; Mwema; 08°13’ S, 27°28’ E; Jul. 1927; A. Bayet leg.; RMCA • 1 ♀; PNU Kabwe sur Muye; 08°47’ S, 26°52’ E, alt. 1320 m; 11 May 1948; Mis. G.F. de Witte leg.; RMCA. – **Haut-Lomami** • 1 ♀; PNU Munoi, bifure Lupiala; 08°50’ S, 26°44’ E, alt. 890 m; 28 May-15 Jun. 1948; Mis. G.F. de Witte leg.; RMCA.

#### *Megachile* (*Callomegachile*) *montibia* Strand, 1911, *Chalicodoma* (*Callomegachile*) *montibia* (Strand, 1911)

**Material.** Cupboard 110, Box 32 (6♀♀). **D.R. CONGO.** – **Haut-Katanga** • 1 ♀; PNU Kabwe sur Muye; 08°47’ S, 26°52’ E, alt. 1320 m; 11 May 1948; Mis. G.F. de Witte leg.; RMCA. – **Nord-Kivu**• 1 ♀; PNA [Parc National Albert]; 00°03’ S, 29°30’ E, 23 Mar. 1954; P. Vanschuytbroeck & H. Synave leg.; RMCA • 2 ♀♀; same location as for preceding; 27 Aug. 1953; P. Vanschuytbroeck & H. Synave leg.; RMCA • 2 ♀♀; same location as for preceding; 28 Mar. 1954; P. Vanschuytbroeck & H. Synave leg.; RMCA.

#### *Megachile* (*Callomegachile*) *punctolineata* Cockerell, 1935, *Chalicodoma* (*Callomegachile*) *punctolineata* (Cockerell, 1935)

**Material.** Cupboard 110, Box 26 (2♀♀). **D.R. CONGO.** Holotype – **Sankuru** • 1 ♀; Sankuru: Komi; 03°23’ S, 23°46’ E; May 1930; J. Ghesquière leg.; RMCA. Paratype – **Sankuru** • 1 ♀; same location as for preceding; Jul. 1930; J. Ghesquière leg.; RMCA.

#### *Megachile* (*Callomegachile*) *rufipennis* (Fabricius, 1793), *Chalicodoma* (*Callomegachile*) *rufipennis* (Fabricius, 1793)

**Material.** Cupboard 110, Box 27 (49 ♀ and 18♂♂). **D.R. CONGO.** – **Bas-Uele** • 1 ♀; Bambesa; 03°28’ N, 25°43’ E; 1938; P. Henrard leg.; RMCA • 1 ♀; same location as for preceding; 1 Oct. 1934; J.V. Leroy leg.; RMCA • 1 ♀; same location as for preceding; 10 May 1937; J. Vrydagh leg.; RMCA • 1 ♀; same location as for preceding; 14 May 1938; J. Vrydagh leg.; RMCA • 1 ♂; same location as for preceding; 14 May 1938; J. Vrydagh leg.; RMCA • 1 ♀; same location as for preceding; 24 Apr. 1937; J. Vrydagh leg.; RMCA • 1 ♀; same location as for preceding; 4 Apr. 1937; J. Vrydagh leg.; RMCA • 2 ♀♀; same location as for preceding; 6 Jul. 1937; J. Vrydagh leg.; RMCA • 2 ♀♀; same location as for preceding; 8 Jul. 1937; J. Vrydagh leg.; RMCA • 5 ♀♀; same location as for preceding; 9 May 1938; P. Henrard leg.; RMCA • 1♀; same location as for preceding; Jan. 1934; H.J. Brédo leg.; RMCA • 3 ♀♀; same location as for preceding; May 1938; P. Henrard leg.; RMCA • 3 ♀♀; same location as for preceding; 14 May 1938; P. Henrard leg.; RMCA • 2 ♂♂; same location as for preceding; 14 May 1938; P. Henrard leg.; RMCA • 1 ♀; same location as for preceding; Jul. 1937; J. Vrydagh leg.; RMCA • 1 ♀; Uele: Bambesa; 03°28’ N, 25°43’ E; 20 Oct. 1933; J.V. Leroy leg.; RMCA • 1 ♀; Uele: Dingila; 03°37’ N, 26°03’ E; Jul. 1933; H.J. Brédo leg.; RMCA • 1 ♂; same location as for preceding; Jul. 1933; H.J. Brédo leg.; RMC• 1 ♀; Uele: Tukpwo; 04°26’ N, 25°51’ E; Aug. 1937; J. Vrydagh leg.; RMCA. – **Haut-Katanga** • 1 ♀; Elisabethville (Lubumbashi); 11°40’ S, 27°28’ E; Jan. 1932; Ch. Seydel leg.; RMCA • 1 ♂; same location as for preceding; Nov. 1934; P. Quarré leg.; RMCA • 1 ♀; Kasenga; 10°22’ S, 28°38’ E; Feb. 1931; R.P. Vankerckhoven leg.; RMCA • 1 ♀; same location as for preceding; Apr. 1931; R.P. Vankerckhoven leg.; RMCA • 1 ♀; Kipushi; 11°46’ S, 27°15’ E; Mar. 1932; G.F. de Witte leg.; RMCA. – **Haut-Lomami** • 1 ♀; Bukama; 09°12’ S, 25°51’ E; 20 Apr. 1911; Dr. J. Bequaert leg.; RMCA • 2 ♀♀; Lualaba: Kabongo; 07°20’ S, 25°35’ E; 16 Nov. 1952; Ch. Seydel leg.; RMCA • 1 ♂; same location as for preceding; 16 Nov. 1952; Ch. Seydel leg.; RMCA • 1 ♂; same location as for preceding; 27 Dec. 1952; Ch. Seydel leg.; RMCA • 1 ♂; same location as for preceding; 27 Dec. 1953; Ch. Seydel leg.; RMCA • 1 ♂; same location as for preceding; 30 Dec. 1952; Ch. Seydel leg.; RMCA •1 ♀; same location as for preceding; 31 Dec. 1952; Ch. Seydel leg.; RMCA • 1 ♀; same location as for preceding; 7 Jan. 1953; Ch. Seydel leg.; RMCA • 3 ♂♂; same location as for preceding; 7 Jan. 1953; Ch. Seydel leg.; RMCA • 2 ♀♀; PNU Mabwe (R.P. Lac Upemba; 585m); 08°39’ S, 26°31’ E; 1-11 Jan. 1949; Mis. G.F. de Witte leg.; RMCA. – **Haut-Uele** • 1 ♀; PNG [Parc National de Garamba]; 03°40’ N, 29°00’ E 5 Oct. 1950; H. De Saeger leg.; RMCA • 1 ♀; Watsa à Niangara; 03°02’ N, 29°32’ E; Jul. 1920; L. Burgeon leg.;

RMCA. – **Kongo Central** • 1 ♀; Boma: Kimbuadi; 05°51’ S, 13°03’ E; 1959; Mme. A. Van Alstein leg.; RMCA • 1 ♂; Kisantu (Madimba); 05°08’ S, 15°06’ E; 1927; R. P. Vanderyst leg.; RMCA • 1 ♀; same location as for preceding; Dr. J. Schouteden leg.; P. Vanderijst RMCA • 1 ♀; Lemfu (Madimba); 05°18’ S, 15°13’ E; Jan. 1945; Rév. P. De Beir leg.; RMCA. – **Lualaba**• 2 ♀♀; Kafakumba (Sandoa); 09°41’ S, 23°44’ E; Apr. 1933; F.G. Overlaet leg.; RMCA • 1 ♀; Kansenia; 10°19’ S, 26°04’ E; Oct. - Dec. 1930; R.P. Vankerckhoven leg.; RMCA • 1 ♀; Lualua: kapanga; 08°21’ S, 22°34’ E; Jan. 1933; F.G. Overlaet leg.; RMCA • 5 ♂♂; same location as for preceding; 7 Oct. 1932; F.G. Overlaet leg.; RMCA. – **Tanganyika** • 2 ♀♀; Vallée Lukuga; 05°40’ S, 26°55’ E; Nov. 1911; Dr. Schwetz leg.; RMCA. – **Tshopo** • 1 ♀; Utisongo (Stanleyville); 00°31’ N, 25°11’ E; 23 Apr. 1932; J. Vrydagh leg.; RMCA.

#### *Megachile* (*Callomegachile*) *rufipes* (Fabricius, 1781), *Chalicodoma* (*Callomegachile*) *rufipes* (Fabricius, 1781)

**Material.** Cupboard 110, Boxes 28, 29 and 30 (223♀♀ and 167♂♂). **D.R. CONGO.** – **Bas-Uele** • 1 ♀; Bambesa; 03°28’ N, 25°43’ E; 14 May 1938; J. Vrydagh leg.; RMCA • 1 ♀; same location as for preceding; 23 Jun. 1937; J. Vrydagh leg.; RMCA • 1 ♀; same location as for preceding; 9 May 1938; P. Henrard leg.; RMCA • 1 ♀; same location as for preceding; Feb. 1934; H.J. Brédo leg.; RMCA • 1 ♀; same location as for preceding; 6 Jul. 1937; J. Vrydagh leg.; RMCA • 1 ♀; Ibembo; 02°38’ N, 23°36’ E; Van Hecks leg.; RMCA • 1 ♂; Bambili (riv. Uele); 03°38’ N, 26°06’ E; Dr. J. Bequaert leg.; RMCA • 1 ♀; Mobwasa; 02°40’ N, 23°11’ E; Sep. 1911; R. Verschueren leg.; RMCA • 1 ♀; Uele: Dingila; 03°37’ N, 26°03’ E; 15 Jul. 1933; J.V. Leroy leg.; RMCA • 1 ♂; same location as for preceding; Aug. 1933; H.J. Brédo leg.; RMCA • 1 ♀; Uele: Ibembo; 02°38’ N, 23°36’ E; 1950; J. Sion leg.; RMCA • 1 ♀; same location as for preceding; 20 May 1950; R. Fr. Hutsebaut leg.; RMCA • 1 ♀; Uele-Poko; 03°09’ N, 26°53’ E; Degreef leg.; RMCA. – **Equateur** • 4 ♀♀; Basankusu; 01°13’ N, 19°49; 1949; Zusters O.L.V. ten Bunderen leg.; RMCA • 2 ♀♀; Coquilhatville (Mbandaka); 00°03’ N, 18°15’ E; 1946; Ch. Scops leg.; RMCA • 2 ♂♂; same location as for preceding; 1946; Ch. Scops leg.; RMCA • 1 ♂; same location as for preceding; 27 Aug. 1930; R. Verschueren leg.; RMCA • 2 ♀♀; Eala (Mbandaka); 00°03’ N, 18°19’ E; 1932; A. Corbisier leg.; RMCA • 3 ♂♂; same location as for preceding; 1932; A. Corbisier leg.; RMCA • 1 ♂; same location as for preceding; 14 Mar. 1933; A. Corbisier leg.; RMCA • 1 ♀; same location as for preceding; 15 Nov. 1929; H.J. Brédo leg.; RMCA • 11 ♀♀; same location as for preceding; 15-30 Nov. 1929; H.J. Brédo leg.; RMCA • 3 ♂♂; same location as for preceding; 15-30 Nov. 1929; H.J. Brédo leg.; RMCA • 5 ♀♀; same location as for preceding; 17 Nov. 1931; H.J. Brédo leg.; RMCA • 1 ♂; same location as for preceding; 17 Nov. 1931; H.J. Brédo leg.; RMCA • 1 ♀; same location as for preceding; 18 Nov. 1929; H.J. Brédo leg.; RMCA • 2 ♀♀; same location as for preceding; 21 Nov. 1931; H.J. Brédo leg.; RMCA • 4 ♂♂; same location as for preceding; 21 Nov. 1931; H.J. Brédo leg.; RMCA • 1 ♂; same location as for preceding; 22 Nov. 1931; H.J. Brédo leg.; RMCA • 1 ♂; same location as for preceding; 28 Oct. 1931; H.J. Brédo leg.; RMCA • 1 ♂; same location as for preceding; 28 Nov. 1929; H.J. Brédo leg.; RMCA • 3 ♀♀; same location as for preceding; 3 Oct. 1931; H.J. Brédo leg.; RMCA • 6 ♂♂; same location as for preceding; 4 Apr. 1932; H.J. Brédo leg.; RMCA • 2 ♀♀; same location as for preceding; 7 Jul. 1932; A. Corbisier leg.; RMCA • 24 ♂♂; same location as for preceding; Mar. 1932; H.J. Brédo leg.; RMCA • 1 ♀; same location as for preceding; Apr. 1932; H.J. Brédo leg.; RMCA • 3 ♀♀; same location as for preceding; Apr. 1933; A. Corbisier leg.; RMCA • 1 ♀; same location as for preceding; May 1932; H.J. Brédo leg.; RMCA • 3 ♂♂; same location as for preceding; May 1932; H.J. Brédo leg.; RMCA • 2 ♀♀; same location as for preceding; May 1935; J. Ghesquière leg.; RMCA • 26 ♀♀; same location as for preceding; Jun. 1932; A. Corbisier leg.; RMCA • 12♂♂; same location as for preceding; Jun. 1932; H.J. Brédo leg.; RMCA • 1 ♂; same location as for preceding; Jul. 1935; J. Ghesquière leg.; RMCA • 2 ♀♀; same location as for preceding; Nov. 1931; H.J. Brédo leg.; RMCA • 5 ♂♂; same location as for preceding; Nov. 1931; H.J. Brédo leg.; RMCA • 2 ♀♀; same location as for preceding; Nov. 1932; H.J. Brédo leg.; RMCA • 3 ♂♂; same location as for preceding; Nov. 1932; H.J. Brédo leg.; RMCA • 1 ♂; same location as for preceding; Nov. 1935; J. Ghesquière leg.; RMCA • 1 ♂; same location as for preceding; Dec. 1929; H.J. Brédo leg.; RMCA • 6 ♂♂; same location as for preceding; 4 Apr. 1932; H.J. Brédo leg.; RMCA • 1 ♀; Lukolela (Bikoro); 01°03’ S, 17°12’ E; 1951; R. Deguide leg.; RMCA • 1 ♂; Ubangi: Yakoma; 04°06’ N, 22°23’ E; 13 Nov. 1932; H.J. Brédo leg.; RMCA • 1 ♀; same location as for preceding; 15 Feb. 1931; H.J. Brédo leg.;

RMCA. – **Haut-Katanga** • 1 ♀; Elisabethville (Lubumbashi); 11°40’ S, 27°28’ E; 1935; Dr. Richard leg.; RMCA • 1 ♀; Kampombwe; 11°40’ S, 27°28’ E; Apr. 1931; H.J. Brédo leg.; RMCA • 1 ♀; Kisamba; 07°20’ S, 24°02’ E; Mar. 1955; Dr. J. Claessens leg.; RMCA • 1 ♀; Mwema; 08°13’ S, 27°28’ E; Jul. 1927; A. Bayet leg.; RMCA • 1 ♂; same location as for preceding; Jul. 1927; A. Bayet leg.; RMCA • 1 ♀; Penge-Bamboli; 01°20’ N, 28°09’ E; Jun. 1933; Putnam leg.; RMCA – **Haut-Lomami** • 3 ♀♀; Kulu-Mwanza; 07°54’ S, 26°45’ E; May 1927; A. Bayet leg.; RMCA • 1 ♀; Lualaba: Kabongo; 07°20’ S, 25°35’ E; 16 Nov. 1952; Ch. Seydel leg.; RMCA • 1 ♀; same location as for preceding; 27 Dec. 1952; Ch. Seydel leg.; RMCA • 1 ♀; same location as for preceding; 3 Jan. 1953; F.G. Overlaet leg.; RMCA • 1 ♀; same location as for preceding; 30 Dec. 1952; Ch. Seydel leg.; RMCA • 2 ♀♀; same location as for preceding; 31 Dec. 1952; Ch. Seydel leg.; RMCA • 1 ♂; same location as for preceding; 31 Dec. 1952; Ch. Seydel leg.; RMCA • 1 ♂; same location as for preceding; 7 Jan. 1953; Ch. Seydel leg.; RMCA • 1 ♀; PNU Munoi, bifure Lupiala; 08°50’ S, 26°44’E, alt. 890 m; 28 May-15 Jun. 1948; Mis. G.F. de Witte leg.; RMCA. – **Haut-Uele** • 1 ♀; Haut-Uele; 02°46’ N, 27°38’ E; Cotonco leg.; RMCA • 1 ♀; PNG [Parc National de Garamba]; 03°40’ N, 29°00’ E 8 Apr. 1952; H. De Saeger leg.; RMCA • 1 ♀; PNG Napokomweli; 03°40’ N, 29°00’ E; 18 Oct. 1950; H. De Saeger leg.; RMCA • 2 ♀♀; Uele: Dungu; 03°37’ N, 28°34’ E; Degreef leg.; RMCA • 1 ♀; Uele: Gangala-na-Bodio; 03°41’ N, 29°08’ E; Nov. 1956; Dr. M. Poll leg.; RMCA • 1 ♀; same location as for preceding; 1956; Dr. M. Poll leg.; RMCA • 1 ♀; Uele: Gingi; 03°34’ N, 25°39’ E; 3 Oct. 1913; R. Verschueren leg.; RMCA • 1 ♀; Uele: Niangara; 03°41’ N, 27°52’ E; Degreef leg.; RMCA • 1 ♀; Uele: Suranga; 04°06’N 22°23’E; Degreef leg.; RMCA. – **Ituri** • 1 ♀; Ituri: Arara-Aru; 02°52’ N, 30°52’ E; 10 Apr. 1952; M. Winand leg.; RMCA • 1 ♀; Ituri: Faradje; 03°44’ N, 29°42’ E; 1944; J. Lisfrane leg.; RMCA • 1 ♀; Kibali-Ituri: Demu; 03°37’ N, 28°34’ E; Feb. – Mar. 1936; Dr. J. Pasteels leg.; RMCA • 1 ♀; Kibali-Ituri: Kilomines; 01°50’ N, 30°09’ E; May 1957; C. Smoor leg.; RMCA • 1 ♀; Kilo: Kere-Kere; 02°42’ N, 30°33’ E; 18 mar. 1950; Dr. Turco leg.; RMCA • 1 ♂; Lac Albert: Kasenyi; 01°24’ N, 30°26’ E; 5 May. 1935; R. Verschueren leg.; RMCA. – **Kasaï** • 2 ♀♀; Luebo; 05°20’ S, 21°24’ E; Jan. – Jul. 1958; F. François leg.; RMCA • 1 ♀; Tshikapa; 06°24’ S, 20°48’ E; Apr. 1939; Mevr. Bequaert leg.; RMCA • 5 ♂♂; same location as for preceding; Apr. 1939; Mevr. Bequaert leg.; RMCA. – **Kasaï Central** • 1 ♂; Luluabourg (Kananga); 05°54’ S, 21°52’ E; 30 Jan. 1963; Jan Deheegher leg.; RMCA • 1 ♀; same location as for preceding; P. Callewaert leg.; RMCA • 1 ♂; same location as for preceding; P. Callewaert leg.; RMCA • 1 ♀; Shenateke; 05°54’ S, 22°25’ E; 27 Jun. 1946; V. Lagae leg.; RMCA. – **Kasaï Oriental** • 1 ♂; Sankuru: Bakwanga (Dibindi); 06°08’ S, 23°38’ E; 18 Jun. 1939; Mevr. Bequaert leg.; RMCA • 1 ♀; same location as for preceding; Jun. 1939; Mevr. Bequaert leg.; RMCA. – **Kinshasa** • 1 ♀; Kalina; 04°18’ S, 15°17’ E; Aug. 1945; Mme. Delsaut leg.; RMCA • 1 ♂; same location as for preceding; Aug. 1945; Mme. Delsaut leg.; RMCA • 1 ♀; Léopoldville (Kinshasa); 04°19’ S, 15°19’ E; 1911; Dr. Mouchet leg.; RMCA • 1 ♀; same location as for preceding; 11 Apr. 1947; Dr. E. Dartevelle leg.; RMCA • 1 ♂; same location as for preceding; 18 Sep. 1910; R. Verschueren leg.; RMCA • 1 ♂; same location as for preceding; Dr. Houssiaux leg.; RMCA • 2 ♂♂; same location as for preceding; Aug. 1949; Dr. A. Dubois leg.;

RMCA. – **Kongo Central** • 1 ♂; Bangu (Mbanza Ngungu); 05°15’S, 14°30’ E; Aug. 1913; R. Verschueren leg.; RMCA • 1 ♀; Boma; 05°51’ S, 13°03’ E; 1935; W. Moreels leg.; RMCA • 13 ♂♂; same location as for preceding; 1937; Dr. Schlesser leg.; RMCA • 1 ♂; same location as for preceding; 7 Nov. 1913; Lt. Styczynski leg.; RMCA • 2 ♀♀; same location as for preceding; 8 Jul. 1920; H. Schouteden leg.; RMCA • 1 ♀; same location as for preceding; Nov. 1950; I. Mesmaekers leg.; RMCA • 1 ♀; same location as for preceding; Dec. 1950; I. Mesmaekers leg.; RMCA • 1 ♂; Kisantu (Madimba); 05°08’ S, 15°06’ E; 1927; R. P. Vanderyst leg.; RMCA • 13 ♂♂; same location as for preceding; 1931; R. P. Vanderyst leg.; RMCA • 5 ♂♂; same location as for preceding; 1932; R. P. Vanderyst leg.; RMCA • 1♂; same location as for preceding; 1 Apr. 1930; R. P. Vanderyst leg.; RMCA • 1 ♂; same location as for preceding; 17 Nov. 1932; R. P. Vanderyst leg.; RMCA • 4 ♂♂; same location as for preceding; Jan. – Apr. 1930; R. P. Vanderyst leg.; RMCA • 1 ♀; same location as for preceding; May 1945; Rév. Fr. Anastase leg.; RMCA • 1 ♀; same location as for preceding; Jun. 1921; P. Van Wing leg.; RMCA • 7 ♂♂; same location as for preceding; Dec. 1927; R. P. Vanderyst leg.; RMCA • 1 ♀; Kitobola; 05°22’ S, 14°31’ E; 1911; Rovere leg.; RMCA • 1 ♀; Lemfu (Madimba); 05°18’ S, 15°13’ E; Feb. 1945; Rév. P. De Beir leg.; RMCA • 1 ♂; same location as for preceding; Jun. 1945; Rév. P. De Beir leg.; RMCA • 1 ♀; same location as for preceding; Oct. – Dec. 1944; Rév. P. De Beir leg.; RMCA • 2 ♂♂; Mayidi (Madimba); 05°11’ S, 15°09’ E; 1942; Rév. P. Van Eyen leg.; RMCA • 1 ♀; Moanda; 05°56’ S, 12°21’ E; 28 Apr. 1970; P.M. Elsen leg.; RMCA • 1 ♀; Thysville (Mbanza Ngungu); 05°15’ S, 14°52’ E; 1948; L. Salathiel leg.; RMCA • 1 ♀; same location as for preceding; Jan. 1953; J. Sion leg.; RMCA • 1 ♂; same location as for preceding; Jan. 1953; J. Sion leg.; RMCA. – **Kwilu** • 1 ♀; Kolo-Kwilu-Madiata; 05°27’ S, 14°52’ E; Sep. 1913; R. Verschueren leg.; RMCA • 2 ♀♀; Leverville (Bulungu); 04°50’ S, 18°44’ E; 1928; Mme. J. Tinant leg.; RMCA • 1 ♀; Moyen Kwilu: Leverville (Bulungu); 04°50’ S, 18°44’ E; Jan. 1914; P. Vanderijst leg.; RMCA. – **Lomami** • 1 ♀; Kabinda; 06°08’ S, 24°29’ E; 1945-1946; Dr. Hautmann leg.; RMCA • 1 ♀; same location as for preceding; Feb. 1933; Mme. Gillardin n leg.; RMCA • 1 ♂; Tshofa; 05°14’ S, 25°15’ E; Dec. 1934; Mme. Gillardin leg.; RMCA. – **Lualaba** • 1 ♂; Kapanga; 08°21’ S, 22°34’ E; Dec. 1932; R. Verschueren leg.; RMCA • 1 ♀; Lulua: Kapanga; 08°21’ S, 22°34’ E; Apr. 1933; F.G. Overlaet leg.; RMCA • 1 ♀; same location as for preceding; Sep. 1932; F.G. Overlaet leg.; RMCA • 4 ♂♂; same location as for preceding; Oct. 1932; F.G. Overlaet leg.; RMCA • 2 ♀♀; same location as for preceding; Nov. 1932; F.G. Overlaet leg.;

RMCA. – **Maï-Ndombe** • 1 ♀; Bolobo (Mushie); 02°10’ S, 16°14’ E; 27 Jul. 1912; Dr. Mouchet leg.; RMCA • 1 ♀; Bolobo: Makamandelo; 02°10’ S, 16°14’ E; 1938; Dr. H. Schouteden leg.; RMCA • 1 ♂; Bomboma; 02°24’ N, 18°53 E; 21 Jul. 1935; A. Bal leg.; RMCA • 2 ♀♀; Kwamouth (Mushie); 03°11’ S, 16°12’ E; 20 Jun. 1913; Dr. J. Maes leg.; RMCA • 2 ♀♀; Lac Léopold II: Bokalakala; 02°00’ S, 18°15’ E; 1957; N’Kele leg.; RMCA • 1 ♀; Yumbi; 01°54’ S, 16°33’ E; 29 Jul. 1912; Dr. Mouchet leg.; RMCA. – **Maniema** • 1 ♀; Kasongo; 04°27’ S, 26°40’ E; Sep. 1959; R. Verschueren leg.; RMCA • 1 ♀; Kasongo: Kahuta; 04°27’ S, 26°40’ E; Sep. 1959; P.L.G. Benoit leg.; RMCA • 1 ♀; Kil. 345 de Kindu; 02°57’ S, 25°55’ E; Dr. Russo leg.; RMCA • 1 ♀; Malela; 04°22’ S, 26°08’ E; Jan. 1914; L. Burgeon leg.; RMCA • 1 ♀; Maniema; 05°00’ S, 29°00’ E; 1936; Guraut leg.; RMCA • 1 ♀; Nyangwe; 04°13’ S, 26°11’ E; 13 May 1918; R. Mayné leg.; RMCA • 2 ♂♂; same location as for preceding; Apr. – May 1918; R. Mayné leg.; RMCA • 1 ♀; Terr. De Kasongo, Riv. Lupaya; 04°27’ S, 26°40’ E; 8 Feb. 1960; P.L.G. Benoit leg.; RMCA • 1 ♀; Terr. de Kasongo, Riv. Lumami; 04°27’ S, 26°40’ E; Feb. 1960; P.L.G. Benoit leg.; RMCA. • 2 ♀♀; Vieux Kassongo (Kasongo); 04°27’ S, 26°40’ E; Dec. 1910; Dr. J. Bequaert leg.; RMCA. – **Mongala** • 1♂; Binga (Lisala); 02°28’ N, 20°31’ E; 1 Mar. 1932; H.J. Brédo leg.; RMCA • 1 ♀; Terr. Lisala; 02°09’ N, 21°30’ E; Mar. 1937; C. Leontovitch leg.; RMCA • 4 ♀♀; Ubangi: Binga (Lisala); 02°28’ N, 20°31’ E; 5 – 12. Mar. 1932; H.J. Brédo leg.; RMCA • 1 ♀; Yambata; 02°26’ N, 22°02’ E; 10 Dec. 1912; R. Mayné leg.; RMCA. – **Nord-Ubangi** • 2 ♀♀; Ubangi: Bosobolo; 04°11’ N, 19°53’ E; 8 – 11 Jan. 1932; H.J. Brédo leg.; RMCA • 1 ♂; same location as for preceding; 8-11 Jan. 1932; H.J. Brédo leg.; RMCA • 1 ♀; Ubangi: Karawa (Businga); 03°21’ N, 20°18’ E; 1936; Rév. Wallin leg.; RMCA • 2 ♀♀; Yakoma; 04°06’ N, 22°23’ E; 4 – 17 Feb. 1932; H.J. Brédo leg.; RMCA • 2 ♀♀; same location as for preceding; 5 – 17 Feb. 1932; H.J. Brédo leg.; RMCA • 1 ♂; same location as for preceding; 5 – 17 Feb. 1932; H.J. Brédo leg.; RMCA. – **Sankuru** • 1 ♀; Sankuru: Tshumbe S. Marie; 04°02’ S, 22°41’ E; May 1948; R. Verschueren leg.; RMCA. – **Sud-Kivu** • 1 ♀; Lolo; 02°16’ S, 27°43’ E; 23 May 1925; Dr. Rodhain leg.; RMCA • 1 ♀; Rég. Des Grands Lacs; 02°29’ S, 28°51’ E; Dr. Sagona leg.; RMCA. – **Sud-Ubangi** • 1 ♀; Kunungu (N’Kele); 02°45’ N, 18°26’ E; 1934; Dr. H. Schouteden leg.; RMCA • 1 ♀; same location as for preceding; 1937; Dr. H. Schouteden leg.; RMCA • 4 ♀♀; Ubangi: Boma-Motenge; 03°15’ N, 18°39’ E; 22 Dec. 1931; H.J. Brédo leg.; RMCA • 2 ♀♀; same location as for preceding; 5 Dec. 1931; H.J. Brédo leg.;

RMCA. – **Tanganyika** • 1 ♂; Katanga: Kabinda (Moba); 07°02’ S, 27°47’ E; Jan. 1926; Ch. Seydel leg.; RMCA • 1 ♀; Kiambi (Manono); 07°19’ S, 28°01’ E; 23 Apr. 1931; G.F. de Witte leg.; RMCA • 1 ♀; same location as for preceding; 26 Apr. 1931; G.F. de Witte leg.; RMCA • 2 ♀♀; same location as for preceding; 5 – 15 May 1931; G.F. de Witte leg.; RMCA • 1 ♀; Kiambi-Baudouinville; 07°19’ S, 28°01’ E; 23 Apr. 1931; R. Verschueren leg.; RMCA • 4 ♀♀; Lukulu (Manono); 07°08’ S, 28°06’ E; 1932; Mr. Ramade leg.; RMCA • • 1 ♀; Lulua: Kalenge; 06°33’ S, 26°35’ E; Feb. 1934; F.G. Overlaet leg.; RMCA • 1 ♂; Mato; 06°07’ S, 26°15’ E; 8 Dec. 1925; Ch. Seydel leg.; RMCA • 1 ♀; same location as for preceding; Jan. 1926; Ch. Seydel leg.; RMCA • 3 ♀♀; Tanganyika: Mpala; 06°45’ S, 29°31’ E, alt. 780 m; Jan. – Feb. 1934; H. Bomans leg.; RMCA • 4 ♀♀; Tanganyika-Moero: Nyunzu; 05°57’ S, 28°01’ E; Jan. – Feb. 1934; H. Bomans leg.; RMCA • 1 ♀; Vallée Lukuga; 05°40’ S, 26°55’ E; Nov. 1911; Dr. Schwetz leg.; RMCA. – **Tshopo** • 1 ♀; Basoko; 01°13’ N, 23°36’ E; May 1948; P.L.G. Benoit leg.; RMCA • 1 ♂; same location as for preceding; May 1948; P.L.G. Benoit leg.; RMCA • 1 ♀; Elisabetha (Yahuma); 01°09’ N, 23°37’ E; Mme. J. Tinant leg.; RMCA • 1 ♀; Miss. St. Gabriel; 00°33’ N, 25°05’ E; M. Torley leg.; RMCA • 1 ♂; Ponthierville (Ubundu); 00°22’ S, 25°29’ E; 22 Oct. 1910; Dr. J. Bequaert leg.; RMCA • 1 ♀; Stanleyville (Kisangani); 00°31’ N, 25°11’ E; 10 Feb. 1928; A. Collart leg.; RMCA • 2 ♀♀; same location as for preceding; 10 – 13 Sep. 1928; A. Collart leg.; RMCA • 2 ♂♂; same location as for preceding; 13 – 23 Aug. 1923 Sep. 1928; A. Collart leg.; RMCA • 1 ♀; same location as for preceding; 2 Aug. 1932; J. Vrydagh leg.; RMCA • 1 ♀; same location as for preceding; 5 Feb. 1928; A. Collart leg.; RMCA. – **Tshuapa** • 1 ♀; Bamania (Mbandaka); 00°01’ N, 18°19’ E; Feb. – Mar. 1958; Rév. P. Hulstaert leg.; RMCA • 1 ♀; Boende; 00°13’ S, 20°52’ E; Jan. 1952; Rév. P. Lootens leg.; RMCA • 1 ♀; Bokuma; 00°06’ S, 18°41’ E; 2 Sep. 1934; R. Verschueren leg.; RMCA • 1 ♂; same location as for preceding; 2 Sep. 1934; R. Verschueren leg.; RMCA • 1 ♂; same location as for preceding; Aug. 1952; R. Verschueren leg.; RMCA • 1 ♀; Tshuapa: Bamanya; 00°01’ N, 18°19’ E; 1968; Rév. P. Hulstaert leg.; RMCA • 1 ♂; Tshuapa: Bokuma; 00°06’ S, 18°41’ E; Feb. 1954; Rév. P. Lootens leg.; RMCA • 1 ♀; same location as for preceding; Mar. 1952; Rév. P. Lootens leg.; RMCA • 1 ♀; same location as for preceding; Mar. 1954; Rév. P. Lootens leg.; RMCA • 1 ♀; Tshuapa: Bokungu; 00°41’ S, 22°19’ E; 1949; Dupuis leg.; RMCA • 1 ♀; Tshuapa: Flandria (Ingende); 00°20’ S, 19°06’ E; 4 May 1946; R. Verschueren leg.; RMCA • 2 ♀♀; same location as for preceding; Jan. – Feb. 1948; Zusters O.L.V. ten Bunderen leg.; RMCA.

#### *Megachile* (*Carinula*) *decemsignata* Rodoszkowski, 1981, *Chalicodoma* (*Carinella*) *decemsignata* (Rodoszkowski, 1981)

**Material.** Cupboard 110, Box 24 (119♀♀ and 39♂♂). **D.R. CONGO.** – **Bas-Uele** • 1 ♀; Bambesa; 03°28’ N, 25°43’ E; Feb. 1934; H.J. Brédo leg.; RMCA • 1 ♀; La Kulu; 07°08’ N, 28°06’ E; 25 Nov. 1930; J. Vrydagh leg.; RMCA • 1 ♀; Uele; 04°06’ N, 22°23’ E; Degreef leg.; RMCA. – **Equateur** • 1 ♂; Coquilhatville (Mbandaka); 00°03’ N, 18°15’ E; 1946; Ch. Scops leg.; RMCA • 1 ♂; Eala (Mbandaka); 00°03’ N, 18°19’ E; 1932; A. Corbisier leg.; RMCA • 2 ♂♂; same location as for preceding; 19 Oct. 1931; H.J. Brédo leg.; RMCA • 1 ♀; same location as for preceding; 20 Apr. 1933; A. Corbisier leg.; RMCA • 1♂; same location as for preceding; 22 Nov. 1931; H.J. Brédo leg.; RMCA • 1 ♂; same location as for preceding; 4 Apr. 1932; H.J. Brédo leg.; RMCA • 5 ♀♀; same location as for preceding; Mar. 1932; H.J. Brédo leg.; RMCA • 4 ♂♂; same location as for preceding; Mar. 1932; H.J. Brédo leg.; RMCA • 1 ♀; same location as for preceding; Apr. 1932; H.J. Brédo leg.; RMCA • 2 ♂♂; same location as for preceding; Apr. 1932; H.J. Brédo leg.; RMCA • 3 ♂♂; same location as for preceding; Apr. 1933; A. Corbisier leg.; RMCA • 3 ♂♂; same location as for preceding; May 1933; A. Corbisier leg.; RMCA • 1 ♀; same location as for preceding; May 1935; J. Ghesquière leg.; RMCA • 1 ♂; same location as for preceding; Nov. 1934; J. Ghesquière leg.; RMCA • 8 ♀♀; same location as for preceding; Jun. 1932; A. Corbisier leg.; RMCA • 1♂; same location as for preceding; Jun. 1932; A. Corbisier leg.; RMCA • 1 ♀; same location as for preceding; Nov. 1931; H.J. Brédo leg.; RMCA • 1 ♀; same location as for preceding; Dec. 1931; H.J. Brédo leg.; RMCA • 1 ♀; Modu (Wamba); 00°36’ N, 20°07’ E; 3 Jun. 1931; H.J. Brédo leg.; RMCA • 1 ♂; same location as for preceding; 5 Jun. 1931; H.J. Brédo leg.; RMCA • 19 ♀♀; Ubangi: Mokolo; 01°35’ N, 29°05’ E; 5 Dec. 1931; H.J. Brédo leg.; RMCA. – **Haut-Lomami** • 1 ♀; PNU Mabwe (R.P. Lac Upemba); 08°47’ S, 26°52’ E, alt. 585 m; 1-11 Jan. 1949; Mis. G.F. de Witte leg.; RMCA • 1 ♀; same location as for preceding; 20-26 Jan. 1949; Mis. G.F. de Witte leg.; RMCA. – **Haut-Uele** • 1 ♀; Haut-Uele: Mauda; 04°46’ N, 27°18’ E; Mar. 1925; H. De Saeger leg.; RMCA • 1 ♀; PNG [Parc National de Garamba]; 03°40’ N, 29°00’ E; 9 Nov. 1950; Dr. J. Schouteden leg.; RMCA • 1 ♂; PNG Nagero (Dungu); 03°45’ N, 29°31’ E; 1 – 30 Jun. 1950; H. De Saeger leg.; RMCA. – **Ituri** • 7 ♀♀; Haut-Ituri, 29°03’, 29°03’ E; 1906; H. Wilmin leg.; RMCA. – **Kasaï** • 1 ♀; Kasaï; 03°11’ S, 16°56’ E; L. Achten leg.; RMCA. • 1 ♂; Luebo rivière; 05°20’ S, 21°24’ E; 25 May 1938; Mevr. Bequaert leg.; RMCA • 1 ♀; same location as for preceding; 25 May 1939; Mevr. Bequaert leg.; RMCA • 1 ♀; Lutzkwadi (Ilebo); 04°20’ S, 20°36’ E; 17 Jun. 1946; V. Lagae leg.;

RMCA. – **Kongo Central** • 1 ♀; Forêt de la Lindi; 05°45’ S, 12°27’ E; 1939; Van den Hirtz leg.; RMCA • 1 ♂; Matadi; 05°49’ S, 13°28’ E; 24 Apr. 1946; Dr. Schlesser leg.; RMCA • 2 ♀♀; Mayidi (Madimba); 05°11’ S, 15°09’ E; 1945; Rév. P. Van Eyen leg.; RMCA. – **Kwilu** • 1 ♀; Leverville (Bulungu); 04°50’ S, 18°44’ E; 1928; Mme. J. Tinant leg.; RMCA. – **Lualaba** • 1 ♀; Kapanga; 08°21’ S, 22°34’ E, Jan. 1933; R.P. Vankerckhoven leg.; RMCA • 1 ♀; Lulua: Kapanga; 08°21’ S, 22°34’ E; Mar. 1933; F.G. Overlaet leg.; RMCA • 1 ♂; same location as for preceding; Sep. 1932; F.G. Overlaet leg.; RMCA • 1 ♂; same location as for preceding; Oct. 1932; F.G. Overlaet leg.; RMCA • 4 ♀♀; same location as for preceding; Dec. 1932; F.G. Overlaet leg.; RMCA • 2 ♂♂; same location as for preceding; Dec. 1932; F.G. Overlaet leg.; RMCA • 1 ♀; Sandoa à Kapanga; 09°41’ S, 22°53’ E; 1929; Dr. Walker leg.; RMCA. – **Maniema** • 2 ♀♀; Kibombo; 03°54’ S, 25°55’ E; 10 Nov. 1910; Dr. J. Bequaert leg.; RMCA • 1 ♀; same location as for preceding; 31 Nov. 1910; Dr. J. Bequaert leg.; RMCA • 1 ♀; Lokandu; 02°31’ S, 25°47’ E; 1939; Lt. Vissers leg.; RMCA • 1 ♀; Malela; 04°22’ S, 26°08’ E; Jan.-Feb. 1913; R. Verschueren leg.; RMCA. – **Mongala** • 1 ♂; Lisala; 02°09’ N, 21°30’ E; 1937; Vermeiren leg.; RMCA. – **Nord-Kivu** • 1 ♀; Beni à Lesse; 00°45’ N, 29°46’ E; Jul. 1911; Dr. Murtula leg.; RMCA. – **Nord-Ubangi** • 1 ♀; Ubangi: Karawa (Businga); 03°21’ N, 20°18’ E; 1937; Rév. Wallin leg.; RMCA • 1 ♀; Ubangi: Mokolo; 01°35’ N, 29°05’ E; 5 Dec. 1931; H.J. Brédo leg.; RMCA. – **Sankuru** • 1 ♀; Sankuru: Komi; 03°23’ S, 23°46’ E; Apr. 1930; J. Ghesquière leg.; RMCA • 2 ♀♀; same location as for preceding; May 1930; J. Ghesquière leg.; RMCA • 3 ♀♀; same location as for preceding; Jun. 1930; J. Ghesquière leg.; RMCA • 4 ♂♂; same location as for preceding; Jun. 1930; J. Ghesquière leg.; RMCA • 1 ♀; same location as for preceding; Jul. 1930; J. Ghesquière leg.; RMCA • 1 ♂; same location as for preceding; Jul. 1930; J. Ghesquière leg.; RMCA. – **Nord-ubangi** • 1 ♂; Kunungu (Nkele); 02°45’ N, 18°26’ E; 1938; Dr. H. Schouteden leg.; RMCA • 12 ♀♀; Ubangi: Boma-Motenge; 03°15’ N, 18°39’ E; Dec. 1931; H.J. Brédo leg.; RMCA • 1 ♀; Ubangi: Nzali; 03°15’ N, 19°47’ E; 3-4 Feb. 1932; H.J. Brédo leg.; RMCA. – **Tanganyika** • 1 ♀; Lulua: Kalenge; 06°33’ S, 26°35’ E; Feb. 1934; F.G. Overlaet leg.; RMCA. – **Tshopo** • 1 ♀; Miss. St. Gabriel; 00°33’ N, 25°05’ E; M. Torley leg.; RMCA • 1 ♀; Ponthierville (Ubundu); 00°22’ S, 25°29’ E; 22 Oct. 1910; Dr. J. Bequaert leg.; RMCA • 1 ♂; same location as for preceding; 22 Oct. 1910; Dr. J. Bequaert leg.; RMCA • 1 ♂; Stanleyville (Kisangani); 00°31’ N, 25°11’ E; Apr. 1915; Lang & Chapin leg.; RMCA • 1 ♀; Terr. de Basoko (forêt); 01°13’ N, 23°36’ E; 1950; R.P. Camps leg.; RMCA • 1 ♀; Yangambi; 00°46’ N, 24°27’ E; 12 Dec. 1936; Lt. J. Ghesquière leg.;

RMCA. – **Tshuapa** • 1 ♀; Bamania (Mbandaka); 00°01’ N, 18°19’ E; Sep.-Nov. 1958; A. Corbisier leg.; RMCA • 1 ♀; Boende; 00°13’ S, 20°52’ E; Jan. 1952; Rév. P. Lootens leg.; RMCA • 1 ♀; Tshuapa: Bamanya; 00°01’ N, 18°19’ E; 1968; Rév. P. Hulstaert leg.; RMCA • 3 ♀♀; Tshuapa: Bokuma; 00°06’ S, 18°41’ E; 1953; Rév. P. Lootens leg.; RMCA • 1 ♀; Bokuma; 00°06’ S, 18°41’ E; May 1958; Rév. P. Hulstaert leg.; RMCA • 1 ♀; same location as for preceding; 00°06’ S, 18°41’ E; Mar. 1952; Rév. P. Lootens leg.; RMCA • 1 ♂; same location as for preceding; Jan.-Feb. 1952; Rév. P. Lootens leg.; RMCA • 11 ♀♀; same location as for preceding; Jul. 1952; Rév. P. Lootens leg.; RMCA • 1 ♀; same location as for preceding; Dec. 1951; Rév. P. Lootens leg.; RMCA.

#### *Megachile* (*Carinula*) *fervida* Smith, 1853, *Chalicodoma* (*Carinella*) *fervida* (Smith, 1853)

**Material.** Cupboard 110, Box 25 (1♀). **D.R. CONGO.** – **Haut-Katanga** • 1 ♀; Elisabethville (Lubumbashi); riv. Kimilolo; 11°40’ S, 27°28’ E; 6 Nov. 1920; Dr. M. Bequaert leg.; RMCA.

#### *Megachile* (*Carinula*) *junodi* Friese, 1904, *Chalicodoma* (*Carinella*) *junodi* (Friese, 1904)

**Material.** Cupboard 110, Box 25 (1♀ and 2♂♂). **D.R. CONGO.** – **Haut-Katanga** • 1 ♀; Elisabethville (Lubumbashi); 11°40’ S, 27°28’ E; 12 Oct. 1934; P. Quarré leg.; RMCA • 1 ♂; PNU Masombwe s. Grde. Kafwe; 09°05’ S, 27°12’ E, alt. 1120 m; 4 – 16 Jan. 1948; Mis. G.F. de Witte leg.; RMCA. – **Haut-Lomami** • 1 ♂; Sankisia; 09°24’ S, 25°48’ E; 4 Sep. 1911; Dr. J. Bequaert leg.; RMCA.

#### *Megachile* (*Carinula*) *silverlocki* Meade-Waldo, 1913, *Chalicodoma* (*Carinella*) *silverlocki* (Meade-Waldo, 1913)

**Material.** Cupboard 110, Box 25 (8♀♀ and 2♂♂). **D.R. CONGO.** Paratypes – **Haut-Lomami** • 1 ♂; Bukama; 09°12’ S, 25°51’ E; 10 Jun. 1911; Dr. M. Bequaert leg.; RMCA • 1 ♀; same location as for preceding; 16 Apr. 1911; Dr. J. Bequaert leg.; RMCA • 2 ♀♀; same location as for preceding; 21 Apr. 1911; Dr. J. Bequaert leg.; RMCA • 1 ♀; same location as for preceding; 23 Apr. 1911; Dr. J. Bequaert leg.; RMCA • 1 ♀; same location as for preceding; 7 Jul. 1911; Dr. J. Bequaert leg.; RMCA. • 1 ♀; Bukama; 09°12’ S, 25°51’ E; 12 Sep. 1911; Dr. J. Bequaert leg.; RMCA • 1 ♂; same location as for preceding; 12 Sep. 1911; Dr. J. Bequaert leg.; RMCA • 2 ♀♀; same location as for preceding; 4 sep. 1911; Dr. J. Bequaert leg.; RMCA.

#### *Megachile* (*Carinula*) *torrida* Smith, 1853, *Chalicodoma* (*Carinella*) *torrida* (Smith, 1853)

**Material.** Cupboard 110, Box 25 (8♀♀ and 2♂♂). **D.R. CONGO.** – **Equateur** • 1 ♂; Beneden-Congo: Tumba; 00°50’ S, 18°00’ E; Sep. 1938; Mevr. Bequaert leg.; RMCA • 3 ♀♀; Eala (Mbandaka); 00°03’ N, 18°19’ E; 1932; A. Corbisier leg.; RMCA • 1 ♂; same location as for preceding; 14 Mar. 1933; A. Corbisier leg.; RMCA • 1 ♂; same location as for preceding; 14 Nov. 1931; H.J. Brédo leg.; RMCA • 1 ♀; same location as for preceding; 17 Nov. 1931; H.J. Brédo leg.; RMCA • 6 ♀♀; same location as for preceding; 21 Nov. 1931; H.J. Brédo leg.; RMCA • 1 ♀; same location as for preceding; 22 Apr. 1932; H.J. Brédo leg.; RMCA • 1 ♀; same location as for preceding; 22 Nov. 1931; H.J. Brédo leg.; RMCA • 3 ♀♀; same location as for preceding; 28 Nov. 1929; H.J. Brédo leg.; RMCA • 1 ♂; same location as for preceding; 28 Nov. 1929; H.J. Brédo leg.; RMCA • 2 ♀♀; same location as for preceding; 4 Apr. 1932; H.J. Brédo leg.; RMCA • 1 ♀; same location as for preceding; 5 Dec. 1931; H.J. Brédo leg.; RMCA • 1 ♀; same location as for preceding; Mar. 1931; H.J. Brédo leg.; RMCA • 21 ♀♀; same location as for preceding; Mar 1932; H.J. Brédo leg.; RMCA • 4 ♂♂; same location as for preceding; Mar 1932; H.J. Brédo leg.; RMCA • 1 ♀; same location as for preceding; Mar 1935; J. Ghesquière leg.; RMCA • 1 ♂; same location as for preceding; Mar 1935; J. Ghesquière leg.; RMCA • 6 ♀♀; same location as for preceding; Apr. 1932; H.J. Brédo leg.; RMCA • 1 ♀; same location as for preceding; Apr. 1933; A. Corbisier leg.; RMCA • 9 ♀♀; same location as for preceding; May 1932; H.J. Brédo leg.; RMCA • 1 ♂; same location as for preceding; May 1932; H.J. Brédo leg.; RMCA • 11 ♀♀; same location as for preceding; Jun. 1932; A. Corbisier leg.; RMCA • 5 ♂♂; same location as for preceding; Jun. 1932; A. Corbisier leg.; RMCA • 7 ♀♀; same location as for preceding; Nov. 1931; H.J. Brédo leg.; RMCA • 2 ♂♂; same location as for preceding; Nov. 1931; H.J. Brédo leg.; RMCA • 5 ♀♀; same location as for preceding; Nov. 1932; A. Corbisier leg.; RMCA • 1 ♀; same location as for preceding; Nov. 1932; A. Corbisier leg.; RMCA • 1 ♀; same location as for preceding; 20 Oct. 1931; H.J. Brédo leg.;

RMCA. – **Haut-Katanga** • 1 ♀; Elisabethville (Lubumbashi); 11°40’ S, 27°28’ E; 1933; De loose leg.; RMCA • 1 ♀; same location as for preceding; 12 Apr. 1926; Dr. M. Bequaert leg.; RMCA • 2♂♂; same location as for preceding; 12 Apr. 1912; Dr. J. Bequaert leg.; RMCA • 2 ♂♂; same location as for preceding; 14 Nov. 1934; Dr. M. Bequaert leg.; RMCA • 1 ♀; same location as for preceding; 17 Sep. 1934; P. Quarré leg.; RMCA • 1 ♀; same location as for preceding; 14 Nov. 1934; P. Quarré leg.; RMCA • 1 ♀; same location as for preceding; 17 Nov. 1934; P. Quarré leg.; RMCA • 1 ♀; same location as for preceding; Feb. 1933; De Loose leg.; RMCA • 1 ♀; same location as for preceding; Feb. 1933; Dr. M. Bequaert leg.; RMCA • 2 ♀♀; same location as for preceding; Feb. 1935; P. Quarré leg.; RMCA • 1 ♂; same location as for preceding; Feb. 1938; H.J. Brédo leg.; RMCA • 1 ♀; same location as for preceding; Sep. 1926; Ch. Seydel leg.; RMCA • 1 ♀; same location as for preceding; De Loose leg.; RMCA • 3 ♂♂; same location as for preceding; De Loose leg.; RMCA • 1 ♀; same location as for preceding; Nov. 1924; P. Quarré leg.; RMCA • 1 ♂; Katumba; 08°32’ S, 25°59’ E; Oct. 1907; Dr. J. Sheffield-Neave leg.; RMCA^5^ • 1 ♀; Lubumbashi; 11°40’ S, 27°28’ E; 22 Mar. 1920; Dr. M. Bequaert leg.; RMCA • 1 ♂; same location as for preceding; 23 May 1920; Dr. M. Bequaert leg.; RMCA • 1 ♀; Mwema; 08°13’ S, 27°28’ E; Jul. 1927; A. Bayet leg.; RMCA • 1 ♂; PNU Lusinga (riv. Kamitungulu); 08°55’ S, 08°55’ S; 16 Apr. 1948; Mis. G.F. de Witte leg.; RMCA • 2 ♀♀; PNU Masombwe s. Grde. Kafwe; 09°05’ S, 27°12’ E, alt. 1120 m; 16 Apr. 1948; Mis. G.F. de Witte leg.; RMCA. – **Haut-Lomami** • 1 ♀; Bukama; 09°12’ S, 25°51’ E; 1947; B. Dewit leg.; RMCA • 1 ♀; same location as for preceding; 21 Apr. 1911; Dr. J. Bequaert leg.; RMCA • 1 ♂; same location as for preceding; 7 Jul. 1911; Dr. J. Bequaert leg.; RMCA • 1 ♀; Lomami: Kishinde; 08°44’ S, 25°00’ E; Sep. 1931; P. Quarré leg.; RMCA • 1 ♂; Lualaba: Kabongo; 07°20’ S, 25°35’ E; Nov. 1953; Ch. Seydel leg.; RMCA • 1 ♂; same location as for preceding; 27 Dec. 1952; Mme. L. Gilbert leg.; RMCA • 1♂; Luashi; 07°31’ S, 24°10’ E; 1935; Freyne leg.; RMCA • 1 ♀; PNU Mabwe (R.P. Lac Upemba; 585m); 08°39’ S, 26°31’ E; 2 Mar. 1949; Mis. G.F. de Witte leg.; RMCA • 1 ♀; same location as for preceding; 6 Mar. 1949; Mis. G.F. de Witte leg.; RMCA • 1 ♂; Sankisia; 09°24’ S, 25°48’ E; 1 Sep. 1911; Dr. J. Bequaert leg.; RMCA • 2 ♂♂; same location as for preceding; 29 Aug. 1911; Dr. J. Bequaert leg.; RMCA • 1 ♀; same location as for preceding; 3 Sep. 1911; Dr. J. Bequaert leg.; RMCA • 1 ♂; same location as for preceding; 3 Sep. 1911; Dr. J. Bequaert leg.; RMCA. – **Haut-Uele** • 1 ♂; Haut-Uele: Paulis; 02°46’ N, 27°38’ E; Aug. 1947; P.L.G. Benoit leg.; RMCA • 1 ♀; PNG [Parc National de Garamba]; 03°40’ N, 29°00’ E; 15 May 1950; H. De Saeger leg.; RMCA • 1 ♂; same location as for preceding; 15 May 1950; H. De Saeger leg.; RMCA • 1 ♂; Uele: Dungu; 03°37’ N, 28°34’ E; 27 Jul. 1937; J.V. Leroy leg.;

RMCA. – **Ituri** • 1 ♀; Kasenyi; 01°24’ N, 30°26’ E; 22 Aug. 1935; H.J. Brédo leg.; RMCA • 1 ♀; same location as for preceding; Dec. 1938; P. Lefèvre leg.; RMCA • 2 ♀♀; Kibali-Ituri: Kilomines; 01°50’ N, 30°09’ E; May 1957; C. Smoor leg.; RMCA • 1 ♀; same location as for preceding; Oct. 1956; C. Smoor leg.; RMCA • 1 ♀; Lac Albert: Kasenyi; 01°24’ N, 30°26’ E; 13 May 1935; H.J. Brédo leg.; RMCA • 1 ♀; Mahagi-Niarembe; 02°15’ N, 31°07’ E; 1935; Ch. Scops leg.; RMCA • 1 ♀; Mongbwalu; 01°50’ N, 30°09’ E; May 1939; Mme. Scheitz leg.; RMCA. – **Kasaï** • 1 ♀; Tshikapa; 06°24’ S, 20°48’ E; 1930; Dr Fourche leg.; RMCA • 1 ♀; same location as for preceding; Apr. 1939; Mevr. Bequaert leg.; RMCA • 1 ♀; same location as for preceding; Apr. – May 1939; Mevr. Bequaert leg.; RMCA. – **Kasaï Central** • 1 ♀; Lula, Terr. Luiza; 07°11’ S, 22°24’ E; Aug. 1958; Dr. M. Poll leg.; RMCA • 1 ♂; same location as for preceding; Aug. 1958; Dr. M. Poll leg.; RMCA • 1 ♂; Luluabourg (Kananga); 05°54’ S, 21°52’ E; P. Callewaert leg.; RMCA • 1 ♀; Luluabourg: Katoka; 05°54’ S, 21°52’ E; 1939; R.P. Vankerckhoven leg.; RMCA. – **Kinshasa** • 1 ♂; Léopoldville (Kinshasa); 04°19’ S, 15°19’ E; 1933; A. Tinant leg.; RMCA • 1 ♂; same location as for preceding; Dr. Houssiaux leg.; RMCA • 4 ♂♂; same location as for preceding; Dr. Houssiaux leg.; RMCA. – **Kongo Central** • 1 ♂; Banana; 06°00’ S, 12°24’ E; 6 Aug. 1920; Dr. H. Schouteden leg.; RMCA • 1 ♀; Boma; 05°51’ S, 13°03’ E; 27 Mar. 1913; Lt. Styczynski leg.; RMCA • 2 ♀♀; same location as for preceding; R.F. Achille leg.; RMCA • 1 ♀; same location as for preceding; 9 Jun. 1915; Dr. M. Bequaert leg.; RMCA • 1 ♀; Cattier (Lufutoto); 05°26’ S, 14°45’ E; 1946-1949; Delafaille leg.; RMCA • 2 ♂♂; Kisantu (Madimba); 05°08’ S, 15°06’ E; 1927; R.P. Vanderyst leg.; RMCA • 1 ♀; same location as for preceding; 1931; R.P. Vanderyst leg.; RMCA • 1 ♂; same location as for preceding; 1931; R.P. Vanderyst leg.; RMCA • 1 ♀; same location as for preceding; Dec. 1927; R.P. Vanderyst leg.; RMCA • 1 ♂; same location as for preceding; 1945; Rév. Fr. Anastase leg.; RMCA • 1 ♀; same location as for preceding; May 1945; Rév. Fr. Anastase leg.; RMCA • 1 ♀; Lemfu (Madimba); 05°18’ S, 15°13’ E; Jan. 1945; Rév. P. De Beir leg.; RMCA • 1 ♂; same location as for preceding; Jun. 1945; Rév. P. De Beir leg.; RMCA • 3 ♀♀; Mayidi (Madimba); 05°11’ S, 15°09’ E; 1942; Rév. P. Van Eyen leg.; RMCA • 2 ♂♂; Thysville (Mbanza Ngungu); 05°15’ S, 14°52’ E; Jan. 1953; J. Sion leg.; RMCA • 1 ♀; same location as for preceding; 3 Mar. 1915; Dr. J. Bequaert leg.; RMCA • 1 ♀; same location as for preceding; 1930; Dr. Vanderhaegen leg.;

RMCA. – **Kwilu** • 1 ♀; Leverville (Bulungu); 04°50’ S, 18°44’ E; 1928; Mme. J. Tinant leg.; RMCA. – **Lomami** • 1 ♀; Kabinda; 06°08’ S, 24°29’ E; Oct. 1934; Mme. Gillardin leg.; RMCA • 1 ♀; Kanda-Kanda: Tshibata; 06°56’ S, 23°37’ E; J. Castelein leg.; RMCA • 2 ♀♀; Lomami: Kambaye; 06°53’ S, 23°44’ E; Oct. 1930; R. Massart leg.; RMCA • 1 ♀; Lomami-Luputa; 11°00’ S, 26°44’ E; Apr. 1934; Dr. Bouvier leg.; RMCA • 1 ♀; Sankuru: Gandajika; 06°45’ S, 23°57’ E; 19 Aug. 1950; P. de Francquen leg.; RMCA • 1 ♀; Ter. Kabinda; 06°08’ S, 24°29’ E; Dec. 1934; Mme. Gillardin leg.; RMCA. – **Lualaba** • 3 ♂♂; Biano; 10°20’ S, 27°00’ E; Oct. 1931; Ch. Seydel leg.; RMCA • 1 ♂; Dilolo; 10°41’ S, 22°21’ E; Sep. – Oct. 1933; H. De Saeger leg.; RMCA • 1 ♀; Haut-Luapula: Kansenia; 10°19’ S, 26°04’ E; 19 Oct. 1929; Dom de Montpellier leg.; RMCA • 1 ♂; Kanzenze; 10°31’ S, 25°12’ E; Aug. 1931; G.F. de Witte leg.; RMCA • 1 ♀; Kapanga; 08°21’ S, 22°34’ E; 4 Dec. 1932; F.G. Overlaet leg.; RMCA • 1 ♂; same location as for preceding; Aug. 1934; F.G. Overlaet leg.; RMCA • 1 ♀; same location as for preceding; Dec. 1932; F.G. Overlaet leg.; RMCA • 1 ♂; Kapiri; 10°18’ S, 26°11’ E; Sep. 1912; Miss. Agric. leg.; RMCA • 1 ♀; Kolwezi (à la lumière); 10°44’ S, 25°28’ E; 18 Oct. 1953; Mme. L. Gilbert leg.; RMCA • 2 ♀♀; Lubudi; 09°55’ S, 25°58’ E; Oct. 1931; Ch. Seydel leg.; RMCA • 1 ♀; Lulua: Kapanga; 08°21’ S, 22°34’ E; 11 Dec. 1932; F.G. Overlaet leg.; RMCA • 2 ♀♀; same location as for preceding; 3 Dec. 1932; F.G. Overlaet leg.; RMCA • 1 ♀; same location as for preceding; 8 Dec. 1932; F.G. Overlaet leg.; RMCA • 2 ♂♂; same location as for preceding; Sep. 1932; F.G. Overlaet leg.; RMCA • 3 ♀♀; same location as for preceding; Dec. 1932; F.G. Overlaet leg.; RMCA • 2 ♂♂; same location as for preceding; Dec. 1932; F.G. Overlaet leg.; RMCA. – **Maï-Ndombe** • 1 ♀; Bokala; 03°07’ S, 17°04’ E; 26 Apr. 1912; Dr. Mouchet leg.; RMCA • 1 ♀; Inongo; 01°57’ S, 18°16’ E; Aug. 1913; Dr. J. Maes leg.; RMCA • 1 ♀; Kwamouth (Mushie); 03°11’ S, 16°12’ E; Aug. 1911; Dr. Mouchet leg.; RMCA • 1 ♀; Yumbi; 01°54’ S, 16°33’ E; 29 Jul. 1912; Dr. Mouchet leg.; RMCA. – **Maniema** • 1 ♀; Kasongo; 04°27’ S, 26°40’ E; Mar. 1960; P.L.G. Benoit leg.; RMCA • 1 ♀; Kibombo; 03°54’ S, 25°55’ E; 18 Nov.1911; Dr. J. Bequaert leg.; RMCA • 1 ♀; same location as for preceding; Sep. – Oct. 1930; H.J. Brédo leg.; RMCA • 2 ♂♂; same location as for preceding; Sep. – Oct. 1930; H.J. Brédo leg.; RMCA • 1 ♀; same location as for preceding; Oct. 1930; H.J. Brédo leg.; RMCA • 1 ♂; Nyangwe; 04°13’ S, 26°11’ E; 1918; R. Mayné leg.; RMCA • 1 ♀; same location as for preceding; Apr. – May 1918; R. Mayné leg.; RMCA • 1 ♀; Terr. De Kasongo, Riv. Lumami; 04°27’ S, 26°40’ E; Sep. 1959; P.L.G. Benoit leg.; RMCA • 1 ♀; Vieux Kasongo; 04°27’ S, 26°40’ E; 17 Dec. 1910; Dr. J. Bequaert leg.; RMCA. – **Nord-Kivu** • 1 ♀; Beni à Lesse; 00°03’ N, 29°41’ E; Jul. 1911; Dr. Murtula leg.; RMCA • 1 ♀; Buseregenye (Rutshuru); 01°11’ S, 29°27’ E; Sep. 1929; Ed. Luja leg.; RMCA. – **Nord-Ubangi** • 1 ♀; Ubangi: Mokolo; 01°35’ N, 29°05’ E; 5 Dec. 1931; H.J. Brédo leg.; RMCA • 1 ♀; Yakoma; 04°06’ N, 22°23’ E; 5 – 17 Feb. 1932; H.J. Brédo leg.; RMCA• 2 ♂♂; same location as for preceding; 5 – 17 Feb. 1932; H.J. Brédo leg.; RMCA. – **Sud-Ubangi** • 2 ♀♀; Kunungu (Nkele); 02°45’ N, 18°26’ E; 1938; Dr. H. Schouteden leg.; RMCA • 1 ♀; M’Paka, Terr. Libenge; 03°39’ N, 18°38’ E; Jul. – Aug. 1959; M. Pecheur leg.; RMCA • 1 ♀; Ubangi: Boma-Motenge; 03°15’ N, 18°39’ E; Dec. 1931; H.J. Brédo leg.; RMCA • 1 ♂; same location as for preceding; Dec. 1931; H.J. Brédo leg.;

RMCA. – **Tanganyika** • 1 ♀; Buli (Kabalo); 05°50’ S, 26°55’ E; 17 Feb. 1911; Dr. J. Bequaert leg.; RMCA • 1 ♀; same location as for preceding; 18 Feb. 1911; Dr. J. Bequaert leg.; RMCA • 1 ♂; Kongolo; 05°24’ S, 27°00’ E; 6 Feb. 1911; R.P. Vankerckhoven leg.; RMCA. – **Tshopo** • 1 ♀; Stanleyville (Kisangani); 00°31’ N, 25°11’ E; 26 Feb. 1928; A. Collart leg.; RMCA • 1 ♀; same location as for preceding; 6-9 Sep. 1928; A. Collart leg.; RMCA • 1 ♀; same location as for preceding; 7 May 1926; Dr. H. Schouteden leg.; RMCA • 2 ♀♀; same location as for preceding; Jun. 1932; J. Vrydagh leg.; RMCA • 1 ♂; same location as for preceding; Jun. 1932; J. Vrydagh leg.; RMCA.

#### *Megachile (Cesacongoa) quadraticauda* (Pasteels, 1965), *Chalicodoma* (*Cuspidella*) *quadraticauda* Pasteels, 1965

**Material.** Cupboard 110, Box 45 (2♀♀). **D.R. CONGO.** Holotype – **Haut-Katanga** • 1 ♀; PNU Munoi bifure Lupiala; 08°50’ S, 26°44’ E, alt. 890 m; 28 May – 15 Jun. 1948; Mis. G.F. de Witte leg.; RMCA. Allotype – **Haut-Katanga** • 1 ♀; PNU Ganza pr. r. Kamandula af.dr. Lukoka; 09°14’ S, 26°42’ E, alt. 860 m; 12 – 18 Jun. 1949; Mis. G.F. de Witte leg.; RMCA.

#### *Megachile (Maximegachile) maxillosa* Guérin-Méneville, 1845, *Chalicodoma* (*Maximegachile*) *maxillosa* (Guérin-Méneville, 1845)

**Material.** Cupboard 110, Box 33 (13♀♀ and 1♂). **D.R. CONGO.** – **Haut-Katanga** • 1 ♂; Kambove-Lukafu; 10°52’ S, 26°38’ E; Apr. 1907; Dr. Sheffield Neave leg.; RMCA • 1 ♀; PNU Kabwe sur Muye; 08°47’ S, 26°52’ E, alt. 1320 m; 11 May 1948; Mis. G.F. de Witte leg.; RMCA • 1 ♀; PNU Masombwe s. Grde. Kafwe; 09°05’ S, 27°12’ E, alt. 1120 m; 16 Apr. 1948; Mis. G.F. de Witte leg.; RMCA. – **Haut-Lomami**• 1 ♀; Sankisia; 09°24’ S, 25°48’ E; 3 Sep. 1911; Dr. J. Bequaert leg.; RMCA. – **Ituri** • 1 ♀; Lac Albert: Ishwa; 02°13’ N, 31°12’ E; Sep. 1935; H.J. Brédo leg.; RMCA • 1 ♀; Mahagi-Niarembe; 02°15’ N, 31°07’ E; Sep. 1935; Ch. Scops leg.; RMCA. – **Lualaba** • 1 ♀; Dilolo; 10°41’ S, 22°21’ E; Aug. 1931; G.F. de Witte leg.; RMCA • 1 ♀; Lulua: Kanzenze; 10°31’ S, 25°12’ E; 1932; F.G. Overlaet leg.; RMCA. – **Tanganyika** • 1 ♀; Tanganyika: Moba; 07°02’ S, 29°47’ E, alt. 780 m; Jul. – Aug. 1953; H. Bomans leg.; RMCA • 5 ♀♀; Tanganyika: Mpala; 06°45’ S, 29°31’ E, alt. 780 m; Jun. 1953; H. Bomans leg.; RMCA.

#### *Megachile (Morphella) ambigua* (Pasteels, 1965), *Chalicodoma* (*Morphella*) *ambigua* Pasteels, 1965

**Material.** Cupboard 110, Box 45 (1♂). **D.R. CONGO.** Holotype – **Equateur** • 1 ♂; Eala (Mbandaka); 00°03’ N, 18°19’ E; Sep. 1935; J. Ghesquière leg.; RMCA.

#### *Megachile (Morphella) biseta* Vachal, 1903, *Chalicodoma* (*Morphella*) *biseta* (Vachal, 1903)

**Material.** Cupboard 110, Box 45 (35♀♀ and 1♂). **D.R. CONGO.** – **Bas-Uele** • 1 ♀; Bambesa; 03°28’ N, 25°43’ E; 4 Jun. 1937; J. Vrydagh leg.; RMCA. – **Equateur** • 2 ♀♀; Eala (Mbandaka); 00°03’ N, 18°19’ E; 15-30 Nov. 1929; H.J. Brédo leg.; RMCA • 1 ♀; same location as for preceding; Oct. 1935; J. Ghesquière leg.; RMCA • 1 ♀; same location as for preceding; Nov. 1935; J. Ghesquière leg.; RMCA • 2 ♀♀; Eala: Boyeka; 00°03’ N, 18°19’ E; 30 Nov. 1929; H.J. Brédo leg.; RMCA. – **Haut-Katanga** • 1 ♀; PNU Kabwe sur Muye; 08°47’ S, 26°52’ E; 11 May 1948; Mis. G.F. de Witte leg.; RMCA. – **Haut-Lomami** • 2 ♀♀; Katanga: Luashi; 07°31’ S, 24°10’ E; 1935; Freyne leg.; RMCA • 1 ♀; Lualaba: Kabongo; 07°20’ S, 25°35’ E; 30 Dec. 1952; Ch. Seydel leg.; RMCA. – **Ituri** • 1 ♀; Kibali-Ituri: Kilomines; 01°50’ N, 30°09’ E; 20 Apr. 1958; I. Mesmaekers leg.; RMCA • 1 ♀; same location as for preceding; Sep. 1957; I. Mesmaekers leg.; RMCA. – **Lualaba** • 19 ♀♀; Lulua: Kapanga; 08°21’ S, 22°34’ E; Oct. 1932; F.G. Overlaet leg.; RMCA. – **Kongo Central** • 1 ♀; Bas-Congo: Kisantu; 05°08’ S, 15°06’ E; Rév. P. Regnier leg.; RMCA. – **Mongala**• 1 ♀; Binga (Lisala); 02°28’ N, 20°31’ E; 1 Mar. 1932; H.J. Brédo leg.; RMCA. – **Nord-Kivu** • 2 ♀♀; Kivu: Tshibinda; 02°19’ S, 28°45’ E; 21-27 Aug.1931; T.D. Cockerell leg.; RMCA. – **Sud-Kivu** • 1♂; Kivu: Mulungu; 02°19’ S, 28°45’ E; 23 Feb.1938; F.L. Hendrickx leg.; RMCA. – **Tshuapa** • 1 ♀; Tshuapa: Bokungu; 00°41’ S, 22°19’ E; 1949; Dupuis leg.; RMCA.

#### *Megachile (Neglectella) mimetica* Cockerell, 1933, *Chalicodoma* (*Neglectella*) *mimetica* (Cockerell, 1933)

**Material.** Cupboard 110, Box 21 (10♂♂). **D.R. CONGO.** – **Haut-Katanga** • 2 ♂♂; Elisabethville (Lubumbashi); 11°40’ S, 27°28’ E; Jun. 1932; De Loose leg.; RMCA • 1 ♂; Kapema; 10°42’ S, 28°40’ E; Sep. 1924; Ch. Seydel leg.; RMCA • 1 ♂; PNU Kilwezi affl.dr. Lufira; 09°05’ S, 26°45’ E, alt. 750 m; 16-21 Aug. 1948; Mis. G.F. de Witte leg.; RMCA. – **Haut-Lomami** • 1 ♂; Luashi; 07°31’ S, 24°10’ E; 1935; Freyne leg.; RMCA. – **Lualaba** • 3 ♂♂; Dilolo; 10°41’ S, 22°21’ E; Aug. 1931; G.F. de Witte leg.; RMCA • 2 ♂♂; Kanzenze; 10°31’ S, 25°12’ E; Aug. 1931; G.F. de Witte leg.; RMCA.

#### *Megachile (Neglectella) scindularia* Buysson, 1903, *Chalicodoma* (*Neglectella*) *scindularia* (Buysson, 1903)

**Material.** Cupboard 110, Box 21(5♀♀ and 8♂♂). **D.R. CONGO.** – **Equateur** • 1 ♀; Eala (Mbandaka); 00°03’ N, 18°19’ E; 7 Jul. 1932; A. Corbisier leg.; RMCA • 2 ♀♀; same location as for preceding; 15 – 30 Nov. 1929; H.J. Brédo leg.; RMCA • 1 ♂; same location as for preceding; 28 Nov. 1929; H.J. Brédo leg.; RMCA • 1 ♂; Lukolela (Bikoro); 01°03’ S, 17°12’ E; Nov. 1934; Dr. Ledoux leg.; RMCA. – **Kasaï Central** • 1 ♂; Luluabourg; 05°54’ S, 21°52’ E; 14 – 17 May 1919; P. Callewaert leg.; RMCA. – **Kongo Central** • 2 ♂♂; Mayidi (Madimba); 05°11’ S, 15°09’ E; 1942; Rév. P. Van Eyen leg.; RMCA. – **Tshopo** • 1 ♀; Ponthierville (Ubundu); 00°22’ S, 25°29’ E; 22 Oct. 1910; Dr. J. Bequaert leg.; RMCA • 2 ♂♂; same location as for preceding; 22 Oct. 1910; Dr. J. Bequaert leg.; RMCA • 1 ♀; same location as for preceding; 24 Oct. 1910; Dr. J. Bequaert leg.; RMCA • 1 ♂; Stanleyville (Kisangani); 00°31’ N, 25°11’ E; 16 Apr. 1915; Exp Lang-Chapin leg.; RMCA.

#### *Megachile (Neglectella) stefenelii* Friese, 1903, *Chalicodoma* (*Neglectella*) *stefenelii* (Friese, 1903)

**Material.** Cupboard 110, Box 21 (1♂). **D.R. CONGO.** – **Ituri** • 1 ♂; Lac Albert: Ishwa; 02°13’ N, 31°12’ E; Sep. 1935; H.J. Brédo leg.; RMCA

#### *Megachile (Pseudomegachile) gibbidens* Vachal, 1910, *Chalicocoma* (*Pseudomegachile*) *gibbidens* (Vachal, 1910)

**Material.** Cupboard 110, Box 20 (6♀♀). **D.R. CONGO.** Holotypes DRC – **Haut-Katanga** • 1 ♀; Kundelungu; 10°40’ S, 28°00’ E; Sep. 1907; Dr. Sheffield-Neave leg.; RMCA. Paratypes – **Haut-Katanga** • 5 ♀♀; Kundelungu; 10°40’ S, 28°00’ E; Sep. 1907; Dr. Sheffield-Neave leg.; RMCA.

#### *Megachile (Pseudomegachile) battorensis* Meade-Waldo, 1912, *Chalicodoma* (*Pseudomegachile*) *battorensis* (Meade-Waldo, 1912)

**Material.** Cupboard 110, Box 20 (13♀♀). **D.R. CONGO.** – **Bas-Uele** • 1 ♀; Bambili (riv. Uele); 03°38’ N, 26°06’ E; Dr. M. Bequaert leg.; RMCA • 1 ♀; Bambili; 03°38’ N, 26°06’ E; 24 Nov. 1913; Dr. Rodhain leg.; RMCA. 1 ♀; Bambili; 03°38’ N, 26°06’ E; 29 Nov. 1913; Dr. Rodhain leg.; – **Haut-Uele** • 1 ♀; PNG [Parc National de Garamba]; 03°40’ N, 29°00’ E; 3 Aug. 1951; H. De Saeger leg.; RMCA. – **Haut-Uele** • 1 ♀; Kibali-Ituri: Abock; 02°00’ N, 31°00’ E; 2 Oct. 1935; Ch. Scops leg.; RMCA • 1 ♀; Kibimbi; 01°22’ N, 28°55’ E; 2 Feb. 1911; Dr. J. Bequaert leg.; RMCA • 1 ♀; Lac Albert: Kasenyi; 01°24’ N, 30°26’ E; 1935; H.J. Brédo leg.; RMCA • 1 ♀; same location as for preceding; 15 May 1935; H.J. Brédo leg.; RMCA • 1 ♀; same location as for preceding; 4 Sep. 1935; H.J. Brédo leg.; RMCA. • 1 ♀; Région d’Abok; 02°00’ N, 31°00’ E; Oct. 1935, M. & Mme. Ch. Scops leg.; RMCA. – **Maniema** • 2 ♀♀; Nyangwe; 04°13’ S, 26°11’ E; 26 Nov. 1910; Dr. J. Bequaert leg.; RMCA. – **Lomami** • 1 ♀; Sankuru: Gandajika; 06°45’ S, 23°57’ E; 27 Jan. 1951; P. de Francquen leg.; RMCA.

#### *Megachile (Pseudomegachile) biloba* Vachal, 1910, *Chalicodoma* (*Pseudomegachile*) *biloba* (Vachal, 1910)

**Material.** Cupboard 110, Box 20 (13♀♀). **D.R. CONGO.** Holotype – **Haut-Katanga** • 1 ♂; Kundelungu; 10°40’ S, 28°00’ E; Sep. 1907; Dr. Sheffield-Neave leg.; RMCA.

– **Haut-Katanga** • 1 ♂; PNU Riv. Lusinga; 08°55’ S, 27°12’ E; 20 Jul. 1945; G.F. de Witte leg.; RMCA.

#### *Megachile (Pseudomegachile) bukamensis* Cockerell, 1935, *Chalicodoma* (*Pseudomegachile*) *bukamensis* (Cockerell, 1935)

**Material.** Cupboard 110, Box 20 (13♀♀). **D.R. CONGO.** Holotype – **Haut-Lomami** • 1 ♂; Bukama; 09°12’ S, 25°51’ E; 14 Jul. 1911; Dr. J. Bequaert leg.; RMCA.

– **Haut-Katanga** • 1 ♂; Elisabethville (Lubumbashi); 11°40’ S, 27°28’ E; 11-17 Sep. 1931; T.D. Cockerell leg.; RMCA • 4 ♀♀; PNU Lusinga (riv. Kamitungulu); 08°55’ S, 08°55’ S, 13 Jun. 1945; G.F. de Witte leg.; RMCA • 1 ♂; PNU Riv. Lusinga; 08°55’ S, 27°12’ E; 20 Jul. 1945; G.F. de Witte leg.; RMCA.

#### *Megachile (Pseudomegachile) congruens* Friese, 1903, *Chalicodoma* (*Pseudomegachile*) *congruens* (Friese, 1903)

**Material.** Cupboard 110, Box 19 (30♀♀ and 15♂♂). **D.R. CONGO.** – **Bas-Uele** • 1 ♂; Bambesa; 03°28’ N, 25°43’ E; 6 Nov. 1937, J. Vrijdagh leg.; RMCA. – **Haut-Katanga** • 1 ♂; Elisabethville (Lubumbashi); 11°40’ S, 27°28’ E; Sep.1931; Ch. Seydel leg.; RMCA • 1 ♂; Elisabethville (60km ruiss. Kasepa); 11°40’ S, 27°28’ E; 23 Sep. 1923; Dr. M. Bequaert leg.; RMCA. – **Haut-Lomami** • 4 ♀♀; Kinda: Kitundu; 09°29’ S, 24°49’ E; 16 Sep. 1914; L. Charliers leg.; RMCA • 13 ♀♀; same location as for preceding; 19 Sep. 1914; L. Charliers leg.; RMCA. – **Haut-Uele** • 1 ♀; PNG [Parc National de Garamba]; 03°40’ N, 29°00’ E; 6 Feb. 1951; H. De Saeger leg.; RMCA • 1 ♂; Uele: Dungu; 03°37’ N, 28°34’ E; 6 Feb. 1951, De Greef leg.; RMCA. – **Ituri** • 2 ♂♂; Kibali-Ituri: Mahagi; 02°18’ N, 30°59’ E; 1932; Ch. Scops leg.; RMCA • 1 ♀; Lac Albert: Ishwa; 02°13’ N, 31°12’ E; Sep. 1935; H.J. Brédo leg.; RMCA • 2 ♂♂; same location as for preceding; Sep. 1935; H.J. Brédo leg.; RMCA • 1 ♂; Lac Albert: Kasenyi; 01°24’ N, 30°26’ E; 1935; H.J. Brédo leg.; RMCA • 2 ♂♂; same location as for preceding; 1 May 1935; H.J. Brédo leg.; RMCA • 1 ♂; Mahagi-Port; 02°09’ N, 31°14’ E; 1935; Ch. Scops leg.; RMCA • 1 ♂; same location as for preceding; Sep. 1934; H.J. Brédo leg.; RMCA • 1 ♀; same location as for preceding; Oct. 1934; H.J. Brédo leg.; RMCA. – **Kasaï Central** • 1 ♂; Kasaï: Lula, Terr. Luiza; 07°11’ S, 22°24’ E; Aug. 1956; Dr. M. Poll leg.; RMCA. – **Lomami** • 1 ♂; Gandajika; 06°45’ S, 23°57’ E; 1952; P. de Francquen leg.; RMCA • 2 ♀♀; Bunkeya; 10°24’ S, 26°58’ E; Oct. 1907; Dr. Sheffield-Neave leg.; RMCA • 1 ♀; Sankuru: Gandajika; 06°45’ S, 23°57’ E; 27 Jan. 1951; P. de Francquen leg.; RMCA. – **Lualaba** • 1 ♀; Lubudi; 09°55’ S, 25°58’ E; 27 Aug. 1923; Ch. Seydel leg.; RMCA • 2 ♀♀; Sandoa; 09°41’ S, 22°53’ E; 1929; Ch. Seydel leg.; RMCA. – **Maniema** • 1 ♂; Nyangwe; 04°13’ S, 26°11’ E; Apr.-May 1918; R. Mayné leg.; RMCA. – **Nord-Kivu** • 1♀; Buseregenye (Rutshuru); 01°11’ S, 29°27’ E; 1930; Ed. Luja leg.; RMCA. – **Tanganyika** • 1 ♂; Bassin Lukuga; 05°40’ S, 26°55’ E; Apr.-Jun. 1934; H. De Saeger leg.; RMCA • 1 ♂; Kongolo; 05°24’ S, 27°00’ E; 23 Jan. 1911; Dr. M. Bequaert leg.; RMCA • 1 ♀; Lukulu (Manono); 07°08’ S, 28°06’ E; 4 – 7 Jul. 1931; G.F. de Witte leg.; RMCA.

#### *Megachile (Pseudomegachile) fastigiata* Vachal, 1910, *Chalicodoma* (*Pseudomegachile*) *fastigiata* (Vachal, 1910)

**Material.** Cupboard 110, Box 20 (2♀♀). **D.R. CONGO.** Holotype – **Haut-Katanga** • 1 ♀; Kambove-Bunkeya; 10°52’ S, 26°38’ E; Oct. 1907; Dr. Sheffield-Neave leg.; RMCA. Paratype – **Haut-Katanga** • 1 ♀; Kambove-Bunkeya; 10°52’ S, 26°38’ E; Oct. 1907; Dr. Sheffield-Neave leg.; RMCA.

#### *Megachile (Pseudomegachile) kigonserana* Friese, 1903, *Chalicodoma* (*Pseudomegachile*) *kigonserana* (Friese, 1903)

**Material.** Cupboard 110, Box 20 (22♀♀ and 5♂♂). **D.R. CONGO.** – **Haut-Katanga** • 1 ♀; Elisabethville (Lubumbashi); 11°40’ S, 27°28’ E; 5 Sep. 1923; Ch. Seydel leg.; RMCA • 1 ♂; same location as for preceding; 1–6 Sep. 1932; De Loose leg.; RMCA • 1 ♂; same location as for preceding; 5 Sep. 1923; Ch. Seydel leg.; RMCA • 1 ♀; same location as for preceding; 25 Sep. 1932; Dr. M. Bequaert leg.; RMCA • 1 ♀; same location as for preceding; Aug. 1923; Dr. M. Bequaert leg.; RMCA • 1 ♀; Kalengalele; 09°55’ S, 26°59’ E; 14 Jul. 1924; Ch. Seydel leg.; RMCA • 1 ♀; Kambove-Lukafu; 10°52’ S, 26°38’ E; Oct. 1907; Dr. Sheffield-Neave leg.; RMCA • 2 ♀♀; Lukonzolwa-Chaka; 08°47’ S, 28°38’ E; 29 Aug. 1907; Dr. Sheffield-Neave leg.; RMCA • 1 ♀; Mwema; 08°13’ S, 27°28’ E; Jul. 1927; A. Bayet leg.; RMCA. – **Haut-Lomami** • 2 ♀♀; Bukama; 09°12’ S, 25°51’ E; 16 Apr. 1911; Dr. J. Bequaert leg.; RMCA • 1 ♀; Kalongwe; 11°01’ S, 25°13’ E; 19 Sep. 1911; Dr. J. Bequaert leg.; RMCA • 1 ♀; Kulu-Mwanza; 07°54’ S, 26°45’ E; May 1927; A. Bayet leg.; RMCA Dr. M. Bequaert • 1 ♀; Mwabo; 08°44’ S, 25°00’ E; Jun. 1932; Ch. Seydel leg.; RMCA Dr. M. Bequaert • 1 ♀; Sankisia; 09°24’ S, 25°48’ E; 25 Sep. 1911; Dr. J. Bequaert leg.; RMCA. – **Lualaba** • 5 ♀♀; Bunkeya; 10°24’ S, 26°58’ E; Oct. 1907; Dr. Sheffield-Neave leg.; RMCA • 1 ♂; Dilolo; 10°41’ S, 22°21’ E; 24-27 Jul. 1931; Mrs. W.P. Cockerell leg.; RMCA • 3 ♀♀; Katentenia; 10°19’ S, 25°54’ E; May 1924; Ch. Seydel leg.; RMCA • 1 ♂; same location as for preceding; May 1924; Ch. Seydel leg.; RMCA • 1 ♂; Lulua: Kapanga; 08°21’ S, 22°34’ E; Dec. 1932; G.F. Overlaet leg.; RMCA.

#### *Megachile (Pseudomegachile) marchalli* Friese, 1904, *Chalicodoma* (*Pseudomegachile*) *marchalli* (Friese, 1904)

**Material.** Cupboard 110, Box 20 (1♀ and 2♂♂). **D.R. CONGO.** – **Haut-Lomami** • 1 ♀; Bukama; 09°12’ S, 25°51’ E; 10 Jun. 1911; Dr. J. Bequaert leg.; RMCA • 1 ♂; same location as for preceding; 12 Jul. 1911; Dr. J. Bequaert leg.; RMCA • 1 ♂; Sankisia; 09°24’ S, 25°48’ E; 30 Jul. 1911; Dr. J. Bequaert leg.; RMCA.

#### *Megachile (Pseudomegachile) mossambica* Gribodo, 1895, *Chalicodoma* (*Pseudomegachile*) *mossambica* (Gribodo, 1895)

**Material.** Cupboard 110, Box 19 (1♀). **D.R. CONGO.** – **Haut-Katanga** • 1 ♀; PNU Kanonga; 09°16’ S, 26°08’ E, alt. 675 m; 14-23 Apr. 1949; Mis. G.F. de Witte leg.; RMCA.

#### *Megachile (Pseudomegachile) natalensis* Friese, 1921, *Chalicodoma* (*Pseudomegachile*) *congruens r. natalensis* (Friese, 1921)

**Material.** Cupboard 110, Box 19 (3♂♂). **D.R. CONGO.** – **Haut-Katanga** • 1 ♂; Mwera; 11°09’ S, 27°19’ E; 1956; R.P.Th. de Caters leg.; RMCA. – **Tanganyika**• 1 ♂; Katompe (Kabalo); 06°11’ S, 26°20’ E; Feb. 1929; Ch. Seydel leg.; RMCA. 1 ♂; Bomokandi (sources); 26 Nov. – 6 Dec. 1925; S. A. R. Prince Léopold leg.; RMCA.

#### *Megachile (Pseudomegachile) neavei* Vachal, 1910, *Chalicodoma* (*Pseudomegachile*) *neavei* (Vachal, 1910)

**Material.** Cupboard 110, Box 20 (21♀♀ and 2♂♂). **D.R. CONGO.** Holotype – **Lualaba** • 1 ♀; Bunkeya-Kambove; 10°24’ S, 26°58’ E; Oct. 1907; Dr. Sheffield-Neave leg.; RMCA. Paratypes – **Haut-Katanga** • 1 ♀; Lukafu; 10°31’ S, 27°33’ E; Oct. 1907; Dr. Sheffield-Neave leg.; RMCA • 2 ♀♀; Lukafu-Bunkeya; 10°31’ S, 27°33’ E; Oct. 1907; Dr. Sheffield-Neave leg.; RMCA. – **Lualaba** • 4♀♀; Bunkeya; 10°24’ S, 26°58’ E; Oct. 1907; Dr. Sheffield-Neave leg.; RMCA • 1 ♂; same location as for preceding; Oct. 1907; Dr. Sheffield-Neave leg.; RMCA • 1 ♂; Bunkeya-Kambove; 10°24’ S, 26°58’ E; Oct. 1907; Dr. Sheffield-Neave leg.; RMCA.

– **Haut-Katanga** • 2 ♀♀; Elisabethville; 11°40’ S, 27°28’ E; De Loose leg.; RMCA • 1 ♀; same location as for preceding; 1 Aug. 1933; Dr. M. Bequaert leg.; RMCA • 1 ♀; same location as for preceding; 15 Aug. 1933; Dr. M. Bequaert leg.; RMCA • 1 ♀; same location as for preceding; 22 Sep. 1923; Dr. M. Bequaert leg.; RMCA • 1 ♀; same location as for preceding; 11-17 Sep. 1931; T.D. Cockerell leg.; RMCA • 1 ♀; Panda (Likasi); 11°00’ S, 26°44’ E; 1 Sep. 1920; Dr. M. Bequaert leg.; RMCA. – **Haut-Lomami** • 1 ♀; Bukama; 09°12’ S, 25°51’ E; Aug. 1923; Ch. Seydel leg.; RMCA • 1 ♂; Sankisia; 09°24’ S, 25°48’ E; Dr. Rodhain leg.; RMCA. – **Lualaba** • 3 ♀♀; Lubudi; 09°55’ S, 25°58’ E; 27 Jul. 1923; Ch. Seydel leg.; RMCA.

– **Tanganyika** • 1 ♀; Lubombo; 08°10’ S, 29°45’ E; Sep. 1928; Ch. Seydel leg.; RMCA.

#### *Megachile (Pseudomegachile) schulthessi* Friese, 1903, *Chalicodoma* (*Pseudomegachile*) *schultessi* (Friese, 1903)

**Material.** Cupboard 110, Box 20 (16♀♀). **D.R. CONGO.** – **Haut-Katanga** • 16 ♀♀; PNU Lusinga (riv. Kamitungulu); 08°55’ S, 08°55’ S; 13 Jun. 1945; G.F. de Witte leg.; RMCA.

#### *Megachile (Pseudmegachile) sinuata* Friese, 1903, *Chalicodoma* (*Pseudmegachile*) *sinuata* (Friese, 1903)

**Material.** Cupboard 110, Box 19 (2♀♀ and 1♂). **D.R. CONGO.** – **Bas-Uele** • 1 ♂; Bambesa; 03°28’ N, 25°43’ E; Feb. 1934; H.J. Brédo leg.; RMCA. – **Haut-Uele** • 2 ♀♀; PNG [Parc National de Garamba]; 03°40’ N, 29°00’ E; 6 Feb. 1951; H. De Saeger leg.; RMCA.

#### *Megachile (Stenomegachile) chelostomoides* Gribodo, 1894, *Chalicodoma* (*Stenomegachile*) *chelostomoides* (Gribodo, 1894)

**Material.** Cupboard 110, Box 33 (6♀♀). **D.R. CONGO.** – **Haut-Katanga** • 1 ♀; PNU Kiamakoto entre Masombwe-Mukana r. dr. Gr. Kafwe; 09°09’ S, 27°10’ E, alt. 1070 m; 20 Sep. 1948; Mis. G.F. de Witte leg.; RMCA • 1 ♀; PNU Kilwezi affl. dr. Lufira; 09°05’ S, 26°45’ E, alt. 750 m; 9-14 Aug. 1948; Mis. G.F. de Witte leg.; RMCA. – **Haut-Lomami** • 1 ♀; Sankisia; 09°24’ S, 25°48’ E; 29 Aug. 1911; Dr. J. Bequaert leg.; RMCA • 2 ♀♀; same location as for preceding; 4 Sep. 1911; Dr. J. Bequaert leg.; RMCA • 1 ♀; same location as for preceding; 13 Sep. 1911; Dr. J. Bequaert leg.; RMCA.

#### *Coelioxys (Allocoelioxys) afra* Lepeltier, 1841

**Material.** Cupboard 110, Box 54 (4♀♀ and 4♂♂). **D.R. CONGO.** – **Equateur** • 1 ♂; Eala (Mbandaka); 00°03’ N, 18°19’ E; Apr. 1935; J. Ghesquière leg.; RMCA • 1 ♀; same location as for preceding; Apr. 1932; A. Corbisier leg.; RMCA. – **Haut-Lomami** • 1 ♂; PNU Mabwe; 08°47’ S, 26°52’ E, alt. 585 m, 22 Jan. 1948; Mis. G.F. de Witte leg.; RMCA. – **Haut-Uele** • 2 ♀♀; PNG Akam; 03°40’ N, 29°00’ E; 21 Apr. 1950; H. De Saeger leg.; RMCA. – **Ituri**, • 1 ♀; Mahagi-Port; 02°09’ N, 31°14’ E; 1929; Ch. Seydel leg.; RMCA. – **Lualaba** • 1 ♂; Ditanto; 10°15’ S, 25°53’ E; Oct. 1925; Ch. Seydel leg.; RMCA. – **Maï-Ndombe** • 1 ♂; Wombali (Mushie); 03°16’ S, 17°22’ E; Jul. 1913; P. Vanderijst leg.; RMCA.

#### *Coelioxys (Allocoelioxys) congoensis* Friese, 1922

**Material.** Cupboard 110, Box 52 (2♀♀ and 2♂♂). **D.R. CONGO.** – **Haut-Uele** • 1 ♀; PNG [Parc National de Garamba]; 03°40’ N, 29°00’ E; 31 Mar. 1951; H. De Saeger leg.; RMCA. – **Kongo Central** • 1 ♂; Mayidi (Madimba); 05°11’ S, 15°09’ E; 1942; Rév. P. Van Eyen leg.; RMCA. – **Lomami** • 1 ♀; Tshofa; 05°14’ S, 25°15’ E; Dec. 1934; Mme. Gillardin n leg.; RMCA. – **Maniema** • 1 ♂; Kibombo; 03°54’ S, 25°55’ E; Sep.-Oct. 1930; Mme. Gillardin leg.; RMCA.

#### *Coelioxys (Allocoelioxys) difformis* Friese, 1904

**Material.** Cupboard 110, Box 52 (1♀). **D.R. CONGO.** – **Sud-Kivu** • 1 ♀; Uvira; 03°25’ S, 29°08’ E; J. Pasteels leg.; RMCA.

#### *Coelioxys (Coelioxys) aurifrons* Smith, 1854

**Material.** Cupboard 110, Box 53 (18♀♀ and 4♂♂). **D.R. CONGO.** – **Bas-Uele** • 2 ♀♀; Bambesa; 03°28’ N, 25°43’ E; 15 Sep. 1933; H.J. Brédo leg.; RMCA • 1 ♀; same location as for preceding; Sep. 1933; H.J. Brédo leg.; RMCA. – **Equateur** • 1 ♀; Eala (Mbandaka); 00°03’ N, 18°19’ E; 22 Nov. 1931; H.J. Brédo leg.; RMCA • 1 ♀; same location as for preceding; 8 Dec. 1932; H.J. Brédo leg.; RMCA • 3 ♀♀; same location as for preceding; Mar. 1932; H.J. Brédo leg.; RMCA • 1 ♂; same location as for preceding; Mar. 1932; H.J. Brédo leg.; RMCA • 1 ♀; same location as for preceding; Apr. 1932; H.J. Brédo leg.; RMCA • 1 ♀; same location as for preceding; Nov. 1934; J. Ghesquière leg.; RMCA • 1 ♀; same location as for preceding; Dec. 1932; H.J. Brédo leg.; RMCA. – **Haut-Katanga** • 1 ♂; Elisabethville (Lubumbashi); 11°40’ S, 27°28’ E; 17 Nov. 1928; Dr. M. Bequaert leg.; RMCA. – **Haut-Lomami** • 1 ♂; Lualaba: Kabongo; 07°20’ S, 25°35’ E; 7 Jan. 1953; Ch. Seydel leg.; RMCA. – **Kinshasa** • 1 ♂; Léopoldville (Kinshasa); 04°19’ S, 15°19’ E; Aug. 1949; Dr. A. Dubois leg.; RMCA. – **Kongo Central** • 1 ♀; Banza Manteka; 05°28’ S, 13°47’ E; 10-15 Jun. 1912; R. Mayné leg.; RMCA. – **Lualaba** • 1 ♀; Kapanga; 08°21’ S, 22°34’ E; Nov. 1933; F.G. Overlaet leg.; RMCA. – **Maniema** • 1 ♀; Kindu; 02°57’ S, 25°55’ E; Nov. 1913; L. Burgeon leg.; RMCA. – **Sud-Kivu** • 1 ♀; Ibanda; 02°29’ S, 28°51’ E; 1935; M. Vandelannoite leg.; RMCA • 2 ♀♀; same location as for preceding; 1952; M. Vandelannoite leg.; RMCA. – **Tshopo** • 1 ♀; Basoko; 01°13’ N, 23°36’ E; Sep. 1948; P.L.G. Benoit leg.; RMCA.

#### *Coelioxys (Coelioxys) capensis* Smith, 1954

**Material.** Cupboard 110, Box 53 (8♀♀). **D.R. CONGO.** – **Haut-Katanga** • 1 ♀; Elisabethville (Lubumbashi); 11°40’ S, 27°28’ E; 1 Feb. 1935; P. Quarré leg.; RMCA • 2 ♀♀; same location as for preceding; Feb. 1935; P. Quarré leg.; RMCA • 1 ♀; same location as for preceding; Oct. 1934; P. Quarré leg.; RMCA • 4 ♀♀; same location as for preceding; Dec. 1934; P. Quarré leg.; RMCA.

#### *Coelioxys (Coelioxys) coeruleipennis* Friese, 1904

**Material.** Cupboard 110, Box 52 (3♀♀ and 1♂). **D.R. CONGO.** – **Equateur** • 1 ♂; Eala (Mbandaka); 00°03’ N, 18°19’ E; Nov. 1931; H.J. Brédo leg.; RMCA. – **Haut-Katanga** • 1 ♀; Elisabethville (Lubumbashi); 11°40’ S, 27°28’ E; Apr. 1933; De Loose leg.; RMCA • 1 ♀; same location as for preceding; Jun. 1946; Ch. seydel leg.; RMCA. – **Haut-Lomami**• 1 ♀; Kulu-Mwanza; 07°54’ S, 26°45’ E; May 1927; A. Bayet leg.; RMCA.

#### *Coelioxys (Coelioxys) cyanura* Cockerell, 1932

**Material.** Cupboard 110, Box 54 (5♀♀ and 3♂♂). **D.R. CONGO.** Holotype – **Haut-Katanga** • 1 ♂; Elisabethville, riv. Kimilolo; 11°40’ S, 27°28’ E; 16 Jan. 1930; Dr. M. Bequaert leg.; RMCA.

– **Bas-Uele** • 1 ♀; Bambesa; 03°28’ N, 25°43’ E; 15 Sep. 1933; H.J. Brédo leg.; RMCA • 1 ♀; same location as for preceding; Dec. 1933; H.J. Brédo leg.; RMCA • 1 ♀; Uele: Bambesa; 03°28’ N, 25°43’ E; 10 Oct. 1933; J.V. Leroy leg.; RMCA. – **Haut-Katanga** • 1 ♂; Elisabethvile: Nord Tshinsangwe; 11°40’ S, 27°28’ E; 4 Feb. 1923; Dr. M. Bequaert leg.; RMCA. – **Kongo Central** • 1 ♀; Lemfu (Madimba); 05°18’ S, 15°13’ E; Oct.-Dec. 1944; Rév. P. De Beir leg.; RMCA. – **Lualaba** • 1 ♂; Lulua: Kapanga; 08°21’ S, 22°34’ E; Nov. 1932; F.G. Overlaet n leg.; RMCA. – **Sud-Kivu** • 1 ♀; Mulungu; 02°20’ S, 28°47’ E; 14 May 1938; Hendrickx leg.; RMCA.

#### *Coelioxys (Coelioxys) foveolata* Smith, 1854

**Material.** Cupboard 110, Box 52 (12♀♀ and 10♂♂). **D.R. CONGO.** – **Equateur** • 1 ♀; Eala (Mbandaka); 00°03’ N, 18°19’ E; Nov. 1934; J. Ghesquière leg.; RMCA. – **Haut-Katanga** • 1 ♀; Kambove; 10°52’ S, 26°38’ E; Feb. 1907; Dr. Sheffield Neave leg.; RMCA • 1 ♀; Kiamanwa; 09°30’ S, 27°57’ E; Nov. 1931; H.J. Brédo leg.; RMCA • 1 ♂; PNU Masombwe S, Grde. Kafwe; 09°05’ S, 27°12’ E, alt. 1120 m; Apr. 1948; Mis. G.F. de Witte leg.; RMCA. – **Haut-Lomami** • 1 ♂; Lomami: Mutombo-Mukulu; 07°58’ S, 24°00’ E; Jun. 1931; P. Quarré leg.; RMCA • 1 ♂; Lualaba: Kabongo; 07°20’ S, 25°35’ E; 30 Dec. 1952; Ch. Seydel leg.; RMCA. – **Haut-Uele** • 1 ♀; Dungu; 03°45’ N, 29°31’ E; Sep. 1919; P. Van den Plas leg.; RMCA. – **Ituri** • 1 ♂; Bunia; 01°34’ N, 30°15’ E; 1937; R.F. Maristes leg.; RMCA • 1 ♂; Ituri: Bunia; 01°34’ N, 30°15’ E; 23 Feb. 1934; J.V. Leroy leg.; RMCA • 1 ♂; same location as for preceding; 1937; 23 Feb. 1934; P. Lefèvre leg.; RMCA. – **Kasaï Central** • 1 ♀; Luluabourg (Kananga); 05°54’ S, 21°52’ E; 1937; P. Callewaert leg.; RMCA • 1 ♂; same location as for preceding; 1937; P. Callewaert leg.; RMCA. – **Kongo Central** • 1 ♀; Kisantu (Madimba); 05°08’ S, 15°06’ E; 1927; R.P. Vanderyst leg.; RMCA • 1 ♂; same location as for preceding; 1927; R. P. Vanderyst leg.; RMCA • 1 ♂; same location as for preceding; Dec. 1927; R. P. Vanderyst leg.; RMCA. – **Lualaba** • 1 ♀; Kafakumba (Sandoa); 09°41’ S, 23°44’ E; 1933; F.G. Overlaet leg.; RMCA • 2 ♀♀; Kapanga; 08°21’ S, 22°34’ E, Jan. 1933; F.G. Overlaet leg.; RMCA. – **Mongala** • 1 ♀; Bumba; 02°11’ N, 22°32’ E; 1939; Rév. P. Lootens leg.; RMCA. – **Nord-Kivu** • 1 ♀; PNA [Parc National Albert]; 00°03’ S, 29°30’ E; 24 Mar. 1954; P. Vanschuytbroeck & H. Synave leg.; RMCA. – **Tanganyika** • 1 ♀; Lukuga r. Niemba; 05°57’ S, 28°26’ E; Oct. 1917-Jan. 1918; Dr. Pons leg.; RMCA • 1 ♂; Tanganyika-Moero: Nyunzu; 05°57’ S, 28°01’ E; Jan. – Feb. 1934; H. Bomans leg.; RMCA.

#### *Coelioxys (Coelioxys) marchalli* Pasteels, 1968, *Coelioxys marchalli* Friese

**Material.** Cupboard 110, Box 54 (1♂). **D.R. CONGO.** – **Haut-Lomami** • 1 ♂; PNU Kapero; 08°56’ S, 27°10’ E, alt. 1760 m; 13 Jan. 1948; Mis. G.F. de Witte leg.; RMCA.

#### *Coelioxys (Coelioxys) odin* Strand, 1912

**Material.** Cupboard 110, Box 53 (2♀♀ and ♂♂). **D.R. CONGO.** – **Bas-Uele** • 1 ♀; Bas-Uele; 04°00’ N, 22°15’ E; Jul.-Aug. 1920; L. Burgeon leg.; RMCA • 1 ♂; Uele: Bambesa; 03°28’ N, 25°43’ E; 30 Oct. 1933; J.V. Leroy leg.; RMCA. – **Haut-Lomami** • 1 ♀; Lualaba: Kabongo; 07°20’ S, 25°35’ E; 1934; Ch. Seydel leg.; RMCA. – **Kinshasa** • 1 ♂; Barumbu; 01°14’ S, 23°31’ E; Aug. 1925; S.A.R. Prince Léopold leg.; RMCA.

#### *Coelioxys (Coelioxys) planidens* Friese, 1904

**Material.** Cupboard 110, Box 53 (5♀♀ and 13♂♂). **D.R. CONGO.** – **Bas-Uele** • 1 ♂; Bambesa; 03°28’ N, 25°43’ E; 4 Nov. 1937; J. Vrydagh leg.; RMCA • 2 ♂♂; Uele: Tukpwo; 04°26’ N, 25°51’ E; Jul. 1937; J. Vrydagh leg.; RMCA. – **Haut-Katanga** • 1 ♂; Kambove-Ruwe; 10°52’ S, 26°38’ E; 2 Mar. 1907; Dr. Sheffield Neave leg.; RMCA. – **Equateur** • 1 ♀; Eala (Mbandaka); 00°03’ N, 18°19’ E; 14 Mar. 1933; A. Corbisier leg.; RMCA • 1 ♀; same location as for preceding; 4 Apr. 1932; H.J. Brédo leg.; RMCA • 1 ♀; same location as for preceding; Mar. 1932; H.J. Brédo leg.; RMCA • 3 ♂♂; same location as for preceding; Mar. 1932; H.J. Brédo leg.; RMCA • 3 ♂♂; same location as for preceding; Jun. 1932; A. Corbisier leg.; RMCA • 2 ♀♀; same location as for preceding; Nov. 1931; H.J. Brédo leg.; RMCA • 1 ♂; same location as for preceding; 4 Apr. 1937; J. Vrydagh leg.; RMCA. – **Kongo Central** • 1 ♂; Congo da Lemba (Songololo); 05°42’ S, 13°42’ E; Jan.-Feb. 1913; R. Mayné leg.; RMCA. – **Tshuapa** • 1 ♂; Bokuma; 00°06’ S, 18°41’ E; Jan. – Feb. 1952; Rév. P. Lootens leg.; RMCA.

#### *Coelioxys (Coelioxys) postponenda* Schulz, 1906

**Material.** Cupboard 110, Box 53 (3♂♂). **D.R. CONGO.** – **Equateur** • 1 ♂; Eala (Mbandaka); 00°03’ N, 18°19’ E; 7 Oct. 1931; H.J. Brédo leg.; RMCA. – **Haut-Uele** • 1 ♂; Mayumbe: Bunda Suindi; 04°54’ S, 13°05’ E; 15 May 1926; A. Collart leg.; RMCA. – **Kasaï** • 1 ♂; Luebo; 05°20’ S, 21°24’ E; 7 Aug. 1921; Lt. J. Ghesquière leg.; RMCA.

#### *Coelioxys (Coelioxys) recusata* Schulz, 1904

**Material.** Cupboard 110, Box 52 (1♂). **D.R. CONGO.** – **Ituri** • 1 ♂; Mahagi-Niarembe; 02°15’ N, 31°07’ E; 1935; Ch. Scops leg.; RMCA.

#### *Coelioxys (Coelioxys) setosa* Friese, 1904

**Material.** Cupboard 110, Box 52 (1♂). **D.R. CONGO.** – **Haut-Katanga** • 1 ♀; Kiamanwa; 09°30’ S, 27°57’ E; Feb. 1931; H.J. Brédo leg.; RMCA. – **Haut-Lomami** • 1 ♂; PNU R. Kapero af. Kafwi; PNU Riv. Lusinga, alt. 1700 m; 21 Jan. 1948; Mis. G.F. de Witte leg.; RMCA. – **Haut-Lomami** • 1 ♂; Mulungu: Tshibinda; 02°19’ S, 28°45’ E; Nov. 1951; P.C. Lefèvre leg.; RMCA.

#### *Coelioxys (Liothyrapis) gracilis* Pasteels, 1968, *Coelioxys* (*Hemicoelioxys*) *gracilis* Pasteels, 1968

**Material.** Cupboard 110, Box 56 (1♀). **D.R. CONGO.** Holotype – **Haut-Katanga** • 1 ♀; Elisabethville (Lubumbashi); 11°40’ S, 27°28’ E; Dec 1934; P. Quarré leg.; RMCA.

#### *Coelioxys (Liothyrapis) junodi* Friese, 1904, *Liothyrapis junodi* Pasteels, 1962

**Material.** Cupboard 110, Box 55A (1♂). **D.R. CONGO.** – **Equateur** • 1 ♂; Eala (Mbandaka); 00°03’ N, 18°19’ E; Mar. 1932; H.J. Brédo leg.; RMCA.

#### *Coelioxys (Liothyrapis) scioensis* Gribodo, 1879, *Liothyrapis scioensis* (Gribodo, 1879)

**Material.** Cupboard 110, Box 55 (12♀♀). **D.R. CONGO.** – **Equateur** • 1 ♀; Eala (Mbandaka); 00°03’ N, 18°19’ E; Jul. 1935; J. Ghesquière leg.; RMCA. – **Haut-Katanga** • 1 ♀; Elisabethville (Lubumbashi); 11°40’ S, 27°28’ E; 1932; De Loose leg.; RMCA • 1 ♀; same location as for preceding; 2 May 1933; Dr. M. Bequaert leg.; RMCA • 1 ♀; same location as for preceding; Mar. 1935; P. Quarré leg.; RMCA • 1 ♀; same location as for preceding; De Loose leg.; RMCA • 1 ♀; same location as for preceding; Dec. 1933; Dr. M. Bequaert leg.; RMCA • 1 ♀; same location as for preceding; Oct. – Dec. 1935; P. Quarré leg.; RMCA • 1 ♀; Munama; 11°40’ S, 27°28’ E; Dec. 1930; Ch. Seydel leg.; RMCA. – **Lualaba** • 1 ♀; Kapanga; 08°21’ S, 22°34’ E; Feb. 1933; F.G. Overlaet leg.; RMCA • 1 ♀; Lulua: Kanzenze; 10°31’ S, 25°12’ E; 1932; F.G. Overlaet leg.; RMCA. – **Nord-Kivu** • 1 ♀; Kivu: Butembo; 00°09’ N, 29°17’ E, alt. 1740 m; May 1965; C. Leontovitch leg.; RMCA. – **Sud-Kivu** • 1 ♀; Mulungu: Tshibinda; 02°19’ S, 28°45’ E; Nov. 1951; P.C. Lefèvre leg.; RMCA.

#### *Coelioxys (Liothyrapis) subdentata* Smith, 1854, *Liothyrapis subdentata* (Smith, 1854)

**Material.** Cupboard 110, Box 55A (2♀♀). **D.R. CONGO.** – **Haut-Katanga** • 1 ♀; Elisabethville (Lubumbashi); 11°40’ S, 27°28’ E; Sep. 1929; Ch. Seydel leg.; RMCA. – **Ituri** • 1 ♀; Bunia; 01°34’ N, 30°15’ E; Fev. 1934; J.V. Leroy leg.; RMCA.

#### *Coelioxys (Liothyrapis) torrida* Smith, 1854, *Liothyrapis* (*Torridapis*) *torrida* (Smith, 1854)

**Material.** Cupboard 110, Box 55 (22♀♀ and 7♂♂). **D.R. CONGO.** – **Bas-Uele** • 1 ♀; Bambesa; 03°28’ N, 25°43’ E; Dec. 1933; H.J. Brédo leg.; RMCA • 1 ♀; same location as for preceding; Feb. 1934; H.J. Brédo leg.; RMCA • 1 ♂; Uele: Tukpwo; 04°26’ N, 25°51’ E; Jul. 1937; J. Vrydagh leg.; RMCA. – **Equateur** • 1 ♀; Eala (Mbandaka); 00°03’ N, 18°19’ E; Mar. 1932; H.J. Brédo leg.; RMCA • 3 ♂♂; same location as for preceding; Mar. 1932; H.J. Brédo leg.; RMCA • 1 ♀; same location as for preceding; May 1932; H.J. Brédo leg.; RMCA. – **Haut-Lomami** • 1 ♀; Lualaba: Kabongo; 07°20’ S, 25°35’ E; 27 Dec. 1952; Ch. Seydel leg.; RMCA • 1 ♀; same location as for preceding; 3 Jan. 1953; Ch. Seydel leg.; RMCA • 2 ♀♀; same location as for preceding; 31 Dec. 1952; Ch. Seydel leg.; RMCA • 1 ♂; same location as for preceding; 31 Dec. 1952; Ch. Seydel leg.; RMCA • 1 ♂; same location as for preceding; Nov. 1953; Ch. Seydel leg.; RMCA • 1 ♂; same location as for preceding; 3 Jan. 1953; F.G. Overlaet leg.; RMCA. – **Haut-Uele** 1 ♀; PNG [Parc National de Garamba]; 03°40’ N, 29°00’ E; 9 Nov 1950; H. De Saeger leg.; RMCA. – **Ituri** • 1 ♀; Lac Albert: Ishwa; 02°13’ N, 31°12’ E; Sep. 1935; H.J. Brédo leg.; RMCA. – **Kasaï** • 1 ♀; Tshikapa; 06°24’ S, 20°48’ E; Apr. 1939; Mevr. Bequaert leg.; RMCA. – **Kasaï Central** • 1 ♀; Luluabourg (Kananga); 05°54’ S, 21°52’ E; 18 May 1919; P. Callewaert leg.; RMCA. – **Kinshasa** • 1 ♀; Léopoldville (Kinshasa); 04°19’ S, 15°19’ E; Dec. 1911; Dr. Mouchet leg.; RMCA. – **Kongo Central** • 2 ♀♀; Boma; 05°51’ S, 13°03’ E; 1935; W. Moreels leg.; RMCA • 1 ♀; same location as for preceding; 1937; Dr. Schlesser leg.; RMCA • 1 ♀; Kisantu (Madimba); 05°08’ S, 15°06’ E; 1931; R.P. Vanderyst leg.; RMCA. – **Lualaba** • 1 ♀; Lulua: Kapanga; 08°21’ S, 22°34’ E; 11 Dec. 1932; F.G. Overlaet leg.; RMCA •1 ♀; same location as for preceding; 8 Dec. 1932; F.G. Overlaet leg.; RMCA. – **Maï-Ndombe** • 1 ♀; Kwamouth (Mushie); 03°11’ S, 16°12’ E; Jun. 1913; Dr. J. Maes leg.; RMCA. – **Maniema** • 1 ♀; Kisamba; 09°12’ S, 25°51’ E; Mar. 1955; Dr. J. Claessens leg.; RMCA. – **Sankuru** • 1 ♀; Lulua: Source Losaka; 04°58’ S, 23°26’ E; 9 Feb. 1932; F.G. Overlaet leg.; RMCA. – **Sud-Ubangi** • 1 ♀; Libenge; 03°39’ N, 18°38’ E; Aug. 1938; C. Leontovitch leg.; RMCA.

#### *Coelioxys (Liothyrapis) verticalis* Smith, 1854, *Liothyrapis verticalis* (Smith, 1854)

**Material.** Cupboard 110, Box 55A (64♀♀ and 4♂♂). **D.R. CONGO.** – **Equateur** • 1 ♀; Basankusu; 01°13’ N, 19°49’ E; 1949; Zusters O.L.V. ten Bunderen leg.; RMCA • 1 ♀; Coquilhatville (Mbandaka); 00°03’ N, 18°15’ E; 1927; Dr. Strada leg.; RMCA • 1 ♀; same location as for preceding; 1945; Rév. P. Hulstaert leg.; RMCA • 1 ♀; Eala (Mbandaka); 00°03’ N, 18°19’ E; 15-30 Nov. 1929; H.J. Brédo leg.; RMCA • 1 ♀; same location as for preceding; Apr. 1932; H.J. Brédo leg.; RMCA • 1 ♂; same location as for preceding; Apr. 1932; H.J. Brédo leg.; RMCA • 1 ♀; same location as for preceding; Sep. 1936; J. Ghesquière leg.; RMCA • 1 ♀; same location as for preceding; May 1935; J. Ghesquière leg.; RMCA • 1 ♀; Flandria; 00°20’ S, 19°06’ E; 1928; Rév. P. Hulstaert leg.; RMCA. – **Haut-Katanga** • 1 ♀; Elisabethville (Lubumbashi); 11°40’ S, 27°28’ E; 2 Apr. 1928; Dr. M. Bequaert leg.; RMCA • 1 ♀; same location as for preceding; Feb. 1935; P. Quarré leg.; RMCA • 1 ♀; same location as for preceding; De Loose leg.; RMCA • 1 ♀; same location as for preceding; Nov. 1934; P. Quarré leg.; RMCA • 1 ♀; same location as for preceding; Dec. 1929; Dr. M. Bequaert leg.; RMCA • 1 ♀; Kampombwe; 11°40’ S, 27°28’ E; Apr. 1931; H.J. Brédo leg.; RMCA • 1 ♀; Kipushi; 11°46’ S, 27°15’ E; Sep. 1931; Ch. Seydel leg.; RMCA. – **Haut-Lomami** • 1 ♀; Kaniama; 07°31’ S, 24°11’ E; May 1932; R. Massart leg.; RMCA • 1 ♀; Lomami: Kamina; 08°44’ S, 25°00’ E; 1930; R. Massart leg.; RMCA • 1 ♀; Lomami: Kaniama; 07°31’ S, 24°11’ E; 1931; R. Massart leg.; RMCA • 1 ♀; same location as for preceding; Mar. – Apr. 1932; R. Massart leg.; RMCA • 1 ♀; same location as for preceding; Apr. 1932; R. Massart leg.; RMCA • 1 ♀; Lualaba: Kaniama; 07°31’ S, 24°11’ E; 15 Dec. 1952; Ch. Seydel leg.; RMCA. – **Haut-Uele**• 1 ♀; Mayumbe: Buku-Lobe; 05°30’ S, 12°53’ E; 22 Nov. 1925; A. Collart leg.; RMCA • 1 ♀; PNG [Parc National de Garamba]; 03°40’ N, 29°00’ E; 15 May 1950; H. De Saeger leg.; RMCA • 1 ♀; Uele: Faradje; 03°44’ N, 29°42’ E; Nov. 1912; Lang & Chapin leg.; RMCA. – **Kasaï** • 1 ♀; Kasaï: Mubanga; 04°20’ S, 21°20’ E; 5 May 1946; V. Lagae leg.; RMCA • 1 ♀; Port-Francqui (Ilebo); 04°20’ S, 20°35’ E; Oct. 1937; Mme. Gillardin leg.; RMCA • 1 ♀; Tshikapa; 06°24’ S, 20°48’ E; Apr. 1939; Mevr. Bequaert leg.; RMCA. – **Kasaï Central** • 3 ♀♀; Gandu; 05°30’ S, 22°00’ E; Apr. 1931; H.J. Brédo leg.; RMCA • 1 ♀; Limbala; 06°07’ S, 22°20’ E; 5 – 8 Aug. 1913; Dr. Rodhain leg.; RMCA • 1♀; Lulua; 05°54’ S, 22°25’ E; 1929; Dr. Walker leg.; RMCA • 1 ♀; Lulua: Riv. Luele; 06°22’ S, 23°51’ E; 1929; Dr. Walker leg.; RMCA • 1 ♀; Luluabourg (Kananga); 05°54’ S, 21°52’ E; 18 May 1919; P. Callewaert leg.; RMCA • 2 ♂♂; same location as for preceding; P. Callewaert leg.; RMCA. – **Kongo Central** • 1 ♂; Kisantu (Madimba); 05°08’ S, 15°06’ E; 1927; R.P. Vanderyst leg.; RMCA • 1 ♀; same location as for preceding; 1932; R.P. Vanderyst leg.; RMCA • 1 ♀; Mayidi (Madimba); 05°11’ S, 15°09’ E; 1945; R.P. Vanderyst leg.; RMCA. – **Kwilu** • 1 ♀; Leverville (Bulungu); 04°50’ S, 18°44’ E; 1928; Mme. J. Tinant leg.; RMCA 1 ♀; Terr. de Banningville (Bandundu); 03°18’ S, 17°21’ E; Apr. 1946; Dr. Fain leg.; RMCA. – **Lualaba** • 2 ♀♀; Kapanga; 08°21’ S, 22°34’ E; Nov. 1934; F.G. Overlaet leg.; RMCA • 2 ♀♀; Lulua: Kapanga; 08°21’ S, 22°34’ E; 11 Dec. 1932; F.G. Overlaet leg.; RMCA • 1 ♀; same location as for preceding; 8 Dec. 1932; F.G. Overlaet leg.; RMCA • 1 ♀; same location as for preceding; Dec. 1932; F.G. Overlaet leg.; RMCA. – **Maï-Ndombe** • 3 ♀♀; Yumbi; 01°54’ S, 16°33’ E; 29 Jul. 1912; Dr. Mouchet leg.; RMCA. – **Maniema** • 2 ♀♀; Kindu; 02°57’ S, 25°55’ E; Dr. Mouchet leg.; RMCA • 1 ♀; Malela; 04°22’ S, 26°08’ E; Dec. 1913; L. Burgeon leg.; RMCA. – **Tanganyika** • 1 ♀; Bassin Lukuga; 05°40’ S, 26°55’ E; Apr. – Jul. 1934; H. De Saeger leg.; RMCA • 1 ♀; Kiambi (Manono); 07°19’ S, 28°01’ E; 18 – 24 Apr. 1931; G.F. de Witte leg.; RMCA. – **Tshopo** • 1 ♀; Mfungwe-Kayumbe; 01°17’ S, 26°16’ E; Jun. 1907; Dr. Sheffield Neave leg.; RMCA • 2 ♀♀; Terr. de Basoko; 01°13’ N, 23°36’ E; 1950; R.P. Camps leg.; RMCA. – **Tshuapa** • 1 ♀; Bokuma; 00°06’ S, 18°41’ E; Feb. – Apr. 1941; Rév. P. Lootens leg.; RMCA • 1 ♀; same location as for preceding; Jul. 1952; Rév. P. Lootens leg.; RMCA • 3 ♀♀; same location as for preceding; Dec. 1952; Rév. P. Lootens leg.; RMCA • 1 ♀; Tshuapa: Bamania (Mbandaka); 00°01’ N, 18°19’ E; Apr. 1960; Rév. P. Hulstaert leg.; RMCA • 1 ♀; same location as for preceding; Oct. 1951; Rév. P. Hulstaert leg.; RMCA • 1 ♀; Tshuapa: Etata; 00°14’ S, 20°43’ E; May 1970; J. Hauwaerts leg.; RMCA.

#### *Coelioxys (Torridapis) maculata* Friese, 1913, *Liothyrapis* (*Torridapis*) *maculata* (Friese, 1913)

**Material.** Cupboard 110, Box 55 (1♀ and 3♂♂). **D.R. CONGO.** – **Haut-Katanga** • 1 ♂; Elisabethville (Lubumbashi); 11°40’ S, 27°28’ E; Nov. 1938; De Loose leg.; RMCA. – **Kongo Central** • 1 ♀; Congo da Lemba (Songololo); 05°42’ S, 13°42’ E; Apr. 1920; R. Mayné leg.; RMCA • 1 ♂; Kisantu (Madimba); 05°08’ S, 15°06’ E; 1931; R.P. Vanderyst leg.; RMCA.

#### *Gronoceras africanibium* (Strand, 1912), *Chalicodoma* (*Gronoceras*) *africanibia* (Strand, 1912)

**Material.** Cupboard 110, Box 44 (12♀♀). **D.R. CONGO.** – **Equateur** • 1 ♀; Eala (Mbandaka); 00°03’ N, 18°19’ E; 15-30. Nov. 1929; H.J. Brédo leg.; RMCA • 1 ♀; same location as for preceding; May 1935; J. Ghesquière leg.; RMCA • 1 ♀; Wendje; 03°59’ S, 23°34’ E; 30 Sep. 1922; Dr. M. Bequaert leg.; RMCA. **Ituri** • 1 ♀; Lac Albert: Ishwa; 02°13’ N, 31°12’ E; Sep. 1935; H.J. Brédo leg.; RMCA. **Sankuru** • 1 ♀; Sankuru: Komi; 03°23’ S, 23°46’ E; May 1930; J. Ghesquière leg.; RMCA. **Tshuapa** • 1 ♀; Likete s/Lomela; 00°43’ S, 21°24’ E; Jun. 1936; J. Ghesquière leg.; RMCA • 2 ♀♀; Likete (Boende); 00°43’ S, 21°24’ E; 15 Juin 1934; J. Ghesquière leg.; RMCA • 3 ♀♀; same location as for preceding; 15 Jun. 1936; J. Ghesquière leg.; RMCA • 1 ♀; Tshuapa: Bokuma; 00°06’ S, 18°41’ E; 1953; Rév. P. Lootens leg.; RMCA.

#### *Gronoceras bombiforme* (Gerstaecker, 1857), *Chalicodoma* (*Gronoceras*) *bombiformis* (Gerstaecker, 1857)

**Material.** Cupboard 110, Box 34 (13♀♀ and 10♂♂). **D.R. CONGO.** – **Haut-Katanga** • 1 ♀; Elisabethville (Lubumbashi); 11°40’ S, 27°28’ E; 3 Feb. 1924; Ch. Seyedel leg.; RMCA • 1 ♀; same location as for preceding; Feb. 1933; De Loose leg.; RMCA. – **Haut-Lomami** • 1 ♂; PNU Mabwe (R.P. Lac Upemba); 08°47’ S, 26°52’ E, alt. 585 m; 1-11 Oct. 1949; Mis. G.F. de Witte leg.; RMCA. – **Kinshasa**• 1 ♀; Kalina; 04°18’ S, 15°17’ E; Jul. 1945; Mme. Delsaut leg.; RMCA • 1 ♀; Léopoldville (Kinshasa); 04°19’ S, 15°19’ E; 17 Apr. 1911; Dr. Mouchet leg.; RMCA • 1 ♀; Léopoldville-Kalina; 04°19’ S, 15°19’ E; Aug. 1945; Mme. Delsaut leg.; RMCA. – **Kongo Central** • 1 ♂; Kisantu (Madimba); 05°08’ S, 15°06’ E; Rév. P. Regnier leg.; RMCA • 1 ♀; Mayidi (Madimba); 05°11’ S, 15°09’ E; 1942; Rév. P. Van Eyen leg.; RMCA • 1 ♀; Mayumbe; 05°08’ S, 12°29’ E; Cabra leg.; RMCA • 1 ♀; same location as for preceding; R. Verschueren leg.; RMCA • 1 ♀; Moanda; 05°56’ S, 12°21’ E; 28 Apr. 1970; Mme. Delsaut leg.; RMCA. – **Kwilu** • 1 ♀; Kolo-Kwilu-Madiata; 05°27’ S, 14°52’ E; Sep. 1913; R. Verschueren leg.; RMCA • 1 ♂; same location as for preceding; Sep. 1913; R. Verschueren leg.; RMCA • 1 ♀; Leverville (Bulungu); 04°50’ S, 18°44’ E; 1928; Mme. J. Tinant leg.; RMCA. – **Lualaba** • 1 ♂; Dilolo; 10°41’ S, 22°21’ E; Aug. 1931; G.F. de Witte leg.; RMCA • 1 ♂; Haut-Luapula: Kansenia; 10°19’ S, 26°04’ E; 14 Oct. 1929; Dom de Montpellier leg.; RMCA • 1 ♂; same location as for preceding; 3 Nov. 1929; Dom de Montpellier leg.; RMCA • 1 ♂; Kansenia; 10°19’ S, 26°04’ E; 15 Sep. 1915-Oct. 1930; G.F. de Witte leg.; RMCA • 1 ♀; Lulua: Kapanga; 08°21’ S, 22°34’ E; Nov. 1932; F.G. Overlaet leg.; RMCA • 1 ♂; Sandu; 09°41’ S, 22°53’ E; Apr. 1931; H.J. Brédo leg.; RMCA. – **Tanganyika** • 1 ♂; Kanikiri; 00°09’ S, 29°32’ E; Oct. 1907; Dr. Sheffield Neave leg.; RMCA • 1 ♂; Vallée Lukuga; 05°40’ S, 26°55’ E; Nov. 1911; Dr. Schwetz leg.; RMCA.

#### *Gronoceras chapini* (Cockerell, 1935), Chalicodoma (*Gronoceras*) chapini (Cockerell, 1935)

**Material.** Cupboard 110, Boxe 34 (1♀). **D.R. CONGO.** – **Equateur** • 1 ♀; Eala (Mbandaka); 00°03’ N, 18°19’ E; 1932; Mar. 1932. H.J. Brédo leg.; RMCA.

#### *Gronoceras cinctum* (Fabricius, 1781), *Chalicodoma* (*Gronoceras*) *cincta* (Fabricius, 1781)

**Material.** Cupboard 110, Boxes 35, 36, 37, 38, 39, 40, 41 and 42 (947♀♀ and 323♂♂). **D.R. CONGO.** – **Bas-Uele** • 3 ♀♀; Bambesa; 03°28’ N, 25°43’ E; 15 Sep. 1933; H.J. Brédo leg.; RMCA • 1 ♀; same location as for preceding; 30 Oct. 1933; H.J. Brédo leg.; RMCA • 1 ♀; same location as for preceding; Dec. 1933; H.J. Brédo leg.; RMCA • 1 ♀; same location as for preceding; 6 Jul. 1937; J. Vrydagh leg.; RMCA • 1 ♀; same location as for preceding; Oct. 1938; P. Lefèvre leg.; RMCA • 1 ♀; same location as for preceding; 10 Sep. 1937; R. Verschueren leg.; RMCA • 1 ♀; same location as for preceding; 11 Jul. 1937; R. Verschueren leg.; RMCA • 1 ♀; same location as for preceding; 14 May 1938; R. Verschueren leg.; RMCA • 2 ♀♀; same location as for preceding; 15 Sep. 1933; R. Verschueren leg.; RMCA • 1 ♀; same location as for preceding; 15 Oct. 1923; R. Verschueren leg.; RMCA • 2 ♀♀; same location as for preceding; 16 Mar. 1940; R. Verschueren leg.; RMCA • 1 ♀; same location as for preceding; 16 Apr. 1937; R. Verschueren leg.; RMCA • 1 ♂; same location as for preceding; 21 Jan. 1933; R. Verschueren leg.; RMCA • 1 ♀; same location as for preceding; 24 Jun. 1937; R. Verschueren leg.; RMCA • 1 ♂; same location as for preceding; 3 Jul. 1937; R. Verschueren leg.; RMCA • 1 ♂; same location as for preceding; 4 May 1937; R. Verschueren leg.; RMCA • 3 ♀♀; same location as for preceding; 6 Jul. 1937; R. Verschueren leg.; RMCA • 9 ♀♀; same location as for preceding; 9 May 1938; R. Verschueren leg.; RMCA • 1 ♀; same location as for preceding; Feb. 1934; R. Verschueren leg.; RMCA • 1 ♀; same location as for preceding; Jun. 1932; R. Verschueren leg.; RMCA • 1 ♀; same location as for preceding; Jun. 1937; R. Verschueren leg.; RMCA • 1 ♀; same location as for preceding; Jul. 1937; R. Verschueren leg.; RMCA • 1 ♂; same location as for preceding; May – Jul. 1937; R. Verschueren leg.; RMCA • 1 ♀; Uele; 04°06’ N, 22°23’ E; 1912; A. Dufrasne leg.; RMCA • 1 ♀; same location as for preceding; Jun. 1932; Degreef leg.; RMCA • 1 ♀; Uele: Bambesa; 03°28’ N, 25°43’ E; 24 Sep. 1943; J. Vrydagh leg.; RMCA • 1 ♀; same location as for preceding; 9 May 1940; J. Vrydagh leg.; RMCA • 1 ♀; same location as for preceding; 20 Oct. 1933; J.V. Leroy leg.; RMCA • 5 ♀♀; same location as for preceding; 30 Oct. 1933; J.V. Leroy leg.; RMCA • 1 ♀; same location as for preceding; 30 Oct. 1933; R. Verschueren leg.; RMCA • 1 ♂; same location as for preceding; 30 Oct. 1933; R. Verschueren leg.; RMCA • 1 ♀; Uele: Bambili; 03°38’ N, 26°06’ E; Degreef leg.; RMCA • 1 ♀; Uele: Baye, terr. Bondo; 03°47’ N, 23°48’ E; Aug. 1956; R. Fr. L. Rooyakkers leg.; RMCA • 1 ♂; Uele: Bondo; 03°47’ N, 23°48’ E; 28 Feb. 1950; R.P. Theunissen leg.; RMCA • 1 ♀; Uele: Dingila; 03°37’ N, 26°03’ E; May 1933; H.J. Brédo leg.; RMCA • 1 ♀; same location as for preceding; 1 Jun. 1933; J.V. Leroy leg.; RMCA • 1 ♀; same location as for preceding; 15 Jul. 1933; J.V. Leroy leg.; RMCA • 1 ♀; Uele: Ibembo; 02°38’ N, 23°36’ E; 1950; R. Fr. Hutsebaut leg.; RMCA • 2 ♀♀; same location as for preceding; 12 Feb. 1950; J.V. Leroy leg.; RMCA • 1 ♀; same location as for preceding; Apr. 1952; J.V. Leroy leg.; RMCA • 2 ♀♀; same location as for preceding; Oct. – Nov. 1951; J.V. Leroy leg.; RMCA • 9 ♀♀; Uele: Tukpwo; 04°26’ N, 25°51’ E; Jul. 1937; J. Vrydagh leg.; RMCA • 2 ♀♀; same location as for preceding; Aug. 1937; J. Vrydagh leg.;

RMCA. – **Equateur** • 1 ♀; Bangala: Lubao; 01°36’ N, 19°09’ E; 1928; Mme. Babilon leg.; RMCA • 1 ♂; Basankusu; 01°13’ N, 19°49; 10 Oct. 1929; De Coninak leg.; RMCA • 1 ♀; same location as for preceding; 1949; Zusters O.L.V. ten Bunderen leg.; RMCA • 1 ♀; Boyeka (Eyala); 00°03’ N, 18°19’ E; 6 Jan. 1915; R. Mayné leg.; RMCA • 5 ♀♀; Coquilhatville (Mbandaka); 00°03’ N, 18°15’ E; 1946; Ch. Scops leg.; RMCA • 2 ♂♂; same location as for preceding; 1946; Ch. Scops leg.; RMCA • 1 ♂; same location as for preceding; 1921; Dr. M. Bequaert leg.; RMCA • 1 ♀; same location as for preceding; 24 Jun. 1924; Dr. M. Bequaert leg.; RMCA • 1 ♀; same location as for preceding; 24 Nov. 1924; Dr. M. Bequaert leg.; RMCA • 1 ♀; same location as for preceding; 1927; Strada leg.; RMCA • 1 ♀; same location as for preceding; 6 Sep. 1948; R. Mouchamps leg.; RMCA • 2 ♀♀; same location as for preceding; 1945; Rév. P. Hulstaert leg.; RMCA • 1 ♀; same location as for preceding; 1959; Rév. P. Hulstaert leg.; RMCA • 1 ♀; same location as for preceding; 18 Feb. 1925; Rév. P. Hulstaert leg.; RMCA • 10 ♀♀; Eala (Mbandaka); 00°03’ N, 18°19’ E; 1932; A. Corbisier leg.; RMCA • 6 ♂♂; same location as for preceding; 1932; A. Corbisier leg.; RMCA • 2 ♀♀; same location as for preceding; 1933; A. Corbisier leg.; RMCA • 1 ♂; same location as for preceding; 1933; A. Corbisier leg.; RMCA • 14 ♀♀; same location as for preceding; 14 Mar. 1933; A. Corbisier leg.; RMCA • 1 ♂; same location as for preceding; 20 Apr. 1933; A. Corbisier leg.; RMCA • 4♀♀; same location as for preceding; 7 Jul. 1932; A. Corbisier leg.; RMCA • 4 ♂♂; same location as for preceding; 7 Jul. 1932; A. Corbisier leg.; RMCA • 16 ♀♀; same location as for preceding; Apr. 1933; A. Corbisier leg.; RMCA • 5 ♂♂; same location as for preceding; Apr. 1933; A. Corbisier leg.; RMCA • 88♀♀; same location as for preceding; Jun. 1932; A. Corbisier leg.; RMCA • 44 ♂♂; same location as for preceding; Jun. 1932; A. Corbisier leg.; RMCA •16 ♀♀; same location as for preceding; Nov. 1932; A. Corbisier leg.; RMCA •9 ♂♂; same location as for preceding; Nov. 1932; A. Corbisier leg.; RMCA • 1 ♀; same location as for preceding; Dec. 1932; A. Corbisier leg.; RMCA •1 ♀; same location as for preceding; Apr. 1933; Corbisier-baland leg.; RMCA • 5 ♂♂; same location as for preceding; Apr. 1933; Corbisier-baland leg.; RMCA •1 ♀; same location as for preceding; 11 Dec. 1931; H.J. Brédo leg.; RMCA •1 ♀; same location as for preceding; 13 Oct. 1931; H.J. Brédo leg.; RMCA •5 ♀♀; same location as for preceding; 13 Nov. 1931; H.J. Brédo leg.; RMCA •1 ♂; same location as for preceding; 13 Nov. 1931; H.J. Brédo leg.; RMCA •1 ♂; same location as for preceding; 14 Oct. 1931; H.J. Brédo leg.; RMCA •13 ♀♀; same location as for preceding; 14 Nov. 1931; H.J. Brédo leg.; RMCA •3 ♂♂; same location as for preceding; 14 Nov. 1931; H.J. Brédo leg.; RMCA • 2 ♀♀; same location as for preceding; 14 Dec. 1931; H.J. Brédo leg.; RMCA • 1 ♂; same location as for preceding; 14 Dec. 1931; H.J. Brédo leg.; RMCA • 1 ♂; same location as for preceding; 15 Nov. 1929; H.J. Brédo leg.; RMCA • 2 ♀♀; same location as for preceding; 15 Nov. 1931; H.J. Brédo leg.; RMCA • 1 ♀; same location as for preceding; 15 – 30 Oct. 1929; H.J. Brédo leg.; RMCA • 14 ♀♀; same location as for preceding; 15 – 30 Nov. 1929; H.J. Brédo leg.; RMCA • 2 ♂♂; same location as for preceding; 15 – 30 Nov. 1929; H.J. Brédo leg.; RMCA • 7 ♀♀; same location as for preceding; 16 Nov. 1929; H.J. Brédo leg.; RMCA • 1 ♂; same location as for preceding; 17 Nov. 1929; H.J. Brédo leg.; RMCA • 5 ♀♀; same location as for preceding; 17 Nov. 1931; H.J. Brédo leg.; RMCA • 1 ♀; same location as for preceding; 18 Nov. 1929; H.J. Brédo leg.; RMCA • 1 ♂; same location as for preceding; 18 – 26 Feb. 1932; H.J. Brédo leg.; RMCA • 1 ♀; same location as for preceding; 2 Oct. 1931; H.J. Brédo leg.; RMCA • 3 ♂♂; same location as for preceding; 2 Oct. 1931; H.J. Brédo leg.; RMCA • 1 ♀; same location as for preceding; 20 Oct. 1931; H.J. Brédo leg.;

RMCA • 26 ♀♀; same location as for preceding; 21 Nov. 1931; H.J. Brédo leg.; RMCA • 10 ♂♂; same location as for preceding; 21 Nov. 1931; H.J. Brédo leg.; RMCA • 1 ♀; same location as for preceding; 22 Apr. 1932; H.J. Brédo leg.; RMCA • 2 ♂♂; same location as for preceding; 22 Apr. 1932; H.J. Brédo leg.; RMCA • 2 ♀♀; same location as for preceding; 28 Nov. 1929; H.J. Brédo leg.; RMCA • 2 ♂♂; same location as for preceding; 28 Nov. 1929; H.J. Brédo leg.; RMCA • 5 ♀♀; same location as for preceding; 28 Nov. 1931; H.J. Brédo leg.; RMCA • 5 ♀♀; same location as for preceding; 3 Oct. 1931; H.J. Brédo leg.; RMCA • 2 ♂♂; same location as for preceding; 3 Oct. 1931; H.J. Brédo leg.; RMCA • 1 ♀; same location as for preceding; 3 Nov. 1931; H.J. Brédo leg.; RMCA • 1 ♂; same location as for preceding; 30 Nov. 1929; H.J. Brédo leg.; RMCA • 1 ♂; same location as for preceding; 4 Apr. 1932; H.J. Brédo leg.; RMCA • 1 ♀; same location as for preceding; 4 Nov. 1931; H.J. Brédo leg.; RMCA • 15 ♀♀; same location as for preceding; 5 Nov. 1931; H.J. Brédo leg.; RMCA • 5 ♂; same location as for preceding; 5 Nov. 1931; H.J. Brédo leg.; RMCA • 6 ♀♀; same location as for preceding; 6 Oct. 1931; H.J. Brédo leg.; RMCA • 6 ♀♀; same location as for preceding; 7 Nov. 1931; H.J. Brédo leg.; RMCA • 1 ♂; same location as for preceding; 7 Nov. 1931; H.J. Brédo leg.; RMCA • 1 ♂; same location as for preceding; 9 Nov. 1931; H.J. Brédo leg.; RMCA • 18 ♀♀; same location as for preceding; Mar. 1932; H.J. Brédo leg.; RMCA • 16 ♂♂; same location as for preceding; Mar. 1932; H.J. Brédo leg.; RMCA • 1 ♀; same location as for preceding; Mar. 1933; H.J. Brédo leg.; RMCA • 7 ♀♀; same location as for preceding; Apr. 1932; H.J. Brédo leg.; RMCA • 3 ♂♂; same location as for preceding; Apr. 1932; H.J. Brédo leg.; RMCA • 14 ♀♀; same location as for preceding; May 1932; H.J. Brédo leg.; RMCA • 12 ♂♂; same location as for preceding; May 1932; H.J. Brédo leg.; RMCA • 1 ♀; same location as for preceding; Oct. 1929; H.J. Brédo leg.; RMCA • 17 ♀♀; same location as for preceding; Nov 1931; H.J. Brédo leg.; RMCA • 7 ♂♂; same location as for preceding; Nov 1931; H.J. Brédo leg.; RMCA • 1 ♀; same location as for preceding; Dec. 1929; H.J. Brédo leg.; RMCA • 1 ♂; same location as for preceding; Dec. 1929; H.J. Brédo leg.; RMCA • 9 ♀♀; same location as for preceding; Dec. 1932; H.J. Brédo leg.; RMCA • 2 ♂♂; same location as for preceding; Dec. 1932; H.J. Brédo leg.; RMCA • 8 ♀♀; same location as for preceding; Mar. 1935; J. Ghesquière leg.; RMCA • 2 ♂♂; same location as for preceding; Mar. 1935; J. Ghesquière leg.; RMCA • 1♀; same location as for preceding; Apr. 1935; J. Ghesquière leg.; RMCA • 2 ♀♀; same location as for preceding; Sep. 1935; J. Ghesquière leg.; RMCA • 4 ♀♀; same location as for preceding; May 1935; J. Ghesquière leg.; RMCA • 1 ♀; same location as for preceding; Jul. 1935; J. Ghesquière leg.; RMCA • 1 ♀; same location as for preceding; Jul. 1936; J. Ghesquière leg.; RMCA • 3 ♀♀; same location as for preceding; Aug. 1935; J. Ghesquière leg.; RMCA • 1 ♀; same location as for preceding; Nov. 1934; J. Ghesquière leg.; RMCA • 1 ♀; same location as for preceding; Nov. 1935; J. Ghesquière leg.;

RMCA • 1 ♀; same location as for preceding; 16 Apr. 1937; J. Vrydagh leg.; RMCA • 1 ♀; same location as for preceding; 25 May 1938; P. Henrard leg.; RMCA • 1 ♀; same location as for preceding; Apr. 1936; P. Henrard leg.; RMCA • 1 ♀; same location as for preceding; Jul. 1936; P. Henrard leg.; RMCA • 1 ♀; same location as for preceding; Jul. 1938; P. Henrard leg.; RMCA • 1 ♀; same location as for preceding; 10 Nov. 1918; R. Mayné leg.; RMCA • 1 ♂; same location as for preceding; Sep. 1912; R. Mayné leg.; RMCA • 1 ♀; Eala: Boyeka; 00°03’ N, 18°19’ E; 30 Nov. 1929; H.J. Brédo leg.; RMCA • 1 ♀; Equateur: Bokote; 00°06’ S, 20°08’ E; 1 Feb. 1926; Rév. P. Hulstaert leg.; RMCA • 1 ♀; Kasaï: cité de Kombo (Ingende); 00°05’ S, 18°37’ E; 21 Mar. 1939; Mevr. Bequaert leg.; RMCA • 1 ♀; Lukolela (Bikoro); 01°03’ S, 17°12’ E; Nov. 1934; Dr. Ledoux leg.; RMCA • 3 ♂♂; same location as for preceding; Nov. 1934; Dr. Ledoux leg.; RMCA • 1 ♀; same location as for preceding; 1930; Dr. Obrassart leg.; RMCA • 1 ♂; Mobeka (Bomongo); 01°53’ N, 19°49’ E; 19 Mar. 1911; L. Burgeon leg.; RMCA • 1 ♂; Modu-Wamba; 00°36’ N, 20°07’ E; 5 Jun. 1931; H.J. Brédo leg.; RMCA • 1 ♂; Neder-Congo: Tumba; 00°50’ S, 18°00’ E; Br. Krist. Scholen leg.; RMCA • 1 ♀; Tshuapa: Basankusu; 01°13’ N, 19°49; 1948; Zusters O.L.V. ten Bunderen leg.;

RMCA. – **Haut-Katanga** • 1 ♀; Elisabethville (Lubumbashi); 11°40’ S, 27°28’ E; 5 Oct. 1923; Ch. Seydel leg.; RMCA • 1 ♀; same location as for preceding; May 1925; Ch. Seydel leg.; RMCA • 1 ♀; same location as for preceding; Nov. 1923; Ch. Seydel leg.; RMCA • 1 ♀; same location as for preceding; Nov. 1923; Ch. Seydel leg.; RMCA • 1 ♀; same location as for preceding; 1932; De Loose leg.; RMCA • 2 ♂♂; same location as for preceding; 1932; De Loose leg.; RMCA • 1 ♂; same location as for preceding; 29 Jul. 1932; De Loose leg.; RMCA • 1 ♀; same location as for preceding; Feb. 1932; De Loose leg.; RMCA • 1 ♀; same location as for preceding; Feb. 1933; De Loose leg.; RMCA • 2 ♂♂; same location as for preceding; Feb. 1933; De Loose leg.; RMCA • 2 ♀♀; same location as for preceding; Mar. 1933; De Loose leg.; RMCA • 1 ♂; same location as for preceding; Mar. 1933; De Loose leg.; RMCA • 4 ♂♂; same location as for preceding; De Loose leg.; RMCA • 1 ♂; same location as for preceding; 21 May 1933; Dr. M. Bequaert leg.; RMCA • 1 ♂; same location as for preceding; 23 Sep. 1923; Dr. M. Bequaert leg.; RMCA • 1 ♂; same location as for preceding; 24 Sep. 1932; Dr. M. Bequaert leg.; RMCA • 1 ♀; same location as for preceding; 24 Oct. 1933; Dr. M. Bequaert leg.; RMCA • 1 ♀; same location as for preceding; Mar. 1931; Dr. M. Bequaert leg.; RMCA • 2 ♀♀; same location as for preceding; Dec. 1933; Dr. M. Bequaert leg.; RMCA • 1 ♀; same location as for preceding; 7 Aug. 1921; Eg. Devroye leg.; RMCA • 1 ♀; same location as for preceding; Mar. 1938; H.J. Brédo leg.; RMCA • 1 ♀; same location as for preceding; May 1946; M. Lips leg.; RMCA • 1 ♀; same location as for preceding; Feb. 1934; P. Quarré leg.; RMCA • 1 ♀; same location as for preceding; Feb. 1935; P. Quarré leg.; RMCA • 2 ♀♀; same location as for preceding; Mar. 1935; P. Quarré leg.; RMCA • 1 ♀; same location as for preceding; Sep. 1934; P. Quarré leg.; RMCA • 4 ♀♀; same location as for preceding; May 1935; P. Quarré leg.; RMCA • 1 ♀; same location as for preceding; Jun. 1932; P. Quarré leg.; RMCA • 2 ♀♀; same location as for preceding; Jun. 1935; P. Quarré leg.; RMCA • 1 ♀; same location as for preceding; Jul. 1935; P. Quarré leg.; RMCA • 1 ♀; same location as for preceding; Jul. 1937; P. Quarré leg.; RMCA • 1 ♀; same location as for preceding; Aug. 1935; P. Quarré leg.; RMCA • 2 ♀♀; same location as for preceding; Oct. 1934; P. Quarré leg.; RMCA • 4 ♀♀; same location as for preceding; Nov. 1934; P. Quarré leg.; RMCA • 1 ♀; same location as for preceding; Dec. 1934; P. Quarré leg.; RMCA • 1 ♀; same location as for preceding; Nov. 1930; R. Massart leg.; RMCA • 1 ♀; Jadotville (Likasi); 10°59’ S, 26°44’ E; 1948; R.S.M. Adelaïde leg.; RMCA • 1 ♀; Kampombwe; 11°40’ S, 27°28’ E; Apr. 1931; H.J. Brédo leg.; RMCA • 1 ♀; Kapema; 10°42’ S, 28°40’ E; Sep. 1924; Ch. Seydel leg.; RMCA • 1 ♀; Katanga; 11°40’ S, 27°28’ E; Lemaire leg.; RMCA • 1 ♀; same location as for preceding; Weyms leg.; RMCA • 1 ♀; Katanga: Kasapa; 11°40’ S, 27°28’ E; 15 – 20 May 1965; W. Verheyen leg.; RMCA • 1 ♀; Katanga: Kasenga; 10°22’ S, 28°38’ E; Apr. 1931; H.J. Brédo leg.; RMCA • 1 ♀; Katanga: Kasinga; 10°22’ S, 28°38’ E; Oct. 1925; Ch. Seydel leg.; RMCA • 1 ♀; Katanga: Mufunga-Sampwe; 09°21’ S, 27°27’ E; 1951; R.S.M. Adelaïde leg.; RMCA • 1 ♀; same location as for preceding; 1955; R.S.M. Faber leg.; RMCA • 1 ♀; Katanga: Mwema; 08°13’ S, 27°28’ E; Jul. 1927; A. Corbisier leg.; RMCA • 1 ♀; Kilwa; 09°18’ S, 28°25’ E; Mar. 1931; H.J. Brédo leg.; RMCA • 1 ♀; Luanza; 10°21’ S, 27°51’ E; Mme. de Paeli leg.; RMCA • 1 ♀; Lubumbashi; 11°40’ S, 27°28’ E; 5 - 9 Oct. 1951; Ch. Seydel leg.; RMCA • 1 ♂; same location as for preceding; 13 Mar. 1921; Dr. M. Bequaert leg.; RMCA • 1 ♀; PNU Lusinga (riv. Kamitungulu); 08°55’ S, 08°55’ S; 13 Jun. 1945; G.F. de Witte leg.;

RMCA. – **Haut-Lomami** • 1 ♀; Katanga: Kilenge; 09°08’ S, 25°52’ E; Apr. 1923; A. Corbisier leg.; RMCA • 2 ♀♀; Katanga: Nyonga; 10°58’ S, 25°40’ E; 22 May 1925; G.F. de Witte leg.; RMCA • 1 ♀; Lomami: Kasese; 07°40’ S, 24°01’ E; Sep. 1948; Dr. Zielinski leg.; RMCA • 1 ♀; Lomami: Kishinde; 08°44’ S, 25°00’ E; Oct. 1931; P. Quarré leg.; RMCA • 1 ♂; Lomami: Mutombo-Mukulu; 07°58’ S, 24°00’ E; Jun. 1931; P. Quarré leg.; RMCA • 2 ♀♀; Lualaba: Kabongo; 07°20’ S, 25°35’ E; 27 Dec. 1952; Ch. Seydel leg.; RMCA • 2 ♂♂; same location as for preceding; 27 Dec. 1952; Ch. Seydel leg.; RMCA • 1 ♂; same location as for preceding; 30 Dec. 1952; Ch. Seydel leg.; RMCA • 1 ♂; same location as for preceding; 5 Jan. 1953; Ch. Seydel leg.; RMCA • 1 ♀; same location as for preceding; 7 Jan. 1953; Ch. Seydel leg.; RMCA • 2 ♂♂; same location as for preceding; 7 Jan. 1953; Ch. Seydel leg.; RMCA • 1 ♂; Sankisia; 09°24’ S, 25°48’ E; 29 Aug. 1911; Dr. J. Bequaert leg.; RMCA • 1 ♀; Sokole; 09°24’ S, 25°48’ E; May 1932 ; Dr. M. Bequaert leg.; RMCA. – **Haut-Uele** • 1 ♀; Dila (Haut-Uele); 02°46’ N, 27°38’ E; 1925; S.A.R. Prince Léopold leg.; RMCA • 1 ♀; Dungu-Nyangara Doruma; 03°37’ N, 28°34’ E; May 1912; Mme. Hutereau leg.; RMCA • 1 ♀; Haut-Uele: Moto; 03°03’ N, 29°28’ E; 1920; L. Burgeon leg.; RMCA • 1 ♀; same location as for preceding; L. Burgeon leg.; RMCA • 1 ♀; same location as for preceding; Oct. – Nov. 1923; L. Burgeon leg.; RMCA • 1 ♀; Ituri: Gangala-na-bodio; 03°41’ N, 29°08’ E; 30 Aug. 1931; H.J. Brédo leg.; RMCA • 1 ♀; Mayumbe: Zobe; 05°08’ S, 12°29’ E; Jan. 1916; R. Mayné leg.; RMCA • 1 ♀; PNG [Parc National de Garamba]; 03°40’ N, 29°00’ E; 23 Jul. 1951; H. De Saeger leg.; RMCA • 2 ♀♀; Uele: Dungu; 03°37’ N, 28°34’ E; Degreef leg.; RMCA • 1 ♀; Uele: Gangala-na-Bodio; 03°41’ N, 29°08’ E; Nov. 1956; Dr. M. Poll leg.; RMCA • 1 ♀; Wamba; 00°36’ N, 20°07’ E; 1936; Dr. Degotte leg.;

RMCA. – **Ituri** • 1 ♀; Bunia; 01°34’ N, 30°15’ E; 30 Jul. 1937; H.J. Brédo leg.; RMCA • 1 ♀; same location as for preceding; Mar. 1934; J.V. Leroy leg.; RMCA • 1 ♀; same location as for preceding; 1939; R.F. Maristetes leg.; RMCA • 1 ♂; same location as for preceding; 1938; Bastiaens leg.; RMCA • 1 ♀; Ituri: Fataki; 02°00’ N, 30°58’ E; 1938; J. Ghesquière leg.; RMCA • 1 ♀; Ituri: Forêt de Kawa; 01°34’ N, 30°32’ E; 13 Apr. 1929; A. Collart leg.; RMCA • 1 ♀; Ituri: Irumu; 01°27’ N, 29°52’ E; 26 Sep. 1931; Mme. L. Lebrun leg.; RMCA • 1 ♀; Kasenyi; 01°24’ N, 30°26’ E; Dec. 1938; H.J. Brédo leg.; RMCA • 2 ♂♂; same location as for preceding; Mar. 1934; P. Lefèvre leg.; RMCA • 1 ♀; Kibali-Ituri: Abock; 02°00’ N, 31°00’ E; 2 Oct. 1935; Ch. Scops leg.; RMCA • 1 ♀; Kibali-Ituri: Geti; 01°12’ N, 30°11’ E; Apr. 1939; R. Randour leg.; RMCA • 1 ♀; Kibali-Ituri: Kilo; 01°50’ N, 30°09’ E; 1930; Mlle. Jordens leg.; RMCA • 2 ♀♀; Kibali-Ituri: Kilomines; 01°50’ N, 30°09’ E; 11 Jan. 1957; C. Smoor leg.; RMCA • 1 ♀; same location as for preceding; Feb. 1956; C. Smoor leg.; RMCA • 1 ♀; same location as for preceding; Sep. 1957; C. Smoor leg.; RMCA • 1 ♀; same location as for preceding; 1955; R. Andry leg.; RMCA • 1 ♂; Kibali-Ituri: Mahagi; 02°18’ N, 30°59’ E; 1934; Ch. Scops leg.; RMCA • 1 ♀; same location as for preceding; 23 Jul. 1931; Ch. Scops leg.; RMCA • 1 ♀; Kilo: Kere-Kere; 02°42’ N, 30°33’ E; 1949; Dr. Turco leg.; RMCA • 1 ♀; same location as for preceding; L. Burgeon leg.; RMCA • 2 ♀♀; Lac Albert: Ishwa; 02°13’ N, 31°12’ E; Sep. 1935; H.J. Brédo leg.; RMCA • 1 ♀; same location as for preceding; Nov. 1935; H.J. Brédo leg.; RMCA • 2 ♀♀; Lac Albert: Kasenyi; 01°24’ N, 30°26’ E; Sep. 1935; H.J. Brédo leg.; RMCA • 3 ♂♂; same location as for preceding; Sep. 1935; H.J. Brédo leg.; RMCA • 1 ♂; Mahagi-Niarembe; 02°15’ N, 31°07’ E; 4 ; Sep. 1935; Ch. Scops leg.; RMCA • 2 ♀♀; same location as for preceding; May 1935; Ch. Scops leg.; RMCA • 1 ♂; same location as for preceding; 14 Sep. 1934; H.J. Brédo leg.; RMCA • 1 ♂; Mahagi-Port; 02°09’ N, 31°14’ E; Sep. 1934; A. Corbisier leg.; RMCA • 1 ♀; Mongbwalu (Kilo); 01°50’ N, 30°09’ E; 1930; Mme. E. Milliau leg.;

RMCA. – **Kasaï** • 1 ♀; Tshikapa; 06°24’ S, 20°48’ E; May 1921; Dr. H. Schouteden leg.; RMCA • 1 ♀; same location as for preceding; Apr. 1939; Mevr. Bequaert leg.; RMCA • 2 ♀♀; same location as for preceding; Apr. – May 1939; Mevr. Bequaert leg.; RMCA. – **Kasaï Central** • 1 ♀; Kasaï: Lula, Terr. Luisa; 07°11’ S, 22°24’ E; Aug. 1956; Dr. M. Poll leg.; RMCA • 1 ♀; Limbala; 06°07’ S, 22°20’ E; 5 Aug. 1913; Dr. Rodhain leg.; RMCA • 1 ♀; Lula (Kasaï); 07°11’ S, 22°24’ E; 1958; A.J. Jobaert leg.; RMCA • 2 ♀♀; same location as for preceding; Jun. 1919; A.J. Jobaert leg.; RMCA • 1 ♀; same location as for preceding; 20 Apr. May 1939; J.J. Deheyn leg.; RMCA • 1 ♀; same location as for preceding; P. Callewaert leg.; RMCA • 1 ♂; same location as for preceding; P. Callewaert leg.; RMCA • 1 ♀; same location as for preceding; Jul. 1936; P. Callewaert leg.; RMCA • 2 ♀♀; same location as for preceding; 1936; Puissant leg.; RMCA • 2 ♂♂; Luluabourg (Kasaï); 05°54’ S, 21°52’ E; 21 Jan. 1963; Jan Deheegher leg.; RMCA. – **Kasaï Oriental** • 1 ♀; Sankuru: Bakwanga; 06°08’ S, 23°38’ E; 12 Jun. 1939; Mevr. Bequaert leg.; RMCA • 1 ♀; same location as for preceding; Jun. 1939; Mevr. Bequaert leg.; RMCA • 1 ♀; Sankuru: Bakwanga (Dibindi); 06°08’ S, 23°38’ E; 18 Jun. 1939; Mevr. Bequaert leg.;

RMCA. – **Kinshasa** • 1 ♀; Bas-Congo: Kalina; 04°18’ S, 15°17’ E; May 1945; Mme. Delsaut leg.; RMCA • 3 ♀♀; same location as for preceding; Jul. 1945; Mme. Delsaut leg.; RMCA • 3 ♂♂; same location as for preceding; Aug. 1945; Mme. Delsaut leg.; RMCA • 2 ♀♀; Congo Belge; 04°19’ S, 15°19’ E; Don Gilson leg.; RMCA • 1 ♂; same location as for preceding; Aug. 1945; Don Gilson leg.; RMCA • 1 ♀; Kinshasa; 04°19’ S, 15°19’ E; A. Tinant leg.; RMCA • 1 ♂; same location as for preceding; A. Tinant leg.; RMCA • 1 ♀; Léopoldville (Kinshasa); 04°19’ S, 15°19’ E; Apr. – May 1911; A. Dubois leg.; RMCA • 2 ♀♀; same location as for preceding; 1933; A. Tinant leg.; RMCA • 1 ♂; same location as for preceding; 1933; A. Tinant leg.; RMCA • 1 ♀; same location as for preceding; 20 Jan. 1912; Dr. A. Dubois leg.; RMCA • 1 ♀; same location as for preceding; Aug. 1949; Dr. A. Dubois leg.; RMCA • 3 ♀♀; same location as for preceding; May – Jun. 1911; Dr. A. Dubois leg.; RMCA • 1 ♀; same location as for preceding; Apr. 1912; Dr. Christy leg.; RMCA • 4 ♀♀; same location as for preceding; Dr. Houssiaux leg.; RMCA • 1 ♀; same location as for preceding; 17 Sep. 1910; Dr. J. Bequaert leg.; RMCA • 1 ♀; same location as for preceding; 1911; Dr. Mouchet leg.; RMCA • 1 ♀; same location as for preceding; 2 Sep. 1911; Dr. Mouchet leg.; RMCA • 1 ♀; same location as for preceding; 7 Jun. 1946; Henrion leg.; RMCA • 1 ♀; same location as for preceding; Jul. 1945; J.M. Berteaux leg.; RMCA • 1 ♂; same location as for preceding; Jul. 1945; J.M. Berteaux leg.; RMCA • 2 ♀♀; same location as for preceding; Aug. 1945; J.M. Berteaux leg.; RMCA • 1 ♂; same location as for preceding; 1957; P. Jobels leg.; RMCA • 2 ♀♀; same location as for preceding; 1942; R. Fiasse leg.; RMCA • 2 ♀♀; same location as for preceding; Jan. – Feb. 1942; R. Fiasse leg.; RMCA • 2 ♀♀; Léopoldville-Kalina; 04°19’ S, 15°19’ E; Apr. 1945; Mme. Delsaut leg.; RMCA • 11 ♀♀; same location as for preceding; Aug. 1945; Mme. Delsaut leg.; RMCA • 8 ♂♂; same location as for preceding; Aug. 1945; Mme. Delsaut leg.; RMCA • 1 ♀; Léo-Stanleyville; 04°19’ S, 15°19’ E; Jan. 1934; Weyms leg.; RMCA • 1 ♀; same location as for preceding; Weyms leg.; RMCA • 1 ♀; Lio-Kalina; 04°19’ S, 15°19’ E; Sep. 1939; J.J. Deheyn leg.; RMCA • 1 ♀; same location as for preceding; Feb. – Mar. 1978; A. Ruwet leg.;

RMCA. – **Kongo Central** • 1 ♀; Banana; 06°00’ S, 12°24’ E; May 1940; A.T. Marré leg.; RMCA • 1 ♀; same location as for preceding; Dr. Etienne leg.; RMCA • 2 ♀♀; same location as for preceding; Aug. 1910; Dr. Etienne leg.; RMCA • 1 ♀; same location as for preceding; 14 Sep. 1913, Dr. J. Bequaert leg.; RMCA • 2 ♀♀; same location as for preceding; Weyms leg.; RMCA • 5 ♀♀; Bangu (Mbanza Ngungu); 05°15’S, 14°30’ E; Aug. 1913; R. Verschueren leg.; RMCA • 1 ♀; Bas-Congo: Banana; 06°00’ S, 12°24’ E; 1951; I. Mesmaekers leg.; RMCA • 1 ♀; Bas-Congo: Boma; 05°51’ S, 13°03’ E; 1950; I. Mesmaekers leg.; RMCA • 1 ♀; same location as for preceding; 1955; R.F. Anselmus leg.; RMCA • 1 ♀; same location as for preceding; Dec. 1955; R.F. Anselmus leg.; RMCA • 1♀; Bas-Congo: Cattier (Lufutoto); 05°26’ S, 14°45’ E; 1946; Delafaille leg.; RMCA • 1 ♀; same location as for preceding; 1946 – 1949; Delafaille leg.; RMCA • 1 ♀; Bas-Congo: Inkisi; 05°08’ S, 15°04’ E; 1951; Mme. Malfeyt leg.; RMCA • 1 ♀; Bas-Congo: Kinkenge; 04°51’ S, 13°37’ E; Feb. – Mar. 1951; Mme. M. Bequaert leg.; RMCA • 1 ♀; Bas-Congo: Kisantu; 05°08’ S, 15°06’ E; May 1945; Rév. Fr. Anastase leg.; RMCA • 1 ♀; Bas-Congo: Kivunda-Luozi; 04°31’ S, 14°14’ E; 19 May 1951; Mme. M. Bequaert leg.; RMCA • 1 ♀; Bas-Congo: Lemfu; (Madimba); 05°18’ S, 15°13’ E; Jan. 1945; Rév. P. De Beir leg.; RMCA • 2 ♂♂; same location as for preceding; Jan. 1945; Rév. P. De Beir leg.; RMCA • 2 ♀♀; same location as for preceding; Apr. 1945; Rév. P. De Beir leg.; RMCA • 2 ♀♀; same location as for preceding; Jun. 1945; Rév. P. De Beir leg.; RMCA • 1 ♀; same location as for preceding; 1931; Rév. P. Van Eyen leg.; RMCA • 1 ♀; Bas-Congo: Lukula; 05°23’ S, 12°57’ E; 1952; Dr. R. Wautier leg.; RMCA • 1 ♀; Bas-Congo: Manzadi; 05°49’ S, 13°28’ E; 1937; Dr. Dartevelle leg.; RMCA • 1 ♀; Bas-Congo: Matadi; 05°49’ S, 13°28’ E; 1959; C. Crespin leg.; RMCA • 1 ♀; Bas-Congo: Mayidi (Madimba); 05°11’ S, 15°09’ E; 1942; Rév. P. Van Eyen leg.; RMCA • 1 ♀; same location as for preceding; 1945; Rév. P. Van Eyen leg.; RMCA • 1 ♀; Bas-Congo: Moanda; 05°56’ S, 12°21’ E; 28 Apr. 1970; P.M. Elsen leg.; RMCA • 1 ♀; Boma; 05°51’ S, 13°03’ E; 15 Nov. 1911; Cambier leg.; RMCA • 1 ♀; same location as for preceding; 16 Jul; 1920; Dr. H. Schouteden leg.; RMCA • 1 ♂; same location as for preceding; 23 Jul. 1920; Dr. H. Schouteden leg.; RMCA •1 ♀; same location as for preceding; 9 Sep. 1920; Dr. H. Schouteden leg.; RMCA • 1 ♂; same location as for preceding; 9 Sep. 1920; Dr. H. Schouteden leg.; RMCA • 4 ♀♀; same location as for preceding; 1937; Dr. Schlesser leg.; RMCA • 8 ♂♂; same location as for preceding; 1937; Dr. Schlesser leg.; RMCA • 1 ♀; same location as for preceding; 1931; H. Derungs leg.; RMCA • 1 ♀; same location as for preceding; 2 Nov. 1945; J. Vrydagh leg.; RMCA • 2 ♀♀; same location as for preceding; 28 Mar. 1913; Lt. Styczynski leg.; RMCA • 1 ♀; same location as for preceding; R.F. Achille leg.; RMCA • 1 ♂; same location as for preceding; R.F. Achille leg.; RMCA • 1 ♀; same location as for preceding; 1935; W. Moreels leg.; RMCA • 2 ♀♀; Boma-Coquilhatville; 05°51’ S, 13°03’ E; Thisquens leg.; RMCA • 1 ♀; Congo da Lemba (Songololo); 05°42’ S, 13°42’ E; Jan. 1913; R. Mayné leg.; RMCA • 1 ♀; same location as for preceding; R. Mayné leg.; RMCA • 2 ♀♀; Kisantu (Madimba); 05°08’ S, 15°06’ E; 26 Dec. 1911; Dr. Mouchet leg.; RMCA • 1 ♀; same location as for preceding; Jun. 1921; P. Gillet leg.; RMCA • 3 ♀♀; same location as for preceding; 1921; P. Van Wing leg.; RMCA • 1 ♂; same location as for preceding; 1921; P. Van Wing leg.; RMCA • 1 ♀; same location as for preceding; 1942; P. Van Wing leg.; RMCA • 1 ♀; same location as for preceding; Jun. 1921; P. Van Wing leg.; RMCA • 7 ♂♂; same location as for preceding; Jun. 1921; P. Van Wing leg.; RMCA • 5 ♀♀; same location as for preceding; 1927; R. P. Vanderyst leg.; RMCA • 7 ♂♂; same location as for preceding; 1927; R.P. Vanderyst leg.; RMCA • 2 ♀♀; same location as for preceding; 1931; R. P. Vanderyst leg.; RMCA • 2 ♂♂; same location as for preceding; 1931; R. P. Vanderyst leg.; RMCA • 1 ♂; same location as for preceding; Jan. – Apr. 1930; R. P. Vanderyst leg.; RMCA • 2 ♀♀; same location as for preceding; Rév. P. Regnier leg.; RMCA • 1 ♀; Kwango: Kiniati-Yasa; 05°20’, 12°56’ E; 18 Jan. 1952; R.P. J. Ruelle leg.; RMCA • 1 ♀; same location as for preceding; Sep. – Oct. 1952; R.P. J. Ruelle leg.; RMCA • 1 ♀; same location as for preceding; Oct. – Nov. 1952; R.P. J. Ruelle leg.; RMCA • 1 ♀; Lukunga (Bas-congo); 05°30’ S, 14°30’ E; 20 Feb. 1968; P.M. Elsen leg.; RMCA • 1 ♀; Matadi; 05°49’ S, 13°28’ E; 1959; Dr. Ch. Wautier leg.; RMCA • 1 ♀; same location as for preceding; Mar. 1937; Dr. Dartevelle leg.; RMCA • 1 ♀; same location as for preceding; Feb. – Mar. 1937; Dr. Dartevelle leg.; RMCA • 16 ♀♀; Mayidi (Madimba); 05°11’ S, 15°09’ E; 1942; Rév. P. Van Eyen leg.; RMCA • 2 ♀♀; same location as for preceding; 1945; Rév. P. Van Eyen leg.; RMCA • 2 ♀♀; Mayumbe; 05°08’ S, 12°29’ E; Cabra leg.; RMCA 2 ♀♀; same location as for preceding; Deleval leg.; RMCA • 1 ♀; Mayumbe: Ganda-Sundi; 05°30’ S, 12°53’ E; Cabra leg.; RMCA • 1 ♀; same location as for preceding; 1915; R. Mayné leg.; RMCA • 1 ♀; Mayumbe: Seke Banza; 05°18’ S, 13°16’ E; 13. Apr. 1924; A. Collart leg.; RMCA • 1 ♀; Mayumbe: Tshela; 04°59’ S, 12°56’ E; 11. May 1924; A. Collart leg.; RMCA • 4 ♂♂; Thysville (Mbanza Ngungu); 05°15’ S, 14°52’ E; Jan. 1953; J. Sion leg.; RMCA • 1 ♀; same location as for preceding; Dec. 1952; N. Leleup leg.; RMCA • 1 ♀; Tshikay (Banana); 06°00’ S, 12°24’ E; Jun. 1949; A.T. Marré leg.;

RMCA. – **Kwilu** • 1 ♀; Kolo-Kwilu-Madiata; 05°27’ S, 14°52’ E; Sep. 1913; R. Verschueren leg.; RMCA • 10 ♂♂; same location as for preceding; Sep. 1913; R. Verschueren leg.; RMCA • 1 ♀; Leverville (Bulungu); 04°50’ S, 18°44’ E; 1928; Mme. J. Tinant leg.; RMCA • 1 ♀; Moyen Kwilu: Leverville (Bulungu); 04°50’ S, 18°44’ E; P. Vanderijst leg.; RMCA • 1 ♀; Mwilambongo (Idiofa); 04°56’ S, 19°48’ E; 1947; Rév. Sœur Imelda leg.;

RMCA. – **Lomami** • 1 ♀; Gadanjika; 06°45’ S, 23°07’ E; 12 Feb. 1948; P. de Francquen leg.; RMCA • 3 ♀♀; same location as for preceding; 18 Dec. 1950; P. de Francquen leg.; RMCA • 1 ♂; same location as for preceding; 18 Dec. 1950; P. de Francquen leg.; RMCA • 2 ♀♀; same location as for preceding; 22 Nov. 1950; P. de Francquen leg.; RMCA • 4 ♀♀; same location as for preceding; 24 Nov. 1950; P. de Francquen leg.; RMCA • 1 ♀; Kabinda; 06°08’ S, 24°29’ E; Dr. Schwetz leg.; RMCA • 1 ♂; Kanda Kanda; 06°56’ S, 23°37’ E; 10 Dec. 1925; Ch. Seydel leg.; RMCA • 1 ♀; same location as for preceding; 10 Dec. 1926; Ch. Seydel leg.; RMCA • 1 ♀; Lomami: Tshofa; 05°14’ S, 25°15’ E; Mar. 1939; Mevr. Bequaert leg.; RMCA • 1 ♀; same location as for preceding; Aug. 1939; Mevr. Bequaert leg.; RMCA • 1 ♀; Sankuru: Gandajika; 06°45’ S, 23°57’ E; 1953; R. Verschueren leg.; RMCA • 1 ♀; same location as for preceding;12 Feb. 1948; R. Verschueren leg.; RMCA • 1 ♀; Sankuru: M’Pemba Zeo (Gandajika); 06°49’ S, 23°58’ E; 11. Jun. 1960; R. Mouchamps leg.; RMCA • 1 ♂; same location as for preceding;19 Dec. 1959; R. Mouchamps leg.; RMCA • 1 ♀; same location as for preceding; 6 Sep. 1959; R. Mouchamps leg.; RMCA • 1 ♀; Station de Gandajika INEAC; 06°49’ S, 23°58’ E; 1957; P. de Francquen leg.; RMCA • 1 ♀; Terr. Kanda Kanda: Gandajika; 06°56’ S, 23°37’ E; 1947; P. Henrard leg.;

RMCA. – **Lualaba** • 4 ♂♂; Dilolo; 10°41’ S, 22°21’ E; Aug. 1931; G.F. de Witte leg.; RMCA • 1 ♀; Ditanto; 10°15’ S, 25°53’ E; Oct. 1925; Ch. Seydel leg.; RMCA • 1 ♂; Haut-Luapula: Kansenia; 10°19’ S, 26°04’ E; 14 Oct. 1929; R. Verschueren leg.; RMCA • 1 ♂; same location as for preceding; 4 Oct. 1929; R. Verschueren leg.; RMCA • 1 ♀; Kafakumba (Sandoa); 09°41’ S, 23°44’ E; Apr. 1933; F.G. Overlaet leg.; RMCA • 1 ♂; same location as for preceding; Apr. 1933; F.G. Overlaet leg.; RMCA • 1 ♀; Kapanga; 08°21’ S, 22°34’ E; Mar. 1933; F.G. Overlaet leg.; RMCA • 1 ♀; same location as for preceding; Nov. 1932; F.G. Overlaet leg.; RMCA • 1 ♀; same location as for preceding; Dec. 1932; F.G. Overlaet leg.; RMCA • 1 ♂; Katanga: Ditanto; 10°15’ S, 25°53’ E; Oct. 1925; Ch. Seydel leg.; RMCA • 1 ♀; Katanga: Lubudi; 09°55’ S, 25°58’ E; Aug. 1945; R. Close leg.; RMCA • 1 ♂; Katanga: Musonoie; 10°42’ S, 25°23’ E; Jul. 1924; Ch. Seydel leg.; RMCA • 1 ♀; Katanga: Kolwezi; 10°44’ S, 25°28’ E; 10 Apr. 1953; Mme. Gilbert leg.; RMCA • 1 ♀; Lulua: Kapanga; 08°21’ S, 22°34’ E; Mar. 1933; Mme. Gilbert leg.; RMCA • 1 ♂; same location as for preceding; Mar. 1933; Mme. Gilbert leg.; RMCA • 1 ♀; same location as for preceding; Oct. 1932; Mme. Gilbert leg.; RMCA • 3 ♂♂; same location as for preceding; Oct. 1932; Mme. Gilbert leg.;

RMCA. – **Maï-Ndombe** • 1 ♀; Kwamouth (Mushie); 03°11’ S, 16°12’ E; 5 Dec. 1911; Dr. A. Dubois leg.; RMCA • 1 ♀; Lac Léopold II: Bolobo (Lac Maï-Ndombe); 02°00’ S, 18°15’ E; 1956; R.J.D. Viccars leg.; RMCA • 1 ♀; Wombali; 03°16’ S, 17°22’ E; Jul. 1913; P. Vanderijst leg.; RMCA. – **Maniema** • 1 ♂; K. 240 de Kindu; 02°57’ S, 25°55’ E; 17 Sep. 1911; L. Burgeon leg.; RMCA • 1 ♂; K. 300 de Kindu; 02°57’ S, 25°55’ E; 6 May 1911; L. Burgeon leg.; RMCA • 1 ♀; Kasongo: Kikwe; 04°27’ S, 26°40’ E; 29 Oct. 1959; P.L.G. Benoit leg.; RMCA • 1 ♂; Kibombo; 03°54’ S, 25°55’ E; Sep. – Nov. 1930; H.J. Brédo leg.; RMCA • 1 ♀; same location as for preceding; May 1930; H.J. Brédo leg.; RMCA • 1 ♀; same location as for preceding; Oct. 1930; H.J. Brédo leg.; RMCA • 1 ♀; Kindu; 02°57’ S, 25°55’ E; Dr. Russo leg.; RMCA • 1 ♀; same location as for preceding; L. Burgeon leg.; RMCA • 1 ♂; Malela; 04°22’ S, 26°08’ E; Jan. 1914; L. Burgeon leg.; RMCA • 1 ♀; Nyangwe; 04°13’ S, 26°11’ E; 19 Nov. 1910; Dr. J. Bequaert leg.; RMCA • 1 ♀; same location as for preceding; Mar. – Apr. 1918; R. Mayné leg.; RMCA • 2 ♀♀; same location as for preceding; Apr. – May 1918; R. Mayné leg.; RMCA • 1 ♀; same location as for preceding; R. Mayné leg.; RMCA • 1 ♀; Vieux Kassongo (Kasongo); 04°27’ S, 26°40’ E; 1910; Dr. J. Bequaert leg.; RMCA • 1 ♀; same location as for preceding; 1 Dec. 1910; Dr. J. Bequaert leg.; RMCA. – **Mongala** • 1 ♀; Bangala: Binga; 02°28’ N, 30°31’ E; 1933; Vandenput leg.; RMCA • 1 ♂; Lisala; 02°09’ N, 21°30’ E; 1930; Mme. Babillon leg.;

RMCA. – **Nord-Kivu** • 1 ♀; Beni; 00°29’ N, 29°28’ E; 18 Aug. 1932; L. Burgeon leg.; RMCA • 1 ♀; same location as for preceding; Lt. Borgerhoff leg.; RMCA • 1 ♀; same location as for preceding; May 1911; Lt. Borgerhoff leg.; RMCA • 1 ♀; Beni; 00°29’ N, 29°28’ E, alt. 1120 m; Feb. 1931; Mme. L. Lebrun leg.; RMCA • 1 ♀; Beni; 00°29’ N, 29°28’ E, alt. 1150 m; 7 Nov. 1931; Mme. L. Lebrun leg.; RMCA • 2 ♀♀; Beni à Lesse; 00°45’ N, 29°46’ E; Jul. 1911; Dr. Murtula leg.; RMCA • 1 ♀; de Beni à Semliki; 00°29’ N, 29°28’ E; 12 Nov. 1931; Mme. L. Lebrun leg.; RMCA • 1 ♀; Ituri: Beni; 00°29’ N, 29°28’ E; Aug. 1914; Dr. J. Bequaert leg.; RMCA • 1 ♀; Ituri: Butembo; 00°09’ N, 29°17’ E; Jul. – Aug. 1914; Mme. Van Riel leg.; RMCA • 1 ♀; Kalonge, Riv Mushuva; 00°21’ N, 29°49’ E; 6 Jun. 1949; G. Marlier leg.; RMCA • 1 ♀; Kivu: Buseregenye (Rutshuru); 01°11’ S, 29°27’ E; Sep. 1929; Ed. Luja leg.; RMCA • 1 ♀; Kivu: Kissengni; 01°41’ S, 29°14’ E; Jan. 1923; R. Van Saceghem leg.; RMCA • 1 ♀; same location as for preceding; Mar. – Apr. 1923; R. Van Saceghem leg.; RMCA • 1 ♀; Kivu: Masisi; 01°24’ S, 28°49’ E, alt. 1200 m; 1951; Dedobeleere leg.; RMCA • 1 ♀; Kivu: Mulo; 00°10’ S, 29°14’ E, alt. 1500 m; Apr. 1950; R.P. Celis leg.; RMCA • 1 ♀; Kivu: Nyabikoro; 01°11’ S, 29°27’ E; Feb. 1957; K. Baeten leg.; RMCA • 1 ♀; Kivu: Nyabikoro (Rutshuru); 01°11’ S, 29°27’ E; Nov. 1956; K. Baeten leg.; RMCA • 1 ♀; N. Kivu: Kisenyi; 01°41’ S, 29°14’ E; Feb. 1927; Ch. Seydel leg.; RMCA • 1 ♂; same location as for preceding; Feb. 1928; Ch. Seydel leg.; RMCA • 1 ♀; N. Kivu: Lulenga (Rutshuru); 01°24’ S, 29°22’ E; Dec. 1927; Ch. Seydel leg.; RMCA • 1 ♂; N.Kivu: Kissenyi; 01°41’ S, 29°14’ E; Feb. 1928; Ch. Seydel leg.; RMCA • 1 ♀; PNA [Parc National Albert]; 00°03’ S, 29°30’ E; 15 Dec. 1957; P. Vanschuytbroeck & H. Synave leg.; RMCA • 1 ♀; same location as for preceding; 23 Oct. 1956; P. Vanschuytbroeck & H. Synave leg.; RMCA • 1 ♀; same location as for preceding; 3 Apr. – 16. Aug. 1957; P. Vanschuytbroeck & H. Synave leg.; RMCA • 1♀; same location as for preceding; 7 Jun. 1957; P. Vanschuytbroeck & H. Synave leg.; RMCA • 1 ♀; same location as for preceding; 7 Nov. 1956; P. Vanschuytbroeck & H. Synave leg.; RMCA • 1 ♀; same location as for preceding; 7 – 15 Jul. 1955; P. Vanschuytbroeck & H. Synave leg.; RMCA • 1 ♀; PNA Lusilube; 00°03’ S, 29°30’ E; 15 Aug. 1946; J. de Wilde leg.; RMCA • 1 ♀; Rutshuru; 01°11’ S, 29°27’ E; Jan. 1934; Dr. De Wulf leg.; RMCA • 1 ♀; same location as for preceding; Mar. 1938; J. Ghesquière leg.; RMCA • 1 ♀; same location as for preceding; Oct. 1937; J. Ghesquière leg.; RMCA • 1 ♀; same location as for preceding; Nov. 1937; J. Ghesquière leg.;

RMCA. – **Nord-Ubangi** • 1 ♀; Abumombazi (Mobayi); 03°34’ N, 22°03’ E; 18 – 26 Feb. 1932; R. Verschueren leg.; RMCA • 1 ♂; same location as for preceding; 18 – 26 Feb. 1932; R. Verschueren leg.; RMCA • 1 ♀; Bosobolo; 04°11’ N, 19°53’ E; Dr. Girling leg.; RMCA • 1 ♀; Ubangi: Bosobolo; 04°11’ N, 19°53’ E; 5 – 11 Jan. 1932; H.J. Brédo leg.; RMCA • 7 ♀♀; same location as for preceding; 8 – 11 Jan. 1932; H.J. Brédo leg.; RMCA • 2 ♂♂; same location as for preceding; 8 – 11 Jan. 1932; H.J. Brédo leg.; RMCA • 1 ♀; Ubangi: Karawa (Businga); 03°21’ N, 20°18’ E; 1935; Rév. Wallin leg.; RMCA • 4 ♀♀; same location as for preceding; 1936; Rév. Wallin leg.; RMCA • 1 ♀; same location as for preceding; 1937; Rév. Wallin leg.; RMCA • 1 ♀; Ubangi: Yakoma; 04°06’ N, 22°23’ E; 15 Feb. 1931; H.J. Brédo leg.; RMCA • 1 ♀; same location as for preceding; Jun. 1932; H.J. Brédo leg.; RMCA • 12 ♀♀; Yakoma; 04°06’ N, 22°23’ E; 5 – 17 Feb. 1932; H.J. Brédo leg.; RMCA • 8 ♂♂; same location as for preceding; 5 – 17 Feb. 1932; H.J. Brédo leg.; RMCA. – **Sankuru** • 1 ♀; Sankuru; 04°58’ S, 23°26’ E; 1910; Dr. Abrassart leg.; RMCA • 1 ♀; Sankuru: Komi; 03°23’ S, 23°46’ E; Dec. 1918; J. Ghesquière leg.; RMCA • 1 ♂; Sankuru: Tshumbe S. Marie; 04°02’ S, 22°41’ E; May 1948; Mevr. Bequaert leg.; RMCA. – **Sud-Kivu** • 1 ♀; Kalembelembe-Baraka; 04°32’ S, 28°47’ E; Jul. 1918; R. Mayné leg.; RMCA • 1 ♀; Kivu: Bobandana; 01°42’ S, 29°01’ E; 1928; O. Douce leg.; RMCA • 1 ♀; Kivu: Bukavu; 02°29’ S, 28°51’ E; Dec. 1953; H. Bomans leg.; RMCA • 1 ♀; Kivu: Katana (Kabare); 02°14’ S, 28°49’ E; Oct. 1932; P Van der Houd leg.; RMCA • 1 ♀; Kivu: Luvungi; 02°52’ S, 29°02’ E; Aug. 1927; Ch. Seydel leg.; RMCA • 1♂; Kivu: Mulungu; 02°19’ S, 28°45’ E; 1949; F.L. Hendrickx leg.; RMCA • 1 ♀; same location as for preceding; 15 Apr. 1938; F.L. Hendrickx leg.; RMCA • 3 ♀♀; same location as for preceding; 2 Aug. 1938; F.L. Hendrickx leg.; RMCA • 2 ♀♀; same location as for preceding; 29 Sep. 1938; F.L. Hendrickx leg.; RMCA • 2 ♀♀; same location as for preceding; Aug. 1938; F.L. Hendrickx leg.; RMCA • 1 ♀; same location as for preceding; 19 Nov. 1932; L. Burgeon leg.; RMCA • 1 ♀; same location as for preceding; 19 – 26 Nov. 1932; L. Burgeon leg.; RMCA • 1 ♀; Kivu: Uvira; 03°25’ S, 29°08’ E; 16 – 23 Mar. 1953; P. Basilewsky leg.; RMCA • 1 ♂; same location as for preceding; 16 – 23 Mar. 1953; P. Basilewsky leg.; RMCA • 1 ♀; same location as for preceding; 24 – 28 Dec. 1952; P. Basilewsky leg.; RMCA • 1 ♀; Mulungu: Tshibinda; 02°19’ S, 28°45’ E; Nov. 1956; P.C. Lefèvre leg.; RMCA • 1 ♀; N. Lac Kivu: Rwankwi; 02°19’ S, 28°45’ E; Apr. 1948; J.V. Leroy leg.; RMCA • 4 ♀♀; Région des Lacs; 02°29’ S, 28°51’ E; Dr. Sagona leg.; RMCA • 3 ♀♀; Uvira; 03°25’ S, 29°08’ E; Nov. 1927; Ch. Seydel leg.; RMCA • 1 ♀; W. Kivu: Ibanda; 02°29’ S, 28°51’ E; 1937; M. Vandelannoite leg.; RMCA • 1 ♀; same location as for preceding; 1935; M. Vandelannoite leg.;

RMCA. – **Sud-Ubangi** • 1 ♀; Kunungu; 02°45’ N, 18°26’ E; 1932; Dr. H. Schouteden leg.; RMCA • 1 ♂; Libenge; 03°39’ N, 18°38’ E; 10 Dec. 1931; H.J. Brédo leg.; RMCA • 1 ♂; same location as for preceding; Dec. 1931; H.J. Brédo leg.; RMCA • 1 ♀; same location as for preceding; 1931; J. Van Gils leg.; RMCA • 1 ♂; same location as for preceding; 1931; J. Van Gils leg.; RMCA • 3 ♀♀; Ubangi: Boma-Motenge; 03°15’ N, 18°39’ E; Dec. 1931; H.J. Brédo leg.; RMCA • 1 ♂; same location as for preceding; Dec. 1931; H.J. Brédo leg.; RMCA • 1 ♀; Ubangi: Duma (Libenge); 03°53’ N, 18°41’ E; 1930; Cap. Van Gils leg.; RMCA • 1 ♀; Ubangi: Libenghi; 03°39’ N, 18°38’ E; Vreurick leg.; RMCA • 5 ♀♀; Ubangi: Motenge-Boma; 03°15’ N, 18°39’ E; 14 Dec. 1931; H.J. Brédo leg.; RMCA • 1 ♀; same location as for preceding; 2 Dec. 1931; H.J. Brédo leg.; RMCA • 2 ♀♀; same location as for preceding; 22 Dec. 1931; H.J. Brédo leg.; RMCA • 1 ♂; same location as for preceding; 22 Dec. 1931; H.J. Brédo leg.; RMCA • 1 ♂; same location as for preceding; Dec. 1931; H.J. Brédo leg.; RMCA. – **Tanganyika** • 1 ♀; Albertville (Kalemie); 05°57’ S, 29°12’ E; Jul. 1954; R. Verschueren leg.; RMCA • 1 ♀; Baudouinville (Moba); 07°02’ S, 29°47’ E; Dec. 1958; H. Bomans leg.; RMCA • 1 ♂; Buli (Kabalo); 05°50’ S, 26°55’ E; 18 Feb. 1911; Dr. J. Bequaert leg.; RMCA • 1 ♀; Katanga: Katompe (Kabalo); 06°11’ S, 26°20’ E; Feb. 1929; Ch. Seydel leg.; RMCA • 4 ♀♀; Katanga: Lukulu (Manono); 07°08’ S, 28°06’ E; Feb. 1929; Mr. Remacle leg.; RMCA • 1 ♀; Kiambi (Manono); 07°19’ S, 28°01’ E; 10 Mar. 1911; Dr. Valdonio leg.; RMCA • 1 ♂; same location as for preceding; 27 Apr. 1931; G.F. de Witte leg.; RMCA • 1 ♀; same location as for preceding; 5 – 15 May 1931; G.F. de Witte leg.; RMCA • 1 ♀; same location as for preceding; May 1931; G.F. de Witte leg.; RMCA • 1 ♀; Lusaka (Baudouinville); 07°02’ S, 29°47’ E; 1937; R.P. Debbaudt leg.; RMCA • 1 ♀; Tanganyika: Moba; 07°02’ S, 29°47’ E; Mar. 1954; H. Bomans leg.; RMCA • 1 ♀; Tanganyika: Moba; 07°02’ S, 29°47’ E, alt. 780 m; Jul. – Aug. 1953; H. Bomans leg.; RMCA • 1 ♂; same location as for preceding; Jul. – Aug. 1953; H. Bomans leg.; RMCA • 1 ♀; Tanganyika: Mpala; 06°45’ S, 29°31’ E, alt. 780 m; Jun. 1953; H. Bomans leg.; RMCA • 1 ♂; same location as for preceding; Jul. – Aug. 1953; H. Bomans leg.; RMCA • 1 ♀; Tanganyika-Moero: Nyunzu; 05°57’ S, 28°01’ E; Jan. – Feb. 1954; H. De Saeger leg.; RMCA • 1 ♀; Vallée Lukuga; 05°40’ S, 26°55’ E; Nov. 1911; Dr. Schwetz leg.; RMCA • 1 ♂; same location as for preceding; Nov. 1911; Dr. Schwetz leg.;

RMCA. – **Tshopo** • 2 ♀♀; Basoko; 01°13’ N, 23°36’ E; Jan. 1948; P.L.G. Benoit leg.; RMCA • 2 ♀♀;; same location as for preceding; Feb. 1948; P.L.G. Benoit leg.; RMCA • 1 ♀; Haut-Uele: Paulis; 02°46’ N, 27°38’ E; Oct. 1947; P.L.G. Benoit leg.; RMCA • 1 ♀; Miss. St. Gabriel; 00°33’ N, 25°05’ E; M. Torley leg.; RMCA • 1 ♀; Stanleyville (Kisangani); 00°31’ N, 25°11’ E; 1936; A. Becquet leg.; RMCA • 2 ♀♀; Terr. de Basoko (forêt); 01°13’ N, 23°36’ E; 1950; R.P. Camps leg.; RMCA • 1 ♀; Yangambi; 00°46’ N, 24°27’ E; Nov. 1936; J. Ghesquière leg.; RMCA • 1 ♂; same location as for preceding; Oct. 1937; P. Henrard leg.; RMCA. – **Tshuapa** • 1 ♀; Bamania (Mbandaka); 00°01’ N, 18°19’ E; 10 Mar; 1936; R. Verschueren leg.; RMCA • 1 ♀; same location as for preceding; Sep. – Nov. 1936; P. Henrard leg.; RMCA • 2 ♀♀; Boende; 00°13’ S, 20°52’ E; Jan. 1952; Rév. P. Lootens leg.; RMCA • 1 ♂; same location as for preceding; Jan. 1952; Rév. P. Lootens leg.; RMCA • 1 ♀; Bokuma; 00°06’ S, 18°41’ E; Feb. – Apr. 1934; R. Verschueren leg.; RMCA • 1 ♀; Equateur: Bamania (Mbandaka); 00°01’ N, 18°19’ E; May 1958; Rév. P. Hulstaert leg.; RMCA • 1 ♀; Equateur: Bokuma; 00°06’ S, 18°41’ E; Jul. 1952; H.J. Brédo leg.; RMCA • 1 ♀; same location as for preceding; 1951; Rév. P. Lootens leg.; RMCA • 1 ♂; same location as for preceding; 4 Jul. 1934; Rév. P. Lootens leg.; RMCA • 2 ♀♀; same location as for preceding; Feb. 1952; Rév. P. Lootens leg.; RMCA • 1 ♀; same location as for preceding; Mar. 1952; Rév. P. Lootens leg.; RMCA • 1 ♀; same location as for preceding; Jul. 1952; Rév. P. Lootens leg.; RMCA • 1 ♀; same location as for preceding; Dec. 1951; Rév. P. Lootens leg.; RMCA • 1 ♀; Equateur: Flandria (Ingende); 00°20’ S, 19°06’ E; Jan. – Feb. 1928; Rév. P. Lootens leg.; RMCA • 1♀; Riv. Busira; 00°06’ S, 19°55’ E; Jun. 1936; J. Ghesquière leg.; RMCA • 1 ♀; Tshuapa: Bamania (Mbandaka); 00°01’ N, 18°19’ E; Sep. 1964; Rév. P. Hulstaert leg.; RMCA • 1 ♀; same location as for preceding; Mar. 1954; Rév. P. Lootens leg.; RMCA • 1 ♀; same location as for preceding; Dec. 1954; Rév. P. Lootens leg.; RMCA • 1 ♀; Tshuapa: Bamanya; 00°01’ N, 18°19’ E; 1960; Rév. P. Hulstaert leg.; RMCA • 1 ♀; same location as for preceding; 1960; Rév. P. Hulstaert leg.; RMCA • 4 ♀♀; same location as for preceding; 1968; Rév. P. Hulstaert leg.; RMCA • 1 ♀; same location as for preceding; Feb. 1962; Rév. P. Hulstaert leg.; RMCA • 1 ♀; Tshuapa: Bokuma; 00°06’ S, 18°41’ E; 1954; Rév. P. Lootens leg.; RMCA • 1 ♀; same location as for preceding; Feb. 1954; Rév. P. Lootens leg.; RMCA • 1 ♀; same location as for preceding; Mar. 1952; Rév. P. Lootens leg.; RMCA • 1 ♂; same location as for preceding; Mar. 1952; Rév. P. Lootens leg.; RMCA • 1 ♀; same location as for preceding; Apr. 1952; Rév. P. Lootens leg.; RMCA • 2♀♀; Tshuapa: Bokungu; 00°41’ S, 22°19’ E; 1949; M. Dupuis leg.; RMCA • 2 ♀♀; Tshuapa: Flandria (Ingende); 00°20’ S, 19°06’ E; Sep. 1946 – Aug. 1947; Rév. P. Lootens leg.; RMCA • 1 ♀; same location as for preceding; Sep. – Dec. 1947; Rév. P. Lootens leg.; RMCA.

#### *Gronoceras felinum* (Gerstaecker, 1857), *Chalicodoma* (*Gronoceras*) *felina* (Gerstaecker, 1857)

**Material.** Cupboard 110, Box 44 (94♀♀ and 21♂♂). **D.R. CONGO.** – **Equateur** • 1 ♀; Basankusu; 01°13’ N, 19°49; 31 Mar. 1911; Dr. Valdonio leg.; RMCA. – **Haut-Katanga** • 1 ♀; Elisabethville (Lubumbashi); 11°40’ S, 27°28’ E; 16 Jan. 1925; Ch. Seydel leg.; RMCA • 1 ♀; same location as for preceding; 7 Aug. 1921; Eg. Devroye leg.; RMCA • 1 ♀; same location as for preceding; 8 Apr. 1928; Dr. M. Bequaert leg.; RMCA • 4 ♀♀; same location as for preceding; Feb. 1935; P. Quarré leg.; RMCA • 1 ♀; same location as for preceding; Mar. 1913; Dr. J. Bequaert leg.; RMCA • 12 ♀♀; same location as for preceding; Mar. 1935; P. Quarré leg.; RMCA • 1 ♀; same location as for preceding; Apr. 1935; De Loose leg.; RMCA • 1 ♀; same location as for preceding; Sep. 1934; P. Quarré leg.; RMCA • 2 ♀♀; same location as for preceding; De Loose leg.; RMCA • 2 ♂♂; same location as for preceding; May 1926; Ch. Seydel leg.; RMCA RMCA • 1♀; same location as for preceding; May 1934; P. Quarré leg.; RMCA • 1 ♀; same location as for preceding; May 1935; P. Quarré leg.; RMCA • 1 ♀; same location as for preceding; Jun. 1926; Dr. M. Bequaert leg.; RMCA • 2 ♂♂; same location as for preceding; Jun. 1926; Dr. M. Bequaert leg.; RMCA • 6 ♀♀; same location as for preceding; Jun. 1935; P. Quarré leg.; RMCA • 3 ♀♀; same location as for preceding; Jul. 1935; P. Quarré leg.; RMCA • 1 ♀; same location as for preceding; Aug. 1935; P. Quarré leg.; RMCA • 10♀♀; same location as for preceding; Oct. 1934; P. Quarré leg.; RMCA • 4 ♀♀; same location as for preceding; Nov. 1934; P. Quarré leg.; RMCA • 2 ♀♀; same location as for preceding; Dec. 1934; P. Quarré leg.; RMCA • 1 ♀; Jadotville (Likasi); 10°59’ S, 26°44’ E; 1948; R. Mouchamps leg.; RMCA • 1 ♀; same location as for preceding; Nov. 1946; R.S.M. Adelaïde leg.; RMCA • 1 ♀; Kalumba-Kilwa; 09°18’ S, 28°25’ E; Aug. 1907; Dr. Sheffield Neave leg.; RMCA • 1 ♂; same location as for preceding; Aug. 1907; Dr. Sheffield Neave leg.; RMCA • 1 ♂; Kambove-Bunkeya; 10°52’ S, 26°38’ E; Oct. 1907; Dr. Sheffield Neave leg.; RMCA • 1 ♀; Kambove; 10°52’ S, 26°38’ E; 27 Jan. 1912; Dr. J. Bequaert leg.; RMCA • 1 ♀; Katanga; 11°07’ S, 27°06’ E; Lemaire leg.; RMCA • 1 ♀; Katanga; 11°07’ S, 27°06’ E; Weym leg.; RMCA • 2 ♂♂; Katumba; 08°32’ S, 25°59’ E; Oct. 1907; Dr. Sheffield Neave leg.; RMCA • 2 ♀♀; Kiamokosa; 11°40’ S, 27°28’ E; Oct. 1907; Dr. Sheffield Neave leg.; RMCA • 1 ♂; same location as for preceding; Oct. 1907; Dr. Sheffield Neave leg.; RMCA • 1 ♀; Kipushi par Sakania; 11°46’ S, 27°15’ E; Jan. 1948; R. Mouchamps leg.; RMCA • 1 ♀; Lualaba: Kapolowe; 11°02’ S, 26°58’ E; 1959; R. P. Th. De Caters leg.; RMCA • 2 ♀♀; Lubumbashi; 11°40’ S, 27°28’ E; 10 Apr. 1921; Dr. M. Bequaert leg.; RMCA • 1 ♀; same location as for preceding; 13 Mar. 1921; Dr. M. Bequaert leg.; RMCA • 1 ♀; same location as for preceding; 15 Jun. 1921; Dr. M. Bequaert leg.; RMCA • 1 ♀; Mbiliwa-Wantu; 11°40’ S, 27°28’ E; Oct. 1907; Dr. Sheffield Neave leg.; RMCA • 1 ♂; same location as for preceding; Oct. 1907; Dr. Sheffield Neave leg.;

RMCA. – **Haut-Lomami** • 1 ♀; Bukama; 09°12’ S, 25°51’ E; 16 Apr. 1911; Dr. J. Bequaert leg.; RMCA • 1 ♀; same location as for preceding; 20 Apr. 1911; Dr. J. Bequaert leg.; RMCA • 1 ♀; Kikondja; 08°11’ S, 26°26’ E; 28 Nov. 1911; Dr. J. Bequaert leg.; RMCA • 2 ♀♀; Kulu-Mwanza; 07°54’ S, 26°45’ E; May 1927; A. Bayet leg.; RMCA • 1 ♂; PNU Bukena près de Mulongo; 07°42’ S, 27°10’ E; Jun. 1949; Mis. G.F. de Witte leg.; RMCA • 1 ♂; PNU Lufira au pied du Mt Sombwe; 10°10’ S, 27°20’ E; 13-15 Jul. 1949; Mis. G.F. de Witte leg.; RMCA • 2 ♀♀; Sankisia; 09°24’ S, 25°48’ E; 13 Sep. 1911; Dr. J. Bequaert leg.; RMCA. – **Haut-Uele** • 1 ♀; PNG Napokomweli; 03°40’ N, 29°00’ E; Oct. 1931; G. Demoulin leg.; RMCA. – **Ituri** • 1 ♀; Ituri: Mahagi; 02°18’ N, 30°59’ E; 1931; Ch. Scops leg.; RMCA • 2 ♀♀; Kibali-Ituri: Kilo; 01°50’ N, 30°09’ E; 1930; G. du Soleil leg.; RMCA • 1 ♀; Kilo (Djugu); 01°50’ S, 30°09’ E; Dec. 1934; G. du Soleil leg.; RMCA. – **Kongo Central** • 1 ♂; Banana; 06°00’ S, 12°24’ E; Sep. 1915; Dr. J. Bequaert leg.; RMCA • 1 ♀; same location as for preceding; Mar. 1951; I. Mesmaekers leg.; RMCA. – **Lualaba** • 2 ♂♂; Biano; 10°20’ S, 27°00’ E; Oct. 1931; Ch. Seydel leg.; RMCA • 1 ♀; Dilolo; 10°41’ S, 22°21’ E; Aug. 1931; G.F. de Witte leg.; RMCA • 1 ♀; Kayambo-Dikuluwe; 10°45’ S, 25°22’ E; Jul. 1907; Dr. Sheffield Neave leg.; RMCA • 1 ♀; La Mulonga; 09°56’ S, 25°53’ E; Aug. 1931; Ch. Seydel leg.; RMCA • 1 ♂; Manika (Kolwezi); 10°20’ S, 27°00’ E; Oct. 1931; Ch. Seydel leg.; RMCA. – **Nord-Kivu** • 1 ♀; PNA [Parc National Albert]; 00°03’ S, 29°30’ E; 20 Dec. 1956; P. Vanschuytbroeck leg.;

RMCA. – **Tanganyika** • 1 ♂; Kanikiri; 00°09’ S, 29°32’ E; Oct. 1907; Dr. Sheffield Neave leg.; RMCA • 1 ♀; Albertville (Kalemie); 05°57’ S, 29°12’ E; May – Jun. 1954; H. Bomans leg.; RMCA • 1 ♀; Kiambi (Manono); 07°19’ S, 28°01’ E; 23 Apr. 1931; G.F. de Witte leg.; RMCA • 2 ♀♀; same location as for preceding; 27 Apr. 1931; G.F. de Witte leg.; RMCA • 2 ♂♂; same location as for preceding; 27 Apr. 1931; G.F. de Witte leg.; RMCA • 1 ♀; same location as for preceding; 5 – 15 May 1931; G.F. de Witte leg.; RMCA • 1 ♂; Tanganyika: Kabalo; 06°03’ S, 26°55’ E; 1 Jul. 1947; H. Bomans leg.; RMCA • 1 ♀; Tanganyika: Mpala; 06°45’ S, 29°31’ E, alt. 780 m; Jun. 1953; H. Bomans leg.; RMCA • 1 ♂; same location as for preceding; Jun. 1953; H. Bomans leg.; RMCA • 1 ♀; Vallée Lukuga; 05°40’ S, 26°55’ E; Nov. 1911; Dr. Schwetz leg.; RMCA.

#### *Gronoceras mutala* (Strand, 1912), *Chalicodoma* (*Gronoceras*) *mutala* (Strand, 1912)

**Material.** Cupboard 110, Box 44 (9♀♀ and 7♂♂). **D.R. CONGO**. – **Equateur** • 1 ♂; Eala (Mbandaka); 00°03’ N, 18°19’ E; 1933; A. Corbisier leg.; RMCA • 3 ♀♀; same location as for preceding; Jun. 1932; A. Corbisier leg.; RMCA • 4 ♂♂; same location as for preceding; Jun. 1932; A. Corbisier leg.; RMCA • 1 ♀; same location as for preceding; 15 – 30 Oct. 1929; H.J. Brédo leg.; RMCA • 1 ♀; same location as for preceding; 5 Nov. 1931; H.J. Brédo leg.; RMCA • 1 ♀; same location as for preceding; Mar. 1932; H.J. Brédo leg.; RMCA • 1 ♂; same location as for preceding; Mar. 1932; H.J. Brédo leg.; RMCA • 1 ♀; same location as for preceding; May 1932; H.J. Brédo leg.; RMCA. – **Nord-Ubangi** • 1 ♀; Yakoma; 04°06’ N, 22°23’ E; 5 – 17 Feb.1932; H.J. Brédo leg.; RMCA • 1 ♂; same location as for preceding; 5 – 17 Feb.1932; H.J. Brédo leg.; RMCA. – **Tshuapa** • 1 ♀; Bamania (Mbandaka); 00°01’ N, 18°19’ E; 22 Jul. 1924; Dr. M. Bequaert leg.; RMCA.

#### *Gronoceras nigrocaudatum* (Friese, 1903), *Chalicodoma* (*Gronoceras*) *nigrocaudata* (Friese, 1903)

**Material.** Cupboard 110, Box 44 (3♀♀ and 2♂♂). **D.R. CONGO**. – **Bas-Uele** • 1 ♂; Uele: Tukpwo; 04°26’ N, 25°51’ E; Aug. 1937; J. Vrydagh leg.; RMCA. – **Ituri** • 1 ♀; Lac Albert: Kasenyi; 01°24’ N, 30°26’ E; May 1935; H.J. Brédo leg.; RMCA • 1 ♀; Mahagi-Niarembe; 02°15’ N, 31°07’ E; 1935; Ch. Scops leg.; RMCA • 1 ♂; same location as for preceding; 1935; Ch. Scops leg.; RMCA. – **Nord-Kivu** • 1 ♀; Beni (1930m); 00°29’ N, 29°28’ E; Mar. 1931; Mme. L. Lebrun leg.; RMCA.

#### *Gronoceras praetextum* (Vachal, 1910), *Chalicodoma* (*Gronoceras*) *praetexta* (Vachal, 1910)

**Material.** Cupboard 110, Box 34 (9♀♀ and 23♂♂). **D.R. CONGO**. Holotype – **Haut-Katanga** • 1 ♂; Katumba; 08°32’ S, 25°59’ E; Oct. 1907; Dr. Sheffield Neave leg.; RMCA. Paratype – **Haut-Katanga** • 1♂; Kiamokosa; 11°40’ S, 27°28’ E; Oct. 1907; Dr. Sheffield Neave leg.; RMCA.

– **Haut-Katanga** • 1 ♂; Elisabethville (Lubumbashi); 11°40’ S, 27°28’ E; 12 Nov. 1933; Dr. M. Bequaert leg.; RMCA • 1 ♂; same location as for preceding; 14 Apr. 1912; Dr. J. Bequaert leg.; RMCA • 1 ♂; same location as for preceding; 2 Oct. 1926; Dr. M. Bequaert leg.; RMCA • 1 ♂; same location as for preceding; 6 Apr. 1912; Dr. M. Bequaert leg.; RMCA • 1 ♀; same location as for preceding; Jul. 1935; P. Quarré leg.; RMCA • 1 ♂; same location as for preceding; Apr. 1933; De Loose leg.; RMCA • 2 ♂♂; same location as for preceding; De Loose leg.; RMCA • 1 ♀; same location as for preceding; Oct. 1934; P. Quarré leg.; RMCA • 1 ♂; Elisabethville (60km ruiss. Kasepa); 11°40’ S, 27°28’ E; 23 Sep. 1923; Dr. M. Bequaert leg.; RMCA • 1 ♂; Panda (Likasi); 11°00’ S, 26°44’ E; 16 Oct. 1920; Dr. M. Bequaert leg.; RMCA • 1 ♀; Lubumbashi; 11°40’ S, 27°28’ E; 12 Mar. 1921; Dr. M. Bequaert leg.; RMCA. – **Haut-Lomami** • 1 ♀; Lomami: Mutombo-Mukulu; 07°58’ S, 24°00’ E; Jun. 1931; P. Quarré leg.; RMCA • 1 ♂; Sankisia; 09°24’ S, 25°48’ E; 13 Jul. 1911; Dr. J. Bequaert leg.; RMCA. – **Lomami** • 2 ♀♀; Kabinda; 06°08’ S, 24°29’ E; Dr. Schwetz leg.; RMCA. – **Lualaba** • 2 ♀♀; Dilolo; 10°41’ S, 22°21’ E; Aug; 1931; G.F. de Witte leg.; RMCA • 1 ♀; same location as for preceding; Sep.-Oct. 1933; H. De Saeger leg.; RMCA • 1 ♂; Kapanga; 08°21’ S, 22°34’ E; Aug. 1934; R. Verschueren leg.; RMCA • 1 ♂; same location as for preceding; Oct. 1932; Dr. J. Schouteden leg.; RMCA • 1 ♂; Kipiri; 07°35’ S, 29°53’ E; Sep. 1912; R. Verschueren leg.; RMCA • 2 ♀♀ and 1 ♂; Lulua: Kapanga; 08°21’ S, 22°34’ E; Nov. 1932; F.G. Overlaet leg.; RMCA • 1 ♂; same location as for preceding; Nov. 1932; Dr. J. Schouteden leg.; RMCA • 1 ♀ and 1 ♂; same location as for preceding; Dec. 1933; F.G. Overlaet leg.; RMCA • 1 ♂; same location as for preceding; Dec. 1932; F.G. Overlaet leg.; RMCA • 2 ♂♂; same location as for preceding; Oct. 1932; F.G. Overlaet leg.; RMCA • 1 ♂; same location as for preceding; Aug. 1934; F.G. Overlaet leg.; RMCA.

#### *Gronoceras quadrispinosa* (Friese, 1904), *Chalicodoma* (*Gronoceras*) *quadrispinosa* (Friese, 1904)

**Material.** Cupboard 110, Box 34 (3♂♂). **D.R. CONGO**. – **Haut-Katanga** • 1 ♂; PNU Kabwe sur Muye; 08°47’ S, 26°52’ E, alt. 1320 m; 11 May 1948; Mis. G.F. de Witte leg.; RMCA • 2 ♂♂; PNU Lusinga (riv. Kamitungulu); 08°55’ S, 08°55’ S; 13 Jun. 1945; Mis. G.F. de Witte leg.; RMCA.

#### *Noteriades clypeatus* Friese, 1904, *Heriades clypeatus* Friese, 1904

**Material.** Cupboard 110, Box 76 (19♀♀ and 2♂♂). **D.R. CONGO**. – **Haut-Katanga** • 2 ♂♂; Elisabethville (Lubumbashi); 11°40’ S, 27°28’ E; 1931, De Loose leg.; RMCA • 1 ♀; same location as for preceding; 19 Aug. 1932, De Loose leg.; RMCA • 1 ♀; same location as for preceding; 27 Nov. 1933; Dr. M. Bequaert leg.; RMCA • 2 ♀♀; same location as for preceding; Sep. 1934; P. Quarré leg.; RMCA • 1 ♀; Kipushi; 11°46’ S, 27°15’ E; Sep. 1931; G.F. de Witte leg.; RMCA. – **Haut-Lomami** • 1 ♀; Lomami: Kaniama; 07°31’ S, 24°11’ E; 1931, R. Massart leg.; RMCA • 1 ♀; same location as for preceding; 1931, R. Massart leg.; RMCA. – **Ituri** • 1 ♀; Kasenyi; 01°24’ N, 30°26’ E; 1935; H.J. Brédo leg.; RMCA. – **Kasaï** • 1 ♀; Tshikapa; 06°24’ S, 20°48’ E; Apr. 1939, Mevr. Bequaert leg.; RMCA. –**Lualaba Province** • 1 ♀; Lulua: Kapanga; 08°21’ S, 22°34’ E; May 1934; G.F. Overlaet leg.; RMCA. – **Maniema** • 2 ♀♀; Kibombo; 03°54’ S, 25°55’ E; Oct. 1930; Ch. Seydel leg.; RMCA • 3 ♀♀; same location as for preceding; Oct. 1930; Ch. Seydel leg.; RMCA • 1 ♀; same location as for preceding; Oct. 1930; H.J. Brédo leg.; RMCA • 3 ♀♀; Kibombo (P. Or); 03°54’ S, 25°55’ E; Oct. 1930; Ch. Seydel leg.; RMCA.

#### *Noteriades quinquecostatus* (Strand, 1912)

**Material.** Cupboard 110, Box 76 (4 ♀). **D.R. CONGO**. – **Haut-Katanga** • 1 ♀; Elisabethville (Lubumbashi); 11°40’ S, 27°28’ E; 8 Sep.1932; De Loose leg.; RMCA. – **Kongo Central** • 1 ♀; Congo da Lemba (Songololo); 05°42’ S, 13°42’ E; Jan. – Feb. 1913; R. Mayné leg.; RMCA. – **Lualaba** • 2 ♀♀; Lulua: Kapanga; 08°21’ S, 22°34’ E; Mar. 1933; F.G. Overlaet leg.; RMCA.

#### *Noteriades tricarinatus* (Bingham, 1903), *Heriades tricarinatus* (Bingham, 1903)

**Material.** Cupboard 110, Box 76 (31♀♀ and 8♂♂). **D.R. CONGO**. – **Haut-Katanga** • 1 ♂; Elisabethville (Lubumbashi); 11°40’ S, 27°28’ E; 14 Apr. 1912; Dr. J. Bequaert leg.; RMCA • 1 ♀; same location as for preceding; 12 Oct. 1923; Ch. Seydel leg.; RMCA • 1 ♂; same location as for preceding; 14 Mar. 1929; Dr. M. Bequaert leg.; RMCA • 1 ♀; same location as for preceding; 2 Apr. 1928; Dr. M. Bequaert leg.; RMCA • 1 ♀; same location as for preceding; Mar. 1931; Dr. M. Bequaert leg.; RMCA • 1 ♀; same location as for preceding; Dr. M. Bequaert leg.; RMCA • 1 ♀; same location as for preceding; 25-30 Nov. 1930; R. Massart leg.; RMCA • 1 ♀; Lubumbashi; 11°40’ S, 27°28’ E; 18 Jan. 1921; Dr. M. Bequaert leg.; RMCA • 1 ♀; same location as for preceding; 22 Mar. 1911; Dr. J. Bequaert leg.; RMCA • 1 ♀; same location as for preceding; 27 Dec. 1920; Dr. M. Bequaert leg.; RMCA • 1 ♀; same location as for preceding; 9 Nov. 1920; Dr. M. Bequaert leg.; RMCA • 1 ♀; Panda (Likasi); 11°00’ S, 26°44’ E; 2 Oct. 1920; Dr. M. Bequaert leg.; RMCA. – **Haut-Lomami** • 1 ♀; Bukama; 09°12’ S, 25°51’ E; 10 Jul. 1911; Dr. J. Bequaert leg.; RMCA • 1 ♂; same location as for preceding; 16 Jul. 1911; Dr. J. Bequaert leg.; RMCA • 1 ♀; Sankisia; 09°24’ S, 25°48’ E; 19 Sep. 1911; Dr. J. Bequaert leg.; RMCA • 1 ♂; same location as for preceding; 23 Sep. 1911; Dr. J. Bequaert leg.; RMCA • 1 ♀; same location as for preceding; 28 Sep. 1911; Dr. J. Bequaert leg.; RMCA. – **Kasaï** • 4 ♀♀; Ikeke; 04°15’ S, 21°15’ E; 8 May 1946; V. Lagae leg.; RMCA. – **Kinshasa** • 1 ♀; Léopoldville (Kinshasa); 04°19’ S, 15°19’ E; 16 Sep. 1910; Dr. J. Bequaert leg.; RMCA • 1 ♂; same location as for preceding; 18 Sep. 1910; Dr. J. Bequaert leg.; RMCA. – **Kongo Central** • 1 ♀; Congo da Lemba (Songololo); 05°42’ S, 13°42’ E Jan. – Feb. 1913; R. Mayné leg.; RMCA • 4 ♀♀; Kisantu (Madimba); 05°08’ S, 15°06’ E; 1927; R.P. Vanderyst leg.; RMCA • 2 ♀♀; Mayidi (Madimba); 05°11’ S, 15°09’ E; 1942; Rév. P. Van Eyen leg.; RMCA. – **Lualaba** • 2 ♀♀; Lulua: Kapanga; 08°21’ S, 22°34’ E; Mar. 1933; G.F. Overlaet leg.; RMCA • 2 ♂♂; Tenke; 10°35’ S, 26°06’ E; 9 Août 1911; J. Ogilvie leg.; RMCA. – **Maï-Ndombe** • 1 ♀; Kwamouth (Mushie); 03°11’ S, 16°12’ E; Jun. 1913; G.F. Overlaet leg.; RMCA. – **Maniema** • 1 ♀; Kibombo; 03°54’ S, 25°55’ E; Oct. 1930; Ch. Seydel leg.; RMCA • 2 ♀♀; same location as for preceding; Jun. 1930; Ch. Seydel leg.; RMCA.

#### *Tribe Anthidiini Afrostelis tegularis* Cockerell, 1931, *Stelis* (*Afrostelis*) *tegularis* (Cockerell, 1931)

**Material.** Cupboard 110, Boxs 60 (2♀♀ and 1♀). **D.R. CONGO**. Holotype – **Sud-Kivu** • 1 ♂; Kamaniola; 02°47’ S, 29°00’ E; 1 Feb. 1927; Dr. M. Bequaert leg.; RMCA.

– **Haut-Lomami** • 1 ♀; Bukama; 09°12’ S, 25°51’ E; 6 May 1911; Dr. J. Bequaert leg.; RMCA. – **Kasaï Central** • 1 ♂; Luluabourg (Kananga); 05°54’ S, 21°52’ E; 12 May 1919; P. Callewaert leg.; RMCA.

#### *Anthidiellum (Pycnanthidium) absonulum* (Cockerell, 1932), *Anthidiellum* (*Pygnanthidiellum*) *absonulum* (Cockerell, 1932)

**Material.** Cupboard 110, Box 58 (5♀♀). **D.R. CONGO**. – **Equateur** • 1 ♀; Eala (Mbandaka); 00°03’ N, 18°19’ E; Mar.1932; H.J. Brédo leg.; RMCA. – **Haut-Katanga** • 1 ♀; Elisabethville (Lubumbashi); 11°40’ S, 27°28’ E; 11-17 Sep.1931; Dr. M. Bequaert leg.; RMCA • 1 ♀; same location as for preceding; Sep. 1934; P. Quarré leg.; RMCA. • 2 ♂♂. – **Haut-Uele** • 1 ♀; Haut-Uele: Polis-Mboli; 02°46’ N, 27°38’ E; Apr.1947; P.L.G. Benoit leg.; RMCA. – **Sud-Kivu** • 1 ♀; Nzombe (Amont, 200 m près de Mwana); 03°10’ S, 28°33’ E; Aug.-Sep. 1950; Ch. Seydel leg.; RMCA.

#### *Anthidiellum (Pycnanthidium) auriscopatum* (Strand), *Anthidiellum* (*Pygnanthidiellum*) *auriscopatum* (*Strand*)

**Material.** Cupboard 110, Box 58 (1♀). **D.R. CONGO**. – **Haut-Katanga** • 1 ♀; Elisabethville (Lubumbashi); 11°40’ S, 27°28’ E; Oct. 1934; Dr. M. Bequaert leg.; RMCA.

#### *Anthidiellum (Pycnanthidium) zebra* (Friese, 1904), *Anthidiellum* (*Pygnanthidiellum*) *zebra* (Friese, 1904)

**Material.** Cupboard 110, Box 58 (1♀). **D.R. CONGO**. – **Tshopo** • 1 ♀; Yangambi; 00°46’ N, 24°27’ E; May 1959; J. Decelle leg.; RMCA.

#### *Anthidium (Anthidium) niveocinctum* Gerstäcker, 1857, *Anthidium* (*Nivanthidium*) *niveocintum* Vachal

**Material.** Cupboard 110, Box 67 (2♀♀). **D.R. CONGO**. – **Kasaï** • 1 ♀; Tshikapa; 06°24’ S, 20°48’ E; Apr. 1939; Mevr. Bequaert leg.; RMCA. – **Lomami** • 1 ♀; Lomami: Luputa; 11°00’ S, 26°44’ E; 1935; Dr. Bouvier leg.; RMCA.

#### *Anthidium (Severanthidium) severini* Vachal, 1903

**Material.** Cupboard 110, Box 67 (5♀♀ and 2♂♂). **D.R. CONGO**. – **Haut-Katanga** • 1 ♂; Elisabethville (Lubumbashi); 11°40’ S, 27°28’ E; 1933; De Loose leg.; RMCA • 1 ♂; same location as for preceding; 11 Nov. 1933; Dr. M. Bequaert leg.; RMCA • 1 ♀; Kilwa; 09°18’ S, 28°25’ E; Apr. 1931; J. Brédo leg.; RMCA • 1 ♀; Lubumbashi; 11°40’ S, 27°28’ E; 10 Apr. 1920; Dr. M. Bequaert leg.; RMCA. – **Lualaba** • 1 ♀; Lulua: r. Kiongwezi; 08°35’ S, 22°31’ E; 20 Sep. 1933; G.F. Overlaet leg.; RMCA. – **Sud-Kivu** • 1 ♀; Uvira; 03°25’ S, 29°08’ E; Sep. 1958; Pasteels leg.; RMCA. – **Tanganyika** • 1 ♀; Vallée Lukuga; 05°40’ S, 26°55’ E; Nov. 1911; Dr. Schwetz leg.; RMCA.

#### *Anthidium (Severanthidium) sudanicum* Mavromoustakis, 1945

**Material.** Cupboard 110, Box 67 (1♀). **D.R. CONGO**. – **Ituri** • 1 ♀; Lac Albert: Ishwa; 02°13’ N, 31°12’ E; Sep. 1935; H.J. Brédo leg.; RMCA.

#### *Cyphanthidium sheppardi* (Mavromoustakis, 1937),*Trianthidiellum sheppardi* (Mavromoustakis, 1937)

**Material.** Cupboard 110, Box 61 (1♂). **D.R. CONGO**. – **Tanganyika**• 1 ♂; Bassin Lukuga; 05°40’ S, 26°55’ E; Apr. – Jun. 1954; H. De Saeger leg.; RMCA.

#### *Eoanthidium (Clistanthidium)* armaticeps (Friese, 1908)

**Material.** Cupboard 110, Box 59 (3♀♀). **D.R. CONGO**. – **Haut-Katanga** • 1 ♀; Elisabethville(Lubumbashi); 11°40’ S, 27°28’ E; 14 Mar. 1929; Dr. M. Bequaert leg.; RMCA • 1 ♀; same location as for preceding; 31 May 1935; P. Quarré leg.; RMCA • 1 ♀; same location as for preceding; 9 Mar. 1938; H.J. Brédo leg.; RMCA.

#### *Euaspis abdominalis* (Fabricius, 1793), Cupboard 110, Boxs 62, 63 and 64 (334♀♀ and 60♂♂), *Euaspis abdominalis subsp. claripennis* Strand, 1911

**Material.** Cupboard 110, Box 64 (4♀♀ and 1♂). **D.R. CONGO**. – **Haut-Katanga** • 1 ♂; Elisabethville (Lubumbashi); 11°40’ S, 27°28’ E; 15 Mar. 1924; Ch. seydel leg.; RMCA • 1 ♀; same location as for preceding; 27 Nov. 1956; Ch. seydel leg.; RMCA • 1 ♀; same location as for preceding; Nov. 1934; P. Quarré leg.; RMCA • 1 ♀; Kasenga; 10°22’ S, 28°38’ E; Apr. 1931; H.J. Brédo leg.; RMCA. – **Haut-Lomami** • 1 ♀; PNU Kanonga; 09°16’ S, 26°08’ E, alt. 695 m; 13-27 Sep. 1947; Mis. G.F. de Witte leg.; RMCA.

#### *Euaspis abdominalis* (Fabricius, 1793)

**Material.** Cupboard 110, Boxs 62 and 63 (321♀♀ and 59♂♂). **D.R. CONGO**. – **Bas-Uele** • 2 ♀♀; Bambesa; 03°28’ N, 25°43’ E; 1938; P. Henrard leg.; Dr. J. Schouteden leg.; RMCA • 1 ♀; same location as for preceding; 10 Apr. 1937; J. Vrydagh leg.; RMCA • 1 ♀; same location as for preceding; 14 May 1938; P. Henrard leg.; RMCA • 1 ♀; same location as for preceding; 15 Sep. 1933; H.J. Brédo leg.; RMCA • 1 ♀; same location as for preceding; 17 Sep. 1940; J. Vrydagh leg.; RMCA • 1 ♀; same location as for preceding; 2 Oct. 1937; J. Vrydagh leg.; RMCA • 1 ♀; same location as for preceding; 11 Jun. 1937; J. Vrydagh leg.; RMCA • 1 ♀; same location as for preceding; 26 Nov. 1937; J. Vrydagh leg.; RMCA • 1 ♀; same location as for preceding; 27 Jan. 1938; J. Vrydagh leg.; RMCA • 1 ♀; same location as for preceding; 3 Jun. 1940; J. Vrydagh leg.; RMCA • 1 ♀; same location as for preceding; 30 Aug. 1933; J.V. Leroy leg.; RMCA • 1♂; same location as for preceding; 30 Oct. 1933; H.J. Brédo leg.; RMCA • 1 ♀; same location as for preceding; 6 Jul. 1937; J. Vrydagh leg.; RMCA • 1 ♀; same location as for preceding; 8 Dec. 1937; J. Vrydagh leg.; RMCA • 1 ♂; same location as for preceding; Jan. 1937; J. Vrydagh leg.; RMCA • 1 ♀; same location as for preceding; Feb. 1934; H.J. Brédo leg.; RMCA • 1 ♂; same location as for preceding; Feb. 1934; H.J. Brédo leg.; RMCA • 1 ♀; same location as for preceding; Feb. 1937; J. Vrydagh leg.; RMCA • 1 ♀; same location as for preceding; Dec. 1933; J. Vrydagh leg.; RMCA • 1 ♀; same location as for preceding; Dec. 1936; J. Vrydagh leg.; RMCA • 1 ♀; same location as for preceding; Dec. 1938; P. Henrard leg.; RMCA • 1 ♀; same location as for preceding; 11 Apr. 1939; P. Henrard leg.; RMCA • 1 ♀; same location as for preceding; 26 Apr. 1939; P. Henrard leg.; RMCA • 1 ♂; same location as for preceding; Dec. 1946; J. Vrydagh leg.; RMCA • 1 ♂; Dingila; 03°37’ N, 26°03’ E; 12 Oct. 1932; J. Vrydagh leg.; Dr. J. Schouteden leg.; RMCA • 1 ♂; Tshuapa: Bambesa; 03°28’ N, 25°43’ E; 9 Oct. 1952; P. Henrard leg.; RMCA • 1 ♀; Aketi; 02°44’ N, 23°46’ E; 1934; C. Johnen leg.; RMCA • 1 ♀; Uele: Bambesa; 03°28’ N, 25°43’ E; 12 Jun. 1940; J. Vrydagh leg.; RMCA • 1 ♀; same location as for preceding; 10 Aug. 1933; J.V. Leroy leg.; RMCA • 1 ♀; same location as for preceding; 20 Oct. 1933; J.V. Leroy leg.; RMCA • 1 ♀; same location as for preceding; 30 Oct. 1933; J.V. Leroy leg.; RMCA • 1 ♀; same location as for preceding; Mar.-Apr. 1938; P. Henrard leg.; RMCA • 1 ♀; Uele: Ibembo; 02°38’ N, 23°36’ E; 1950; R. Fr. Hutsebaut leg.; RMCA • 1 ♀; Uele: Bondo; Uele: Bondo; 28 Jan. 1950; R.P. Theunissen leg.; RMCA • 1 ♀; same location as for preceding; 1946-1948; R. Fr. Hutsebaut leg.; RMCA • 2 ♀♀; same location as for preceding; 20 May 1950; R. Fr. Hutsebaut leg.; RMCA • 1 ♀; same location as for preceding; 30 Dec. 1951; R. Fr. Hutsebaut leg.; RMCA • 1 ♀; same location as for preceding; Feb. 1952; R. Fr. Hutsebaut leg.; RMCA • 1 ♀; same location as for preceding; Sep. 1949; R. Fr. Hutsebaut leg.; RMCA • 1 ♀; same location as for preceding; Jun. 1952; R. Fr. Hutsebaut leg.; RMCA • 1 ♀; same location as for preceding; Oct. 1949; R. Fr. Hutsebaut leg.; RMCA • 3 ♀♀; Lukulu (Manono); 07°08’ S, 28°06’ E; 1928 – 1932; Vandenbranden leg.; RMCA • 2 ♀♀; Pawa; 02°30’ N, 27°41’ E; 1938; Dr. A. Dubois leg.; RMCA • 1 ♂; Uele-Itimbiri: Aketi; 02°03’ N, 22°45’ E; 1934; C. Johnen leg.;

RMCA. – **Equateur** • 1 ♂; Basankusu; 01°13’ N, 19°49’ E; 1949; Zusters O.L.V. ten Bunderen leg.; RMCA • 4 ♀♀; Coquilhatville (Mbandaka); 00°03’ N, 18°15’ E; 1946; Ch. Scops leg.; Dr. J. Schouteden leg.; RMCA • 1 ♂; same location as for preceding; 1946; Ch. Scops leg.; RMCA • 4 ♀♀; same location as for preceding; 1927; Dr. Strada leg.; RMCA • 1 ♂; same location as for preceding; 1927; Dr. Strada leg.; RMCA • 1 ♂; same location as for preceding; 21 Dec. 1929; D. Bourginuofuon leg.; RMCA • 1 ♀; Wafanya (Monkoto); 01°21’ S, 20°20’ E; 1927; Rév. P. Hulstaert leg.; Dr. J. Schouteden leg.; RMCA • 2 ♀♀; Eala (Mbandaka); 00°03’ N, 18°19’ E; 1937; G. Couteaux leg.; Dr. J. Schouteden leg.; RMCA • 1 ♀; same location as for preceding; 1 Jan. 1921; R. Mayné leg.; RMCA • 2 ♀♀; same location as for preceding; 10 Nov. 1918; R. Mayné leg.; RMCA • 1 ♀; same location as for preceding; 13 Jan. 1921; R. Mayné leg.; RMCA • 1 ♀; same location as for preceding; 13 Nov. 1931; H.J. Brédo leg.; RMCA • 1 ♂; same location as for preceding; 13 Nov. 1931; H.J. Brédo leg.; RMCA • 1 ♀; same location as for preceding; 14 Nov. 1931; H.J. Brédo leg.; RMCA • 1 ♀; same location as for preceding; 19 Oct. 1931; H.J. Brédo leg.; RMCA • 1 ♂; same location as for preceding; 3 Oct. 1931; H.J. Brédo leg.; RMCA • 1 ♀; same location as for preceding;7 Oct. 1931; H.J. Brédo leg.; RMCA • 3 ♀♀; same location as for preceding; 7 Nov. 1931; H.J. Brédo leg.; RMCA • 1 ♂; same location as for preceding; 7 Nov. 1931; H.J. Brédo leg.; RMCA • 1 ♂; same location as for preceding; Jan. 1936; J. Ghesquière leg.; RMCA • 1 ♀; same location as for preceding; Feb. 1935; J. Ghesquière leg.; RMCA • 2 ♀♀; same location as for preceding; Mar. 1932; H.J. Brédo leg.; RMCA • 1 ♀; same location as for preceding; Mar. 1935; J. Ghesquière leg.; RMCA • 2 ♀♀; same location as for preceding; Apr. 1932; H.J. Brédo leg.; RMCA • 1 ♀; same location as for preceding; Apr. 1933; A. Corbisier leg.; RMCA • 3 ♀♀; same location as for preceding; Apr. 1935; J. Ghesquière leg.; RMCA • 2 ♂♂; same location as for preceding; Apr. 1935; J. Ghesquière leg.; RMCA • 1 ♀; same location as for preceding; Sep. 1935; J. Ghesquière leg.; RMCA • 1 ♂; same location as for preceding; May 1932; H.J. Brédo leg.; RMCA • 2 ♂♂; same location as for preceding; May 1935; J. Ghesquière leg.; RMCA • 5 ♀♀; same location as for preceding; Jun. 1932; A. Corbisier leg.; RMCA • 1 ♀; same location as for preceding; Jun. 1935; J. Ghesquière leg.; RMCA • 2 ♀♀; same location as for preceding; Jul. 1935; J. Ghesquière leg.; RMCA • 1 ♂; same location as for preceding; Jul. 1936; J. Ghesquière leg.; RMCA • 1 ♀; same location as for preceding; Aug. 1935; J. Ghesquière leg.; RMCA • 1 ♂; same location as for preceding; Oct. 1929; H.J. Brédo leg.; RMCA • 3 ♀♀; same location as for preceding; Oct. 1935; J. Ghesquière leg.; RMCA • 1 ♀; same location as for preceding; Nov. 1931; H.J. Brédo leg.; RMCA • 5 ♀♀; same location as for preceding; Nov. 1934; J. Ghesquière leg.; RMCA • 1 ♀; same location as for preceding; Nov. 1935; J. Ghesquière leg.; RMCA • 1 ♀; same location as for preceding; Dec. 1932; H.J. Brédo leg.; RMCA • 1 ♂; same location as for preceding; Dec. 1932; H.J. Brédo leg.; RMCA • 2 ♀♀; same location as for preceding; Dec. 1934; J. Ghesquière leg.; RMCA • 1 ♂; Flandria (Ingende); 00°20’ S, 19°06’ E; Jan.-Feb. 1928; Rév. P. Hulstaert leg.; RMCA • 1 ♀; same location as for preceding; Nov. 1931; Rév. P. Hulstaert leg.; RMCA • 1 ♀; Lukolela (Bikoro); 01°03’ S, 17°12’ E; 1951; R. Deguide leg.; RMCA • 2 ♀♀; same location as for preceding; Nov. 1934; J. Ghesquière leg.; RMCA • 1 ♀; Mobeka (Bomongo); 01°53’ N, 19°49’ E; 11 Aug. 1947; Dr. M. Poll leg.; Dr. J. Schouteden leg.; RMCA • 2 ♀♀; Nouvelle Anvers (Bomongo); 01°36’ N, 19°09’ E; 9 Dec. 1952; P. Basilewsky leg.;

RMCA. – **Haut-Katanga** • 1 ♂; Elisabethville (Lubumbashi); 11°40’ S, 27°28’ E; De Loose leg.; Dr. J. Schouteden leg.; RMCA • 1 ♂; same location as for preceding; Dec. 1914; Dr. J. Bequaert leg.; RMCA • 1 ♀; same location as for preceding; Dec. 1914; Dr. J. Bequaert leg.; RMCA • 1 ♂; same location as for preceding; Dec. 1934; Dr. M. Bequaert leg.; RMCA • 1 ♀; same location as for preceding; Dec. 1934; P. Quarré leg.; RMCA. – **Haut-Lomami** • 1 ♀; Kaniama; 07°31’ S, 24°11’ E; 1932; R. Massart leg.; Dr. J. Schouteden leg.; RMCA • 1 ♀; Kisamba; 07°20’ S, 24°02’ E; 1955; Dr. J. Claessens leg.; Dr. J. Schouteden leg.; RMCA. – **Haut-Uele** • 1 ♂; Abimva; 03°38’ N, 30°06’ E; 1925; L. Brgeon leg.; RMCA • 2 ♀♀; Yebo Moto; 03°34’ N, 28°20’ E; 1926; Dr. H. Schouteden leg.; RMCA • 1 ♀; PNG Nagero (Dungu); 03°45’ N, 29°31’ E; 3-29 May 1954; C. Nebay leg.; RMCA • 1 ♀; Bayenga (Wamba); 02°09’ N, 28°00’ E, alt. 810 m; 10 May 1956; R. Castelain leg.; RMCA • 1 ♀; same location as for preceding; 1-10 May 1957; R. Castelain leg.; RMCA • 1 ♂; same location as for preceding; 15 Jul. 1956; R. Castelain leg.; RMCA • 1 ♂; Uele: Bayenga, terr.Wamba (riv Tele); 02°09’ N, 28°00’ E, alt. 820 m, Oct. 1956; R. Castelain leg.; RMCA. – **Ituri** • 1 ♀; Kibali-Ituri: Bayenga, terr. Wamba; 02°09’ N, 28°00’ E, alt. 810 m; Nov. 1955; R. Castelain leg.; RMCA • 1 ♂; Kibali-Ituri: Kilo; 01°50’ N, 30°09’ E; Dec. 1931; Ch. Scops leg.; leg.; RMCA. – **Kasaï** • 1 ♀; Brabanta; 04°20’ S, 20°36’ E; Apr.-May 1949; P. Henrard leg.; RMCA • 1 ♀; Luebo; 05°20’ S, 21°24’ E; Feb. 1931; J.P. Colin leg.; RMCA • 2 ♀♀; Tshikapa; 06°24’ S, 20°48’ E; Apr.1939; Mevr. Bequaert leg.; RMCA • 1 ♀; same location as for preceding; Apr.-May 1939; Mevr. Bequaert leg.; RMCA • 1 ♀; same location as for preceding; Sep. 1921; Mevr. Bequaert leg.;

RMCA. – **Kasaï Central** • 1 ♀; Katoka-Luluabourg (Kananga); 05°54’ S, 21°52’ E; 1933; R.P. Vankerckhoven leg.; RMCA • 1 ♂; same location as for preceding; 1933; R.P. Vankerckhoven leg.; RMCA • 1 ♀; Lulua: r. Luiza; 07°34’ S, 22°40’ E; 15 Oct. 1933; F.G. Overlaet leg.; RMCA • 1 ♀; Luluabourg (Kananga); 05°54’ S, 21°52’ E; P. Callewaert leg.; RMCA • 1 ♀; same location as for preceding; Apr. 1935; Mme. Gillardin leg.; RMCA • 1 ♀; Luluabourg-Katoka (Kananga); 05°54’ S, 21°52’ E; 1939; R.P. Vankerckhoven leg.; RMCA • 1 ♀; Tshibala; 06°21’ S, 21°52’ E; 1949; Zusters van Heule leg.; RMCA. – **Kinshasa** • 3 ♀♀; Kimwenza (Kinshasa); 04°28’ S, 15°17’ E; Jan.-Apr. 1956; Rév. P. Van Eyen leg.; RMCA • 1 ♀; same location as for preceding; May 1956; Rév. P. Van Eyen leg.; RMCA • 1 ♀; Kinshasa; 04°19’ S, 15°19’ E; Nov. 1923; Mme. J. Tinant leg.; RMCA • 1 ♀; Léopoldville (Kinshasa); 04°19’ S, 15°19’ E; 1933; A. Tinant leg.; RMCA • 1 ♂; same location as for preceding; 19 Mar. 1912; Dr. A. Dubois leg.; RMCA • 1 ♀; same location as for preceding; 22 Apr. 1911; Dr. Mouchet leg.; RMCA • 1 ♂; same location as for preceding; 30 Jul. 1911; Dr. A. Dubois leg.; RMCA • 1 ♀; same location as for preceding; 30 Nov. 1911; Dr. A. Dubois leg.; RMCA • 4 ♀♀; same location as for preceding; Dr. Houssiaux leg.; RMCA • 1 ♀; same location as for preceding; May-Jun. 1911; Dr. A. Dubois leg.; RMCA • 2 ♀♀; same location as for preceding; Oct.1955; P. Jobels leg.; RMCA • 1 ♀; same location as for preceding; 1942; R. Fiasse leg.; RMCA. – **Kongo Central** • 1 ♀; Boma; 05°51’ S, 13°03’ E; 1933; Dr. Hallet leg.; RMCA • 1 ♀; Cattier (Lufutoto); 05°26’ S, 14°45’ E; 1946-1949; Delafaille leg.; RMCA • 1 ♀; Kinkenge; 04°51’ S, 13°37’ E; Feb.-Mar. 1951; Mme. M. Bequaert leg.; RMCA • 15 ♀♀; Kisantu (Madimba); 05°08’ S, 15°06’ E; 1927; R. P. Vanderyst leg.; RMCA • 1 ♂; same location as for preceding; 1927; R. P. Vanderyst leg.; RMCA • 41 ♀♀; same location as for preceding; 05°08’ S, 15°06’ E; 1931; R. P. Vanderyst leg.; RMCA • 1 ♂; same location as for preceding; 1931; R. P. Vanderyst leg.; RMCA • 8 ♀♀; same location as for preceding; 1932; R. P. Vanderyst leg.; RMCA • 1 ♀; same location as for preceding; 1 Apr. 1930; R. P. Vanderyst leg.; RMCA • 1 ♀; same location as for preceding; 26 Dec. 1911; Dr. Mouchet leg.; RMCA • 1 ♀; same location as for preceding; Jan. 1950; Rév. Fr. Anastase leg.; RMCA • 2 ♀♀; same location as for preceding; P. Vanderijst leg.; RMCA • 1 ♂; same location as for preceding; May 1945; Rév. Fr. Anastase leg.; RMCA • 1 ♂; same location as for preceding; May 1946; Rév. Fr. Anastase leg.; RMCA • 3 ♀♀; same location as for preceding; Dec. 1927; R.P. Vanderyst leg.; RMCA • 1 ♀; Kivunda Luozi; 04°32’ S, 14°14’ E; 19 May 1951; Mme. M. Bequaert leg.; RMCA • 1 ♀; Lemfu (Madimba); 05°18’ S, 15°13’ E; Jan. 1945; Rév. P. De Beir leg.; RMCA • 1 ♀; same location as for preceding; Feb. 1945; Rév. P. De Beir leg.; RMCA • 1 ♀; same location as for preceding; Dec. 1945; Rév. P. De Beir leg.; RMCA • 1 ♀; Lukula; 05°23’ S, 12°57’ E; Jan. 1952; Dr. R. Wautier leg.; RMCA • 1 ♀; Mayidi (Madimba); 05°11’ S, 15°09’ E; 1912; Rév. P. Van Eyen leg.; RMCA • 5 ♀♀; same location as for preceding; 1942; Rév. P. Van Eyen leg.; RMCA • 1 ♂; same location as for preceding; 1942; Rév. P. Van Eyen leg.; RMCA • 1 ♀; same location as for preceding; 1945; Rév. P. Van Eyen leg.; RMCA • 1 ♀; Thysville (Mbanza Ngungu); 05°15’ S, 14°52’ E; Jan. 1951; P. Basilewsky leg.; RMCA • 1 ♀; same location as for preceding; 1953; J. Sion leg.; RMCA • 1 ♀; Tshela; 04°59’ S, 12°56’ E; 10 Nov. 1920; Dr. H. Schouteden leg.;

RMCA. – **Kwango** • 1 ♀; Kasongolunda; 06°29’ S, 16°49’ E; Jul. 1971; P.V. Van Haelst leg.; RMCA • 1 ♂; Kingungi; 04°51’ S, 16°58’ E; Aug. 1939; R.P. Renier leg.; RMCA • 1 ♂; Mukila (Kenge); 05°01’ S,16°59’ E; Oct. 1952; Dr. M. Bequaert leg.; RMCA • 1 ♀; same location as for preceding; Nov. 1952; Dr. M. Bequaert leg.; RMCA • 1 ♀; Panzi (Kasongo Lunda); 07°13’ S, 17°58’ E; 12 Feb. 1939; Mevr. Bequaert leg.; RMCA • 1 ♀; Popokabaka; 05°42’ S, 16°34’ E; Mar. 1952; L. Pierquin leg.; RMCA. – **Kwilu** • 1 ♀; Banningville (Bandundu); 03°18’ S, 17°21’ E; 1939; Dr. H. Schouteden leg.; RMCA. – **Lomami** • 1 ♂; Lomami: Luputa: Luputa; 07°10’ S, 23°43’ E; Apr. 1934; Dr. Bouvier leg.; RMCA • 1 ♀; Sankuru: M’Pemba Zeo (Gandajika); 06°49’ S, 23°58’ E; 1 Sep. 1958; R. Maréchal leg.; RMCA • 1 ♂; Seke (Lubao); 05°08’ S, 25°44’ E; 26 Jun. 1911; R. Mayné leg.; RMCA. – **Lualaba** • 1 ♀; Kasaï: Lubudi; 09°55’ S, 25°58’ E; 6 Sep. 1929; Legros leg.; RMCA • 1 ♀; Kapanga; 08°21’ S, 22°34’ E; Oct. 1932; F.G. Overlaet leg.; RMCA • 1 ♂; same location as for preceding; Oct. 1932; F.G. Overlaet leg.; RMCA • 1 ♀; same location as for preceding; Oct. 1933; F.G. Overlaet leg.; RMCA • 1 ♀; same location as for preceding; Nov. 1933; F.G. Overlaet leg.; RMCA. – **Maï-Ndombe** • 1 ♀; Bolobo (Mushie); 02°10’ S, 16°14’ E; May 1921; Dr. Mouchet leg.; RMCA • 1 ♀; Inongo; 01°57’ S, 18°16’ E; Aug. 1913; Dr. J. Maes leg.; RMCA • 1 ♂; Kutu; 02°44’ S, 18°45’ E; Oct. 1913; R. Mayné leg.; RMCA • 1 ♀; Kwamouth (Mushie); 03°11’ S, 16°12’ E; Jun. 1913; Dr. J. Maes leg.; RMCA • 1 ♀; Wombali (Mushie); 03°16’ S,17°22’ E; Jul. 1913; P. Vanderijst leg.; RMCA. – **Maniema** • 1 ♀; Maniema; 05°00’ S, 29°00’ E; 1936; Guraut leg.; RMCA. – **Mongala** • 1 ♀; Lisala; 02°09’ N, 21°30’ E; 1940; C. Leontovitch leg.; RMCA • 1 ♀; Terr. Lisala; 02°09’ N, 21°30’ E; Mar. 1937; C. Leontovitch leg.; RMCA • 1 ♀; Ubangi: Binga (Lisala); 02°28’ N, 20°31’ E; 5-12 Mar. 1932; H.J. Brédo leg.; RMCA • 1 ♀; Ubangi: Bumba; 02°11’ N, 22°32’ E; 14 Oct. 1939; C. Leontovitch leg.; RMCA. – **Nord-Kivu** • 1 ♂; Buseregenye (Rutshuru); 01°11’ S, 29°27’ E; 1930; Ed. Luja leg.; RMCA • 1 ♀; Cité de Kimbulugu (Lubero); 00°10’ S, 29°14’ E; Mar. 1939; Mevr. Bequaert leg.; RMCA • 1 ♀; PNA [Parc National Albert]; 00°03’ S, 29°30’ E; Mar. 25 Jun. 1957; P. Vanschuytbroeck & H. Synave leg.; RMCA. – **Nord-Ubangi** • 1 ♀; Ubangi: Karawa (Businga); 03°21’ N, 20°18’ E; 1936; Rév. Wallin leg.; RMCA. – **Sankuru** • 1 ♀; Djeka; 05°00’ S, 23°28’ E; Jan. 1954; R. Roiseux leg.; RMCA • 2 ♀♀; Kasaï: Kondoue (Lusambo); 04°58’ S, 23°16’ E; Ed. Luja leg.; RMCA • 1 ♀; Lusambo; 04°58’ S, 23°26’ E; 19 Sep. 1949; Dr. M. Fontaine leg.; RMCA. – **Sud-Kivu** • 1 ♀; Costermansville (Bukavu); 02°29’ S, 28°51’ E; 1939; Dr. Hautmann leg.; RMCA • 1 ♂; same location as for preceding; 1939; Dr. Hautmann leg.; RMCA • 2 ♀♀; Lubungola pr. Shabunda; 02°41’ S, 21°21’ E; Feb. 1939; Dr. Hautmann leg.; RMCA • 1 ♂; same location as for preceding; Feb. 1939; Dr. Hautmann leg.; RMCA • 1 ♀; Région des lacs, 02°29’ S, 28°51’ E; Dr. Sagona leg.; RMCA • 1 ♀; Uvira; 03°25’ S, 29°08’ E; Oct. 1949; G. Marlier leg.; RMCA. – **Sud-Ubangi** • 1 ♀; Kunungu; 02°45’ N, 18°26’ E; 1932; Dr. H. Schouteden leg.; RMCA • 5 ♀♀; Libenge; 03°39’ N, 18°38’ E; 1931; J. Van Gils leg.; RMCA • 1 ♀; same location as for preceding; Aug. 1937; C. Leontovitch leg.; RMCA • 1 ♂; same location as for preceding; Dec. 1937; C. Leontovitch leg.; RMCA • 1 ♀; Ubangi: Libenge; 03°39’ N, 18°38’ E; Mar. 1936; C. Leontovitch leg.; RMCA • 1 ♂; Ubangi: Libenghi; 03°39’ N, 18°38’ E; 29 Aug. 1926; Cap. Van Gils leg.; RMCA • 2 ♀♀; same location as for preceding; Vreurick leg.; RMCA • 2 ♀♀; Zulu sur Lua (Ubangi); 02°45’ N, 18°26’ E; 1931; J. Van Gils leg.; RMCA • 2 ♂♂; same location as for preceding; 1931, J. Van Gils leg.;

RMCA. – **Tanganyika** • 1 ♀; Katanga: Kabinda (Moba); 07°02’ S, 27°47’ E; Jun. 1930; Ch. Seydel leg.; RMCA. – **Tshopo** • 1 ♀; Aruwimi; 01°13’ N, 23°36’ E; Jun. 1926; Dr. H. Schouteden leg.; RMCA • 1 ♀; Basoko; 01°13’ N, 23°36’ E; Sep. 1948; P.L.G. Benoit leg.; RMCA • 1 ♀; same location as for preceding; May 1949; P.L.G. Benoit leg.; RMCA • 1 ♀; Basoko Yaeberomans, 01°13’ N, 23°36’ E; Apr. 1949; P.L.G. Benoit leg.; RMCA • 1 ♀; Elisabetha (Yahuma); 01°09’ N, 23°37’ E; Mme. J. Tinant leg.; RMCA • 1 ♀; Stanleyville (Kisangani); 00°31’ N, 25°11’ E; 1936; A. Becquet leg.; RMCA • 1 ♀; same location as for preceding; 13 – 23 Aug. 1928; A. Collart leg.; RMCA • 1 ♀; same location as for preceding; Jan. 1926; Lt. J. Ghesquière leg.; RMCA • 1 ♀; same location as for preceding; Feb. 1948; Dr. R. Mouchamps leg.; RMCA • 1 ♀; same location as for preceding; Mar. 1926; J. Ghesquière leg.; RMCA • 1 ♂; same location as for preceding; Mar. 1926; J. Ghesquière leg.; RMCA. – **Tshuapa** • 1 ♀; Bokuma; 00°06’ S, 18°41’ E; 1938; Rév. P. Hulstaert leg.; RMCA • 2 ♀♀; same location as for preceding; Feb. – Apr. 1941; Rév. P. Hulstaert leg.; RMCA • 1 ♂; same location as for preceding; May 1935; Rév. P. Hulstaert leg.; RMCA • 1 ♀; same location as for preceding; Feb. 1952; Rév. P. Lootens leg.; RMCA • 1 ♀; same location as for preceding; Mar. 1952; Rév. P. Lootens leg.; RMCA • 1 ♀; same location as for preceding; Aug. – Sep. 1951; Rév. P. Lootens leg.; RMCA • 1 ♀; same location as for preceding; Dec. 1951; Rév. P. Lootens leg.; RMCA • 1 ♀; Bolima (Bolomba); 00°03’ N, 19°23’ E; 17 – 28 Feb. 1939; Rév. P. Hulstaert leg.; RMCA • 1 ♀; Tshuapa: Bamania (Mbandaka); 00°01’ N, 18°19’ E; Jan. 1954; Rév. P. Hulstaert leg.; RMCA • 1 ♀; same location as for preceding; Mar. 1954; Rév. P. Hulstaert leg.; RMCA • 2 ♀♀; same location as for preceding; Jan. – Jun. 1965; Rév. P. Hulstaert leg.; RMCA • 1 ♀; same location as for preceding; May 1960; Rév. P. Hulstaert leg.; RMCA • 1 ♀; same location as for preceding; Oct. 1951; Rév. P. Hulstaert leg.; RMCA • 1 ♀; same location as for preceding; Nov. 1951; Rév. P. Hulstaert leg.; RMCA • 3 ♀♀; Tshuapa: Bamanya (Mbandaka); 00°01’ N, 18°19’ E; 1968; Rév. P. Hulstaert leg.; RMCA • 1 ♀; same location as for preceding; Jan. 1963; Rév. P. Hulstaert leg.; RMCA • 1 ♀; same location as for preceding; Jan. – Jun. 195; Rév. P. Hulstaert leg.; RMCA • 1 ♀; same location as for preceding; May 1960; Rév. P. Hulstaert leg.; RMCA • 1 ♀; Tshuapa: Boende; 00°13’ S, 20°52’ E; 15 – 30 May 1951; Rév. P. Hulstaert leg.; RMCA • 1 ♀; Tshuapa: Bokuma; 00°06’ S, 18°41’ E; 1952; Rév. P. Lootens leg.; RMCA • 1 ♀; same location as for preceding; Mar. 1954; Rév. P. Lootens leg.; Dr. J. Schouteden leg.; RMCA • 1 ♂; same location as for preceding; Mar. 1954; Rév. P. Lootens leg.; Dr. J. Schouteden leg.; RMCA • 1 ♀; same location as for preceding; Jun. 1952; Rév. P. Lootens leg.; RMCA • 1 ♀; Tshuapa: Flandria; 00°20’ S, 19°06’ E; 1940; Rév. P. Hulstaert leg.; RMCA • 1 ♂; same location as for preceding; 1946; Rév. P. Hulstaert leg.; RMCA • 1 ♂; same location as for preceding; Jan. 1946; Rév. P. Hulstaert leg.; RMCA • 2 ♀♀; same location as for preceding; Apr.-May 1948; Rév. P. Hulstaert leg.; RMCA • 2 ♀♀; same location as for preceding; Sep. – Dec. 1947; Rév. P. Hulstaert leg.; RMCA.

#### *Euaspis abdominalis subsp. martini* Vachal, 1910

**Material.** Cupboard 110, Box 64 (9♀♀). **D.R. CONGO**. – **Equateur** • 1 ♀; Eala (Mbandaka); 00°03’ N, 18°19’ E; Sep.1935; J. Ghesquière leg.; RMCA. – **Haut-Katanga** • 2 ♀♀; Elisabethville (Lubumbashi); 11°40’ S, 27°28’ E; Jun.1926; Dr. M. Bequaert leg.; RMCA, • 1 ♀; Kasenga; 10°22’ S, 28°38’ E; Jun.1926; H.J. Brédo leg.; RMCA. – **Nord-Uele** • 1 ♀; Wamba (Uele Nepoko); 00°20’ S, 29°50’ E, alt. 1400 m; 25 Apr.1931; J. Vrydagh leg.; RMCA. – **Sud-Kivu** • 1 ♀; Lubungola pr. Shabunda; 02°41’ S, 21°21’ E; 1939; Dr. Hautmann leg.; RMCA, • 1 ♀; Mulungu; 02°20’ S, 28°47’ E; Jun. 1935; J.V. Leroy leg.; RMCA, • 1 ♀; Uvira: dans habitation; 03°25’ S, 29°08’ E; Aug.-Dec. 1949; G. Marlier leg.; RMCA. –**Tanganyika** • 1 ♀; Tanganyika: Moba; 07°02’ S, 29°47’ E; 1939; H. Bomans leg.; RMCA.

#### *Euaspis erythros* (Meunier, 1890)

**Material.** Cupboard 110, Box 64 (43♀♀ and 5♂♂). **D.R. CONGO**. – **Haut-Katanga** • 1 ♀; Elisabethville (Lubumbashi); 11°40’ S, 27°28’ E; 1 Feb. 1935; P. Quarré leg.; RMCA • 1 ♀; same location as for preceding; 3 Mar. 1921; Dr. M. Bequaert leg.; RMCA • 1 ♀; same location as for preceding; 31 Jan. 1921; Dr. M. Bequaert leg.; RMCA • 1 ♀; same location as for preceding; Mar. 1935; P. Quarré leg.; RMCA • 1♀; same location as for preceding; Apr. 1928; Dr. M. Bequaert leg.; RMCA • 1 ♀; same location as for preceding; Oct. 1934; P. Quarré leg.; RMCA • 2 ♀♀; Kampombwe; 11°40’ S, 27°28’ E; Apr. 1931; H.J. Brédo leg.; RMCA • 1 ♀; same location as for preceding; 5 – 9 Oct. 1951; Ch; seydel leg.; RMCA. – **Haut-Lomami** • 1 ♀; Lulua: Kabongo; 07°20’ S, 25°35’ E; 31 Jul. 1952; Ch. Seydel leg.; RMCA. – **Haut-Uele** • 1 ♀; Haut-Uele: Abimva; 03°44’ N, 29°42’ E; 1925; L. Burgeon leg.; RMCA • 1 ♀; Haut-Uele: Yebo Moto; 02°46’ N, 27°38’ E; Dec. 1926; L. Burgeon leg.; RMCA. – **Ituri** • 2 ♀♀; Kasenyi; 01°24’ N, 30°26’ E; 22 Aug.1935; H.J. Brédo leg.; RMCA • 3 ♂♂; same location as for preceding; 22 Aug.1935; H.J. Brédo leg.; RMCA • 2 ♀♀; Lac Albert: Kasenyi; 01°24’ N, 30°26’ E; 15 May 1935; H.J. Brédo leg.; RMCA • 1♀; same location as for preceding; Dec.1935; H.J. Brédo leg.; RMCA. – **Kinshasa** • 2 ♀♀; Léopoldville (Kinshasa); 04°19’ S, 15°19’ E; 1933; A. Tinant leg.; RMCA • 6 ♀♀; same location as for preceding; Aug. 1949; Dr. A. Dubois leg.; RMCA • 1 ♂; same location as for preceding; 1948; Dr. Richard leg.; RMCA • 1♂; Limete (Léo); 04°19’ S, 15°19’ E; 1958; A. Froidebise leg.; RMCA. – **Kongo Central** • 1 ♀; Banana; 06°00’ S, 12°24’ E; Mar. 1951; I. Mesmaekers leg.; RMCA • 1 ♀; Boma; 05°51’ S, 13°03’ E; 1935; W. Moreels leg.; RMCA • 2 ♀♀; same location as for preceding; Aug. 1937; Dr. Schlesser leg.; RMCA • 2 ♀♀; same location as for preceding; 31 Mar. 1913; Lt. Styczynski leg.; RMCA • 1 ♀; same location as for preceding; Jan. 1951; I. Mesmaekers leg.; RMCA • 1 ♀; Boma-Coquilhatville; 05°51’ S, 13°03’ E; Thisquens leg.; RMCA • 2 ♀♀; Kisantu (Madimba); 05°08’ S, 15°06’ E; 1931; R.P. Vanderyst leg.; RMCA • 1 ♀; same location as for preceding; 4 Jan. 1926; R.P. Vanderyst leg.; RMCA. – **Kwango** • 1 ♀; Kwango: Wamba; 04°51’ S, 16°58’ E; 1938; A. Henrion leg.; RMCA. – **Lualaba** • 1 ♀; Lulua: Kapanga; 08°21’ S, 22°34’ E; Oct. 1932; F.G. Overlaet leg.; RMCA. – **Sud-Kivu** • 2 ♀♀; Uvira: dans habitatioN, 03°25’ S, 29°08’ E; Aug. – Dec. 1949; G. Marlier leg.; RMCA. – **Tanganyika** • 1 ♀; Albertville (Kalemie); 05°57’ S, 29°12’ E; May – Jun. 1954; H. Bomans leg.; RMCA. • 2 ♀♀; Tanganyika: Moba; 07°02’ S, 29°47’ E, alt. 780 m; Jul. – Oct. 1953; H. Bomans leg.; RMCA. – **Tshopo** • 1 ♂; Mfungwe-Kayumbe; 01°17’ S, 26°16’ E; Jul. 1907; Dr. Sheffield Neave leg.; RMCA.

#### *Euaspis rufiventris subsp. uvirensis* Cockerell, 1933

**Material.** Cupboard 110, Box 64 (1♀). **D.R. CONGO**. Holotype – **Sud-Kivu** • 1 ♀; Uvira; 03°25’ S, 29°08’ E; 1927; Ch. Seydel leg.; RMCA

#### *Pachyanthidium africanum* (Smith, 1854)

**Material.** Cupboard 110, Box 65 (1♂). **D.R. CONGO**. – **Tanganyika** • 1 ♂; Tanganyika-Moero: Nyunzu; 05°57’ S, 28°01’ E; Jan. – Feb. 1934; H. De Saeger leg.; RMCA.

#### *Pachyanthidium (Trichanthidium) benguelense* (Vachal, 1903)

**Material.** Cupboard 110, Box 66 (6♀♀ and 2♂♂). **D.R. CONGO**. – **Haut-Katanga** • 1 ♂; Elisabethville (Lubumbashi); 11°40’ S, 27°28’ E; De Loose leg.; RMCA • 1 ♀; same location as for preceding; 2 May 1933; Dr. M. Bequaert leg.; RMCA • 1 ♀; same location as for preceding; 20 Sep. 19323; Dr. M. Bequaert leg.; RMCA • 1 ♀; same location as for preceding; 8 Apr. 1933; Dr. M. Bequaert leg.; RMCA • 1 ♀; same location as for preceding; Mar. 1931; Dr. M. Bequaert leg.; RMCA • 1 ♂; Lubumbashi; 11°40’ S, 27°28’ E; 10 Jun. 1920; Dr. M. Bequaert leg.; RMCA • 1 ♀; same location as for preceding; 24 Apr. 1920; Dr. M. Bequaert leg.; RMCA. – **Haut-Lomami** • 1 ♀; Lualaba: Kabongo; 07°20’ S, 25°35’ E; 18 Nov. 1952; Ch. Seydel leg.; RMCA.

#### *Pachyanthidium (Pachyanthidium) bouyssoui* (Vachal, 1903)

**Material.** Cupboard 110, Box 65 (27♀♀ and 12♂♂). **D.R. CONGO**. – **Bas-Uele** • 1 ♂; Bambesa; 03°28’ N, 25°43’ E; 15 Sep. 1933; H.J. Brédo leg.; RMCA • 3 ♀♀; same location as for preceding; 25 Sep. 1934;

H.J. Brédo leg.; RMCA • 1 ♀; same location as for preceding; 30 Oct. 1933; H.J. Brédo leg.; RMCA • 1 ♀; same location as for preceding; Dec. 1933; H.J. Brédo leg.; RMCA • 1 ♀; Uele: Bambesa; 03°28’ N, 25°43’ E; 10 Oct. 1933; J.V. Leroy leg.; RMCA. – **Equateur** • 1 ♂; Eala (Mbandaka); 00°03’ N, 18°19’ E; 16 Oct. 1936; G. Couteaux leg.; RMCA • 2 ♀♀; same location as for preceding; Apr. 1932; H.J. Brédo leg.; RMCA • 1 ♂; same location as for preceding; May 1932; H.J. Brédo leg.; RMCA • 1 ♀; same location as for preceding; Apr. 1932; A. Corbisier leg.; RMCA • 2 ♀♀; same location as for preceding; Jun. 1932; A. Corbisier leg.; RMCA • 6 ♂♂; same location as for preceding; Jun. 1932; A. Corbisier leg.; RMCA • 1 ♀; same location as for preceding; May 1935; J. Ghesquière leg.; RMCA • 1 ♀; same location as for preceding; Jul. 1932; J. Ghesquière leg.; RMCA • 1 ♀; Eala-Bokatola-Bikoro; 00°03’ N, 18°19’ E; Sep. – Oct. 1930; Dr. P. Staner leg.; RMCA. – **Haut-Lomami** • 1 ♀; PNU Gorge de la Petenge; 08°56’ S, 27°12’ E, alt. 1150 m; 19 Jun. 1947; Mis. G.F. de Witte leg.; RMCA. – **Ituri** • 1 ♀; Ituri: Bunia; 01°34’ N, 30°15’ E; 1938; P. Lefèvre leg.; RMCA. – **Kasaï Central** • 1 ♀; Lulua; 05°54’ S, 22°25’ E; 1919; Dr. Walker leg.; RMCA • 1♂; Luluabourg (Kananga); 05°54’ S, 21°52’ E; P. Callewaert leg.; RMCA. – **Lualaba** • 1 ♀; Lulua: Kapanga; 08°21’ S, 22°34’ E; 30 Nov. 1932; F.G. Overlaet leg.; RMCA. – **Maï-Ndombe** • 1 ♂; Bena Bendi (Oshwe); 04°18’ S, 20°22’ E; May 1915; R. Mayné leg.; RMCA • 2 ♀♀; Bomboma; 02°24’ N, 18°53 E; 21 Jul. 1935; A. Bal leg.; RMCA. – **Maniema** • 1 ♀; Malela; 04°22’ S, 26°08’ E; 30 Feb. 1913; L. Burgeon leg.; RMCA. – **Sankuru** • 1 ♀; Sankuru: Komi; 03°23’ S, 23°46’ E; 13 Mar. 1930; J. Ghesquière leg.; RMCA. – **Sud-Kivu** • 1 ♀; Lolo; 02°16’ S, 27°43’ E; 23 May 1925; Dr. Rodhain leg.; RMCA. – **Tanganyika** • 1 ♂; Bassin Lukuga; 05°40’ S, 26°55’ E; Apr. – Jul. 1934; H. De Saeger leg.; RMCA. – **Tshopo** • 1 ♀; Ile Bertha; 00°33’ N, 24°56’ E; 18 Oct. 1910; Dr. J. Bequaert leg.; RMCA. – **Tshuapa** • 1 ♀; Tshuapa: Bokuma; 00°06’ S, 18°41’ E; Mar. 1952; Rév. P. Lootens leg.; RMCA.

#### *Pseudoanthidium (Branthidium) braunsi* (Friese, 1904)

**Material.** Cupboard 110, Box 59 (1♀). **D.R. CONGO**. – **Haut-Katanga** • 1 ♀; Elisabethville (Lubumbashi); 11°40’ S, 27°28’ E; 17 Oct. 1923; Ch. Seydel leg.; RMCA.

#### *Pachyanthidium (Pachyanthidium) katangense* Cockerell, 1930

**Material.** Cupboard 110, Box 65 (4♀♀ and 2♂♂). **D.R. CONGO**. Holotype – **Haut-Katanga** • 1 ♂; Lubumbashi; 11°40’ S, 27°28’ E; 23 May 1920; Dr. M. Bequaert leg.; RMCA.

– **Haut-Katanga** • 1 ♀; PNU Kabwe sur Muye; 08°47’ S, 26°52’ E, alt. 1320 m; 11 May 1948; Mis. G.F. de Witte leg.; RMCA. – **Haut-Lomami** • 1 ♀; Kilenge; 09°08’ S, 25°52’ E; Apr. 1923; Ch. Seydel leg.; RMCA. – **Lualaba** • 1 ♂; Katentenia; 10°19’ S, 25°54’ E; Oct. 1931; Ch. Seydel leg.; RMCA. – **Sud-Kivu** • 1 ♀; Uvira; 03°25’ S, 29°08’ E; Nov. 1927; Ch. Seydel leg.; RMCA. – **Tanganyika** • 1 ♀; Tembwe (Tanganyika); 06°30’ S, 29°26’ E; Feb. 1926; Dr. H. Schouteden leg.; RMCA.

#### *Pachyanthidium (Pachyanthidium) paulinieri* (Guérin-Méneville, 1845), *Pachyanthidium paulinieri* Smith^6^

**Material.** Cupboard 110, Box 65 (5♀♀ and 3♂♂). **D.R. CONGO**. – **Bas-Uele** • 1 ♂ ; Bambesa ; 03°28’ N, 25°43’ E ; 10 Jul. 1933 ; J.V. Leroy leg. ; RMCA. – **Ituri** • 1 ♀ ; Ituri : Bunia ; 01°34’ N, 30°15’ E ; 1938 ; P. Lefèvre leg. ; RMCA. – **Kongo Central** • 1 ♂ ; Bateke-Nord ; 03°30’ S, 15°45’ E ; 1942 ; R.C. Eloy leg. ; RMCA • 1 ♀ ; Mayidi (Madimba) ; 05°11’ S, 15°09’ E ; 1942 ; 1942 ; Rév. P. Van Eyen leg. ; RMCA. – **Kwilu** • 1 ♀ ; Terr. de Baningville (riv Lune, Mpo) ; 03°18’ S, 17°21’ E; 1945; Dr. Fain leg.; RMCA. – **Lomami** • 1 ♀ ; Sankuru : M’Pemba Zeo ; 06°49’ S, 23°58’ E ; 3 Jun. 1960; R. Maréchal leg.; RMCA. – **Maï-Ndombe** • 1 ♀; Bumbuli; 03°24’ S, 20°31’ E; 1915; R. Mayné leg.; RMCA • 1 ♀; Wombali (Mushie); 03°16’ S, 17°22’ E; 30 Sep. 1913; P. Vanderijst leg.; RMCA.

#### *Pseudoanthidium* (*Immanthidium*) *integrum* (Friese, 1905), *Immanthidium integrum* (Friese, 1905)

**Material.** Cupboard 110, Box 72 (1♀). **D.R. CONGO**. – **Haut-Uele** • 1 ♀; PNG [Parc National de Garamba]; 03°40’ N, 29°00’ E; 29 Nov. 1951; H. De Saeger leg.; RMCA.

#### *Pseudoanthidium* (*Micranthidium*) *lanificum* (Smith, 1879), *Micranthidium lanificum* Smith, 1879

**Material.** Cupboard 110, Box 73 (84♀♀ and 7♂♂). **D.R. CONGO**. – **Bas-Uele** • 2 ♂♂; Bambesa; 03°28’ N, 25°43’ E; 1933; H.J. Brédo leg.; RMCA • 3 ♀♀; same location as for preceding; 15 Sep. 1933; H.J. Brédo leg.; RMCA • 3 ♀♀; same location as for preceding; Dec. 1933; H.J. Brédo leg.; RMCA • 1 ♀; same location as for preceding; 31 Aug. 1933; J.V. Leroy leg.; RMCA • 2 ♀♀; same location as for preceding; Sep. 1933; J.V. Leroy leg.; RMCA • 1 ♂; Bas-Uele; 04°00’ N, 22°15’ E; Jul.-Aug. 1920; L. Burgeon leg.; RMCA • 4 ♀♀; Uele: Bambesa; 03°28’ N, 25°43’ E; 10 Oct. 1933; J.V. Leroy leg.; RMCA • 1 ♀; Uele: Dingila; 03°37’ N, 26°03’ E; Aug. 1933; H.J. Brédo leg.; RMCA • 1 ♀; same location as for preceding; 15 Jul. 1933; J.V. Leroy leg.; RMCA • 1 ♀; same location as for preceding; 27 Jul. 1933; J.V. Leroy leg.; RMCA. – **Equateur** • 2 ♀♀; Eala (Mbandaka); 00°03’ N, 18°19’ E; 1932; A. Corbisier leg.; RMCA • 1 ♀; same location as for preceding; Mar. 1935; A. Corbisier leg.; RMCA • 2 ♀♀; same location as for preceding; Nov. 1931; H.J. Brédo leg.; RMCA • 1 ♀; same location as for preceding; Jan. 1936; J. Ghesquière leg.; RMCA • 1 ♀; same location as for preceding; Mar. 1935; J. Ghesquière leg.; RMCA • 1 ♀; same location as for preceding; Sep. 1935; J. Ghesquière leg.; RMCA • 3 ♀♀; same location as for preceding; May 1935; J. Ghesquière leg.; RMCA • 1 ♀; same location as for preceding; Jun. 1935; J. Ghesquière leg.; RMCA • 1 ♂; same location as for preceding; Aug. 1938; J. Ghesquière leg.; RMCA • 3 ♀♀; same location as for preceding; Nov. 1935; J. Ghesquière leg.; RMCA • 1 ♂; same location as for preceding; Nov. 1935; J. Ghesquière leg.; RMCA • 2 ♀♀; Ubangi: Nouvelle Anvers (Bomongo); 00°03’ N, 18°19’ E; 9 Dec. 1952; P. Basilewsky leg.; RMCA. – **Haut-Lomami** • 8 ♀♀; Lualaba: Kaniama; 07°31’ S, 24°11’ E; 19 Dec. 1952; Ch. Seydel leg.; RMCA. – **Haut-Uele** • 1 ♂; Haut-Uele: Abimva; 03°44’ N, 29°42’ E; 1925; L. Burgeon leg.; RMCA • 1 ♀; Mayumbe; 05°30’S, 12°53’E; 1917; R. Mayné leg.; RMCA • 1 ♀; Mayumbe Hiolo; 05°30’S, 12°53’E; 19 Jun. 1924; A. Collart leg.; RMCA • 1 ♀; Uele: Bayenga, terr. Wamba; 02°09’ N, 28°00’ E, alt. 810 m; 25 Nov. 1956; R. Castelain leg.; RMCA • 1 ♀; Watsa à Niangara; 03°02’ N, 29°32’ E; Jul. 1920; L. Burgeon leg.;

RMCA. – **Kongo Central**• 1 ♀; Banza Manteka; 05°28’ S, 13°47’ E; 10-15 Jun. 1912; R. Mayné leg.; RMCA • 1 ♀; Congo da Lemba (Songololo); 05°42’ S, 13°42’ E; Jan.-Feb. 1913; R. Mayné leg.; RMCA • 2 ♀♀; Luali (Tshela); 05°06’S, 12°28’E; 26 Aug. 1913; Dr. J. Bequaert leg.; RMCA • 1 ♂; same location as for preceding; 26 Aug. 1913; Dr. J. Bequaert leg.; RMCA • 1 ♀; same location as for preceding; Aug. 1913; Dr. J. Bequaert leg.; RMCA. – **Lualaba** • 1 ♂; Lulua: Kapanga; 08°21’ S, 22°34’ E; Mar. 1933; G.F. Overlaet leg.; RMCA • 2 ♀♀; same location as for preceding; Apr. 1933; G.F. Overlaet leg.; RMCA • 1 ♀; same location as for preceding; Sep. 1928; G.F. Overlaet leg.; RMCA • 1 ♀; same location as for preceding; May 1933; G.F. Overlaet leg.; RMCA • 1 ♀; same location as for preceding; Nov. 1932; G.F. Overlaet leg.; RMCA. – **Maï-Ndombe** • 3 ♀♀; Bena Bendi (Oshwe); 04°18’ S, 20°22’ E; May 1915; R. Mayné leg.; RMCA. – **Maniema** • 2 ♀♀; Lomami: Kisamba; 07°20’ S, 24°02’ E; Jan. 1931; P. Quarré leg.; RMCA • 2 ♀♀; Maniema; 05°00’ S, 29°00’ E; R. Mayné leg.; RMCA. • 1 ♀; Terr. De Kasongo, Riv. Lumami; 04°27’ S, 26°40’ E; Jan. 1950; P.L.G. Benoit leg.; RMCA. – **Mongala** • 1 ♀; Binga (Lisala); 02°28’ N, 20°31’ E²; 1 Mar. 1932; H.J. Brédo leg.; RMCA • 1 ♀; Bumba; 02°11’ N, 22°32’ E; Dec. 1939-Jan. 1940; H. De Saeger leg.; RMCA • 1 ♀; Ubangi: Binga (Lisala); 02°28’ N, 20°31’ E; 5-12 Mar. 1932; H.J. Brédo leg.; RMCA. – **Nord-Kivu** • 1 ♀; Masisi-Walikale; 01°24’ S, 28°49’ E; 4 Jan. 1915; Dr. J. Bequaert leg.; RMCA • 1 ♀; N’Guli (Lubero); 00°10’ S, 29°20’ E; Sep. 1913; Dr. Rodhain leg.; RMCA • 1 ♀; PNA [Parc National Albert]; 00°03’ S, 29°30’ E; 8 Jan. 1954; H. Synave leg.; RMCA • 1 ♀; same location as for preceding; 7-15 Jul. 1955; P. Vanschuytbroeck leg.; RMCA • 1 ♀; Rutshuru; 01°11’ S, 29°27’ E; Nov. 1937; J. Ghesquière leg.;

RMCA. – **Nord-Ubangi** • 1 ♀; Abumombazi (Mobayi); 03°34’ N, 22°03’ E; 18-26 Feb. 1932; H.J. Brédo leg.; RMCA • 1 ♀; same location as for preceding; 23 Nov. 1932; H.J. Brédo leg.; RMCA. – **Sud-Kivu** • 1 ♀; Lac Kivu: Rwankwi; 01°20’ S, 29°22’ E; Jul. 1951; J.V. Leroy leg.; RMCA. – **Tanganyika** • 1 ♀; Bassin Lukuga; 05°40’ S, 26°55’ E; Apr.-Aug. 1934; H. De Saeger leg.; RMCA • 1 ♀; Kampunda; 08°14’ S, 29°46’ E; 10 Nov. 1914; Dr. Mouchet leg.; RMCA. – **Tshopo** • 1 ♀; Basoko Yamabuti; 01°13’ N, 23°36’ E; 16 Mar. 1948; P.L.G. Benoit leg.; RMCA. – **Tshuapa** • 1 ♀; Itoka (Djolu); 00°10’ S, 23°05’ E; Oct. 1912; R. Mayné leg.; RMCA • 1 ♀; Lokolenge (Lulonga); 01°11’ N, 22°40’ E; 19 May 1927; J. Ghesquière leg.; RMCA • 1 ♀; same location as for preceding; 00°06’ S, 18°41’ E; 1953; R.P. Lootens leg.; RMCA • 1 ♀; Tshuapa: Bokungu; 00°41’ S, 22°19’ E; 1949; Dupuis leg.; RMCA.

#### *Pseudoanthidium* (*Gnathanthidium*) *prionognathum* (Mavromoustakis, 1938), *Gnathanthidium prionognathum* Friese

**Material.** Cupboard 110, Box 72 (2♀♀). **D.R. CONGO**. – **Maniema** • 1 ♀; Kasongo; 04°27’ S, 26°40’ E; Sep. 1959; P.L.G. Benoit leg.; RMCA. – **Maniema** • 1 ♀; Tshuapa: Bamania (Mbandaka); 00°01’ N, 18°19’ E; Nov. 1951; R.P. Hulstaert leg.; RMCA.

#### *Pseudoanthidium* (*Immanthidium*) *sjoestedti* (Friese, 1909), *Immanthidium sjoestedti* (Friese, 1909)

**Material.** Cupboard 110, Box 72 (4♀♀ and 4♂♂). **D.R. CONGO**. – **Haut-Katanga** • 1 ♂; Elisabethville (Lubumbashi); 11°40’ S, 27°28’ E; 25-30 Nov. 1930; R. Massart leg.; RMCA. – **Haut-Lomami** • 1 ♀; PNU Mbuye-Bala; 08°44’ S, 25°00’ E, alt. 1750 m; 24-31 Nov. 1948; Mis. G.F. de Witte leg.; RMCA. – **Ituri** • 1 ♂; Mahagi-Niarembe; 02°15’ N, 31°07’ E; Sep. 1935; Ch. Scops leg.; RMCA. – **Lualaba** • 1 ♀; Tenke; 10°35’ S, 26°06’ E; 30 Jul. – 9 Aug. 1931; T.D. Cockerell leg.; RMCA. – **Nord-Kivu** • 2 ♀♀; Beni; 00°29’ N, 29°28’ E; Aug. 1914; Dr. J. Bequaert leg.; RMCA. • 1 ♂; Rutshuru; 01°11’ S, 29°27’ E; Sep. – Oct. 1936; Dr. Delville leg.; RMCA • 1 ♂; Terr. Rutshuru; 01°11’ S, 29°27’ E; 1937; Miss. Prophylactique leg.; RMCA.

#### *Pseudoanthidium* (*Micranthidium*) *truncatum* (Smith, 1854), *Micranthidium truncatum* (Smith, 1854)

**Material.** Cupboard 110, Box 74 (239♀♀ and 30♂♂). **D.R. CONGO**. – **Bas-Uele** • 1 ♀; Uele: Ibembo; 02°38’ N, 23°36’ E; 1933; H.J. Brédo leg.; RMCA. • 1 ♀; Bambesa; 03°28’ N, 25°43’ E; 1933; R.F. Hutsebaut leg.; RMCA. – **Equateur** • 5 ♀♀; Bikoro; 00°45’ S, 18°07’ E; May 1936; J. Ghesquière leg.; RMCA • 1 ♀; same location as for preceding; May 1954; R.P. Hulstaertleg.; RMCA • 1 ♀; same location as for preceding; 1953; R.P. Lootens leg.; RMCA • 1 ♀; same location as for preceding; Feb. 1952; R.P. Lootens leg.; RMCA • 1 ♀; same location as for preceding; Jan. – Feb. 1952; R.P. Lootens leg.; RMCA • 2♀♀; same location as for preceding; Mar. 1954; R.P. Lootens leg.; RMCA • 1 ♂; same location as for preceding; Jun. 1952; R.P. Lootens leg.; RMCA • 3 ♀♀; same location as for preceding; Jul. 1952; R.P. Lootens leg.; RMCA • 1 ♀; same location as for preceding; Dec. 1951; R.P. Lootens leg.; RMCA • 1 ♀; Coquilhatville (Mbandaka); 00°03’ N, 18°15’ E; 12 Nov. 1922; Dr. M. Bequaert leg.; RMCA • 1 ♀; same location as for preceding; 24 Jun. 1924; Dr. M. Bequaert leg.; RMCA • 1 ♀; same location as for preceding; 1927; Strada leg.; RMCA • 4 ♀♀; Eala (Mbandaka); 00°03’ N, 18°19’ E; 1932; A. Corbisier leg.; RMCA • 3 ♀♀; same location as for preceding; 1933; A. Corbisier leg.; RMCA • 1 ♀; same location as for preceding; Mar. 1935; A. Corbisier leg.; RMCA • 1 ♀; same location as for preceding; Apr. 1933; A. Corbisier leg.; RMCA • 5 ♀♀; same location as for preceding; Jun. 1932; A. Corbisier leg.; RMCA • 5 ♀♀; same location as for preceding; 28 Feb. 1939; Coll. Ghesquière leg.; RMCA • 3 ♂♂; same location as for preceding; 28 Feb. 1939; Coll. Ghesquière leg.; RMCA • 1 ♀; same location as for preceding; 15 Oct. 1936; G. Couteaux leg.; RMCA • 4 ♀♀; same location as for preceding; 15 Nov. 1931; H.J. Brédo leg.; RMCA • 1 ♀; same location as for preceding; 19 Nov. 1931; H.J. Brédo leg.; RMCA • 2 ♀♀; same location as for preceding; 20 Oct. 1931; H.J. Brédo leg.; RMCA • 1 ♀; same location as for preceding; 22 Apr. 1932; H.J. Brédo leg.; RMCA • 1 ♀; same location as for preceding; 22 Nov. 1931; H.J. Brédo leg.; RMCA • 1 ♀; same location as for preceding; 28 Nov. 1931; H.J. Brédo leg.; RMCA • 2 ♀♀; same location as for preceding; 5 Nov. 1931; H.J. Brédo leg.; RMCA • 2 ♀♀; same location as for preceding; Mar. 1932; H.J. Brédo leg.; RMCA • 3 ♀♀; same location as for preceding; Apr. 1932; H.J. Brédo leg.; RMCA • 6 ♀♀; same location as for preceding; May 1932; H.J. Brédo leg.; RMCA • 3 ♀♀; same location as for preceding; Nov. 1931; H.J. Brédo leg.; RMCA • 1 ♀; same location as for preceding; Jan. 1936; J. Ghesquière leg.; RMCA • 2 ♀♀; same location as for preceding; Feb. 1935; H.J. Brédo leg.; RMCA • 1 ♀; same location as for preceding; Apr. 1955; H.J. Brédo leg.; RMCA • 2 ♀♀; same location as for preceding; May 1935; H.J. Brédo leg.; RMCA • 1 ♀; same location as for preceding; May 1936; H.J. Brédo leg.; RMCA • 5 ♀♀; same location as for preceding; Jun. 1935; H.J. Brédo leg.; RMCA • 1 ♀; same location as for preceding; Oct. 1935; H.J. Brédo leg.; RMCA • 1 ♀; same location as for preceding; 18 Apr. 1928; Lt. J. Ghesquière leg.; RMCA • 1♀; same location as for preceding; 26 Feb. 1918; R.P. Hulstaert leg.; RMCA • 3 ♀♀; Equateur; 00°00’ N, 18°14’ E; Verlaine leg.; RMCA • 2 ♀♀; Flandria; 00°20’ S, 19°06’ E; 1931; R.P. Hulstaert leg.; RMCA • 1 ♀; same location as for preceding; 24 Nov. 1931; R.P. Hulstaert leg.; RMCA • 1 ♀; same location as for preceding; Sep. 1931; R.P. Hulstaert leg.; RMCA • 3 ♀♀; Lukolela (Bikoro); 01°03’ S, 17°12’ E; Nov. 1934; Dr. Ledoux leg.; RMCA • 9 ♀♀; Ubangi: Nouvelle Anvers (Bomongo); 00°03’ N, 18°19’ E; 9 Dec. 1952; P. Basilewsky leg.; RMCA • 6 ♂♂; same location as for preceding; 9 Dec. 1952; P. Basilewsky leg.; RMCA • 6 ♀♀; Ubangi: Nouvelle Anvers (Bomongo); 00°03’ N, 18°19’ E; 9-11 Dec. 1952; P. Basilewsky leg.;

RMCA. – **Haut-Katanga** • 1 ♀; Elisabethville (Lubumbashi); 11°40’ S, 27°28’ E; Jun. 1929; Ch. Seydel leg.; RMCA • 1 ♀; same location as for preceding; 16 Août 1931; De Loose leg.; RMCA • 1 ♀; same location as for preceding; 11 Sep. 1932; Dr. M. Bequaert leg.; RMCA • 1 ♀; same location as for preceding; 15 Aug. 1933; Dr. M. Bequaert leg.; RMCA • 1 ♂; same location as for preceding; 20 Sep. 1926; Dr. M. Bequaert leg.; RMCA • 1 ♀; same location as for preceding; 21 Dec. 1930; Dr. M. Bequaert leg.; RMCA • 1 ♀; same location as for preceding; 26 Feb. 1933; Dr. M. Bequaert leg.; RMCA • 1 ♀; same location as for preceding; 5 Sep. 1929; Dr. M. Bequaert leg.; RMCA • 1 ♀; same location as for preceding; Mar. 1926; Dr. M. Bequaert leg.; RMCA • 1 ♀; Elisabethville, riv. Kimilolo; 11°40’ S, 27°28’ E; 14 Jun. 1920; Dr. M. Bequaert leg.; RMCA • 2 ♂♂; same location as for preceding; 19 Jun. 1920; Dr. M. Bequaert leg.; RMCA • 1 ♀; same location as for preceding; 27 May 1920; Dr. M. Bequaert leg.; RMCA • 1 ♀; Jadoville: Numbi; 10°59’ S, 26°44’ E; May 1957; R.P.Th. de Caters leg.; RMCA • 1 ♂; Kambove-Ruwe; 10°52’ S, 26°38’ E; Feb. – Mar. 1907; Dr. Sheffield-Neave leg.; RMCA^7^ • 1 ♀; Kilwa-Lukonzolwa; 08°47’ S, 28°38’ E; Aug. 1907; Dr. Sheffield-Neave leg.; RMCA^8^. – **Haut-Lomami** • 1 ♀; Lualaba: Kalombo; 08°18’ S, 26°19’ E; 29 May 1947; Dr. M. Poll leg.; RMCA • 1 ♀; PNU Kalumengongo; 08°00’ S, 27°05’ E, alt. 1780 m; 21 Jan. 1948; Mis. G.F. de Witte leg.; RMCA • 1 ♀; PNU Mbuye-Bala; 08°44’ S, 25°00’ E, alt. 1750 m; 25-31 Mar. 1948; Mis. G.F. de Witte leg.; RMCA. – **Haut-Uele** • 2 ♀♀; PNG [Parc National de Garamba]; 03°40’ N, 29°00’ E; 29 Nov. 1951; H. De Saeger leg.; RMCA. – **Ituri** • 8 ♀♀; Ituri: Bunia; 01°34’ N, 30°15’ E; 29 Nov. 1951; P. Lefèvre leg.; RMCA. – **Kasaï Central** • 1 ♀; Luluabourg (Kananga); 05°54’ S, 21°52’ E; P. Callewaert leg.; RMCA. – **Kinshasa** • 1 ♀; Kinkole (Stanley Pool); 04°12’ S, 15°33’ E; 2 Aug. 1958; J. Pasteels leg.; RMCA • 1 ♀; Léopoldville (Kinshasa); 04°19’ S, 15°19’ E; 20 Sep. 1923; A. Collart leg.; RMCA • 1 ♀; same location as for preceding; 4 Aug. 1958; J. Pasteels leg.; RMCA. – **Kongo Central** • 1 ♀; Congo da Lemba (Songololo); 05°42’ S, 13°42’ E; 1911; R. Mayné leg.; RMCA • 1 ♀; same location as for preceding; Jan. – Feb. 1913; R. Mayné leg.; RMCA • 1 ♀; Kisantu (Madimba); 05°08’ S, 15°06’ E; Sep. 1920; P. Vanderijst leg.; RMCA • 1 ♀; same location as for preceding; 1911; Rovere leg.; RMCA • 1 ♀; Kitobola; 05°22’ S, 14°31’ E; 1911; Rovere leg.; RMCA • 14 ♀♀; Mayidi (Madimba); 05°11’ S, 15°09’ E; 1945; Rév. P. Van Eyen leg.; RMCA • 1 ♂; same location as for preceding; 1945; Rév. P. Van Eyen leg.;

RMCA. – **Lomami** • 1 ♀; Sankuru: M’Pemba Zeo (Gandajika); 06°49’ S, 23°58’ E; 17 Oct. 1958; Don R. Maréchal leg.; RMCA • 1 ♀; same location as for preceding; 24 Aug. 1959; Don R. Maréchal leg.; RMCA. – **Lualaba** • 1 ♀; Lulua: Kapanga; 08°21’ S, 22°34’ E; 19 Nov. 1932; G.F. Overlaet leg.; RMCA. – **Maï-Ndombe** • 1 ♀; Bokoro (Kutu); 02°50’S, 18°23’ E; 20 Mar. 1915; R. Mayné leg.; RMCA. – **Maniema** • 1 ♀; Kasongo; 04°27’ S, 26°40’ E; Aug. 1959; P.L.G. Benoit leg.; RMCA • 1 ♀; Lokandu; 02°31’ S, 25°47’ E; Mar. 1939; Capt. Marée leg.; RMCA • 1 ♂; Nyangwe; 04°13’ S, 26°11’ E; 1918; R. Mayné leg.; RMCA. – **Mongala** • 1 ♀; Bumba; 02°11’ N, 22°32’ E; Dec. 1939 – Jan. 1940; H. De Saeger leg.; RMCA. – **Nord-Kivu** • 1 ♀; Beni à Lesse; 00°03’ N, 29°41’ E; Sep. 1911; Dr. Martula leg.; RMCA • 1 ♀; Burunga; 01°20’ S, 29°02’ E; 1914; Dr. J. Bequaert leg.; RMCA • 1 ♀; Kinyamahura (Djomba); 01°18’ S, 29°33’ E, alt. 1800 m; 23 Aug. 1934; G.F. de Witte leg.; RMCA • 1 ♀; Lesse; 00°45’ N, 29°46’ E; 21 Jul. 1914; Dr. J. Bequaert leg.; RMCA • 3 ♀♀; Nyabikoro (Rutshuru); 01°11’ S, 29°27’ E; 21 Jul. 1914; K. Baeten leg.; RMCA • 1 ♀; PNA [Parc National Albert]; 00°03’ S, 29°30’ E; 19 Jul. 1954; P. Vanschuytbroeck & H. Synave leg.; RMCA • 1 ♀; same location as for preceding; 13 Apr. 1953; P. Vanschuytbroeck & H. Synave leg.; RMCA • 3 ♀♀; Rutshuru; 01°11’ S, 29°27’ E; Jan. 1938; J. Ghesquière leg.; RMCA • 8 ♀♀; same location as for preceding; Dec. 1937; J. Ghesquière leg.; RMCA • 3 ♂♂; same location as for preceding; Dec. 1937; J. Ghesquière leg.; RMCA • 1 ♀; same location as for preceding; 26 May 1936; L. Lippens leg.; RMCA • 1 ♀; same location as for preceding; 28 May 1936; L. Lippens leg.; RMCA • 1 ♀; Rutshuru (riv. Musugereza, 1100m); 01°11’ S, 29°27’ E; 9 Jul. 1935; G.F. de Witte leg.; RMCA • 1 ♀; Rwankwi; 01°20’ S, 29°22’ E; 3 Mar. 1946; J.V. Leroy leg.; RMCA • 1 ♀; Terr. Rutshuru; 01°11’ S, 29°27’ E; 18 Aug. 1937; Miss. Prophylactique leg.; RMCA. – **Nord-Ubangi** • 3 ♀♀; Abumombazi (Mobayi); 03°34’ N, 22°03’ E; 18-26 Feb. 1932; H.J. Brédo leg.; RMCA • 1 ♀; same location as for preceding; 8 – 11 Jan. 1932; H.J. Brédo leg.; RMCA • 2 ♀♀; Yakoma; 04°06’ N, 22°23’ E; 5-17 Feb. 1932; H.J. Brédo leg.;

RMCA. – **Sud-Kivu** • 1 ♀; Kalonge, Riv. Mushuva; 00°21’ N, 29°49’ E; 6 Jun. 1949; G. Marlier leg.; RMCA • 1 ♀; Mulungu; 02°20’ S, 28°47’ E; 1938; Hendrickx leg.; RMCA • 1 ♀; same location as for preceding; May 1954; J. Decelle leg.; RMCA • 1 ♀; Mulungu: Tshibinda; 02°19’ S, 28°45’ E; Nov. 1951; P.C. Lefèvre leg.; RMCA • 1 ♀; Région des lacs; 02°29’ S, 28°51’ E; Dr. Sagona leg.; RMCA. – **Tanganyika** • 1 ♀; Kiambi (Manono); 07°19’ S, 28°01’ E; 19 Feb. 1911; Dr. Valdonio leg.; RMCA • 1 ♀; Tang. Moero: Nyunzu; 05°57’ S, 28°01’ E; 1935; H. De Saeger leg.; RMCA. – **Tshopo** • 1♀; Mfungwe-Kayumbe; 01°17’ S, 26°16’ E; Jun. 1907; Dr. Sheffield-Neave leg.; RMCA^9^ • 2 ♂♂; same location as for preceding; Jun. 1907; Dr. Sheffield-Neave leg.; RMCA^10^ • 2 ♀♀; Miss. St. Gabriel; 00°33’ N, 25°05’ E; M. Torley leg.; RMCA • 1 ♀; Stanleyville (Kisangani); 00°31’ N, 25°11’ E; 26 Sep. – 17 Oct. 1928; A. Collart leg.; RMCA • 1 ♀; same location as for preceding; 7 Jan. 1930; A. Collart leg.; RMCA • 1♀; same location as for preceding; Nov. 1925; J. Ghesquière leg.; RMCA • 1 ♀; Yangambi; 00°46’ N, 24°27’ E; Jan. 1960; J. Decelle leg.; RMCA • 1 ♀; same location as for preceding; 15 Jun. 1948; P.L.G. Benoit leg.; RMCA. – **Tshuapa** • 1 ♀; Bokuma; 00°06’ S, 18°41’ E; Feb. – Apr. 1941; R.P. Hulstaert leg.; RMCA • 1 ♀; Tshuapa: Bamanya; 00°01’ N, 18°19’ E; Aug. 1963; R.P. Hulstaert leg.; RMCA • 1 ♀; same location as for preceding; Oct. 1961; R.P. Hulstaert leg.; RMCA • 1 ♂; same location as for preceding; Oct. 1961; R.P. Hulstaert leg.; RMCA • 1 ♀; same location as for preceding; Dec. 1952; R.P. Hulstaert leg.; RMCA • 1 ♀; same location as for preceding; 1968; Rév. P. Hulstaert leg.; RMCA • 1 ♀; Tshuapa: Bokuma; 00°06’ S, 18°41’ E; Apr. 1955; R.P. Hulstaert leg.; RMCA • 2 ♀♀; same location as for preceding; 1953; R.P. Lootens leg.; RMCA • 6 ♀♀; same location as for preceding; Mar. 1954; R.P. Lootens leg.; RMCA • 2 ♂♂; same location as for preceding; Mar. 1954; R.P. Lootens leg.; RMCA • 1 ♀; same location as for preceding; Sep. 1954; R.P. Lootens leg.; RMCA • 11 ♀♀; same location as for preceding; Jun. 1952; R.P. Lootens leg.; RMCA • 1 ♀; Tshuapa: Flandria; 00°20’ S, 19°06’ E; 1946; Rév. P. Hulstaert leg.; RMCA • 1 ♀; Tshuapa: Flandria-Bolima; 00°03’ N, 19°23’ E; Oct. 1941; Rév. P. Hulstaert leg.; RMCA • 1 ♀; Tshuapa: Iyonda; 00°17’ S, 20°52’ E; 1953; R.P. Michielsen t leg.; RMCA.

#### *Serapista denticulata* (Smith, 1854)

**Material.** Cupboard 110, Box 75 (22♀♀ and 23♂♂). **D.R. CONGO**. – **Equateur** • 1 ♀; Eala (Mbandaka); 00°03’ N, 18°19’ E; 22 Apr. 1932; H.J. Brédo leg.; RMCA • 1 ♂; same location as for preceding; 22 Apr. 1932; H.J. Brédo leg.; RMCA. – **Haut-Katanga** • 1 ♂; Elisabethville (Lubumbashi); 11°40’ S, 27°28’ E; 12 Nov. 1933; Ch. Seydel leg.; RMCA • 2 ♀♀; same location as for preceding; Oct. 1935; Ch. Seydel leg.; RMCA • 1 ♀; same location as for preceding; Jun. 1932, De Loose leg.; RMCA • 1 ♀; same location as for preceding; 21 Aug. 1930; Dr. M. Bequaert leg.; RMCA • 4 ♂♂; Elisabethville: Munama; 11°40’ S, 27°28’ E; Aug. 1931; G.F. de Witte leg.; RMCA • 1 ♀; Kabunda; 12°25’ S, 29°22’ E; Dec. 1928; Ch. Seydel leg.; RMCA • 1 ♂; Lubumbashi; 11°40’ S, 27°28’ E; 8 Apr. 1921; Dr. M. Bequaert leg.; RMCA • 3 ♂♂; Munama; 11°40’ S, 27°28’ E; 26 Jul. 1926; Ch. Seydel leg.; RMCA. – **Ituri** • 6 ♀♀; Ituri: Logo; 02°11’ N, 30°56’ E; 15 Nov. 1926, R.P. Thalman leg.; RMCA • 2 ♂♂; same location as for preceding; 15 Nov. 1926, R.P. Thalman leg.; RMCA • 1 ♀; Mahagi-Niarembe; 02°15’ N, 31°07’ E; Nov. 1935, M. & Mme. Ch. Scops leg.; RMCA • 1 ♂; same location as for preceding; Nov. 1935, M. & Mme. Ch. Scops leg.; RMCA. – **Lualaba** • 1 ♂; Ditanto; 10°15’ S, 25°53’ E; Oct. 1925; Ch. Seydel leg.; RMCA • 2 ♀♀; Lulua: Kapanga; 08°21’ S, 22°34’ E; Nov. 1932; G.F. Overlaet leg.; RMCA. – **Nord-Kivu** • 1 ♂; Buseregenye (Rutshuru); 01°11’ S, 29°27’ E; Sep. 1929, Ed. Luja leg.; RMCA • 1 ♀; Mukule; 01°20’ S, 29°15’ E; Sep. 1914; Dr. J. Bequaert leg.; RMCA • 3 ♀♀; Ngesho; 01°17’ S, 29°06’ E; Sep. 1937; J. Ghesquière leg.; RMCA • 2 ♂♂; same location as for preceding; Sep. 1937; J. Ghesquière leg.; RMCA • 1 ♂; PNA [Parc National Albert]; 00°03’ S, 29°30’ E; 16 Mar. 1954, P. Vanschuytbroeck & H. Synave leg.; RMCA • 1 ♂; Rutshuru; 01°11’ S, 29°27’ E; Aug. 1937; J. Ghesquière leg.; RMCA. – **Sud-Kivu**• 1 ♀; Ibanda; 02°29’ S, 28°51’ E; 1952, M. Vandelannoite leg.; RMCA • 1 ♂; Mulungu: Tshibinda; 02°19’ S, 28°45’ E; Nov. 1951, P.C. Lefèvre leg.; RMCA • 1 ♀; Mushoko; 02°29’ S, 28°51’ E; May 1937; J. Ghesquière leg.; RMCA • 1 ♂; same location as for preceding; May 1937; J. Ghesquière leg.; RMCA • 1 ♀; same location as for preceding; Jun. 1937; J. Ghesquière leg.; RMCA • 1 ♀; same location as for preceding; Aug. 1937; J. Ghesquière leg.; RMCA • 1♀; Mushoko (Lac); 02°29’ S, 28°51’ E; Aug. 1937; J. Ghesquière leg.; RMCA.

#### *Heriades (Heriades) bequaerti* Cockerell, 1931, *Heriades bequaerti* Ckll. Schlett

**Material.** Cupboard 110, Box 77 (1♀). **D.R. CONGO**. Holotype – **Nord-Kaivu** • 1 ♀; Masisi; 01°24’ S, 28°49’ E; 30 Dec. 1914; Dr. J. Bequaert leg.; RMCA.

#### *Heriades frontosus* Schletterer, 1989, *Heriades frondosus* [sic] Schlett

**Material.** Cupboard 110, Box 77 (4♀♀). **D.R. CONGO**. – **Equateur** • 1 ♀; Coquilhatville (Mbandaka); 00°03’ N, 18°15’ E; Oct. 1922; Dr. M. Bequaert leg.; RMCA • 1 ♀ ; same location as for preceding; ; 1946 ; Ch. Scops leg. ; RMCA • 1 ♀ ; Eala (Mbandaka) ; 00°03’ N, 18°19’ E ; 20 Oct. 1931 ; H.J. Brédo leg. ; RMCA. – **Mongala** • 1 ♀ ; Ubangi : Burubu ; 02°09’ N, 21°30’ E ; 6 Dec. 1929 ; H.J. Brédo leg. ; RMCA.

#### *Heriades hercule* Strand, 1911, *Heriades herculus* Strd. Schlett

**Material.** Cupboard 110, Box 77 (1♀). **D.R. CONGO**. – **Nord-Kivu** • 1 ♀; Mt. Ruwenzori; 00°26’ S, 29°50’ E, alt. 2300 m; 19 Apr. 1914; Dr. J. Bequaert leg.; RMCA.

#### *Heriades impressus* Schletterer, 1889, *Heriades impressus* Schlett

**Material.** Cupboard 110, Box 77 (1♀). **D.R. CONGO**. – **Nord-Kivu** • 1 ♀; Mukule; 01°20’ S, 29°15’ E; Sep. 1914; Dr. J. Bequaert leg.; RMCA.

#### *Heriades nitescens* Cockerell, 1931, *Heriades nitescens* Ckll. Schlett

**Material.** Cupboard 110, Box 77 (1♀). **D.R. CONGO**. Holotype – **Nord-Kivu** • 1 ♀; Kasindi-Beni; 00°29’ N, 29°28’ E; Aug. 1914; Dr. J. Bequaert leg.; RMCA.

#### *Heriades perminutus* Cockerell, 1935, *Heriades perminuta* Ckll. Schlett

**Material.** Cupboard 110, Box 77 (1♂). **D.R. CONGO**. Paratypes – **Sankuru** • 1 ♂; N’Kole, Makarikari; 03°27’ S, 22°26’ E; 6-23 Aug. 1930; T.D. Cockerell leg.; RMCA.

#### *Heriades spiniscutis* (Cameron, 1905)

The name *Heriades spiniscutis* was given as the oldest synonym of 16 other species names in the genus *Heriades* (Michener 1968), including also *Heriades communis*, which Pasteels has retained as separate species, preserved in the RMCA collections.

#### *Heriades spiniscutis* (Cameron, 1905)

**Material.** Cupboard 110, Boxes 76 and 77 (31♀♀ and 26♂♂). **D.R. CONGO**. – **Bas-Uele** • 1 ♀; Bambesa; 03°28’ N, 25°43’ E; 16 May 1938; P. Henrard leg.; RMCA. – **Equateur** • 1 ♀; Eala (Mbandaka); 00°03’ N, 18°19’ E; 20 Oct. 1931; H.J. Brédo leg.; RMCA • 1 ♂; same location as for preceding; Nov. 1936; J. Ghesquière leg.; RMCA. – **Haut-Katanga** • 1 ♀; Elisabethville (Lubumbashi); 11°40’ S, 27°28’ E; Aug. 1923; Dr. M. Bequaert leg.; RMCA • 1 ♀; Lubumbashi; 11°40’ S, 27°28’ E; 16 Jan. 1921; Dr. M. Bequaert leg.; RMCA • 1 ♂; Kasenga; 10°22’ S, 28°38’ E; 1 Feb. 1912; Dr. J. Bequaert leg.; RMCA • 1 ♀; Mufunga-Sampwe; 09°21’ S, 27°27’ E; 1 – 16. Dec. 1911; Dr. J. Bequaert leg.; RMCA. – **Haut-Lomami** • 1 ♀; Bukama; 09°12’ S, 25°51’ E; 29 May 1911; Dr. J. Bequaert leg.; RMCA • 1 ♂; Lomami: Kaniama; 07°31’ S, 24°11’ E; 1930; R. Massart leg.; RMCA •1 ♀; Sankisia; 09°24’ S, 25°48’ E; 4 Oct. 1911; Dr. J. Bequaert leg.; RMCA. – **Haut-Uele** • 1 ♂; Haut-Uele: Mauda; 04°46’ N, 27°18’ E; Mar. 1925; Dr. J. Schouteden leg.; RMCA • 1 ♀; Haut-Uele: Paulis; 02°46’ N, 27°38’ E; Jun. 1947; P.L.G. Benoit leg.; RMCA • 3 ♂♂; same location as for preceding; Jun. 1947; P.L.G. Benoit leg.; RMCA. – **Ituri** • 1 ♀; Ituri: Bunia; 01°34’ N, 30°15’ E; Jun. 1938; P. Lefèvre leg.; RMCA • 1 ♀; Ituri: Nduye-Makara; 01°50’ N, 28°59’ E; Sep. – Sep. 1921; A. Pilette leg.; RMCA • 1 ♀; Kibimbi; 01°22’ N, 28°55’ E; 3 Feb. 1911; Dr. J. Bequaert leg.; RMCA. – **Kasaï** • 1 ♀; Luebo; 05°20’ S, 21°24’ E; Aug. 1921; Lt. J. Ghesquière leg.; RMCA. – **Kasaï Central** • 1 ♀; Luluabourg (Kananga); 05°54’ S, 21°52’ E; 14-17 May 1919; P. Callewaert leg.; RMCA. – **Kinshasa** • 2 ♀♀; Léopoldville (Kinshasa); 04°19’ S, 15°19’ E; 15 Sep. 1910; Dr. J. Bequaert leg.; RMCA • 1 ♂; same location as for preceding; 1910; Dr. J. Bequaert leg.; RMCA • 1 ♀; same location as for preceding; 16 Sep. 1910; Dr. J. Bequaert leg.; RMCA • 1 ♂; same location as for preceding; 18 Sep. 1910; Dr. J. Bequaert leg.; RMCA • 1 ♀; same location as for preceding; 25 Sep. 1910; Dr. J. Bequaert leg.; RMCA • 1 ♀; same location as for preceding; Aug. 1949; R. Dubois leg.; RMCA. – **Kongo Central** • 2 ♀♀; Congo da Lemba (Songololo); 05°42’ S, 13°42’ E; 1913; R. Mayné leg.; RMCA • 3 ♂♂; same location as for preceding; Jan. – Feb. 1913; R. Mayné leg.; RMCA • 1 ♀; same location as for preceding; Jan. – Apr. 1913; R. Maynéleg.; RMCA • 1 ♀; Kisantu (Madimba); 05°08’ S, 15°06’ E; 1927; R.P. Vanderyst leg.; RMCA • 2 ♀♀; same location as for preceding; 1931; R.P. Vanderyst leg.; RMCA. – **Maï-Ndombe** • 2 ♂♂; Wombali (Mushie); 03°16’ S, 17°22’ E; Jul. 1913; P. Vanderijst leg.; RMCA. – **Maniema** • 2 ♀♀; Kibombo; 03°54’ S, 25°55’ E; 1 Nov. 1910; Dr. J. Bequaert leg.; RMCA • 1 ♂; Maniema; 05°00’ S, 29°00’ E; Jul. 1913; R. Mayné leg.; RMCA • 1 ♂; Nyangwe; 04°13’ S, 26°11’ E; 12 Nov. 1910; Dr. J. Bequaert leg.; RMCA • 1 ♀; same location as for preceding; Apr. 1918; R. Mayné leg.; RMCA. – **Nord-Kivu** • 4 ♂♂; Rutshuru; 01°11’ S, 29°27’ E; Sep. –Oct. 1936; Dr. Delville leg.; RMCA • 2 ♂♂; same location as for preceding; 17 May 1936; L. Lippens leg.; RMCA • 1 ♂; same location as for preceding; 01°11’ S, 29°27’ E; 13 Aug. 1937; Miss. Prophylactique leg.; RMCA • 2 ♂♂; same location as for preceding; 2 Jul. 1937; Miss. Prophylactique leg.; RMCA. – **Sud-Ubangi** • 1 ♀; Ubangi: Nzali; 03°15’ N, 19°47’ E; 3-4 Feb. 1932; H.J. Brédo leg.; RMCA. – **Tanganyika** • 1 ♀; Baudouinville (Moba); 07°02’ S, 29°47’ E; 19 Jan. 1933; L. Burgeon leg.; RMCA. – **Tshopo** • 1 ♀; Stanleyville (Kisangani); 00°31’ N, 25°11’ E; Mar. 1926; Lt. J. Ghesquière leg.; RMCA • 1 ♀; same location as for preceding; Jun. 1926; Lt. J. Ghesquière leg.; RMCA.

#### *Heriades communis* Cockerell, 1931^11^

**Material.** Cupboard 110, Box 77 (4♀♀ and 8♂♂). **D.R. CONGO**. Paratypes: – **Haut-Katanga** • 1 ♀; Elisabethville (Lubumbashi); 11°40’ S, 27°28’ E; 13 May 1920; Dr. M. Bequaert leg.; RMCA • 1 ♂; Elisabethville; riv. Kimilolo; 11°40’ S, 27°28’ E; 6 Nov. 1920; Dr. M. Bequaert leg.; RMCA • 1 ♀; Lubumbashi; 11°40’ S, 27°28’ E; 1 Jan. 1921; Dr. M. Bequaert leg.; RMCA •1 ♂; same location as for preceding; 23 Dec. 1920; Dr. M. Bequaert leg.; RMCA •1 ♂; same location as for preceding; 3 Jun. 1920; Dr. M. Bequaert leg.; RMCA •1 ♂; same location as for preceding; 31 Jan. 1921; Dr. M. Bequaert leg.; RMCA •1 ♂; same location as for preceding; Feb.-Apr. 1921; Dr. M. Bequaert leg.; RMCA. – **Ituri** • 1 ♀; Irumu: Lesse; 01°27’ N, 29°52’ E; 21 Jul. 1914; Dr. J. Bequaert leg.; RMCA • 1 ♀; Irumu: Penge; 01°27’ N, 29°52’ E; 1 Mar. 1914; Dr. J. Bequaert leg.; RMCA.

– **Haut-Katanga** • 1 ♀; Sakania; 12°45’ S, 28°33’ E; Sep. 1931, J. Ogilvie leg.; RMCA. – **Sud-Kivu** • 1 ♂; Uvira; 03°25’ S, 29°08’ E; 29 Aug. 1931; T.D. Cockerell leg.; RMCA. – **Tanganyika** • 1 ♂; Albertville (Kalemie); 05°57’ S, 29°12’ E; 1 Sep. 1931; T.D. Cockerell leg.; RMCA.

#### *Heriades sulcatulus* Cockerell, 1931, *Heriades sulcatulus* Ckll. Schlett

**Material.** Cupboard 110, Box 77 (4♀). **D.R. CONGO**. Holotype – **Ituri** • 1 ♀; Ituri: Lesse; 00°45’ N, 29°46’ E; 21 Jun. 1914; Dr. J. Bequaert leg.; RMCA.

– **Ituri** • 1 ♀; Ituri: La Moto (Madyu); 01°34’ N, 30°14’ E; L. Burgeon leg.; RMCA. – **Sankuru** • 1 ♀; Sankuru: Komi; 03°23’ S, 23°46’ E; 17 Mar. 1930; J. Ghesquière leg.; RMCA. – **Tshopo** • 1 ♀; Terr. Stan. -Epulu (Apayo); 01°23’ N, 28°36’ E; 4 Dec. 1939; Dr. Van Breuseghem leg.; RMCA.

#### *Heriadopsis striatulus* Cockerell, 1931

**Material.** Cupboard 110, Box 45A (1♀ and 22♂♂). **D.R. CONGO**. Holotype – **Haut-Katanga** • 1 ♂; Elisabethville (Riv.Kimololo); 11°40’ S, 27°28’ E; 6 Nov. 1920; Dr. M. Bequaert leg.; RMCA. Paratype – **Haut-Katanga** • 1 ♂; Elisabethville (Riv.Kimololo); 11°40’ S, 27°28’ E; 6 Nov. 1920; Dr. M. Bequaert leg.; RMCA.

– **Haut-Katanga** • 1 ♂; Elisabethville (Lubumbashi); 11°40’ S, 27°28’ E; 1931; De Loose leg.; RMCA • 1 ♂; same location as for preceding; 1932; De Loose leg.; RMCA • 2 ♂♂; Elisabethville (Riv.Kimololo); 11°40’ S, 27°28’ E; 6 Nov. 1920; Dr. M. Bequaert leg.; RMCA. • 2 ♂♂; same location as for preceding; Nov. 1928; Ch. Seydel leg.; RMCA • 2 ♂♂; Kapema; 10°42’ S, 28°40’ E; Sep. 1924; Ch. Seydel leg.; RMCA. – **Haut-Lomami** • 2 ♂♂; Sankisia; 09°24’ S, 25°48’ E; 10 Aug. 1911; Dr. J. Bequaert leg.; RMCA • 1 ♂; same location as for preceding; 30 Aug. 1911; Dr. J. Bequaert leg.; RMCA • 1 ♀; same location as for preceding; 4 Sep. 1911; Dr. J. Bequaert leg.; RMCA • 9 ♂♂; same location as for preceding; 4 Sep. 1911; Dr. J. Bequaert leg.; RMCA.

#### *Osmia reginae* Cockerell, 1932

**Material.** Cupboard 110, Box 77 (1♀). **D.R. CONGO**. – **Haut-Katanga** • 1 ♀; Elisabethville (Lubumbashi); 11°40’ S, 27°28’ E; 11-17 Sep. 1911; J. Ogilvie leg.; RMCA.

#### Lost or loaned materials

The following species, recorded from the RMCA by the cited species authorities, are believed to be lost as they were no longer in the collections at the time of our study. The type specimens of two other species supposedly curated at the RMCA were also absent from the collections. These are: *Coelioxys ateneata* Strand, 1920 and *Coelioxys neavei* Vachal, 1910.

We resume these omissions below in order to complete the list of Megachilidae taxa curated at RMCA.

#### *Coelioxys ateneata* Strand, 1920^12^

**D.R. CONGO**. Holotype – **Kwango** • 1 ♀; Kwango: Atene; 05°23’ S, 19°24’ E; Charliers leg.; RMCA.

#### *Coelioxys aspericauda* Cockerell, 1935 (= *Coelioxys congoensis* Friese, 1922)

**D.R. CONGO**. – **Kwango** • 1 ♀; Kwango; 08°00’ S, 20°00’ E; 1925; Vanderijst leg.; RMCA^13^.

#### *Coelioxys neavei* Vachal, 1910^14^

**D.R. CONGO**. Holotype – **Lualaba** • 1 ♀; Bunkeya; 10°24’ S, 26°58’ E; Oct. 1907; Dr. Sheffield-Neave leg.; RMCA.

#### *Coelioxys piliventris* Vachal, 1910 (= *Coelioxys planidens* Friese, 1904)

**D.R. CONGO**. – **Lualaba** • 1 ♂; Kambove-Ruwe; 10°52’ S, 26°38’ E; 27. Nov. 1907; Dr. Sheffield-Neave leg.; RMCA^15^.

#### *Coelioxys acuticauda* Cockerell, 1935 (= *Coelioxys planidens* Friese, 1904)

**D.R. CONGO**. – **Haut-Katanga** • 1 ♀; Elisabethville (Lubumbashi); 11°40’ S, 27°28’ E; 27 Apr. 1930; Dr. M. Bequaert leg.; RMCA^16^.

#### *Coelioxys rotundicauda* Cockerell, 1935 (= *Coelioxys umbripennis* Friese, 1922)

**D.R. CONGO**. – **Haut-Katanga** • 1 ♀; La Kasepa (Lubumbashi); 11°40’ S, 27°28’ E; 23 Sep. 1923; Ch. Seydel leg.; RMCA^17^.

#### *Megachile flexa* Vachal, 1910 (= *Megachile (Pseudomegachile) schulthessi* Friese, 1903)

**D.R. CONGO**. – **Lualaba** • 1 ♀; Bunkeya; 10°24’ S, 26°58’ E; Oct. 1907; Dr. Sheffield-Neave leg.; RMCA^18^ • 1 ♀; Bunkeya-Lukafu; 10°24’ S, 26°58’ E; Oct. 1907; Dr. Sheffield-Neave leg.; RMCA^19^.

#### *Megachile corneipalmis* Vachal, 1910 (= *Megachile (Anodonteutricharaea) ungulata* Smith, 1853)

**D.R. CONGO**.– **Lualaba** • 1 ♂; Kambove-Bunkeya; 10°24’ S, 26°58’ E; Dr. Sheffield-Neave leg.; RMCA^20^.

### Spatial patterns of Megachilidae species and specimen records

We recorded and digitized the data associated with a total of 6,490 specimens relevant to 195 wild bee species grouped in 18 genera within the family Megachilidae curated at RMCA. All specimens examined were collected between 1905 and 1978.

The species and specimen count per administrative unit are summarized in Tables 1 and 2 below.

**Table 1.** Distribution of species richness, specimen abundance and their ratios relative to the administrative entities of the DRC. **a)** The former provinces (up to 1960) include: Equateur (Equateur, Mongala, Nord-Ubangi, Sud-Ubangi, Tshuapa), Kasaï (Kasaï, Kasaï Central, Kasaï Oriental, Lomami, Sankuru), Katanga (Haut-Katanga, Haut-Lomami, Lualaba, Tanganyika), Kivu (Maniema, North Kivu, South Kivu), Leopoldville (Kinshasa, Kongo Central, Kwango, Kwilu, Maï-Ndombe) and Province Orientale (Bas-Uele, Haut-Uele, Ituri, Tshopo); **b)**26 Present-day provinces.

| Administrative units |  | N species | N specimens | N species/N specimens (%) |
| --- | --- | --- | --- | --- |
| **a) Historical Provinces (n=6)** |  |  |  |  |
| 1 | Kivu | 82 | 443 | 18.5 |
| 2 | Kasai | 52 | 297 | 17.5 |
| 3 | Orientale | 88 | 815 | 10.7 |
| 4 | Katanga | 147 | 1,801 | 8.1 |
| 5 | Léopoldville | 47 | 891 | 5.2 |
| 6 | Equateur | 55 | 2,243 | 2.4 |
| **b) Present-day provinces (n = 26)** |  |  |  |  |
| 1 | Kasai Oriental | 3 | 7 | 42.8 |
| 2 | Mai-Ndombe | 20 | 47 | 42.5 |
| 3 | Kasai | 20 | 50 | 40.0 |
| 4 | Mongala | 15 | 41 | 36.5 |
| 5 | Sud-Kivu | 42 | 116 | 36.2 |
| 6 | Kwango | 6 | 17 | 35.2 |
| 7 | Kwilu | 13 | 38 | 34.2 |
| 8 | Kasai Central | 26 | 77 | 33.7 |
| 9 | Lomami | 27 | 87 | 31.0 |
| 10 | Maniema | 43 | 143 | 30.0 |
| 11 | Haut-Uele | 47 | 155 | 30.3 |
| 12 | Sankuru | 22 | 76 | 28.9 |
| 13 | Tshopo | 30 | 108 | 27.7 |
| 14 | Tanganyika | 50 | 191 | 26.2 |
| 15 | Ituri | 53 | 220 | 24.1 |
| 16 | Haut-Lomami | 79 | 355 | 22.3 |
| 17 | Nord-Kivu | 41 | 184 | 21.9 |
| 18 | Nord-Ubangi | 17 | 97 | 17.5 |
| 19 | Sud-Ubangi | 16 | 95 | 16.8 |
| 20 | Haut-Katanga | 113 | 768 | 14.7 |
| 21 | Kinshasa | 20 | 152 | 13.2 |
| 22 | Lualaba | 64 | 487 | 13.1 |
| 23 | Bas-Uele | 38 | 332 | 11.4 |
| 24 | Tshuapa | 29 | 361 | 8.0 |
| 25 | Kongo Central | 38 | 637 | 5.9 |
| 26 | Equateur | 46 | 1,649 | 2.7 |

**Table 2.** Distribution of species richness, specimen abundance and their ratios relative to the phytogeographical entities of the DRC. The Coastal district shows more diversity within its small amount of collected bee specimens.

| Phytogeographical districts |  | N species | N specimens | N species/N specimens (%) |
| --- | --- | --- | --- | --- |
| I | Coastal | 9 | 17 | 52.9 |
| IX | Lakes Edouard and Kivu | 135 | 291 | 46.4 |
| VIII | Lake Albert | 36 | 104 | 34.6 |
| IV | Kasaï | 88 | 284 | 30.9 |
| VII | Ubangi-Uele | 47 | 153 | 30.7 |
| II | Mayumbe | 6 | 22 | 27.2 |
| V | Low-Katanga | 135 | 513 | 26.3 |
| III | Low-Congo | 42 | 750 | 5.6 |
| VI | Central Forestry | 156 | 2,938 | 5.3 |
| X | Upper-Katanga | 63 | 1,418 | 4.4 |

The administrative entities of the colonial era that hold more specimens are not the most diverse in terms of species sampled. The ratio between species richness and the number of specimen records is higher for the historical Kivu province (18.5% of its 443 specimens). These proportions are also observed in Kasaï, where 17.5% of species were found represented in the collections by 297 specimens. The Province Orientale is ranked third with a ratio of 10.7% (88 species and 815 specimens), followed by the Katanga Province (8.1%), Leopoldville (5.2%) and the Equateur which has a relatively low species exploration rate as its ratio is only 2.4%.

Following the present-day provinces, the lowest species richness is observed in the Kasaï Oriental Province. This is the smallest of the provinces where averages of only 3 species and 7 specimens have been recorded. A similar observation is made in the apparently less explored Kwango province which encompasses 6 species out of only 17 historical specimens.

The Coastal phytogeographical district is relatively richer in species. This species richness seems to be the highest in proportion to the number of records, as 9 species are separated out of 17 specimens. The Coastal (I) and Lakes Edouard and Kivu (IX) districts thus show a ratio of more than 40%. The phytogeographical districts of Mayumbe (II), Kasaï (IV), Low-Katanga (V) and Lake Albert (VIII) are between 20 and 40 % of these ratios. The 3 other districts (III, VI and X) have fewer species in all the related records, their ratio is less than 10%.

All of these data are illustrated on the maps of Figure 1 below.

**Figure 1.**
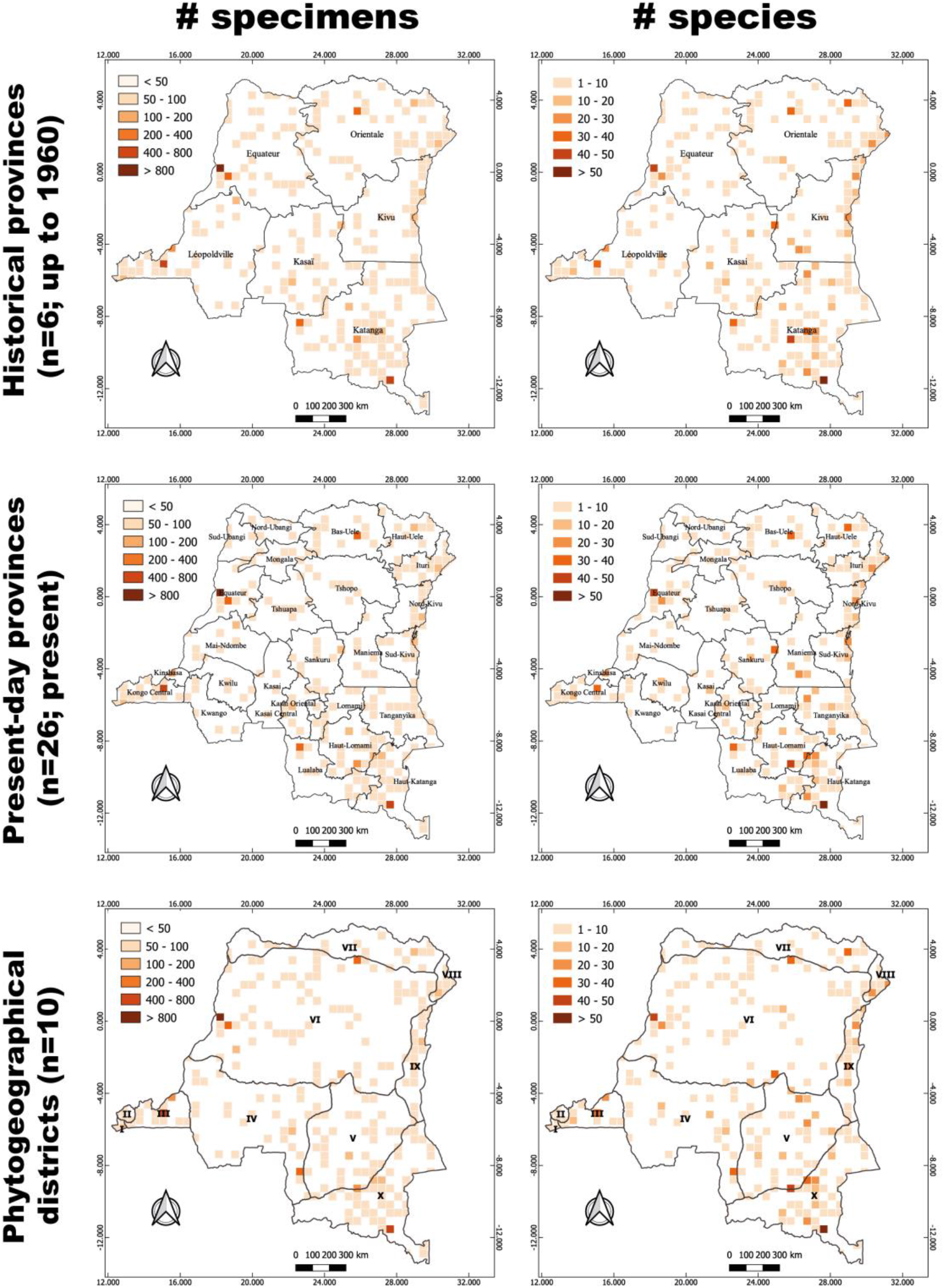
Map of all Megachilidae records illustrating the number of specimens and species for each 50 km x 50 km grid cell in (i) the six historical provinces in DRC (top row), (ii) the 26 present-day provinces (middle row), and (iii) the 10 known phytogeographical districts (bottom row) (I = Coastal; II = Mayumbe; III = Low-Congo; IV = Kasaï; V = Low-Katanga; VI = Central Forestry; VII = Ubangi-Uele; VIII = Lake Albert; IX = Lakes Edouard and Kivu; X = Upper-Katanga) defined within the country. See methods section for details on the administrative units.

The number of historical collections and their associated species richness varies according to the administrative/natural units. An average of 408.3 ± 305.3 specimens for 78.5 ± 37.5 species is counted between the 6 historical provinces, while 186.3 ± 195.4 specimens for 35.3 ± 23.9 species are obtained by following the current provinces. Between phytogeographic districts, an average of 213.8 ± 251.4 specimens corresponding to 71.7 ± 54.2 species is recorded. The linear correlation, which reflects the gain in species richness for more collection abundance (Figure 2), describes a notably significant model between the 26 current DRC provinces (R-squared = 27%, p < 0.001) and the 10 phytogeographic districts (R-square = 38%, p < 0.05). A non-significant model was derived from the correlation based on the 6 former provinces (R-squared = 8%, p > 0.05). The low variance of these models explains the outlier deviations of some units from the mean values. Thus, the greater collection abundance in Province Equateur ultimately reveals a species richness close to the average among the 6 historical provinces; just as the triple number of species in Katanga was collected with half as many specimens as in Province Orientale. Although less significant, the linear pattern between phytogeographic districts also explains the double species recorded in the large Central Forestry district (VI), on five more specimens than in the Kasaï district (IV).

**Figure 2.**
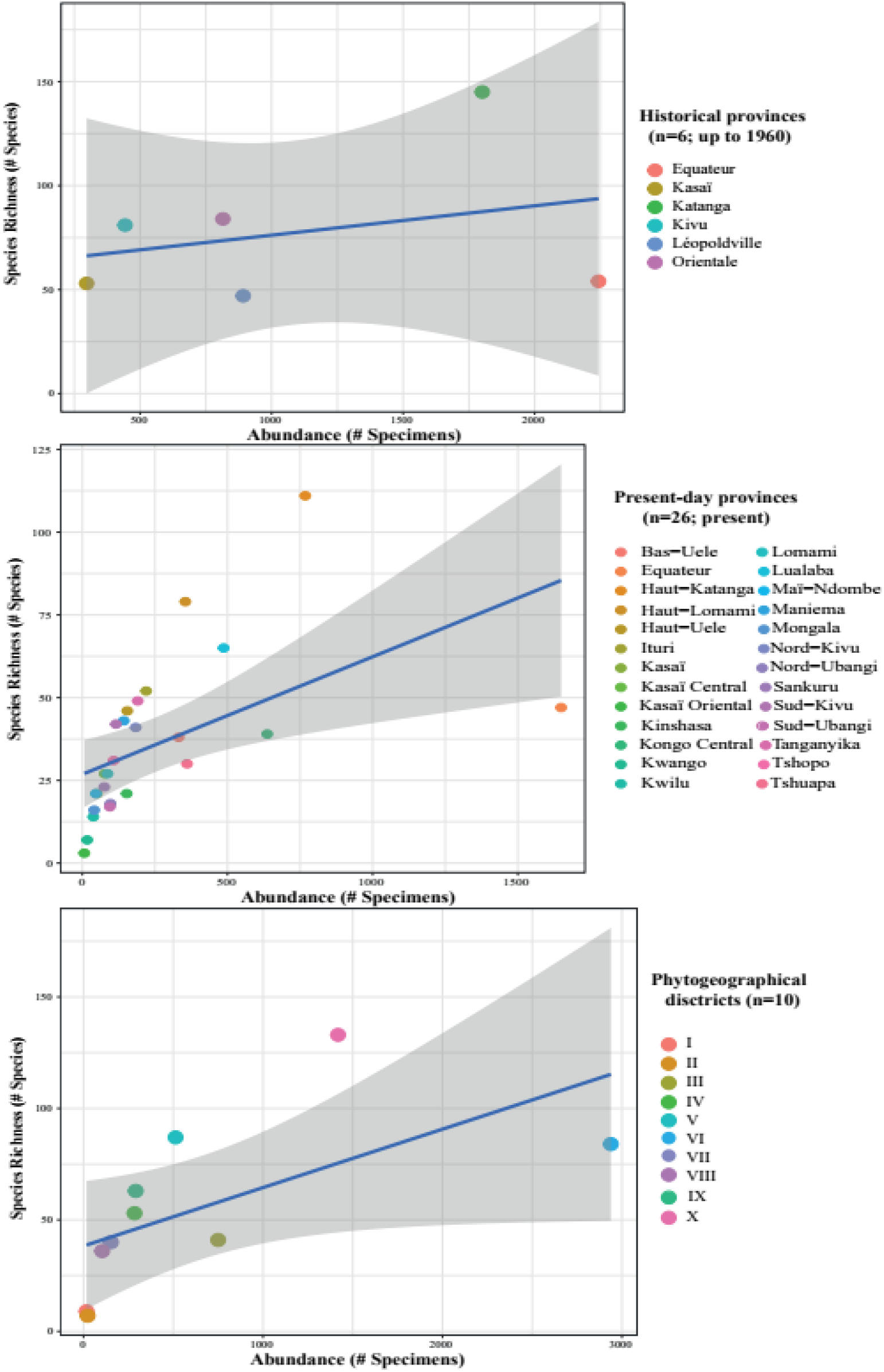
Linear models between collection abundance and associated species richness in (i) the six historical provinces in DRC (top row), (ii) the 26 present-day provinces (middle row), and (iii) the 10 known phytogeographical districts (bottom row) (I = Coastal; II = Mayumbe; III = Low-Congo; IV = Kasaï; V = Low-Katanga; VI = Central Forestry; VII = Ubangi-Uele; VIII = Lake Albert; IX = Lakes Edouard and Kivu; X = Upper-Katanga) defined within the country. Some administrative/natural units deviate from the overall attributable values. The Province Equaeur deviates from the number of collections for species richness close to the mean.

The ANOSIM analysis of community composition among the historical six provinces delimited during the colonial period did not show significant differences in taxonomic composition of communities (ANOSIM: 0.06, p= 0.15). However, this composition of historically collected bees in the DRC shows a strong dissimilarity within the boundaries of the present-day 26 provinces (ANOSIM: 0.05, p < 0.001). Taking phytogeography into account, we found an overall similarity in Megachilidae communities sampled across the known phytogeographical districts at the national scale (ANOSIM: 0.05, p = 0.2).

Our analyses of community turnover among administrative units confirm the results above with congruent results between the taxonomic and phylogenetic beta diversity approaches by showing that the total turnover (βsør) observed among the six historical provinces of DRC and the phytogeographical districts is primarily due to the nestedness of communities (βsne), with comparatively little replacement (βsim) (Table 3). A more balanced mix of turnover was found when repeating the analysis using the 26 contemporary provinces where the higher taxonomic turnover (βsør = 0.74) is due almost as much to community nestedness as to replacement. This implies more heterogeneity in the known fauna of Megachilidae species among the 26 present-day provinces than among the six historical provinces.

**Table 3.** Wild bee community differentiation based on 6,490 specimens relevant to 196 wild bee species grouped in 18 genera and seven phylogenetic groups within the family Megachilidae curated at RMCA (Tervuren, Belgium) and collected between 1905 and 1978. Community changes were assessed through beta diversity analyses and their partitioning among three categories of administrative or natural units: (i) the six historical provinces in DRC, (ii) the 26 present-day provinces, and (iii) the 10 known phytogeographical districts defined within the country. The metric βsør represents a measure of total dissimilarity, and its partitioning into its two major components: (1) species replacement among sampling units, i.e., turnover: βsim; and (2) species loss/gain, i.e., nestedness: βsne; following the formula βsør = βsim + βsne (Baselga 2010, Legendre 2014).

| | $\beta_{sor}$ | | $\beta_{sim}$ | | $\beta_{sne}$ | |
| --- | --- | --- | --- | --- | --- | --- |
|  | Taxonomic | Phylogenetic | Taxonomic | Phylogenetic | Taxonomic | Phylogenetic |
| Historical provinces (up to 1960) | $0.72 \pm 0.0$ | $0.56 \pm 4.3$ | $0.21 \pm 0.0$ | $0.05 \pm 0.4$ | $0.50 \pm 0.0$ | $0.51 \pm 3.9$ |
| Present-day provinces | $0.94 \pm 6.9$ | $0.74 \pm 4987.8$ | $0.52 \pm 3.8$ | $0.32 \pm 2180.0$ | $0.41 \pm 3.0$ | $0.41 \pm 2807.7$ |
| Phytogeographical districts | $0.81 \pm 0.0$ | $0.64 \pm 15.0$ | $0.23 \pm 0.0$ | $0.06 \pm 1.6$ | $0.57 \pm 0.0$ | $0.57 \pm 13.4$ |

A detailed examination of the occurrence records indicates that only the present-day provinces of Haut-Katanga, Kongo Central and Lualaba harbour bees from all seven taxonomics groups described above. The provinces of Kasaï-Oriental and Kwango are each home to species belonging to two groups within the Megachilidae, the province of Nord-Ubangi has three groups, while the remaining others (Kasaï, Kinshasa, Kwilu, Maï-Ndombe, Maniema, Tanganyika, Tshopo, Tshuapa, Ituri, Sankuru, Bas-Uele, Haut-Uele, Sud-Kivu, Nord-Kivu, Haut-Lomami, Kasaï-Central, Mongala, and Lomami) have species belonging to either five or six groups.

### Taxonomic and temporal patterns of Megachilidae specimen records

The analysis of the specimen records reveals that the most frequently collected species from DRC and curated at RMCA (Tervuren, Belgium) include the resin-dauber bee *Gronoceras cinctum* (Fabricius, 1781) (n=1,270 specimens), followed (in much lower numbers) by its specific cleptoparasitic bee *Euaspis abdominalis* (Fabricius, 1793) (n=394 specimens), and then by several species in the genus *Megachile sensu lato* such as *M. rufipes* (Fabricius, 1781) (n=390 specimens) and *Megachile bituberculata* Ritsema, 1880 (n=281) (Figure 2). Collectively, the four species mentioned above encompass 2,335 specimens and make up 36.0% of the dataset. At the other end of the abundance spectrum, we found 26 species that were represented in the RMCA collections only as type specimens (holotype and paratype).

Our specimen accumulation curves show several interesting patterns, such as the fact that 99.1% of the Megachilidae specimens currently curated at RMCA were collected strictly between 1905 and 1960 (the year the country gained political and administrative Independence on June 30, 1960), with a peak of specimen collection during the decade of 1930-40, particularly in 1932 when 1,188 specimens (belonging to 198 species in 18 genera) were collected by various entomologists (see below). The *Gronoceras* specimens are still represented in the collection up to 1978, whereas the last specimens collected in other groups are much older, such as the Lithurgini (last caught in 1955) and the *Noteriades* species (last caught in 1946).

Furthermore, the accumulation curves illustrate that the *Megachile sensu lato* species have been regularly collected and in higher numbers during the period 1905-1920; the sharp-tailed bees (cleptoparasites in the genus *Coelioxys*), the Osmiini species and the *Noteriades* (three species represented out of 11 or more known in Africa (Eardley *et al*. 2010)) have received comparatively less attention during these periods and are therefore less well represented in the RMCA specimen collection.

### The entomologists behind the historical Megachilidae specimen records at RMCA

Collectively, we found that the names of about 286 entomologists appeared on collector labels (legit) associated with Megachilidae specimens in the RMCA collection. Overall, and similar to the situation observed for the species (Figure 4), we found that some names appeared with a greater frequency than others. Of these, as shown by our rank abundance plot (Figure 5), ten collectors accounted for nearly 60 % of the specimens preserved. We found that Mr. Hans Joseph Brédo was the most frequent name, appearing on the labels of 1,057 specimens; this is almost double the number of specimens attributed to the following entomologists such as Mr. Anatole Corbisier (ranked second with n=536 specimens) and Mr. Frans Guillaume Overlaet (ranked third with n=530 specimens).

**Figure 3.**
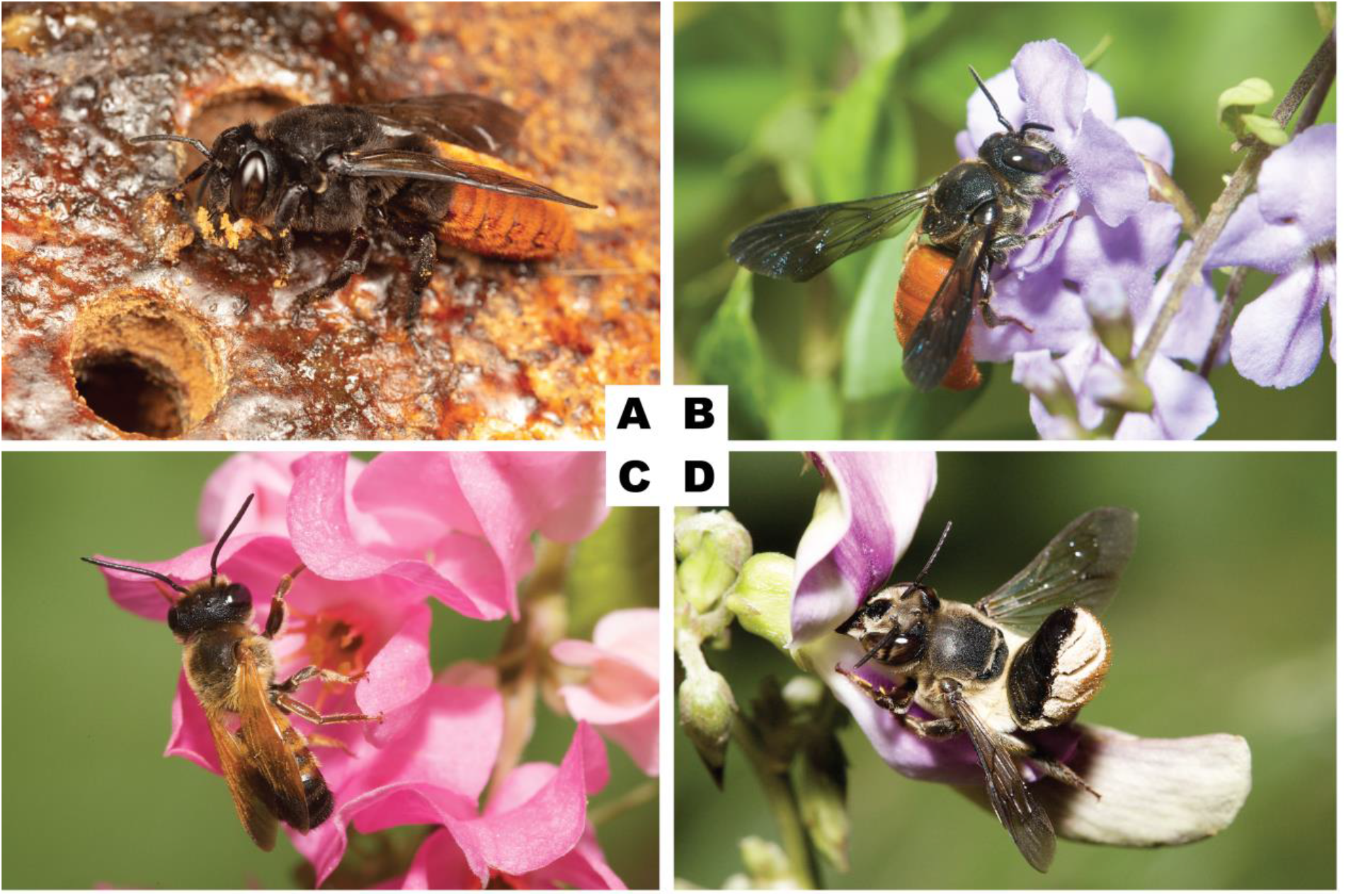
Top 4 of the Megachilidae species represented by the highest number of specimens in the RMCA collection (Tervuren, Belgium). **A.** *Gronoceras cinctum* (Fabricius, 1781) (female at nest entrance) (n=1,270 specimens); **B.** *Euaspis abdominalis* (Fabricius, 1793) (female on *Duranta erecta* (Verbanaceae)) (n=394 specimens); **C.** *Megachile rufipes* (Fabricius, 1781) (male on *Antigonon leptopus* (Polygonaceae)) (n=390 specimens); **D.** *Megachile bituberculata* Ritsema, 1880 (female on *Pueraria javanica* (Fabaceae)) (n=281). All photographs by NJ Vereecken.

**Figure 4.**
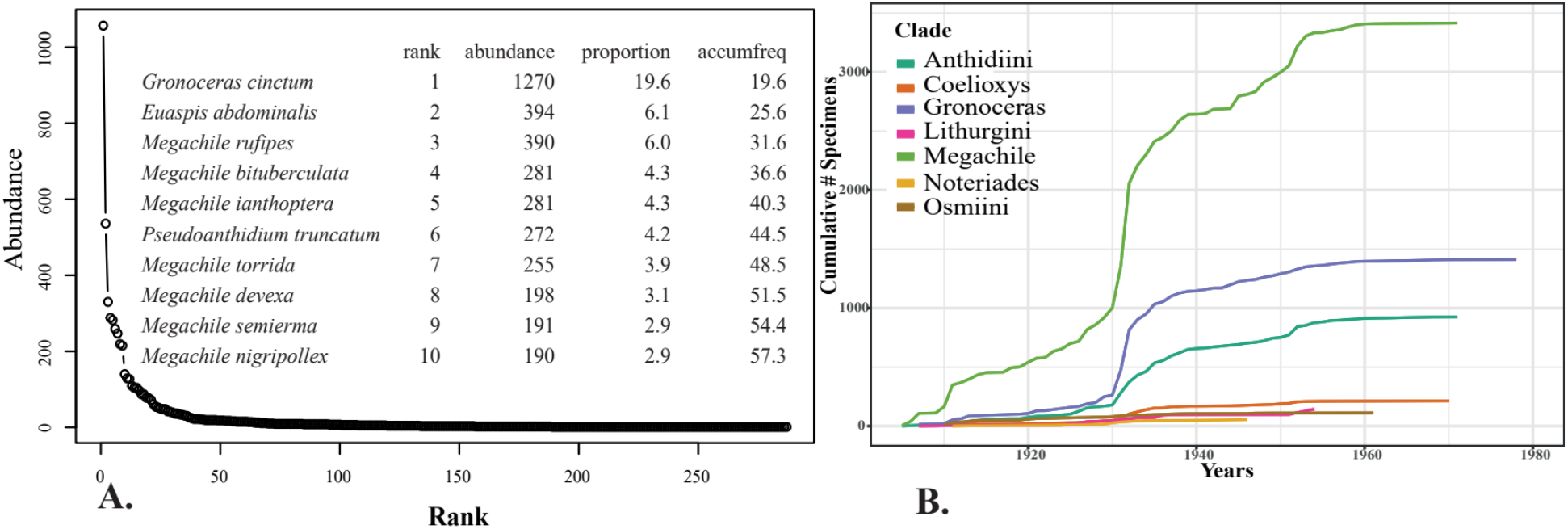
Taxonomic and temporal patterns of Megachilidae specimen records in the RMCA collection (Tervuren, Belgium). **A.** Rank abundance plot and associated data for the 10 most frequently collected species showing that the top 8 species make up to 50% of the dataset. The distribution of species along the curve shows a very long “tail”, implying that many of the species curated are only represented by a low or a very low number of species. For example, 26 species of Megachilidae were represented in the RMCA collections only as type specimens (holotype and paratype); **B.** Specimen accumulation curve through time (1905-1978) for the seven groups of species in the family Megachilidae showing that the best represented groups are the *Megachile sensu lato*, the *Gronoceras*, the Lithurgini and the Anthidiini species (by decreasing order of magnitude). It is noteworthy that the *Megachile* sensu lato have been regularly collected and in higher numbers during the period 1905-1920, and that all the above groups have received more attention during the decade 1930-40 compared to the other groups of Megachilidae.

**Figure 5.**
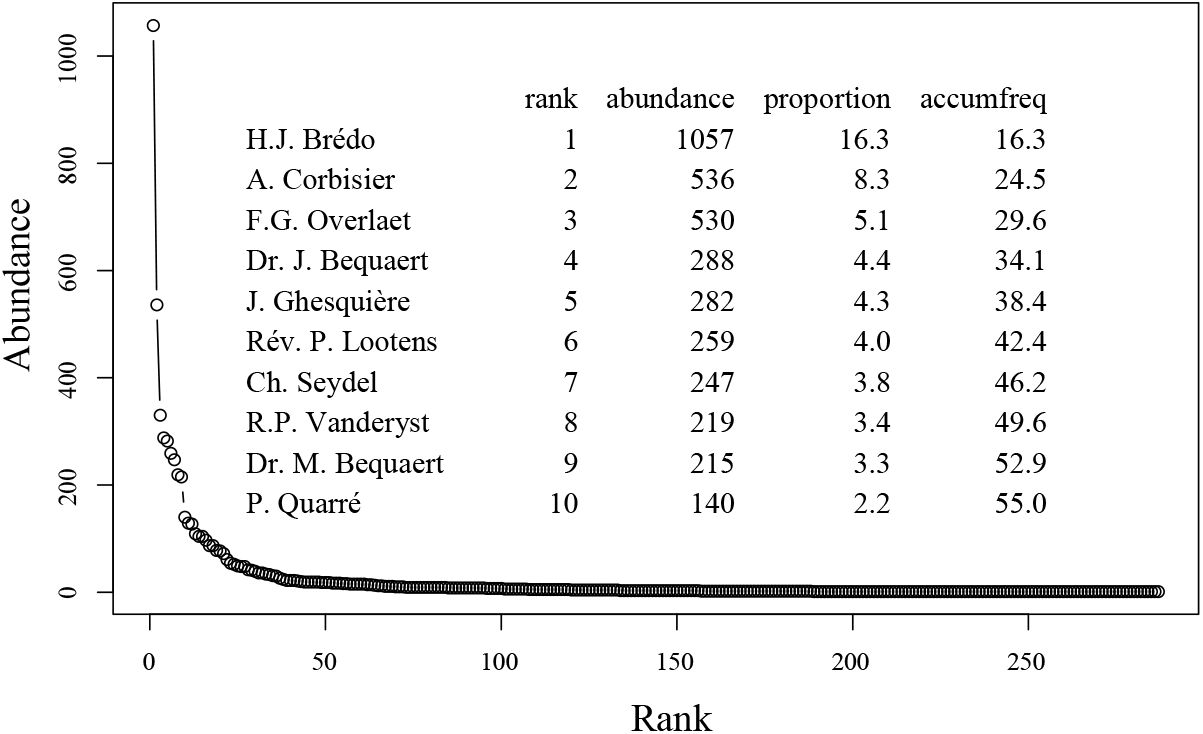
The entomologists behind the collection of Megachilidae specimen records at RMCA (Tervuren, Belgium).

A professional entomologist by training, Mr. Hans Joseph Brédo (1903-1991) was Belgian and studied mainly at the University of Louvain in Belgium. He later travelled Africa and DRC in particular as entomologist for the Belgian government where he worked on locusts, primarily on the environmental drivers of their swarming behaviour, and their control for crop protection. He contributed several papers during his time in DRC, *La lutte biologique et son importance économique au Congo Belge* (Brédo 1934), and he managed to collect most specimens of bees during his multiple field surveys across the provinces of DRC, in addition to locusts, other insects and thousands of herbaria on the flora of the 11 sub-Saharan countries (Harroy 1985).

Anatole Corbisier (Corbisier Baland, Anatole Antoine Pierre-Joseph-Ghislain) (1881-1950) was an agronomist and arboriculturist hired by the Belgian colony to supervise crops in Elisabethville (now Lubumbashi), where he transformed this chief town of the Katanga province into a city with beautiful streets and well laid-out parks. He was also appointed as director to the Botanical Garden of Eala, in the area of Coquilhatville (now Mbandaka), capital city of the Équateur Province along the Congo river, from 1916 until the end of his career in 1933. In addition to the numerous herbaria of species collected, he also provided important collections of insects from these regions of the Congo (Staner 1972). The Botanical Garden of Eala was run in turn by Belgians until Congo’s independence and managed by the INERA (Institut National pour l’Etude et les Recherches Agronomiques) until it became part of the Congolese Institute for Nature Conservation (ICCN) in 2010.

While very little information is available on the Belgian missionary Frans Guillaume Overlaet (1887-1956), we found that other catholic missionaries were also actively involved in the collection of Megachilid bees. Such is the case for Reverend Father Germain Lootens (1910-1976) (n=259 specimens, ranked sixth in Figure 5) and also for Reverend Father Hyacinth Julien Robert Vanderyst (1860-1934) (n=219 specimens, ranked eighth in Figure 5), the latter being described as an anthropologist, agronomist, botanist and naturalist, and who reported more than 3 % of Megachilidae specimens kept at the RMCA. Interestingly, we owe the earliest specimens of Megachilidae from DRC from as early as in 1905 in DRC to the Reverend Father Van Eyen and other anonymous entomologists; some of these voucher specimens were then sent to the French entomologist Joseph Vachal (1838-1911) by Dr. Henri Schouteden (1881-1972) who had been hired in the Natural Sciences section of the then-new Museum of Belgian Congo in 1910. These specimens described by Vachal (1910) also included series of wild bees collected by the British naturalist and entomologist Dr. Sheffield Airey Neave (1879-1961) during expeditions to Rhodesia and the Congo Free State between 1904 and 1913 (e.g., Neave 1907, 1910) where Dr. Neave was working on ticks, sleeping sickness and insect-borne diseases (Anonymous 1962; Baker & Bayliss 2009; Gilbert 2010). Dr. Neave is another classic example of an entomologist working on insect pests and disease vectors, similar to Dr. Joseph Charles Bequaert (1886-1982), a Belgian-born American naturalist who spent seven years (1909-1916) in DRC, first as entomologist working on the Belgian Sleeping Sickness Commission and other topics (e.g., Bequaert 1913, see also Pauly 2001). He was later head of botanical explorations in the Congo for the Belgian Colonial Government (Carpenter 1982). His brother Michel Bequaert also collected specimens in Elisabethville (now Lubumbashi) and at other localities in the former Katanga province (now Haut-Katanga province).

Last, it is important to note that only Theodore D. Cockerell (Cockerell 1932; 1933a, 1933b, 1933c, 1935a, 1935b, 1935c, 1938) has both travelled to DRC and described new species based on the collected specimens belonging to the family Megachilidae. Another important character in the study of DRC Megachilidae was the late Jean Jules Pasteels (1906-1991), Belgian Professor at the Université libre de Bruxelles (ULB) where he led research on embryology and entomology. Pasteels is also the authority behind the description of several Megachilidae species curated in the RMCA collection, and a pioneer in the taxonomy and systematics of sub-Saharan bees and other insects which he studied by examining the rich collections at RMCA, including material collected during the renowned “Mission G. F. de Witte (1946-1949) - Exploration of the National Park Upemba”.

The Table 4 below summarises the rate of specific variability of Megachilidae for the major collectors.

**Table 4.**
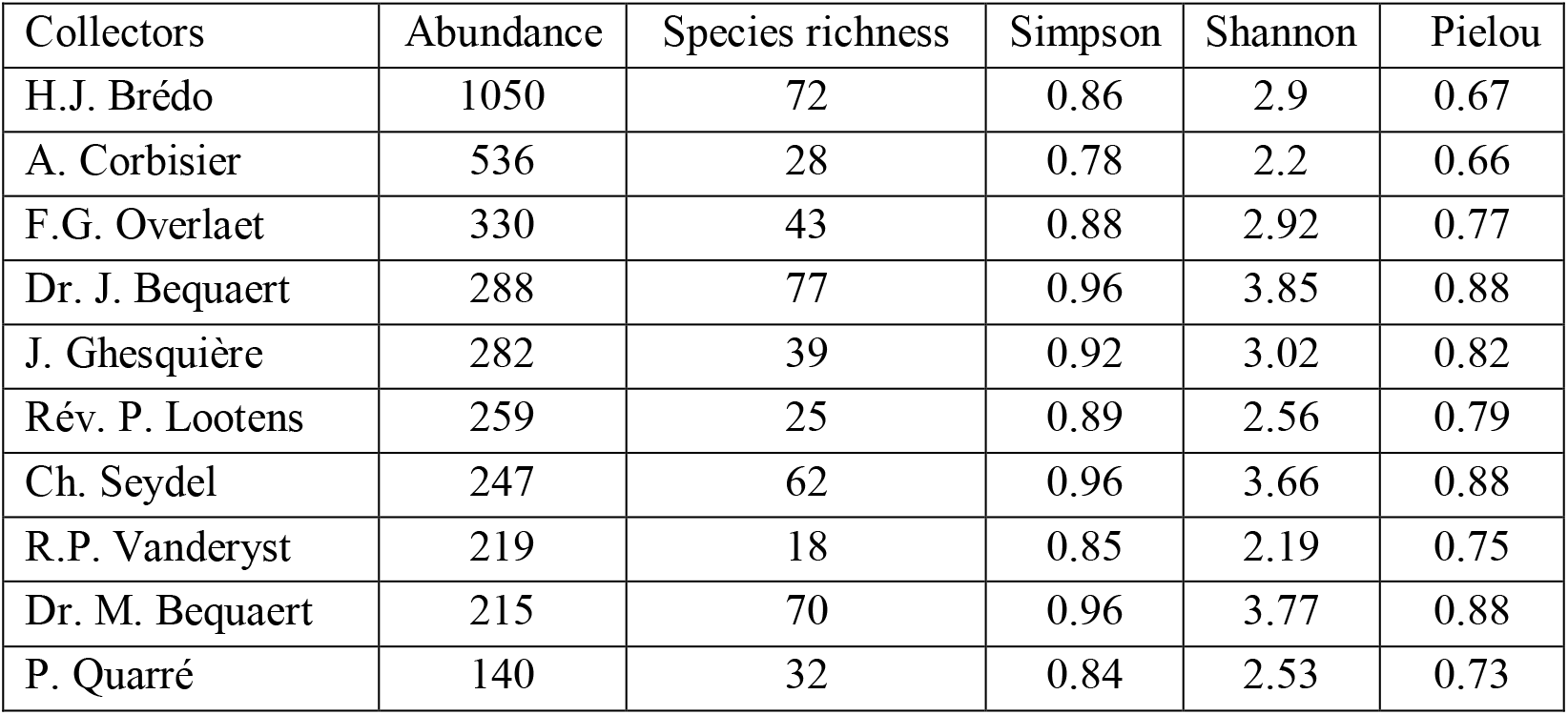
Diversity indicies of bee collecting by major entomologists. Some entomologists collected a relatively smaller number of species per specimens than others, reflecting low relative species diversity. For example, Mr. Hans Joseph Bredo (n = 1,050) collected the greatest number of sepcimens, yet a smaller number of species than did Dr. Joseph Charles Bequaert (ranked fourth in terms of abundance), deliberately recording the highest species richness on his 288 specimens, so great species diversity.

Although having collected the largest number of specimens, Hans Joseph Bredo’s collections are mainly dominated by the species *Gronoceras cinctum* (Fabricius, 1781), which accounts for 32.4% and secondarily *Euaspis abdominalis* (Fabricius, 1793) accounting for 9.5% of his 1,057 specimens, which justifies a poor distribution of relative specific abundances. Anatole Corbisier shows a lower diversity of collection with also a dominance of *Gronoceras cinctum* (Fabricius, 1781) (41%) on the 28 species sampled. In contrast to the best collector, Joseph Bequaert reported a greater variety of species with evenly distributed abundances, hence the high species diversity associated with his 288 specimens.

A gallery of portraits of the abovementioned entomologists is shown on Figure 6.

**Figure 6.**
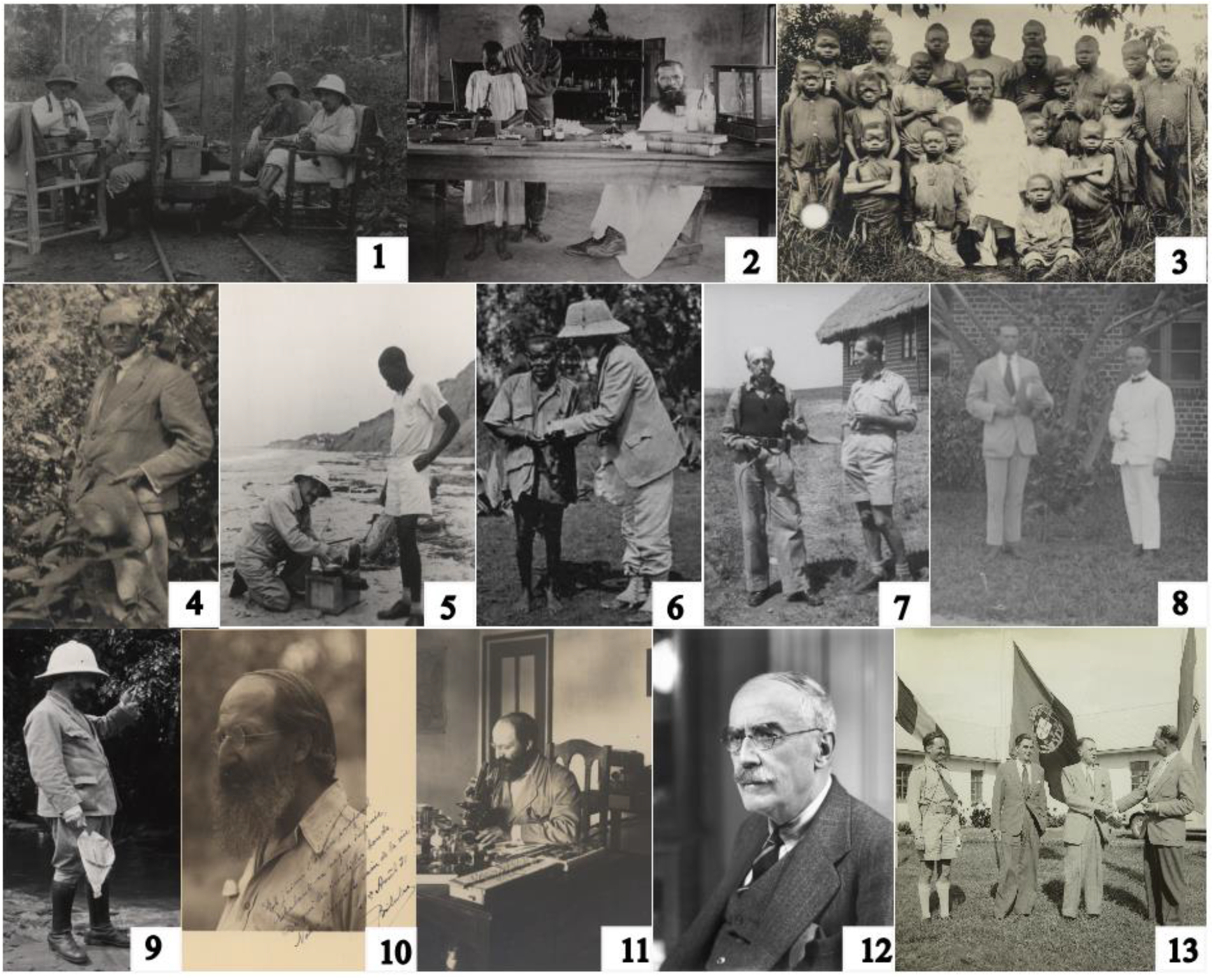
The diversity of entomologists behind the collection of Megachilidae specimen records at RMCA (Tervuren, Belgium). We list here the inventory number corresponding to each photograph, as well as the names of the photographers, for the images kindly provided by the RMCA archives department. **1.** Mrs. Lebrun and Vrydagh (*left*) and Mrs. Corbisier and Staner (*right*) (AP.0.0.29742; photo by P. Staner 1930); **2.** Reverend Father Hyacinthe Vanderyst in Kisantu (AP.0.0.23368; photo by “Jesuit Order”, 1910-1925); **3.** Reverend Father Hyacinthe Vanderyst (R.P. Vanderyst) in Ipamu (AP.0.2.368; photo by R.P. Delaere, 1922); **4.** Frans Guillaume Overlaet (AP.0.0.25895; photo G. Fr. de Witte, 1931); **5.** Maurice Bequaert (HP.1950.7.37, photographer not identified, 1950); **6.** Professor Theodore D. Cockerell and the pygmies chief Kasulo, who lived in the forest near Tshibinda in the Sud-Kivu Province (photo by Alice Mackie); **7.** Gaston-François De Witte (*left*) and Hans Joseph Brédo (*right*) (HP.2011.62.20-221; photo G. Fr. De Witte, 1947); **8.** Prince Léopold III of Belgium (*left*) and Mr Ghesquière (*right*) in front of the laboratory in Stanleyville (now Kisangani) (AP.0.0.25895; photo Ghesquière, 1925); **9**. Charles Seydel *alias* “Bwana Bilulu” (HP.1953.21.8, photo E. Devroey, 1922); **10.** Charles Seydel *alias* “Bwana Bilulu” (HP.2011.62.13-226, photo Gaston-François De Witte, 1931) ; **11.** Charles Seydel *alias* “Bwana Bilulu” (HP. 2011.62.5-145; photo Gaston-François De Witte, 1925) ; **12.** Sheffield Airey Neave; **13.** Hans Joseph Brédo on the right, Director of the International Locust Laboratory in Abercorn (Northern Rhodesia), welcomes Belgian personalities on an official visit to the Laboratory (2017.24.30; photo Infocongo, 1952). All rights reserved for photographs **1**, **2**, **3**, **5**, **8** and **9**; photographs **4**, **7**, **10**, **11** and **13** under Creative Commons license (CC-BY 4.0); photograph **6** after Cockerell (1932) and **12** ©National Portrait Gallery.

## Discussion

### Wild bees in the family Megachilidae in DRC

All these collections of Megachilidae from the RMCA were identified by Jean Jules Pasteels and the species described in his publications. However, this author did not place much importance on citing these data, which most of the time he mentioned only the country, “all of Africa”, or even one of its parts. Other researchers have repeated Pasteels’ work without citing his data in detail. For example, Liongo li Enkulu (1988) limited himself to producing maps for the ancient Chalicodoma group. Far from being a personal contribution to the identification of new material, the present work is simply a publication of the data identified by Pasteels and verified in part by Liongo li Enkulu, to which are added patterns of diversity by tracking their distribution in the Congolese space. It supports 6,490 specimens representing 195 species, constituting a valuable natural history legacy from the colonial era of the DRC and represents the largest dataset on wild bees in the country. Indeed, only a small number and partial occurrence records of bees from DRC have been digitized and made available to date. For example, at the time of adding the last edits to this manuscript, no specimen record of any Megachilidae species from DRC was available on the iNaturalist platform (https://www.inaturalist.org); only 443 confirmed specimen records associated with both preserved specimens and coordinates, primarily from institutions and museums in the USA, were available on GBIF (https://www.gbif.org). Of these, 286 are Megachilidae specimens (i.e., 64.56%) curated at the American Museum of Natural History (AMNH, New York, USA), unfortunately lacking the collection month and year. Another 104 specimen records of Megachilidae from DRC (i.e., 23.47%) have been shared on GBIF by curators of the Snow Entomological Museum Collection (SEMC) at the University of Kansas Biodiversity Institute (Kansas, USA), 34 specimen records (i.e., 7.67%) by the USDA-ARS Pollinating Insects Research Unit of the United States Geological Survey (USGS) in Logan (Utah, USA). The only African institution contributing to this publicly available dataset is the Iziko South African Museum in Cape Town (South Africa) which has shared the details relevant to 10 specimen records (i.e., 2.25%). A comprehensive synthesis of the Megachilidae bees of DRC was out of the scope of the present study, as it would require digitizing all congolese bee specimens curated in other international depositories among those listed by Coetzer & Eardley (2019). However, the figures presented above show that because of the historical colonial context of DRC and Belgium, and because virtually no other field surveys that we are aware of have been published before or ever since the Independence of DRC in 1960, we reckon that we have presumably covered the vast majority of all specimen records ever collected for wild bees in the family Megachilidae in DRC through the present work.

The rich collections from the Equateur Province were obtained through surveys at the Botanical Garden of Eala, the headquarters that coordinated most biological expeditions until the end of the DRC’s colonial era (e.g., Harroy 1985). To date, the catalogue of Coetzer & Eardley (2019) lists only 72 species of Megachilidae known from Katanga and 93 reported at the national level in DRC. In our catalogue, we recorded up to 147 different Megachilidae species from the Katanga Province alone (Table 1). The Katanga province has arguably been one of the most scientifically explored regions, including in other entomological groups (Vachal 1910, Cockerell 1937), but it shows that this province in particular is home to a rich fauna of Megachilidae bees. While the catalogue of Coetzer & Eardley (2019) listed 93 species of Megachilidae reported from the whole of the DRC, the present study now brings this figure up by another 102 additional species, reaching 195 species in total for the country. It should be noted here that the existence of another three species (*Coelioxys ateneata* Strand, 1920; *Coelioxys neavei* Vachal, 1910; *Coelioxys umbripennis* Friese, 1922) described and supposedly curated at the RMCA according to historical publications (Strand 1921; Vachal 1910a; Cockerell 1935b), but which were not found there, raises the species richness to 198 species of Megachilidae in DRC.

Different sampling intensities can be observed depending on the administrative/natural unit considered. The most explored provinces, such as the current Kongo Central, also exhibits significantly higher levels of Megachilidae species richness. The species sampling rate (Figure 1) seems to have been driven by the detectability of species (many of which are large and conspicuous), as well as by the accessibility of phytogeographic areas and the historicity of localities within provinces (Monsarrat *et al*. 2019; Wang *et al*. 2022). Some taxa were associated with less records, such as the smaller bees of the genus *Noteriades* or the subgenus *Megachile* (*Eutricharaea*), have been comparatively less frequently sampled by the entomologists during the colonial period. Likewise, the comparatively less accessibility of the Central Forestry District (VI) (Figure 1), which is dominated by tropical rainforest and therefore quite difficult to survey, is likely to explain the lower number of specimens and species of Megachilidae sampled in this region. The historicity of localities refers to the succession of administrative and scientific activities over time. For example, the locality of Eala (Province of Equateur) is better represented in terms of abundance of collections because of the location of the botanical garden originally called Bokoto and then inaugurated as the Eala Botanical Garden on February 2, 1900, founded by Emile Laurent of the Faculté Universitaire des Sciences Agronomiques de Gembloux (Belgium) who was then employed by what was still the Congo Free State of King Leopold II of Belgium. This botanical garden was once considered one of the most important tropical gardens of the world, a major scientific hub for agricultural research on exotic species of plants and trees with economic potential introduced to Africa, and key headquarters from which many scientists went on expeditions across DRC during the colonial era (Harroy 1885).

Finally, the spatial scale of sampling (Rahbek 2005; Wang *et al*. 2022) is important in the distribution of abundances and species richness of Megachilidae. Among the cells (50 km x 50 km) within phytogeographic regions, some appear to be more explored than others. This is the case in the locality of Eala (Mbadaka) which has got almost all the specimens reported from the Equateur Province, and almost half of the specimens collected in the vast “Central Forestry” phytogeographical district (VI). Lubumbashi also dominates the abundance of collections in the Upper Katanga phytogeographical district (X). By contrast, the provinces Haut-Katanga, Lualaba, and Kongo Central comprise species from all taxonomic groups as described above, and at least one representative of the 7 taxonomic groups of Megachilidae, thus appear to be more sampled than the rest. Generally speaking, the neighbouring provinces are compositionally different, except for the two provinces Kwango and Kwilu that exhibit more similar in their Megachilidae fauna and which are found within the same phytogeographical districts.

### Diversity and endemism of wild bees in DRC

Many species described from the Congo are not currently reported from other countries in Sub-Saharan Africa (Liongo li Enkulu 1988; Eardley & Urban, 2010). The works of Cockerell (1932; 1933a, 1933b, 1933c, 1935a, 1935b, 1935c, 1938), Pasteels (1965, 1966, 1968), as well as those of Eardley and colleagues (Eardley & Urban, 2010; Eardley & Griswold, 2015; 2016; 2017) have highlighted the taxonomic originality and the high endemism of the DRC wild bee fauna. However, we have reservations about the endemism of the DRC, given that it appears relatively more prospected than other Central African countries (e.g. Gabon, Central African Republic, Republic of Congo), but also the fact that these specimens were mostly identified by J.J. Pasteels. Within the bee family Megachilidae, putative DRC endemic species include the cleptoparasitic species *Euaspis rufiventris* subsp. *uvirensis* Cockerell, 1933 and *Coelioxys* (*Allocoelioxys*) *congoensis* Friese, 1922, as well as the Anthidiine *Pachyanthidium* (*Pachyanthidium*) *katangense* Cockerell, 1930 and two *Megachile* species, namely *M.* (*Eurymella*) *kimilolana* Cockerell, 1931, and *M.* (*Pseudomegachile*) *bukamensis* Cockerell, 1935. According to Pauly (2015), a series of 13 species of *Megachile sensu lato* seem to exhibit a distribution restricted to the Congo Basin and DRC in particular, with no records outside the country’s present-day administrative boundaries. Of these putative endemic species, eight belong to *Megachile sensu stricto*, including seven that are recorded in our catalogue (*Megachile akamiella, M. brochidens, M. maculosella, michaelis, M. niveicauda* and *M. paupera*) as well as *M. pinguicula* Pasteels, 1965 (Pasteels, 1965). Another three species displaying a similar geographic distribution belong to the group of dauber bees, treated by by Pasteels (1965) as the genus *Chalicodoma*, namely *M. ambigua* (Eardley & Urban 2010), *M. biloba* and *M. biseta*. Similarly, we found a potential endemic species from the genus *Gronoceras* (*Gronoceras chapini*) historically designed as *Chalicodoma* (*Gronoceras*) *chapini* Pasteels, 1965, as well as another species from the subgenus *Creightonella*, namely *Megachile* (*Creightonella*) *alternans* Friese, 1922, which was not represented in the RMCA collections. Overall, we support the suggestions made by previous authors on the potentially high rate of endemism among the Conglese bees as reflected by the current state of knowledge, but more bee surveys in neighbouring countries should be conducted to test the extent of this DRC endemism across bee families (Ferreira de Lima *et al*. 2020; Santos & Ribeiro 2022; Shipley & McGuire 2022).

At the scale of DRC, habitat specialist bees might also be encountered, including for Megachilidae species such as *M.* (*Eurymella*) *kimilolana* Cockerell, 1931 that appears to be intimately linked to the savannah-forest mosaic of the lower (V) and upper Katanga (X) phytogeographical districts. Unlike the central basin of the Congo, which is covered by dense forests, the former province of Katanga has a savannah whose fauna would probably be found in neighboring countries such as Angola, Zambia and Tanzania (Eardley & Urban 2010; Ascher & Pickering 2020). Future studies should explore ecological affinities of wild bees in DRC, including their (micro-)habitat preferences, but also their biotic interactions, from host plants to cuckoo-host species pairs, which remain poorly known and documented to the present day (Kuhlmann, 2009; Gous *et al*. 2021).

### Contemporary research on wild bees in DRC: where to start?

The production of the present catalogue on the large family Megachilidae from Congo raises several challenges. Among the most important, it seems to us a priority to consider a contemporary re-sampling of historically well-explored localities and biodiverse habitats such as the Eala Botanical Garden near Mbandaka (Province Orientale), as well as the area in and around the Kisantu Botanical Garden (Kongo Central Province). This research activity is able to facilitate the state of knowledge of local diversity for possible disappearances or discoveries of new species. In the same vein, the Garamba and Upemba national parks are of great interest for contemporary bee exploration. These protected areas, as well as the mountains bordering Lake Tanganyika, are home to lesser-known savannah fauna that should also be explored. Another priority sampling area would be the former experimental station of Mulungu-Tshibinda on the SE shore of Lake Kivu in the eastern highlands, where an Afromontane botanical garden and agricultural research station was established by the Belgian government authorities during the colonial period. The Mulungu-Tshibinda station and its surroundings were successfully explored by Professor Theodore D. Cockerell, who described several endemic species from this locality (Cockerell 1932). Resampling of taxa can provide insights into possible biodiversity loss or other functional changes (Graham *et al*. 2021) in these well-sampled areas of DRC and could help collecting more biological material and generate DNA barcodes for species known only from their type specimens which are now already more than 60 years old. The almost inevitable degradation of type specimens would mean the loss of biological material essential for the development of follow-up taxonomic studies. This is particularly relevant when considering the various taxonomic challenges. For example, Pasteels (1965) reported, before his multiple revisions, at least 500 valid species names of Afrotropical Megachilidae, not counting synonymies. Contemporary taxonomic revisions of the Megachilidae as well as the other five bee families (Apidae, Colletidae, Halictidae, Melittidae), of DRC and Sub-Saharan Africa are much needed to provide the basis for future research on basic and applied aspects of the ecology and evolution of wild bees and their biotic interactions.

Secondly, research in this area should be directed towards the implementation of regular monitoring of known and unknown hotspots, both in the remaining pristine tropical forests and in areas subjected to anthropogenic land use change. Of particular interest should be regions of DRC that are part of the Eastern Afromontane biodiversity hotspot, one of the Earth’s 35 biodiversity hotspots, the most biologically rich yet threatened areas around the globe where access is reportedly compromised by the proliferation of armed groups and inter-ethnic conflicts (Dorsouma & Bouchard 2010; Pourtier 2009). As much as the knowledge of poorly known wild bee species is compromised by the ravages of war, some of them could disappear altogether with the unstoppable loss of key components of their Afromontane habitat. We therefore advocate here for a renewed interest and financial support for scientific research in such regions that are likely among the most biodiverse tropical hotspots of diversity for wild bees and other pollinators.

### Concluding remarks

This study is an important first step to establish a historical baseline on the diversity of wild bees in the family Megachilidae, and it contributes to the recent wave of museum decolonization efforts through the use of digital technologies to enable democratizing and repatriating important aspects of DRC’s natural heritage and potential downstream value (Berents *et al*. 2010; Beaman & Cellinese 2012; Ströbel *et al*. 2018; Hedrick *et al*. 2020; Msila 2021). NHCs contain a diversity of information beyond their conservation over time (McLean *et al*. 2016; Thomson *et al*. 2018). Given their inaccessibility to many scientists, their digitisation is essential to facilitate understanding of biodiversity as it evolves in the face of environmental and land use change (Miller & Rogo 2001; Drew *et al*. 2017; Hedrick *et al*. 2020). These efforts lay a solid foundation to stimulate a new wave of contemporary research to assess the present-day status of wild pollinators in DRC and neighbouring regions, and to explore the biodiversity-(agro) ecosystem productivity nexus as well as its impact on human welfare (Potts *et al*. 2016b; Nayak *et al*., 2021). Should the national and provincial authorities of DRC take these advances into account, logistical and financial allocations for scientific research would provide DRC researchers interested in wild bee diversity and functions with a significant head start in the Afrotropical region. Far from the vision of a proliferation of parachute science, whereby Northern scientists engage in Southern issues “out-of-the-blue” (Raja *et al*. 2022) the present study also exemplifies a fruitful and sustainable partnership (Akena 2012; Tobiasz. *et al*. 2019) among a diversity of co-authors from the North and the South and between research institutions already engaged through privileged academic partnerships (e.g., ULB and UNILU). We hope that this study will inspire other similar initiatives across Africa and beyond, and that this is also a first step towards a more in-depth study of the major bee groups in DRC, and the perspectives of gaining more fine-grained insights into the distribution, ecology, threats and trends of these essential pollinators.

## Authors’ contribution

The study was conceptualised by NJV. ATN digitized and analysed the data. NJV and ATN led the writing of the manuscript, with valuable inputs from AP and AD. All authors have read and approved the latest version of the manuscript.

## Acknowledgements

We are grateful to the RMCA staff, particularly Didier Van der Spiegel, Stéphane Hanot, Christophe Allard, Patricia Van Schuylenbergh and Anne Welschen for their availability, valuable advice, as well as for sharing photographs from the historical archives of insect collectors in DRC. We thank Stéphane De Greef (ULB) for providing logistic help with the initiation of the specimen records digitization, and Leon Marshall (ULB) for his help with the data formatting. We thank also Prof. Mylor Ngoy Shutcha (UNILU) for his support throughout the study. This study was supported by the ARES-CCD grant to ATN in the framework of his PhD thesis at ULB and UNILU on wild bees and crop pollination in the Katanga Province of DRC.

## Appendices State of knowledge on the endemism of bees within the family Megachilidae in the DRC

**Appendix 1.**
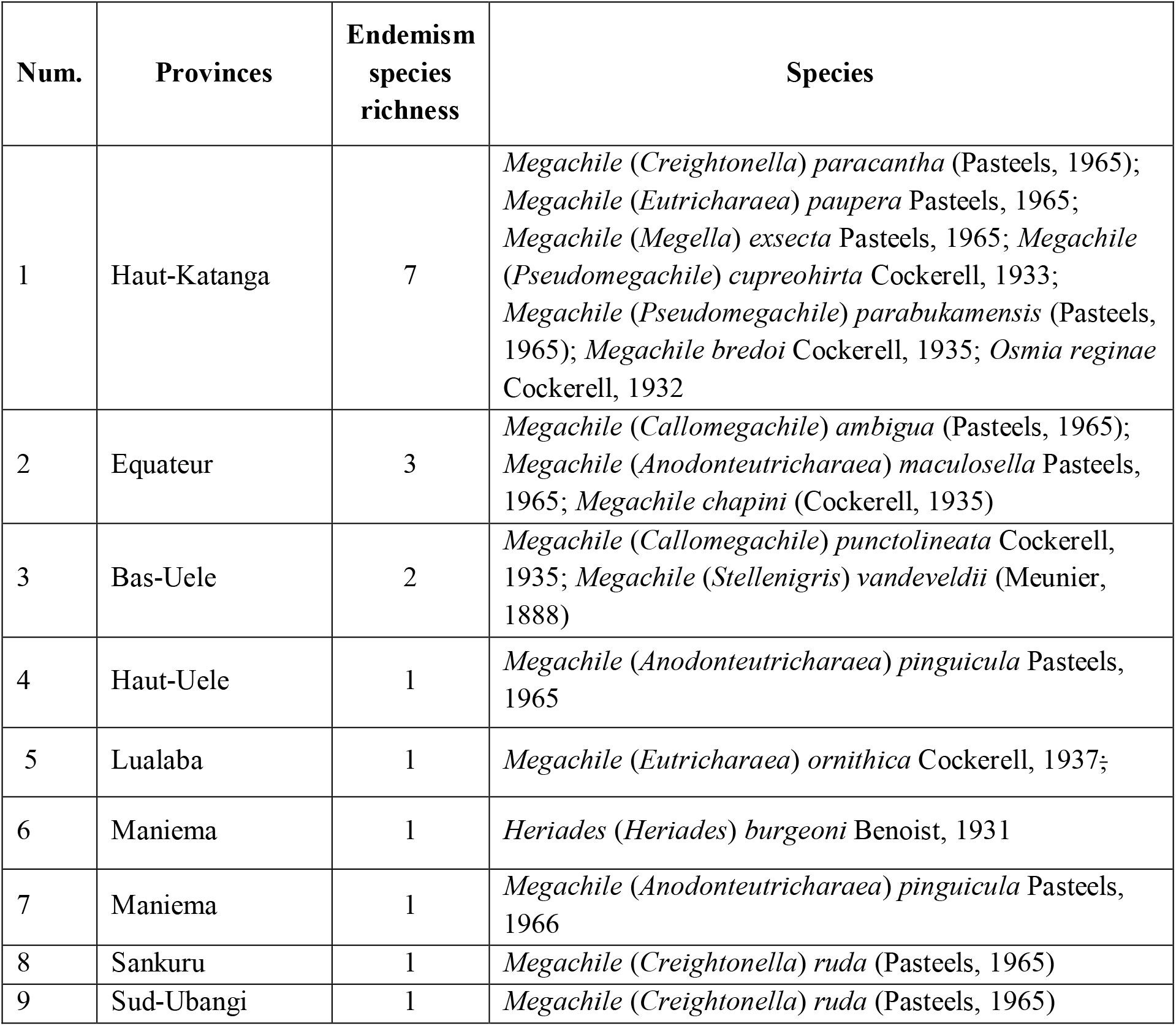
DRC bee endemism by province. The province of Haut-Katanga shows the highest level of endemism

**Appendix 2.**
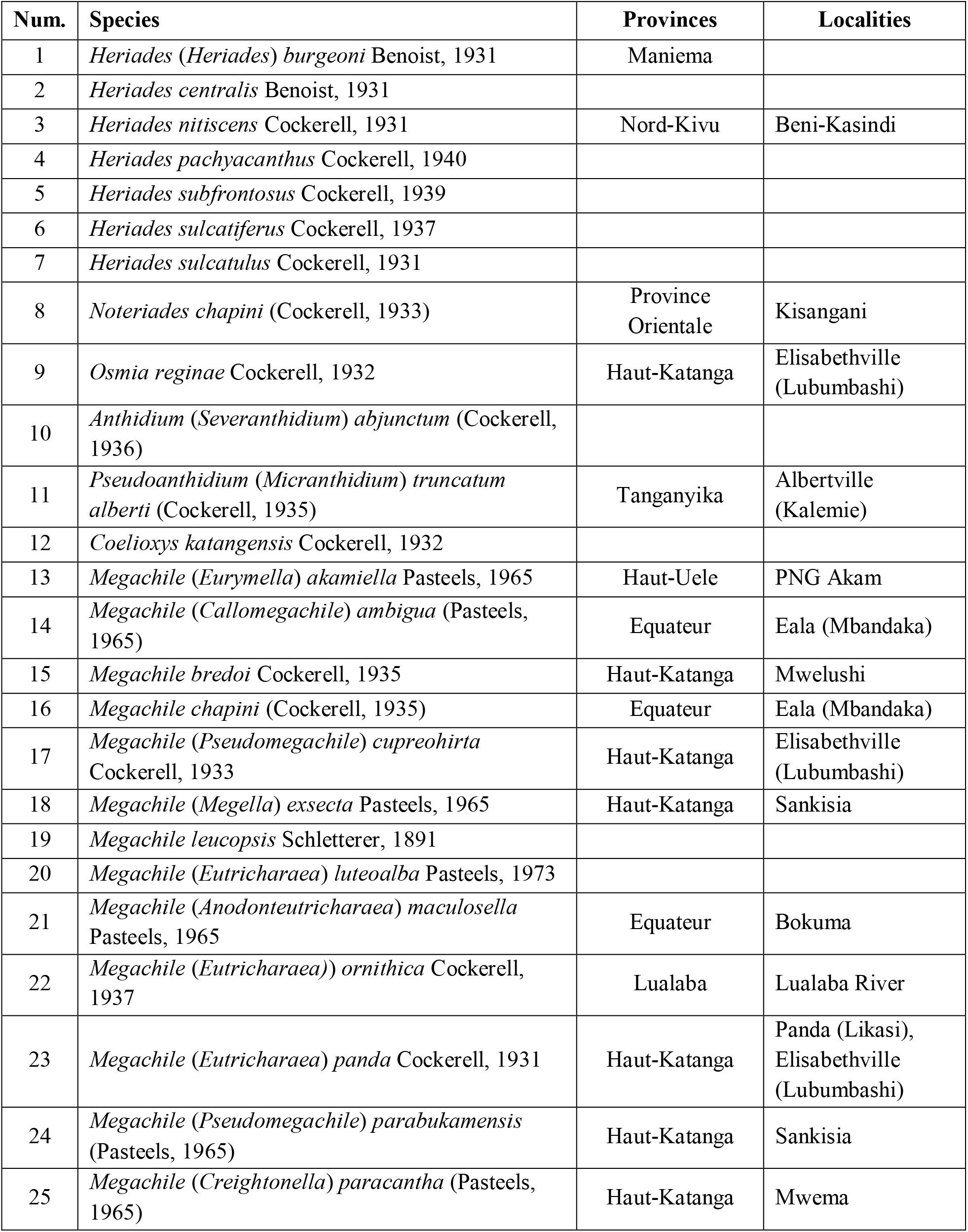
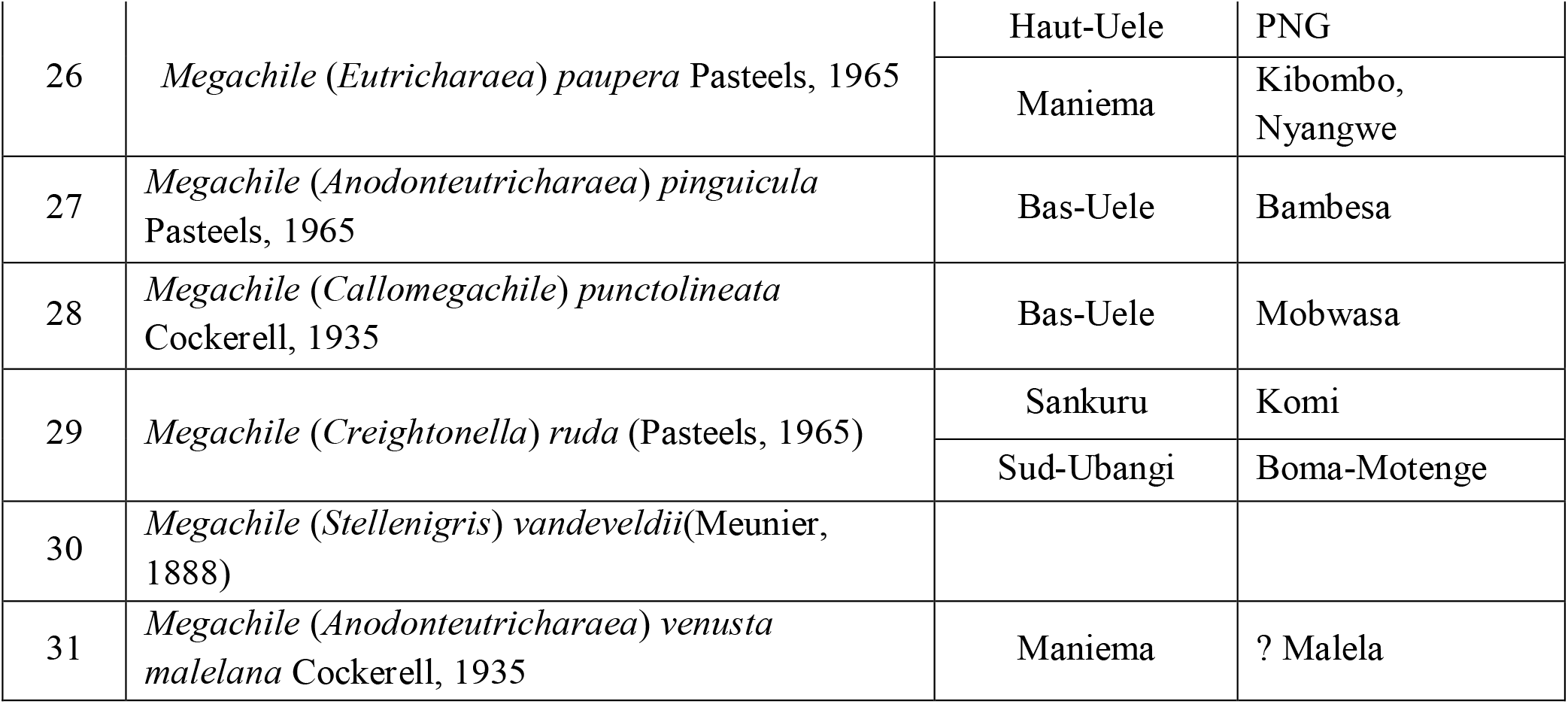
Endemism of DRC bees within the family Megachilidae following Provinces and localities. Some locations have not been specified since the data source.

## Footnotes

1 Holotype for Megachile fimbriata kiambensis Cockerell, 1933

2 This specimen is gynandromorphic

3 Published as Megachile vanderysti holotype in Cockerell, T.D.A. (1935) Bees of the genus Megachile in the Congo Museum. Revue de Zoologie et de Botanique Africaines, 26, 239–246.

4 Published as Megachile ruficauda holotype in Cockerell, T.D.A. (1935) Bees of the genus Megachile in the Congo Museum. Revue de Zoologie et de Botanique Africaines, 26, 239–246.

5 Described as Megachile torula Vachal, 1910

6 The attribution of this species to Smith as descriptor in the RMCA collections is erroneous.

7 Published as Anthidium glomerosum holotype in Vachal (1910). Diagnoses d’insectes nouveaux recueillis dans le Congo belge par le Dr. Sheffield-Neave (Musée du Congo, Tervueren), pp306-328.

8 Other specimen of Anthidium glomerosum Vachal, 1910.

9 Published as Anthidium neavei holotype in Vachal (1910). Diagnoses d’insectes nouveaux recueillis dans le Congo belge par le Dr. Sheffield-Neave (Musée du Congo, Tervueren), pp306-328.

10 Published as Anthidium neavei Vachal, 1910 paratype

11 This species is given as synonym of Heriades spiniscutis (Cameron, 1905) in Michener (1968).

12 Strand, E. (1921). Notes sur quelques Apides de Congo belge. Revue de Zoologie et de Botanique Africaines, 8, pp.104-106.

13 A junior synonym of Coelioxys congoensis Friese, 1922, in Cockerell, (1935b). African bees of the genus Coelioxys. Revue de Zoologie et de Botanique Africaines, 26, pp.443-444.

14 Vachal, (1910). Diagnose d’insectes nouveaux recueillis par le Dr. Sheffield-Neave: Hymenoptera, Apidae. Annales de la Société Entomologique de Belgique 54, pp.315-316.

15 A junior synonym of Coelioxys planidens Friese, 1904, etc. Vachal, (1910). Diagnose d’insectes nouveaux recueillis par le Dr. Sheffield-Neave: Hymenoptera, Apidae. Annales de la Société Entomologique de Belgique, 54, pp.316-317.

16 A junior synonym of Coelioxys planidens Friese, 1904, etc. Cockerell, T.D.A. (1935b). African bees of the genus Coelioxys, Revue de Zoologie et de Botanique Africaines, 26, p.442.

17 A junior synonym of Coelioxys umbripennis Friese, 1922, in Cockerell (1935b). African bees of the genus Coelioxys. Revue de Zoologie et de Botanique Africaines, 26, p.437.

18 A junior synonym of Megachile schulthessi Friese, 1903, in Vachal (1910). Diagnoses d’insectes nouveaux recuellis dans le Congo belge par le Dr. Sheffield-Neave: Hymenoptera, Apidae. Annales de la Société Entomologique de Belgique 54, p.308.

19 Another specimen was identified as Megachile flexa in Vachal (1910). Diagnoses d’insectes nouveaux recuellis dans le Congo belge par le Dr. Sheffield-Neave: Hymenoptera, Apidae. Annales de la Société Entomologique de Belgique 54, p.308.

20 A junior synonym of Megachile ungulata Smith, 1853, in Vachal (1910). Diagnoses d’insectes nouveaux recuellis dans le Congo belge par le Dr. Sheffield-Neave: Hymenoptera, Apidae. Annales de la Société Entomologique de Belgique 54, pp.311-312.

## Notes

### Competing Interest Statement

The authors have declared no competing interest.

